# Biosynthesis of bisbenzylisoquinoline alkaloids

**DOI:** 10.64898/2026.05.22.726869

**Authors:** Qishuang Li, Xinyi Li, Xiang Jiao, Guanghong Cui, Xiangmei Tan, Ying Ma, Yanduo Wang, Yaqiu Zhao, Jian Wang, Wenbo Xu, Tong Chen, Yating Hu, Ping Su, Yifeng Zhang, Jens Nielsen, Yun Chen, Juan Guo, Luqi Huang

## Abstract

Bisbenzylisoquinoline alkaloids (bisBIAs) are pharmacologically valuable plant metabolites with complex stereochemical architectures, yet the catalytic principles governing their assembly have remained largely unclear. Here, we elucidate the enzymatic pathway to cyclic bisBIAs and uncover a non-canonical redox-mediated mechanism for post-assembly stereochemical control. We identify cytochrome P450 enzymes that catalyze regioselective oxidative dimerization and macrocyclization of benzylisoquinoline monomers, establishing the macrocyclic scaffold. Subsequent stereochemical specification is achieved by a paired oxidase-reductase module that selectively epimerizes a single stereocenter through a transient imine formation, converting (*R*,*S*)-configured intermediates to (*S*,*S*)-products. Reconstitution of the pathway in yeast enabled production of both native bisBIAs and non-natural analogs, demonstrating pathway modularity and engineering potential. These results establish the biochemical principle underlying bisBIA biosynthesis and provide a framework for programmable biosynthesis of these complex natural products.

## Main Text

Benzylisoquinoline alkaloids (BIAs) constitute a chemically coherent group of plant specialized metabolites that includes widely used drugs such as morphine and noscapine^1^. Among more than 2,500 structurally distinct BIAs identified to date^2^, bisbenzylisoquinoline alkaloids (bisBIAs) represent a particularly elaborate subclass, generated through oxidative phenol coupling between two 1-benzylisoquinoline alkaloids (1-BIA) units (Fig. 1a)^3^. Members of this family have played prominent roles in medicine and pharmacology, ranging from early neuromuscular blockers such as tubocurarine to widely used therapeutic agents including cepharanthine and semisynthetic derivatives such as cisatracurium^4,5^. In addition, several bisBIAs have demonstrated clinical or preclinical utility, including tetrandrine, which inhibits Ebola virus replication by targeting lysosomal integral membrane protein-2 to disrupt sphingosine-mediated calcium signaling^6,7^; cepharanthine, which interferes with viral entry, positioning it as a promising therapeutic lead against SARS-CoV-2^8,9^; and methylated derivatives of daurisoline, which modulate GLP-1-mediated glucose metabolism^10^. It is thus evident that the bisBIA scaffold holds high promise for therapeutic applications by targeting diverse pathologies ranging from viral infections to metabolic disorders.

**Fig. 1.**
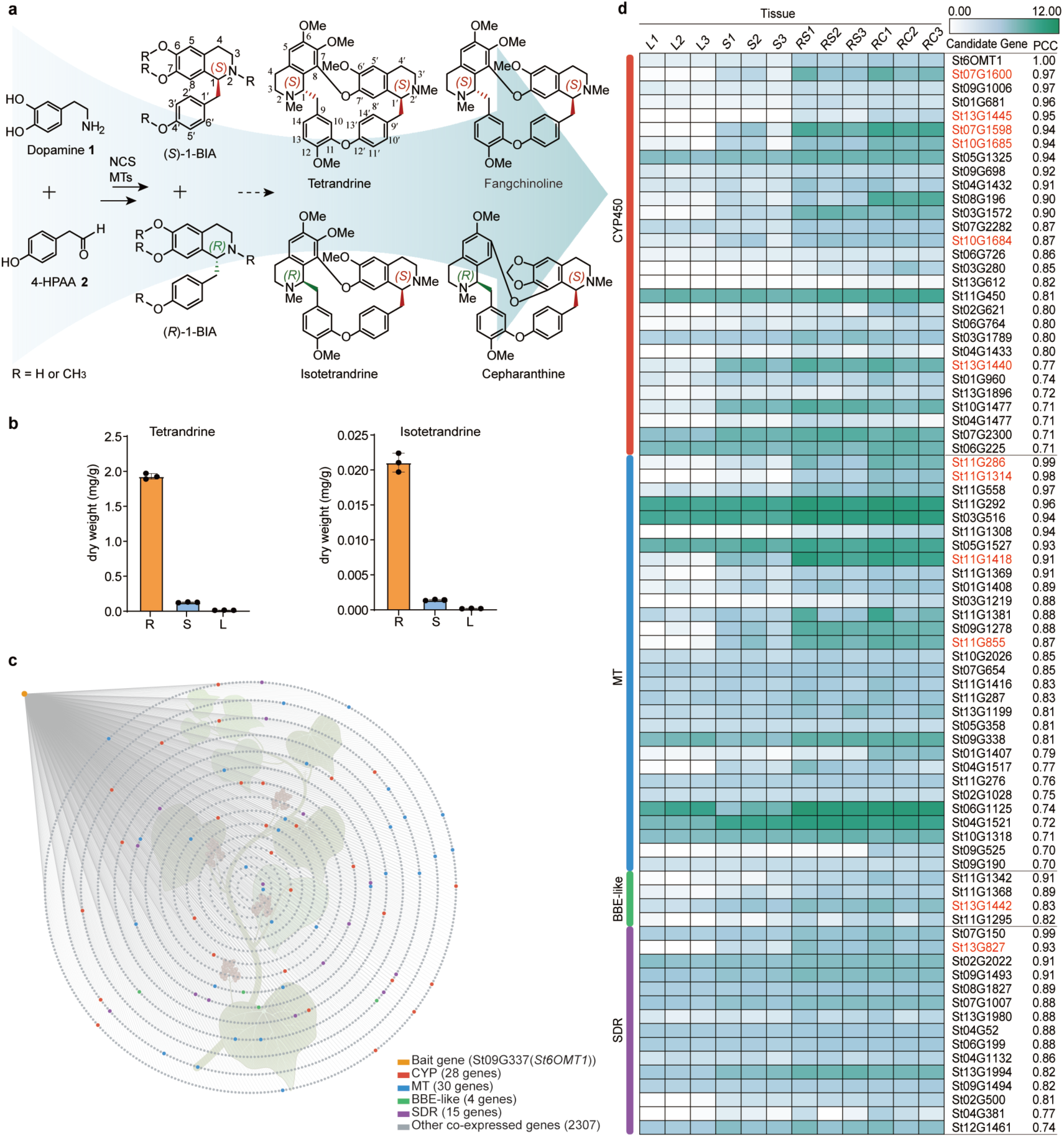
Tissue-specific co-expression profiling and genome sequencing enable the identification of candidate enzymes for bisbenzylisoquinoline alkaloid biosynthesis in *Stephania tetrandra.* **a**, Proposed biosynthetic pathway to representative bisbenzylisoquinoline alkaloids (bisBIAs). The pathway proceeds from dopamine **1** and 4-hydroxyphenylacetaldehyde (4-HPAA, 2) to 1-BIA monomers, followed by sequential intermolecular and regioselective intramolecular C-O oxidative coupling to form cepharanthine-type (C7-O-C8′) and tetrandrine/isotetrandrine-type (C7′-O-C8) scaffolds. A detailed illustration of the previously proposed biosynthetic pathway is provided in Supplementary Fig. 1. NCS, norcoclaurine synthase; 1-BIA, 1-benzylisoquinoline alkaloid. b, Tissue-specific accumulation of major bisBIAs in *S. tetrandra*. Quantitative liquid chromatography-tandem mass spectrometry (LC–MS) analysis of roots (mixed root cortex and root stele), stems, and leaves from one-year-old plants revealed that both tetrandrine and isotetrandrine accumulate predominantly in the roots, with levels ∼100-fold higher than in aerial tissues. Tetrandrine is the major bisBIA, accumulating to levels ∼100-fold higher than isotetrandrine in the roots. **c**, Co-expression analysis with *St6OMT1* as the bait gene identifies a suite of candidate biosynthetic genes. Using the previously characterized *norcoclaurine 6-O-methyltransferase* (*St6OMT1*) as the bait, we identified 2,384 co-expressed genes based on the Pearson correlation coefficient (PCC) across the leaf, stem, root stele, and root cortex transcriptomes, mapping to the newly sequenced and assembled *S. tetrandra* genome. Colored dots indicate candidates from key enzymatic families. CYP, cytochrome P450; MT, methyltransferase; BBE-like, berberine bridge enzyme-like oxidases; SDR, short-chain dehydrogenases/reductases. Values in parentheses indicate the total number of identified candidates within each respective enzyme family. **d**, Expression profiles of selected candidate genes across different tissues and respective biological replicates (1–3). The heatmap displays log_2_-transformed FPKM values for 77 candidate genes annotated as encoding CYPs, MTs, BBE-like, and SDRs. Genes functionally characterized in this work are highlighted in red. L, Leaf; S, stem; RC, root cortex; RS, root stele.

Structural diversity within the bisBIA class arises from a limited set of 1-BIA precursors^3,11^. Oxidative phenol coupling generates dimeric scaffolds that differ in linkage position, macrocyclization, and stereochemical configuration (Fig. 1a, Supplementary Fig. 1), producing multiple chiral centers and conformationally flexible biaryl frameworks^3,12,13^. These features generate discrete three-dimensional geometries with measurable differences in aromatic stacking, electronic delocalization, and solvent accessibility, which directly influence physicochemical behavior and ligand-binding potential across the subclass^14,15^. Despite extensive structural characterization, the biosynthetic logic governing the assembly of bisBIAs, particularly the origin and control of stereochemistry, has remained poorly understood^16–18^. Current models propose that biosynthesis begins with cytochrome P450 (CYP)-mediated intermolecular oxidative phenol coupling of two benzylisoquinoline monomers, followed by a series of downstream transformations, including intramolecular cyclization, isomerization, and regioselective methylation (Fig. 1a, Supplementary Fig. 1)^16,19^. Although CYP80 enzymes have been shown to catalyze the initial dimerization^16–18^, the enzymatic basis of macrocycle formation and subsequent stereochemical diversification remains elusive. In particular, the mechanisms governing configurational isomerization and methylation, the key drivers of bisBIA diversity-represent major gaps in the pathway. This contrasts sharply with other BIA classes, such as morphinan^20^, phthalideisoquinoline^21,22^, protoberberine^23,24^, and aporphine^25,26^, for which biosynthetic pathways have been elucidated and successfully reconstructed in microbial systems.

BisBIAs occur predominantly in plants of the Menispermaceae, Berberidaceae, Lauraceae, and Ranunculaceae families^3^. The herbaceous perennial vine *Stephania tetrandra* (Menispermaceae) stands out as a representative species in this group, synthesizing both (*S*,*S*)-configured tetrandrine and (*R*,*S*)-configured bisBIAs such as isotetrandrine and cepharanthine (Supplementary Fig. 2)^27^. However, access to these compounds at scale is constrained by slow-growing plant sources, e.g., Menispermaceae species require several years of cultivation before harvest^27^; and the difficulty of chemical synthesis^28^. These constraints have hindered systematic exploration of their therapeutic potential and strongly motivated the need for biosynthetic elucidation and heterologous production. In this work, we elucidate the biosynthetic pathway of bisBIAs in *S. tetrandra*, identifying CYP enzymes responsible for regioselective dimerization and macrocyclization, and revealing a redox-mediated mechanism that controls stereochemical configuration of the macrocyclized products. We further achieve *de novo* biosynthesis of these complex alkaloids in a heterologous host, establishing a scalable platform for their production and diversification.

### Genome sequencing of *Stephania tetrandra* and co-expression analysis empower identification of candidate enzymes for bisBIA biosynthesis

To elucidate the biosynthetic pathway of bisBIAs, we first performed transcriptome and whole-genome sequencing of *S. tetrandra*. By integrating Nanopore long-read sequencing and Hi-C technologies, we generated a chromosome-level genome assembly of 924 Mb (Supplementary Table 1), comprising 13 chromosomes -consistent with flow cytometry and karyotype analyses (Supplementary Figs. 3 and 4). Transcriptomes were analyzed from leaves (L), stems (S), root cortex (RC), and root stele (RS). Previously characterized upstream genes committed to the BIA biosynthetic pathway, including *StNCS2/4* (encoding the norcoclaurine synthases which catalyze the condensation of dopamine (**1**) and 4-hydroxyphenylacetaldehyde (4-HPAA, **2**) to form norcoclaurine (**3**))^29,30^ and *St6OMT1* (encoding the norcoclaurine 6-*O*-methyltransferase 1 which catalyzes C6-*O*-methylation of **3** to yield coclaurine (**4**))^31,32^ (Supplementary Fig. 1), were markedly enriched in transcriptomes from underground tissues compared with their aboveground counterparts, consistent with tissue-specific accumulation of bisBIAs (Fig. 1b)^29,31^. We used *St6OMT1* as a bait to identify 2,384 correlated co-expressed candidate genes involved in bisBIA biosynthesis (Pearson’s *r* > 0.7) (Fig. 1c). This dataset was rich in enzymatic families putatively involved in bisBIA biosynthesis, including CYPs, methyltransferases (MTs), berberine bridge enzyme (BBE)-like oxidases, and short-chain dehydrogenases/reductases (SDRs), which were compiled into a co-expression dataset for subsequent functional analysis (Fig. 1d).

### CYP80Q4 is an intermolecular C-O phenol-coupling enzyme for linear (*R*,*S*)-bisBIA formation

CYP80 family enzymes have been reported to catalyze phenol coupling of the benzylisoquinoline monomers (*R*)- and (*S*)-*N*-methylcoclaurine (**5**) to form (*R*,*R*)- and (*R*,*S*)-configured linear bisBIAs (Supplementary Fig. 1)^16^. However, given the confirmed presence of bisBIA structures lacking *N*-methyl groups in *S. tetrandra* such as 2′-*N*-norfangchinoline (Supplementary Fig. 2)^27^, we hypothesized that the *N*-nonmethyl 1-benzylisoquinoline alkaloid (1-BIA) monomers **3** and **4** may also serve as precursors towards linear bisBIA coupling in this species. Since linear bisBIAs are biosynthesized by phenol coupling of two phenol monomers^16^, we focused on genes encoding CYPs in the co-expression dataset. A total of 28 candidates (Fig. 1d) were first expressed in WAT11 yeast strain, and the microsomes were extracted for *in vitro* assays^33^. Both the (*R*)- and (*S*)- configured **3**, **4**, and **5** were tested as potential substrates in the assays. LC-MS analysis revealed that two of the 28 screened CYPs yielded detectable products. These two enzymes share 99% amino acid sequence identity and are encoded by tandemly duplicated loci *St07G1598* and *St07G1600*, on chromosome 7 (Supplementary Fig. 5). The enzyme encoded by *St07G1598* was subsequently designated as CYP80Q4.

CYP80Q4 catalyzed the coupling of (*R*)-**5**/(*S*)-**5** pairs to yield C3′-O-C4′ phenol-coupled product (*R*,*S*)-berbamunine ((*R*,*S*)-**6**), as assigned by comparison with the product of the well-characterized CYP80A1 from *Berberis stolonifera*^16^ (Fig. 2, Supplementary Figs. 6 and 7a-c). This reaction also produced a minor amount of (*R*,*R*)-guattegaumerine ((*R*,*R*)-**6**), a product resulting from the analogous coupling of two (*R*)-**5** monomers (Supplementary Figs. 6 and 7a-c)^16^. While (*R*)-**5**/(*S*)-**5** has been the most extensively studied substrate pair in previous reports^16,18^, our results reveal that (*S*)-**4**/(*R*)-**4** also serves as a highly robust substrate pair for CYP80Q4-mediated coupling (Fig. 2, Supplementary Fig. 6 and 7d-f). Upon utilizing (*S*)-**4**/(*R*)- **4** pairs as substrates, we detected a product mass matching the dimer of **4** ([M + H]^+^ = *m*/*z* 569), along with its characteristic doubly protonated ion ([M + 2H]^2+^ = *m*/*z* 285), typical of bisBIAs^34^. Subsequent comparison with synthetic standards using chiral stationary-phase chromatography confirmed this product as the C3′-O-C4′ phenol-coupled (*R*,*S*)-lindoldhamine ((*R*,*S*)-**8**) (Supplementary Fig. 8; see Supplementary Information for chemical synthesis procedures, and Supplementary Table 2 for the Nuclear Magnetic Resonance (NMR) data of (*R*,*S*)-**8**). CYP80Q4 also accepted heterologous monomer pairs. It converted (*R*)-**5**/(*S*)-**4** into (*R*,*S*)-configured 2′-norberbamunine ((*R*,*S*)-**7**), as confirmed by NMR analysis (Fig. 2, Supplementary Fig. 6 and 7g-j, NMR data for (*R*,*S*)-**7** in Supplementary Table 3), and (*R*)-**4**/(*S*)-**5** into 2-norberbamunine ((*R*,*S*)-**9**), tentatively assigned by MS/MS analysis (Fig. 2, Supplementary Fig. 6 and 7k-m, see Supplementary Information for structural elucidation). In contrast, no detectable activity was observed with (*R*)-**3**/(*S*)-**3** (Supplementary Fig. 6). Minor quantities of additional putative coupling products were also detected for other substrate combinations ((*R*)-**4**/(*S*)-**3,** (*R*)-**5**/(*S*)-**3** and (*R*)-**3**/(*S*)-**4** pairs), as indicated by characteristic doubly protonated ions (Supplementary Fig. 6 and 9a-c). Together, these results demonstrate that CYP80Q4 exhibits broad substrate promiscuity, enabling the formation of a diverse array of bisBIA scaffolds. These findings suggest an alternative metabolic trajectory toward bisBIAs in addition to the *N*-methylated intermediate **5**.

**Fig. 2.**
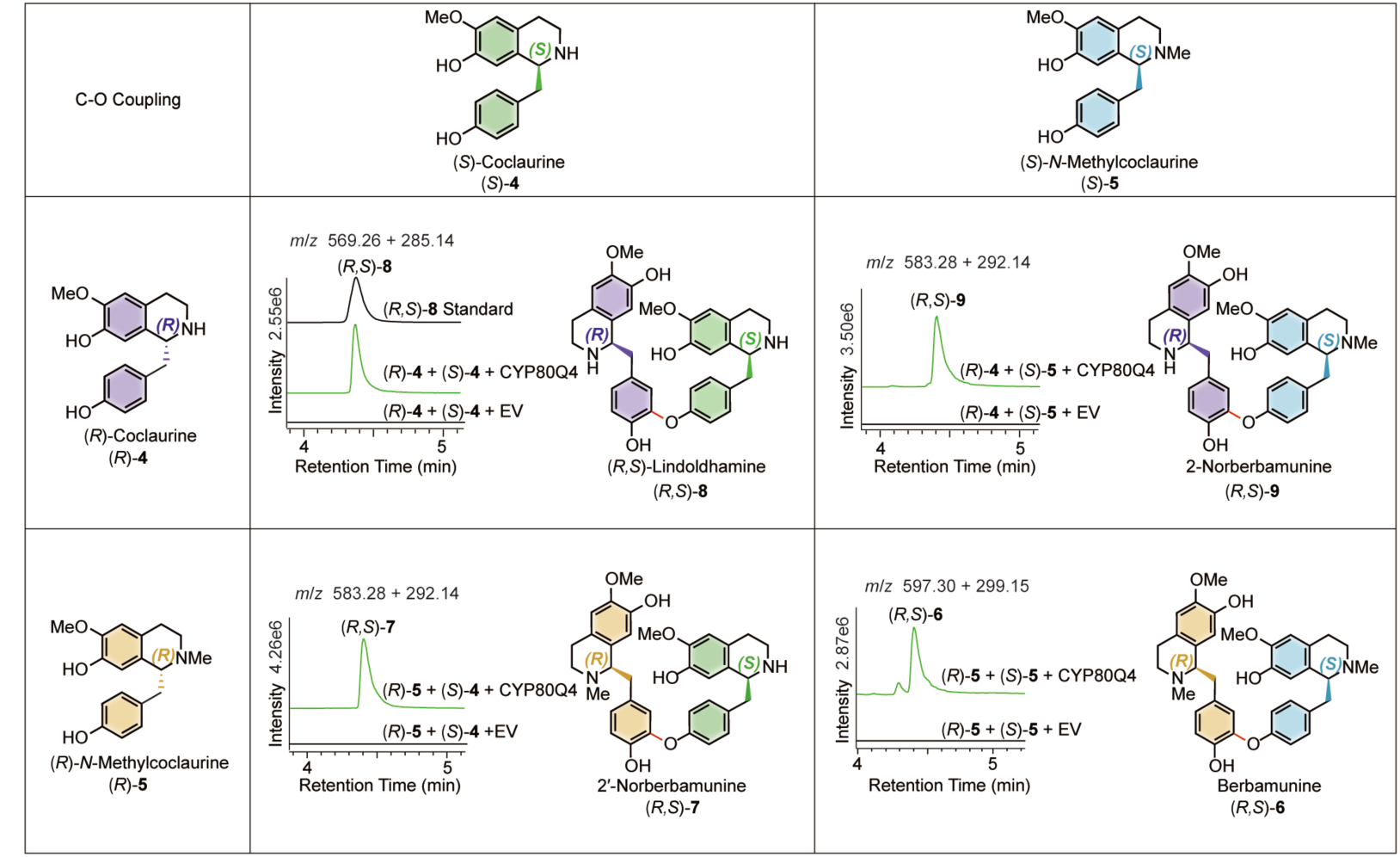
CYP80Q4 catalyzes intermolecular C-O phenol coupling to generate linear (*R*,*S*)-bisBIAs. *In vitro* microsomal assays using (*R*)- and (*S*)-configured coclaurine (**4)** and *N*-methylcoclaurine (**5)** demonstrate that CYP80Q4 efficiently mediates the C3′-O-C4′ intermolecular phenol coupling of hetero-enantiomeric substrate pairs, yielding four distinct linear bisBIAs. LC–MS analysis reveals that these bisBIAs exhibit a characteristic ionization pattern consisting of both singly ([M + H]^+^) and doubly ([M + 2H]^2+^) charged species^34^. Extracted ion chromatograms (EICs) are displayed for both ions; for example, (*R*,*S*)-**8** is detected at *m*/*z* 569.27 ([M + H]^+^) and *m*/*z* 285.14 ([M + 2H]^2+^). Hereafter, all displayed EICs for bisBIAs encompass both the [M+H]^+^ and [M + 2H]^2+^ ions.

To compare the activity of CYP80Q4 against the previously reported bisBIA-dimerizing enzymes, we synthesized *BsCYP80A1*^16^, *NnCYP80Q2*^17^, and *Bs-MaCYP80A1*^18^ and assayed these CYP80 homologs under identical conditions. Analysis of a comprehensive substrate panel revealed that all enzymes accept a broader range of 1-BIA substrates than previously recognized (Supplementary Fig. 6)^16–18^. CYP80Q4 predominantly generated (*R*,*S*)-configured dimers, whereas the other CYP80s exhibited a marked preference for coupling two (*R*)-enantiomers to form (*R*,*R*)-configured products (Supplementary Fig. 6 and 9). Notably, none of the tested CYP80 enzymes catalyzed detectable (*S*,*S*)-configured bisBIAs. This observation aligns with the presence of (*R*,*S*)-configured isotetrandrine and cepharanthine in *S. tetrandra*, yet it fails to explain the accumulation of the more abundant (*S*,*S*)-configured tetrandrine.

### CYP82BCs catalyze intramolecular phenol coupling to form cyclic bisBIA scaffolds

The biosynthesis of bioactive macrocyclic bisBIAs, such as tetrandrine, isotetrandrine, and cepharanthine, necessitates the cyclization of linear bisBIA precursors, which likely occurs via enzyme-mediated intramolecular phenol coupling (Supplementary Fig. 1)^3^. Based on the structures of bisBIAs known to accumulate in *S. tetrandra*, we inferred that there are at least two types of intramolecular coupling reactions, C7-O-C8′ phenol coupling (towards cepharanthine) and C7′-O-C8 phenol coupling (towards tetrandrine and isotetrandrine), for the assembly of complex cyclic bisBIA scaffolds (Fig. 1, Supplementary Fig. 2). To elucidate the subsequent macrocyclization steps, we established a microsomal cascade in which linear (*R*,*S*)-bisBIA intermediates were generated *in situ* and subsequently subjected to intramolecular phenol coupling to form cyclic bisBIAs. Assuming that CYPs would be plausible for such intramolecular phenol-coupling reactions, the remaining 26 CYP candidates were screened using microsomal cascade reactions (Fig. 1d). Microsomes expressing each candidate were co-incubated with microsomes expressing CYP80Q4, which served to generate the requisite linear substrates. The reactions were initiated by the addition of 1-BIA monomer pairs ((*R*)-**4**/(*S*)-**4**, (*R*)-**4**/(*S*)-**5**, (*R*)-**5**/(*S*)-**4**, and (*R*)-**5**/(*S*)-**5**) designed to produce the linear intermediates (*R*,*S*)-**8**, (*R*,*S*)-**9**, (*R*,*S*)-**7**, and (*R*,*S*)-**6**, respectively (Fig. 2).

This screening identified four active enzymes belonging to the CYP82 family, designated as CYP82BC1-4. In combination with CYP80Q4, these enzymes generated new products with a characteristic mass shift of –2 Da relative to the linear intermediates, consistent with the loss of two protons during intramolecular phenol coupling (Fig. 3, Supplementary Fig. 10). CYP82BC1 and CYP82BC4, which share 92.3% (Supplementary Fig. 11a) amino acid sequence identity and are encoded by tandemly duplicated loci *St10G1685* and *St10G1684* on chromosome 10 (Supplementary Fig. 5), displayed dual regioselectivity in mediating intramolecular phenol coupling. Large-scale purification and NMR analysis confirmed that both enzymes catalyzed C7-O-C8′ coupling and C7′-O-C8 coupling of (*R*,*S*)-**8** and (*R*,*S*)-**7** to yield cepharanthine-type scaffolds bisnoraromoline ((*R*,*S*)-**16**) and racemosinine A ((*R*,*S*)-**18**), as well as isotetrandrine-type scaffolds bisnorobamegine ((*R*,*S*)-**17**) and 2′-norobamegine ((*R*,*S*)-**19**) (Fig. 3, Supplementary Fig. 10a-h, NMR data for (*R*,*S*)-**16**, (*R*,*S*)-**17**, (*R*,*S*)-**18**, (*R*,*S*)-**19** in Supplementary Table 4–7). In contrast, CYP82BC2 and CYP82BC3, which share 80.6% (Supplementary Fig. 11b) amino acid sequence identity and are encoded by tandem loci *St13G1440* and *St13G1445* on chromosome 13 (Supplementary Fig. 5), showed strong regioselectivity. These enzymes predominantly catalyzed C7′-O-C8-coupling, converting (*R*,*S*)-**8** and (*R*,*S*)-**7** into the corresponding isotetrandrine-type macrocycles (*R*,*S*)-**17** and (*R*,*S*)-**19**, with only trace amounts of the C7-O-C8′ isomers detected (Fig. 3, Supplementary Fig. 10a-h). Notably, the cyclization efficiency was substrate-dependent, with only trace amounts of the cyclized product detected from the *N*,*N*′-dimethylated substrate (*R*,*S*)-**6** (Supplementary Fig. 10i-l) and none from the *N*′-methylated substrate (*R*,*S*)-**9**. This enzymatic preference suggests that the *N*′-methylation of bisBIAs is likely after macrocyclization. Moreover, we also challenged CYP82BC enzymes with synthetic (*R*,*R*)-, (*S*,*R*)-, and (*S*,*S*)-**8** diastereomers (see Supplementary Information for chemical synthesis procedures) to define stereochemical specificity. No cyclized products were detected in these reactions, indicating that CYP82BC enzymes are highly selective for (*R*,*S*)-configured linear bisBIA substrates.

**Fig. 3.**
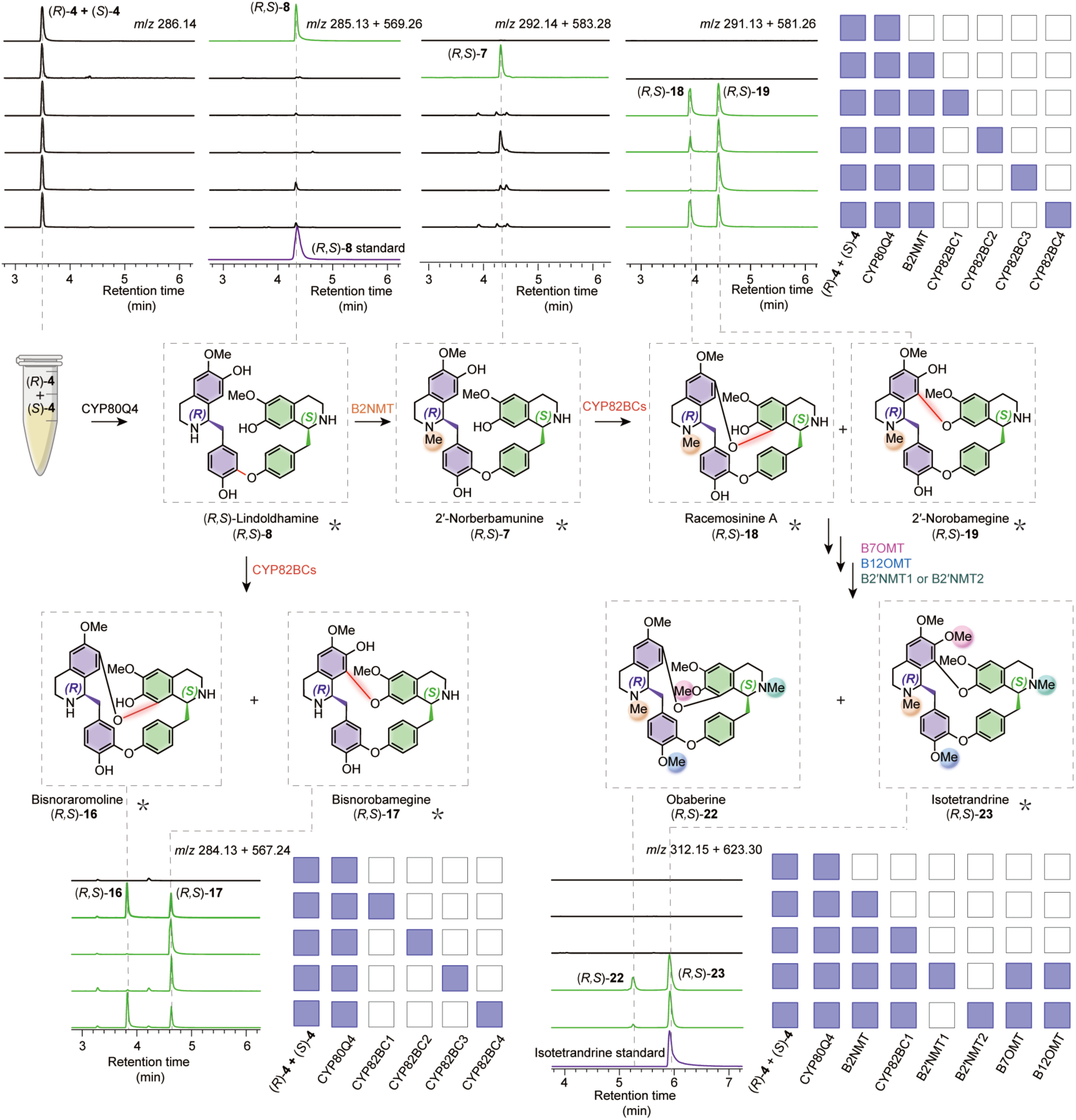
Elucidation of biosynthetic pathways leading to (*R*,*S*)-configured bisBIAs obaberine and isotetrandrine. Center: Schematic representation of the complete biosynthetic route to obaberine ((*R*,*S*)-**22**) and isotetrandrine ((*R*,*S*)-**23**) in *S. tetrandra*, as deciphered by multienzyme cascade assays *in vitro*. Using (*R*)-**4** and (*S*)-**4** as starting substrates, the cascade was initiated by CYP80Q4 to generate the linear intermediate (*R*,*S*)-**8**, followed by the sequential action of CYP82BCs and MTs. CYP82BCs mediate the intramolecular cyclization of linear precursors (*R*,*S*)-**8** and (*R*,*S*)-**7** via distinct C7-O-C8′ and C7′-O-C8 regiochemistries. Among the MTs, B2NMT specifically methylates the linear substrate (*R*,*S*)-**8** at the *N*2 position, whereas B7OMT, B2′NMT1/2, and B12OMT catalyze the methylation of the cyclic intermediates (*R*,*S*)-**18** and (*R*,*S*)-**19** at the C7/C7′-hydroxyl, *N*2′, and C12-hydroxyl positions, respectively, in a flexible metabolic grid to yield (*R*,*S*)-**22** and (*R*,*S*)-**23**. Top and Bottom Panels: LC–MS analysis of *in vitro* cascade reactions. Each row represents a distinct reaction mixture, with the matrix indicating the presence (purple squares) or absence (white squares) of the indicated enzymes and substrates. Green traces represent EICs of enzymatic products; purple traces represent authentic standards. Colored spheres on the chemical structures correspond to the specific methyltransferases catalyzing methylation at those positions. Asterisks indicate compounds structurally verified by NMR or confirmed by comparison with authentic standards; (*R*,*S*)-**22** was identified by comparing its MS/MS spectrum with that of the (*R*,*S*)-**23** standard (see the Supplementary Information for detailed structural elucidation).

Our analysis of the CYP82BC enzymes in *S. tetrandra* revealed that metabolite abundance correlates with enzyme regioselectivity. While all four CYP82BCs catalyze the formation of C7′-O-C8 linkage, only two show activity toward C7-O-C8′ coupling matching the phytochemical profile of *S. tetrandra*, where isotetrandrine/tetrandrine-type bisBIAs predominate over cepharanthine congeners^35^.

### Five methyltransferases catalyze heteroatom methylation in bisBIA biosynthesis

To identify the enzymes bridging the gap between the cyclized scaffolds and methyl-decorated bisBIAs, we screened 30 candidate MTs from the co-expression dataset (Fig. 1d). The *N*-nonmethyl monomers (*R*)-**4** and (*S*)-**4** were used to initiate the cascade containing CYP80Q4, CYP82BC1 and each candidate MT. LC-MS analysis revealed that reactions containing five candidates yielded new peaks exhibiting a +14 Da mass shift relative to the cyclic scaffolds (*R*,*S*)-**16**, (*R*,*S*)-**17**, consistent with single methylation events (Fig. 3, Supplementary Fig. 12). Based on chromatographic retention time shifts reflecting changes in polarity, these MTs were provisionally classified as three *N*-methyltransferases (NMTs) and two *O*-methyltransferases (OMTs).

We performed large-scale reactions and purified the resulting compounds for NMR analysis, enabling precise assignment of NMT activities. One enzyme, designated as bisbenzylisoquinoline 2-*N*-methyltransferase (B2NMT), catalyzed methylation at the *N*2 position to yield (*R*,*S*)-**18** and (*R*,*S*)-**19**, as confirmed by comparison with the cyclized products generated from (*R*)-**5** and (*S*)-**4** via the CYP80Q4 and CYP82BC1 catalytic cascade. (Supplementary Fig. 12a, b). Similarly, NMR characterization of (*R*,*S*)-**35** and (*R*,*S*)-**36** defined two paralogous enzymes, B2′NMT1 and B2′NMT2 (71.6% shared amino acid sequence identity) (Supplementary Fig. 13), as 2′-*N-*methyltransferase (Supplementary Fig. 12a, b, g, h; NMR data for (*R*,*S*)-**35** and (*R*,*S*)-**36** in Supplementary Tables 8 and 9). For the OMTs, catalytic positions were inferred from diagnostic MS/MS fragmentation patterns. One candidate OMT produced a +14 Da shift in the dimeric isoquinoline fragment (*m*/*z* 353) relative to the substrates, indicating methylation at either the *O-*7 or *O-*7′ position to yield (*R*,*S*)-**30** and (*R*,*S*)-**31** (Supplementary Fig. 12a-d; see Supplementary Information for structural elucidation). We consequently designated this enzyme bisbenzylisoquinoline 7-*O*-methyltransferase (B7OMT). Its ability to methylate either position likely reflects the close spatial proximity of the *O*7 and *O*7′ hydroxyl groups within the macrocycle. Conversely, products of the second OMT retained the unshifted isoquinoline fragment (*m*/*z* 339), while the diphenyl ether-associated fragment (*m*/*z* 227) exhibited a +14 Da shift. This fragmentation signature localized the methylation to the *O-*12 position, yielding (*R*,*S*)-**33** and (*R*,*S*)-**34**, characterizing the enzyme as bisbenzylisoquinoline 12-*O*-methyltransferase (B12OMT) (Supplementary Fig. 12a, b, e, f; see Supplementary Information for structural elucidation).

In contrast to the positional promiscuity reported for 1-BIA MTs^36^, bisBIA MTs display high regioselectivity, likely reflecting the greater spatial separation of reactive sites in the dimeric scaffold. An exception is B7OMT, consistent with the proximity of its target hydroxyl groups. Finally, by integrating CYP80Q4, CYP82BC1, and four MTs (B7OMT, B12OMT, B2NMT, either B2′NMT1 or B2′NMT2) into a cascade reaction, we successfully obtained the fully methylated products, isotetrandrine ((*R*,*S*)-**23**), as confirmed by comparison with an authentic standard (Fig. 3, Supplementary Fig. 14). In parallel, we detected a second product, tentatively assigned as obaberine ((*R*,*S*)-**22**), whose structure was inferred from MS/MS analysis and the established regioselectivities of the combined MTs (Fig. 3, Supplementary Fig. 14, see Supplementary Information for structural elucidation).

To further determine substrate specificity, recombinant MTs were assayed *in vitro* using both linear (*R*,*S*)-**8** and its cyclic scaffolds (*R*,*S*)-**16** and (*R*,*S*)-**17** as substrates. Four MTs (B2′NMT1, B2′NMT2, B7OMT, and B12OMT) acted on both linear (*R*,*S*)-**8** and cyclic scaffolds (*R*,*S*)-**16**, and (*R*,*S*)-**17** (Supplementary Figs. 15-17; see Supplementary Information for structural elucidation of linear bisBIA products), whereas B2NMT exhibited strict specificity for the linear substrate (*R*,*S*)-**8** to produce (*R*,*S*)-**7** (Supplementary Fig. 18). This indicates that *N*2-methylation precedes intramolecular cyclization in the pathway.

### A dual-enzyme redox system catalyzes stereochemical inversion of (*R*,*S*)- to (*S*,*S*)-bisBIAs

Although the (*S*,*S*)-configured tetrandrine is the most abundant bisBIA in *S. tetrandra*, CYP80Q4 does not generate (*S*,*S*)-configured linear bisBIAs, and CYP82BCs do not catalyze their subsequent cyclization. This indicates that cyclic (*S*,*S*)-bisBIAs are likely formed through the epimerization of cyclic (*R*,*S*)-bisBIAs. Mechanistically, such a stereochemical inversion typically requires a two-step redox process involving an initial oxidative desaturation to a planar intermediate, followed by a stereoselective reduction. A notable parallel exists in opium poppy, where reticuline epimerase (REPI, a CYP-aldo-keto reductase fusion protein) drives a redox-mediated configurational reset via a transient 1,2-dehydroreticuline intermediate^37,38^. However, no REPI-like fusion protein homologs were identified in the *S. tetrandra* genome, and assays of discrete CYP and aldo-keto reductase homologs using cyclic (*R*,*S*)-**17** as a substrate yielded no detectable products. This suggests that a different class of enzymes is responsible for this conversion.

Guided by this observation, we examined our co-expression dataset for BBE-like oxidases, which are known to catalyze C-N bond desaturation in BIA biosynthesis^24,39^. This led to the identification of four candidate enzymes (Fig. 1d), including *St13G1442,* which is genomically clustered with *CYP82BC2* and *CYP82BC3* on chromosome 13 (Supplementary Fig. 5). *In vitro* assays using (*R*,*S*)-**17** as substrate revealed that St13G1442 catalyzes the formation of a dehydrogenated product showing a –2 Da mass shift, consistent with formation of the planar imine intermediate, 1,2-dehydrobisnorobamegine (**24**). The identity of this intermediate was further supported by its subsequent stereochemical inversion to (*S*,*S*)-**17** upon reduction (Fig. 4a, Supplementary Fig. 19). We therefore designated this enzyme as bisbenzylisoquinoline oxidase (BBOX). BBOX activity is strictly dependent on the C7′-O-C8 macrocycle and the presence of a secondary amine, showing no activity toward the C7-O-C8′ regioisomer (*R*,*S*)-**16** or the *N*2-methylated analog (*R*,*S*)-**19** (Supplementary Fig. 19d-g). This specificity reflects the instability of the corresponding iminium intermediate from tertiary amines under physiological conditions^40^, and implies that oxidative activation precedes *N*2-methylation in the pathway.

**Fig. 4.**
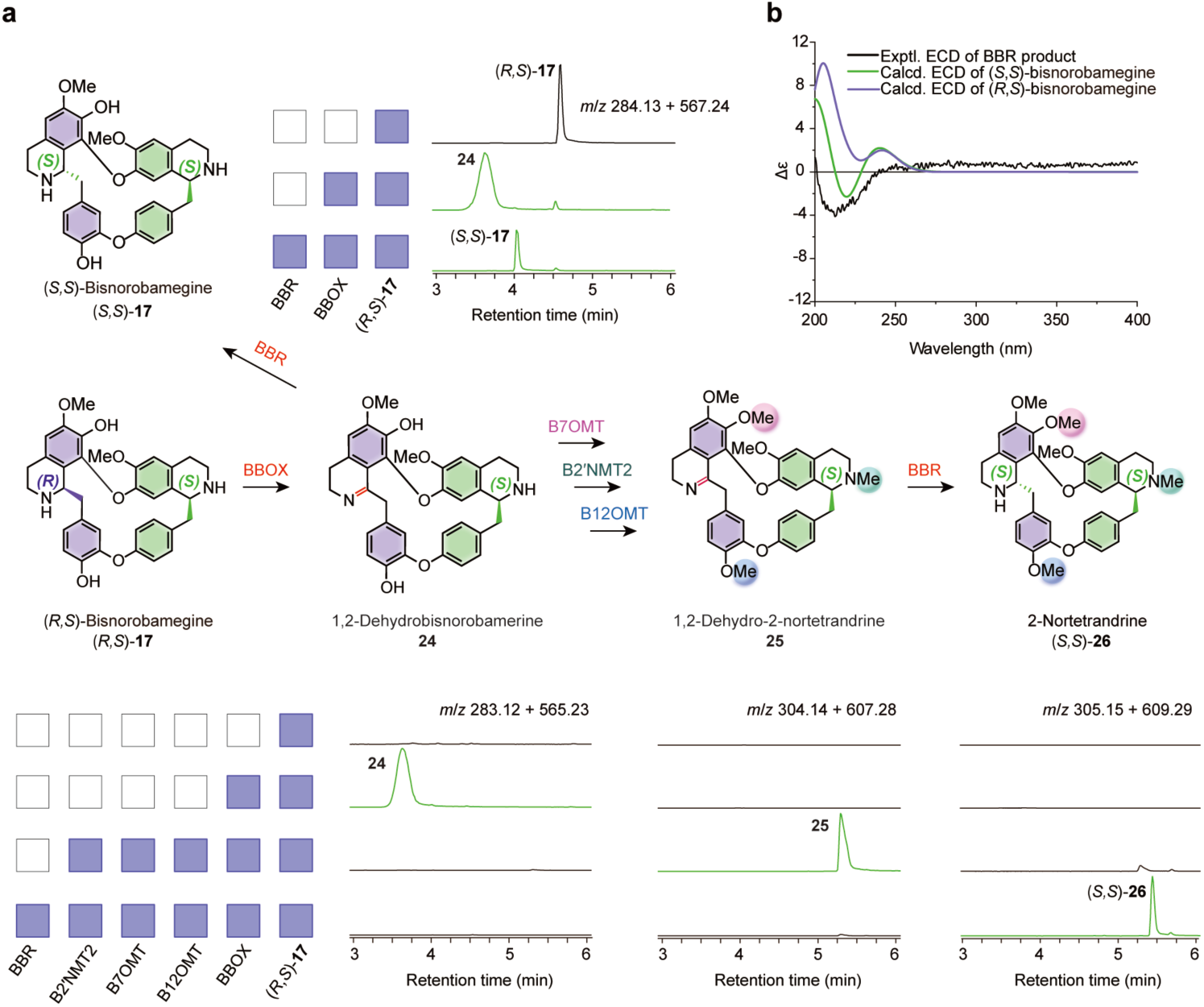
BBOX-BBR catalyze the redox-mediated epimerization of cyclic (*R,S*)- to (*S,S*)-bisBIA. **a**, *In vitro* reconstitution of the BBOX-BBR-catalyzed epimerization cascade. Incubation of (*R*,*S*)-**17** with yeast-expressed BBOX led to its efficient conversion into the oxidized intermediate **24**, as shown by EICs. Subsequent addition of bacterially expressed BBR catalyzed the stereospecific reduction of **24** to a new product, (*S*,*S*)-**17**, which eluted at a distinct retention time relative to (*R*,*S*)-**17**. Intermediate **24** can be intercepted by tailoring MTs (B7OMT, B2′NMT2, B12OMT) to form the trimethylated intermediate **25**. BBR also accepted **25** as a substrate, converting it to (*S*,*S*)-**26**. Each row represents a distinct reaction mixture, with the matrix indicating the presence (purple squares) or absence (white squares) of the indicated enzymes and substrates. Green traces represent EICs of enzymatic products. Colored spheres on the chemical structures indicate the specific sites methylated by the corresponding MTs. **b**, Absolute stereochemistry of the BBR product confirmed by electronic circular dichroism (ECD) spectroscopy. The experimental ECD spectrum of enzymatically produced (*S,S*)-17 closely matches the calculated ECD spectrum for the (*S,S*)-configuration and is distinct from that of the (*R,S*)-**17** substrate, establishing the (*S,S*)-stereochemistry.

To identify the reductase responsible for the stereospecific reduction of intermediate **24**, we screened 15 SDRs from the co-expression dataset (Fig. 1d). These candidates were tested in coupled assays with BBOX using (*R*,*S*)-**17** as substrate. One candidate, an atypical SDR (encoded by *St13G827*), converted the planar imine intermediate **24** to a product with a +2 Da mass shift (Fig. 4a, Supplementary Fig. 20). Although this product shared identical mass and MS/MS fragmentation with (*R*,*S*)-**17**, it exhibited a distinct retention time (Supplementary Fig. 20). Electronic circular dichroism (ECD) spectroscopy confirmed this product as the (*S*,*S*)-configured diastereomer, confirming inversion at the C1 stereocenter (Fig. 4b). We designated this enzyme as bisbenzylisoquinoline reductase (BBR). Notably, BBR is also encoded on chromosome 13, although outside the *BBOX/CYP82BC* cluster (Supplementary Fig. 5), suggesting coordinated but spatially distributed genetic control.

In contrast to fused epimerization enzymes, this two-enzyme architecture creates a temporal window in which the planar iminium intermediate (**24**) can be intercepted by tailoring MTs. Indeed, incubation of the BBOX reaction mixture with B2′NMT2, B7OMT and B12OMT -either individually or in combination -generated mono- and trimethylated oxidized products, reflected by +14 Da and +42 Da mass shifts, respectively. MS/MS analysis confirmed that each MT maintained its regioselectivity on the oxidized scaffold (Fig. 4 and Supplementary Fig. 21; see Supplementary Information for structural elucidation). Subsequent reduction of the trimethylated intermediate (**25)** by BBR successfully yielded the fully elaborated (*S*,*S*)-configured product (*S*,*S*)-**26** (Fig. 4, Supplementary Fig. 22). This oxidation–methylation–reduction sequence defines a redox-mediated “stereochemical editing” mechanism that couples epimerization with late-stage methylation, thereby expanding the accessible chemical space of (*S*,*S*)-configured bisBIAs.

### *De novo* biosynthesis of canonical and novel bisBIAs in yeast reveals modular methylation control

To evaluate whether the pathway logic elucidated in *S. tetrandra* could be transferred to a heterologous host, we reconstituted the bisBIA pathway in *Saccharomyces cerevisiae* to establish a microbial production platform. The pathway was assembled stepwise by introducing discrete enzymatic modules for monomer supply, oxidative coupling, macrocyclization, methylation, and stereochemical editing. As a starting chassis, we used strain XJ0636, which produces over 300 mg/L (*S*)-**3** in shake-flask cultures^24^. Genotypes of all constructed strains are shown in Supplementary Table 10.

Chromosomal integration of *6OMT* from *P. somniferum* (*Ps6OMT*) yielded strain XJ06360, which produced (*S*)-**4** (Fig. 5a, Supplementary Table 10)^32^. Given that bisBIA formation requires both (*R*)- and (*S*)-configured monomers, we next introduced the DRS-DRR enzyme module (PrDCS from *P. rhoeas* and PbDCR from *P. bracteatum*), previously shown to catalyze the epimerization of (*S*)-**5** to (*R*)-**5** and (*S*)-reticuline to (*R*)-reticuline^18^. Since no homologs have been identified, whether the PrDCS-PbDCR module could similarly mediate the epimerization of *N*-nonmethyl 1-BIA monomers (*S*)-**4** to (*R*)-**4** within our engineered context was thus explored. Integration of *PrDCS* and *PbDCR*, together with *CYP80Q4* in XJ06360 yielded strain XJ06388 (XJ06360 + PrDCS + CYP80Q4 + PbDCR), which produced the linear bisBIA (*R*,*S*)-**8**, indicating successful *in vivo* epimerization of (*S*)-**4** to (*R*)-**4** and confirming that both enantiomers act as necessary substrates for CYP80Q4-mediated coupling (Fig. 5b, Supplementary Fig. 23). Increasing the copy number of *PrDCS*, *PbDCR*, and *CYP80Q4* further improved production, yielding strain XJ06399 that efficiently produced the linear (*R*,*S*)-bisBIA precursor (Fig. 5b).

**Fig. 5.**
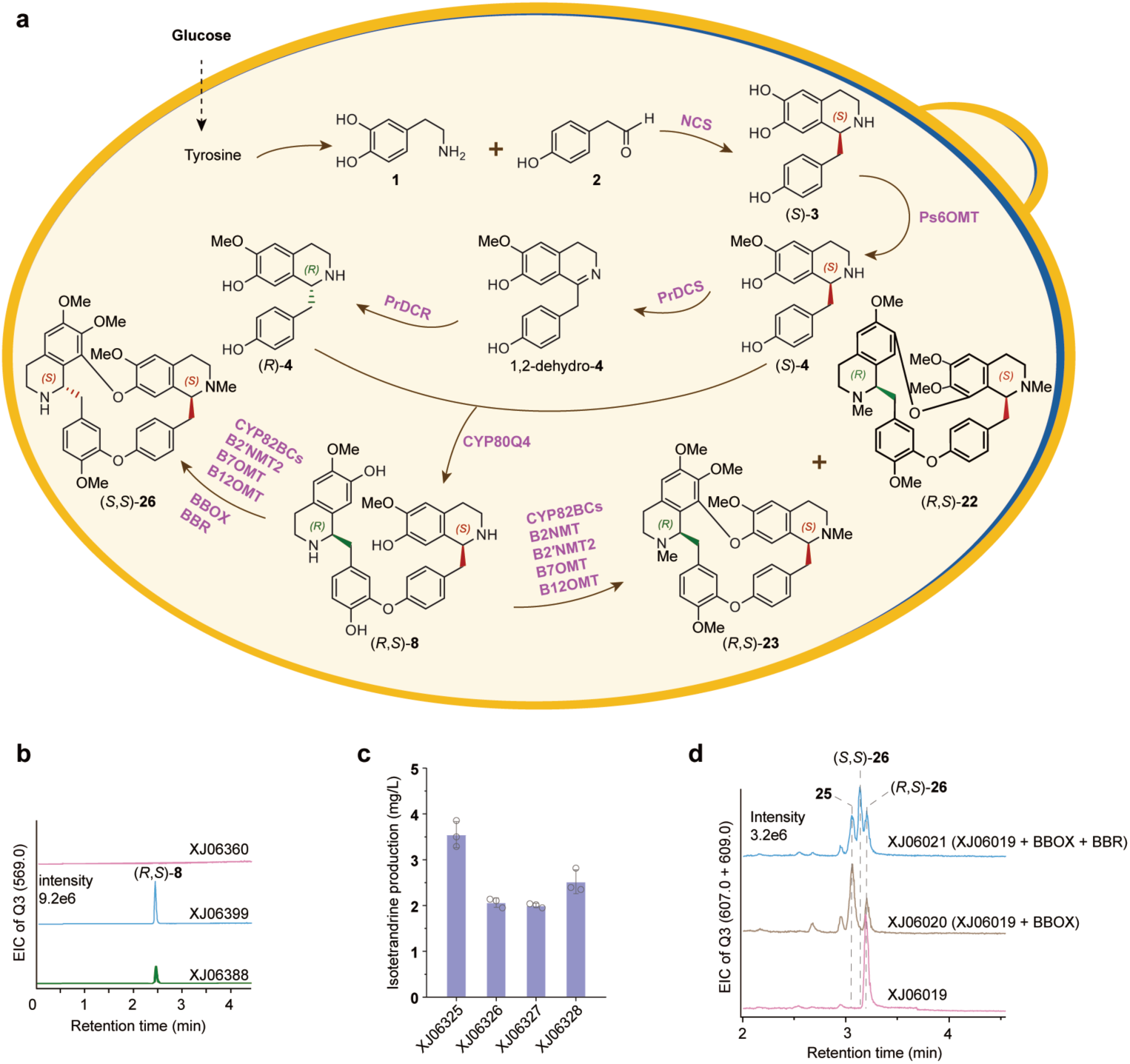
Modular expression in yeast enables efficient *de novo* production and structural diversification of bisBIAs. **a**, Pathway design for the production of diverse bisBIAs in yeast. The optimized chassis XJ06399 was developed by enhancing the precursor supply in strain XJ06388 through the integration of a single copy *Ps6OMT* and multiple copies of *PrDCS, PbDCR*, and *CYP80Q4*. The systematic addition of CYP82BCs, MTs, and the BBOX-BBR epimerization module enables the tailored production of both (*R*,*S*)- and (*S*,*S*)-configured cyclic bisBIAs. **b**, Total ion chromatogram (TIC) of Q3 (*m*/*z* 569.0 [M + H]^+^) comparing the production of the pivotal linear intermediate (*R*,*S*)-**8** in the starting strain XJ06360 (derived from XJ0636 to produce (*S*)-**4**) and the optimized chassis XJ06399. The accumulation of (*R*,*S*)-**8** in XJ06399 validates the success of the multi-copy integration and pathway-optimization strategy. **c**, Comparative evaluation of four CYP82BCs in the background strain XJ06320 carrying B2NMT, B12OMT, B7OMT, and B2′NMT2. The evaluated strains include XJ06325 (XJ06320 + *CYP82BC3*), XJ06326 (XJ06320 + *CYP82BC1*), XJ06327 (XJ06320 + *CYP82BC2*), and XJ06328 (XJ06320 + *CYP82BC4*). Among these, the strain expressing CYP82BC3 (XJ06325) exhibited the highest catalytic efficiency for the formation of cyclic bisBIAs, resulting in the highest accumulation of (*R*,*S*)-**23**. Data represent the mean ± s.d. from *n* = 3 independent biological replicates. **d**, TIC of Q3 multiple ions (MI) (*m*/*z* 607.0 and 609.0 [M + H]^+^) from strains XJ06019 (derived from XJ06399 by integrating *CYP82BC3*, *B12OMT*, *B7OMT*, and *B2′NMT2*), XJ06020 (XJ06019 + *BBOX*), and XJ06021 (XJ06019 + *BBOX* + *BBR*), demonstrating the stereochemical transition from (*R*,*S*)- to (*S*,*S*)-configuration. Heterologous expression of the *BBOX*-*BBR* module resulted in the formation of the transient imine intermediate **25** (XJ06020) and the final (*S*,*S*)-configured product (*S*,*S*)-**26** (XJ06021), confirming successful redox-controlled stereochemical editing.

Subsequent introduction of CYP82BC enzymes (CYP82BC1-4) together with four bisBIA MTs (*B2NMT, B2*′*NMT2, B7OMT and B12OMT*) in strain XJ06399 enabled conversion of the linear (*R*,*S*)-configured intermediate into cyclic bisBIAs, demonstrating that both oxidative macrocyclization and downstream tailoring reactions are functional in yeast and preserve the stereochemical constraints observed *in vitro* (Supplementary Fig. 24). Across the resulting strains, bisBIA production was readily detected, including the fully methylated cyclic bisBIAs (*R*,*S*)-**22** and (*R*,*S*)-**23**. Among these strains, XJ06325 expressing *CYP82BC3* in combination with the four MTs yielded the highest accumulation (3.6 mg/L) of the fully methylated cyclic bisBIA (*R*,*S*)-**23** (Fig. 5c, Supplementary Fig. 24).

To investigate how methylation contributes to structural diversification, we systematically examined combinations of MTs using strain XJ06322, which produces both (*R*,*S*)-**16** and (*R*,*S*)-**17** (Supplementary Fig. 23). Expression of individual MTs revealed distinct regioselectivity: *B2NMT*, *B2′NMT2*, or *B12OMT* each resulted in two methylated bisBIAs, whereas *B7OMT* yielded a single product (Supplementary Fig. 25). Combinatorial expression of MT pairs further expanded product diversity, generating multiple derivatives with mass shifts corresponding to sequential methylation events ([M + H]^+^ = *m*/*z* 595.0; Supplementary Fig. 26). Addition of a third MT yielded further derivatives with *m*/*z* 609.0 ([M + H]^+^), consistent with further heteroatom methylation (Supplementary Fig. 27). These results demonstrate that modular assembly of MTs enables predictable expansion of bisBIA chemical diversity *in vivo*, providing a versatile platform for generating structurally and potentially functionally diverse bisBIA derivatives (Supplementary Fig. 28).

To reconstitute stereochemical editing in yeast, we introduced the *BBOX-BBR* redox module into strain XJ06019 (XJ06321 + B2′NMT2 + B7OMT + B12OMT), a cyclic (*R*,*S*)-bisBIA-producing strain lacking B2NMT. Expression of BBOX alone (strain XJ06020, Supplementary Table 10) produced an oxidized intermediate with *m*/*z* 607.0 [M + H]^+^, while co-expression of BBOX and *BBR* (strain XJ06021, Supplementary Table 10) yielded cyclic (*S*,*S*)-**26**, as confirmed by MS/MS spectral matching with (*R*,*S*)-**26** (Fig. 5d, Supplementary Fig. 29).

Collectively, these results establish a modular yeast platform for bisBIAs biosynthesis, enabling production of both native and novel compounds with diverse methylation patterns and stereochemistries. Starting from (*S*)-**4**, the engineered strains produced at least 21 cyclic (*R*,*S*)-configured bisBIAs, with further diversification achievable via combinatorial methylation.

### Conclusions

Here, using genome and transcriptome data from *S. tetrandra*, we identified and characterized five CYPs, one BBE-like oxidase, one SDR, and five methyltransferases involved in bisBIA biosynthesis, successfully elucidating the complete pathways to multiple (*R*,*S*)- and (*S*,*S*)-configured bisBIAs. This establishes the biochemical logic underlying cyclic bisBIA formation, revealing how post-cyclization redox reactions control stereochemistry within a dimeric scaffold. By defining the mechanism that converts (*R*,*S*) to (*S*,*S*)-configured bisBIA and demonstrating its functional integration in a reconstructed pathway, we unify previously discrete aspects of bisBIA biosynthesis -oxidative coupling, redox inversion, and methylation diversification -into a single coherent framework. These findings expand the known strategies by which plant alkaloid pathways generate molecular complexity and create new opportunities to explore stereochemical control, enzymatic modularity, and metabolite diversification in engineered systems.

The successful reconstitution of this pathway in a heterologous yeast host marks a transition from strictly descriptive biochemistry to synthetic biology. By integrating the CYP82BCs, the promiscuous methyltransferase network, and the BBOX-BBR editing module, we have a new biosynthetic platform. This modular system allows for the combinatorial assembly of pathway enzymes to generate not only the naturally occurring (*S*,*S*)- and (*R*,*S*)-bisBIAs but also a library of previously undescribed congeners with novel methylation patterns and stereochemical configurations. Given the potent antiviral and anti-inflammatory properties of tetrandrine and cepharanthine, this pathway engineering overcomes the supply bottleneck associated with harvesting slow-growing wild plants. Moreover, it provides immediate access to a diverse array of “new-to-nature” derivatives, serving as a sustainable approach for structure-activity relationship studies and the development of next-generation therapeutics.

## Data availability

All gene sequences analyzed in this study are provided in the main text or Supplementary Materials. The raw genomic and transcriptomic sequencing data, along with the chromosome-level genome assembly generated in this study, have been deposited in the National Genomics Data Center (NGDC) under BioProject accession code PRJCA056975. The genome assembly and annotation of *S. tetrandra* have been deposited in Figshare (https://doi.org/10.6084/m9.figshare.31220782). All datasets will be publicly accessible upon publication.

## Author contributions

L.H., J.G., Y.C., J.N. and Q.L. conceived and designed the entire research plans; Q.L., X.L. and X.J. performed most of the experiments; X.L., X.T., and Y.W. helped Q.L. to identify the structure of enzyme-catalyzed reaction products; Y.M., J.W., T.C., P.S. and Y.Z. participated in some of the experiments; Q.L., X.L., X.J. and W.X. analyzed results of the experiments; Q.L., X.L., J.G., X.J., Y.C. and L.H. wrote the manuscript; G.C., J.N. and Y.H. revised the manuscript. All authors read and approved the contents of this paper.

## Acknowledgements

This work was supported by the Scientific and Technological Innovation Project of CACMS (CI2023D002), the National Natural Science Foundation of China (82504968, 82404789, 82011530137, 31961133007), the National Key R&D Program of China (2020YFA0908000), the Postdoctoral Fellowship Program of CPSF under Grant Number GZC20252587, Key project at central government level: The ability to establish sustainable use of valuable Chinese medicine resources (2060302). The authors would like to acknowledge funding from Vetenskapsrådet, Stiftelsen för internationalisering av högre utbildning och forskning.

## Methods

### Plant Materials

Cultivated *Stephania tetrandra* plants (1 year old) were collected from the traditional Chinese medicinal herb cultivation base in Yichun City, Jiangxi Province, China. For genome sequencing, fresh young leaves were collected for DNA extraction. For transcriptome profiling, root cortex, root stele, stem, and leaf tissues were collected with three biological replicates per tissue, flash-frozen in liquid nitrogen, and stored at −80 °C until RNA extraction. High quality DNA and RNA samples were sent to Wuhan Benagen Technology Co., Ltd. (Wuhan, China) for genome sequencing.

### BIA standards

Authentic standards of (*R*)- and (*S*)-norcoclaurine (**3**), (*R*)- and (*S*)-coclaurine (**4**), and (*R*)-and (*S*)-*N*-methylcoclaurine (**5**) were obtained from WuXi AppTec. The (*R*,*S*)-, (*S*,*R*)-, (*R*,*R*)-, and (*S*,*S*)-stereoisomers of lindoldhamine (**8**) were synthesized as detailed in the Supplementary Information. All other authentic standards were purchased from Shanghai Yuanye Bio-Technology Co., Ltd.

### Genome sequencing, assembly, and chromosome-scale scaffolding

Short-read libraries were sequenced on an Illumina NovaSeq platform. After filtering, 68.93 Gb clean Illumina data were obtained, with a Q30 value of 93.49%. K-mer analysis estimated a genome size of 927.61 Mb. Long-read sequencing was performed on an Oxford Nanopore PromethION platform. A total of 65.12 Gb filtered Nanopore reads were generated, about 70x coverage based on the estimated genome size. The final genome assembly was 924.44 Mb in 923 contigs (GC 37.25%), with a contig N50 of 2.07 Mb and a maximum contig length of 8.74 Mb. Hi-C scaffolding (ALLHiC)^41^ grouped sequences into 13 pseudochromosomes (chr01-chr13), with an additional unanchored set (chrUnK). Assembly accuracy was supported by an Illumina read mapping rate of 99.82%, with an average depth of 73.97x and genome coverage of 92.38%. To improve transcript evidence for genome annotation and downstream expression analyses, PacBio Iso-Seq full-length transcriptome sequencing was performed on a SMRTbell library sequenced using the PacBio Sequel platform^42^. Protein-coding genes were predicted using an evidence-guided approach (BRAKER integrating GeneMark-ET^43^ and AUGUSTUS^44^) trained with transcriptome evidence, yielding 22,226 predicted protein-coding genes. Illumina RNA-seq reads from each sample were aligned to the reference genome using STAR^45^. Gene-level read counts were obtained with HTSeq^46^, and expression levels were estimated using RSEM. Expression values were normalized (e.g., FPKM/TPM as appropriate) for downstream analyses. Differential expression analysis across tissues was performed using DESeq2^47^.

### Gene selection

Co-expression analysis of the entire S. tetrandra gene set was performed using the scipy library (v1.5.3) in Python. Genes exhibiting a Pearson correlation coefficient (r) > 0.7 with the bait gene *St6OMT1* (locus St09G337) were defined as the primary co-expression pool. This dataset was subsequently filtered based on functional annotations to retain candidates annotated as “cytochrome P450”, “methyltransferase”, “berberine bridge enzyme”, or “short-chain dehydrogenase/reductase”. Candidates were further curated by excluding sequences with truncated or aberrant open reading frames (ORFs) to generate the final primary candidate list. Upon functional validation of hits from this primary screen, an iterative homology-based mining strategy was employed: validated enzymes served as queries for local BLAST searches to retrieve paralogs sharing >70% amino acid sequence identity, which constituted a secondary candidate pool for subsequent functional characterization.

### Gene Cloning and Plasmid Construction

Total RNA was isolated from the root cortices and steles of S. tetrandra using an RNA Isolation Kit (Huayueyang Biotechnology, Beijing, China). First-strand cDNA was synthesized using TransScript One-Step gDNA Removal and cDNA Synthesis SuperMix (TransGen Biotech, Beijing, China). Candidate genes were amplified from cDNA using PrimeSTAR® Max DNA Polymerase (Takara Bio, Japan) and gene-specific primers (Supplementary Table 11). Full-length synthetic genes for Bs*CYP80A1*^16^, Nn*CYP80Q2*^17^, and *Bs-MaCYP80A1*^18^ (SinoGenoMax, Beijing, China) served as positive controls. *Bs-MaCYP80A1* is a chimeric construct derived from a *BsCYP80A1* ortholog in *Mahonia aquifolium*^18^. As the initial sequence lacked the N-terminal signal peptide and integral membrane anchor characteristic of plant CYP enzymes, it was completed by fusing the first 86 amino acids from the N-terminus of BsCYP80A1 to its N-terminal end. The resulting recombinant gene was designated Bs-MaCYP80A1. PCR amplicons encoding CYPs and BBE-like enzymes were gel-purified and cloned into the *Bam*HI-linearized pESC-Ura vector (New England Biolabs, USA) via Gibson assembly (TransGen Biotech, China). Similarly, methyltransferase and SDR genes were cloned into the *Bam*HI-linearized pET-32a(+) vector using the same assembly method. Recombinant plasmids were transformed into chemically competent *E. coli* Trans1 T1 cells (TransGen Biotech). Positive clones were identified by colony PCR and verified by Sanger sequencing (Ruibiotech, Beijing, China). Validated plasmids were extracted using the Magen HiPure Plasmid Miniprep Kit (Magen, Guangzhou, China). For *S. cerevisiae* strain engineering, all pathway genes were codon-optimized for yeast expression and chemically synthesized by GenScript (Nanjing, China).

### Heterologous expression of CYPs in yeast

Recombinant pESC-Ura (Agilent) plasmids harboring candidate CYP genes were individually transformed into S. cerevisiae strain WAT11 using the Frozen-EZ Yeast Transformation II Kit (Zymo Research). The WAT11 strain was selected for its engineered co-expression of the Arabidopsis thaliana cytochrome P450 reductase ATR1, which supports efficient plant CYP catalysis^33^. An empty pESC-Ura vector control was generated in parallel. Transformants were selected on synthetic dropout medium lacking uracil (SD-Ura, FunGenome) supplemented with 20 g/L glucose. Validated clones were cultured at 30 °C with shaking until reaching an OD_600_ of 2-3. Cells were then harvested by centrifugation, washed three times with sterile ddH_2_O to deplete residual glucose, and resuspended in induction medium (YPL: 10 g/L yeast extract, 20 g/L peptone, 20 g/L galactose). Protein expression was induced by culturing overnight at 30 °C.

### Microsomes extraction and *in vitro* enzymatic activity assay

Yeast microsomes were prepared according to established protocols^48^, which have been successfully employed for the characterization of plant CYPs^49–51^. Following induction, yeast cells were harvested by centrifugation (6,000 × *g*, 5 min) and conditioned by resuspension in TEK buffer (50 mM Tris-HCl, 1 mM EDTA, 100 mM KCl, pH 7.5; 0.5 *g* wet cells/mL) for 5 min at room temperature. Cells were re-pelleted and resuspended in ice-cold TESB buffer (50 mM Tris-HCl, 1 mM EDTA, 600 mM sorbitol, pH 7.5). Cell lysis was achieved using a high-pressure homogenizer (10 cycles at 1,000 psi, 2-4 °C). The lysate was centrifuged at 20,000 × *g* for 20 min at 4 °C to remove cell debris. Microsomes were precipitated from the supernatant by the addition of NaCl (150 mM final) and PEG-4000 (0.1 *g*/mL final). After incubation on ice for 1 h with periodic mixing, microsomal fractions were pelleted (20,000 × *g*, 20 min, 4 °C) and resuspended in 1 mL TEG buffer (50 mM Tris-HCl, 1 mM EDTA, 20% glycerol, pH 7.5).

In vitro enzymatic assays were conducted in a 500 μL reaction volume containing 100 mM Tris-HCl (pH 7.5), 0.5 mg microsomal protein, 0.2 mM substrate, 500 μM NADPH, and an NADPH-regenerating system (5 mM glucose-6-phosphate, 1 U glucose-6-phosphate dehydrogenase, 5 μM FAD, 5 μM FMN). Reactions were incubated at 30 °C for 3 h with shaking (180 rpm). To quench the reaction and facilitate alkaloid extraction, the pH was adjusted to ∼10 by the addition of 10 μL ammonium hydroxide, followed by extraction with 500 μL ethyl acetate. For coupled assays involving multiple enzymes, the total reaction volume was maintained constant using Tris-HCl buffer. The organic phase was evaporated to dryness, reconstituted in 150 μL methanol, and centrifuged (20,000 × *g*, 15 min) prior to LC-MS analysis.

### Heterologous expression and recombinant protein extraction of BBE-like

Heterologous expression of BBE-like enzymes was performed in the *S. cerevisiae* strain WAT11, following the transformation, cultivation, and induction protocols described above for CYPs. Post-induction, cells were harvested (6,000 × *g*, 5 min) and conditioned in TEK buffer (50 mM Tris-HCl, 1 mM EDTA, 100 mM KCl, pH 7.5; 0.5 *g* wet cells/mL, 5 min, room temperature). The cell pellet was resuspended in one-fifth volume of ice-cold TESB buffer (50 mM Tris-HCl, 1 mM EDTA, 600 mM sorbitol, pH 9) and disrupted using a high-pressure homogenizer (>1,000 psi, 15 cycles, 2-4 °C). The lysate was clarified by centrifugation at 20,000 × *g* for 20 min at 4 °C. The resulting supernatant, containing the soluble recombinant BBE-like protein, was harvested as the crude enzyme fraction. For enzymatic assays, 100 μL of this crude lysate was used per reaction, with the total volume normalized across samples using Tris-HCl buffer.

### Heterologous expression of methyltransferases and SDR in *E. coli*

Recombinant pET-32a(+) plasmids harboring candidate methyltransferase and reductase genes were transformed into E. coli BL21(DE3) (TransGen) for heterologous expression. Transformants were cultured in 100 mL Luria-Bertani (LB) broth supplemented with 100 μg/mL ampicillin at 37 °C with shaking (220 rpm). Upon reaching an OD_600_ of 0.6-0.8, protein expression was induced by the addition of isopropyl-β-D-thiogalactopyranoside (IPTG) to a final concentration of 0.3 mM. Cultures were incubated for 16 h at 16 °C (160 rpm). Cells were harvested by centrifugation (5,000 × *g*, 5 min, 4 °C) and resuspended in 5 mL of 100 mM Tris-HCl buffer (pH 7.5). Prior to lysis, phenylmethylsulfonyl fluoride (PMSF) was added to a final concentration of 1 mM (from a 100 mM stock) to inhibit proteolysis. Cell disruption was performed by sonication on ice (40% amplitude, 2 s on/2 s off, 5 min total). The lysate was clarified by centrifugation at 12,000 × *g* for 20 min, and the supernatant was collected as the crude enzyme fraction.

### *In vitro* enzymatic activity assay of methyltransferases and reductases

Standard in vitro assays were performed in a reaction buffer containing 50 mM Tris-HCl (pH 7.5), 0.02 mM substrate, and 100 μL of crude enzyme lysate. Cofactors were supplemented based on the enzyme class: 0.2 mM S-adenosylmethionine (SAM) for methyltransferase reactions and 0.2 mM NADPH for reductase reactions. For coupled assays involving multiple enzymes (including CYPs with their requisite regenerating systems), 100 μL of each crude lysate was added, and the final volume was normalized across all samples using Tris-HCl buffer. To interrogate the reaction hierarchy of methyltransferases, a sequential workup protocol was employed: products from the initial enzymatic step were extracted, evaporated to dryness, and reconstituted in fresh reaction buffer containing the subsequent enzyme and cofactor. This iterative extraction and reconstitution cycle was repeated for each step. All reactions were incubated at 30 °C for 3 h. Metabolite extraction and sample preparation for LC-MS analysis followed the procedures described above.

### LC-MS analysis of enzymatic reaction products

Chromatographic separation was performed on a Waters Acquity UPLC system equipped with an ACQUITY UPLC HSS T3 column (2.1 × 100 mm, 1.8 μm particle size) maintained at 38 °C. The mobile phase consisted of acetonitrile (A) and 0.1% aqueous formic acid (B) delivered at a flow rate of 0.4 mL/min. The elution gradient was programmed as follows: 5-30% A (0-6.0 min), 30-60% A (6.0-8.0 min), 60-90% A (8.0-8.5 min), 90% A isocratic hold (8.5-9.5 min), 90-5% A (9.5-10.0 min), and 5% A re-equilibration (10.0-11.0 min).

Mass spectrometry was conducted on a Waters Xevo G2-S Q-TOF mass spectrometer equipped with an electrospray ionization (ESI) source operating in positive ion mode. Data were acquired over a mass range of *m*/*z* 50-1500. Optimized source parameters were: capillary voltage, 0.5 kV; sample cone voltage, 40 V; extraction cone voltage, 4 V; source temperature, 100 °C; desolvation temperature, 300 °C; and desolvation gas flow, 800 L/h. MS/E acquisition was performed with a low-energy trap collision energy of 6 eV and a high-energy ramp of 30-50 eV^52^. Data acquisition and processing were controlled by MassLynx software (Waters).

### Stereochemical analysis via Electronic Circular Dichroism (ECD) and Chiral HPLC

The absolute configurations of the chemically synthesized lindoldhamine **8** stereoisomers were determined by comparing experimental ECD spectra, acquired on a JASCO J-815 spectropolarimeter (JASCO, Easton, MD, USA), with theoretically calculated spectra. This validated computational ECD workflow was subsequently applied to establish the absolute stereochemistry of the BBR-catalyzed reduction product (*S*,*S*)-**17**. For the stereochemical assignment of the CYP80Q4 product, the enzymatic (*R*,*S*)-**8** (derived from the coupling of (*R*)-**4** and (*S*)-**4**) was analyzed against a panel of four synthetic lindoldhamine diastereomers using chiral HPLC. Separation was achieved on a CHIRALPAK IJ column (4.6 × 250 mm, 5 μm; Daicel Chiral Technologies) maintained at 35 °C. The mobile phase consisted of an isocratic mixture of methanol/acetonitrile/diethylamine (95:5:0.1, v/v/v) delivered at 1 mL/min. Chromatograms were recorded at 282 nm using an Agilent 1260 Infinity II HPLC system equipped with a diode array detector.

### Computational electron circular dichroism analysis

Conformational analyses were carried out via Monte Carlo searching using molecular mechanics with MMFF force field in the Spartan′18 program. The obtained conformers were reoptimized using DFT at the B3LYP/6-31G(d) level in gas phase using the Gaussian 09 program, and selected relative Gibbs free energies in the range of 0-2 kcal/mol were chosen for subsequent calculations. The B3LYP/6-31G(d) harmonic vibrational frequencies were further calculated to confirm their stability. The energies, oscillator strengths, and rotational strengths of the first 60 electronic excitations were calculated using the TDDFT methodology at the M062X/TZVP level. The ECD spectra were simulated by the overlapping Gaussian function^53^.

To get the final ECD spectra, the simulated spectra of the lowest energy conformers were averaged according to the Boltzmann distribution theory and their relative Gibbs free energy (ΔG).

### Isolation of enzymatic reaction products and nuclear magnetic resonance analysis

The reaction system was scaled up to 200 mL, with products extracted using the established method^54^. In multistep enzymatic reaction systems conducted at scale, buffer volumes were adjusted during microsomal disruption or dissolution to increase enzyme concentration prior to each reaction phase, ensuring the final volume consistently approximated 200 mL. The completely evaporated products were dissolved in 2 mL methanol and purified via semi-preparative HPLC equipped with an MWD UV detector (SEP LC-52 system, Separation Technologies). Separation employed a CSH Prep C18 column (10 × 250 mm, 5 μm; Waters) using a mobile phase of acetonitrile and 0.1% aqueous formic acid at 2 mL/min. A linear gradient progressed from 10% to 20% acetonitrile between 1-20 min, with UV monitoring at 282 nm. Structural characterization was performed through ¹H NMR, ¹³C NMR, and 2D NMR spectroscopy (Bruker DRX Avance 500/600/800 MHz). Chemical shifts are reported in parts per million (ppm) relative to tetramethylsilane. All NMR spectra are shown in Supplementary Fig. 30 to 67.

### Strains, Media, and Reagents for Metabolic Engineering

All strains and plasmids used in this study are listed in Supplementary Table 10. *E. coli* DH5α strain was used for routine plasmid assembly. The norcoclaurine-producing platform yeast XJ0636^24^, constructed in our prior study, was employed as starting strain to engineer for bisBIAs biosynthesis.

YPD medium (20 g/L peptone (Difco), 10 g/L yeast extract (Merck Millipore), and 20 g/L glucose (VWR)) was used for routine yeast cultivation and competent cells preparation. Synthetic dropout minus uracil medium, consisting of 6.7 g/L yeast nitrogen base (YNB) without amino acids, (Formedium), 0.77 g/L complete supplement mixture without uracil (CSM-URA, Formedium) and 20 g/L glucose, was used for screening positive colonies transformed with *URA3* marker plasmid. CSM + 5-FOA medium, containing 6.7 g/L YNB without amino acids, 0.79 g/L complete supplement mixture (CSM, Formedium) and 0.8 g/L 5-fluoroorotic acid (5-FOA, Sigma) was used for recycling the URA3 marker. If necessary for preparing plates, 20 g/L agar (Merck Millipore) was added. Delft medium (7.5 g/L (NH_4_)_2_SO_4_, 14.4 g/L KH_2_PO_4_, 0.5 g/L MgSO_4_·7H_2_O, 20 g/L glucose, 2 ml/L trace metal solutions and 1 ml/L vitamin solutions ^55^, supplemented with 60 mg/L uracil if required) was used for shake flask fermentation for producing bisBIAs.

Gibson Assembly Kit and Golden Gate Assembly Kit were obtained from NEB. PrimeSTAR HS polymerase and Phusion polymerase were purchased from Takara and Thermo Fisher Scientific, respectively.

### Genetic manipulations

All primers used in this work are listed in Supplementary Table 11 and 12. Since starting strain XJ0636 carried chromosomally integrated *Cas9* gene under *TEF1* promoter, the CRISPR-Cas9 based genome editing method was used for genetic manipulations as described before^56^. All promoters and terminators were amplified from XJ0636 genomic DNA by using PrimeSTAR HS or Phusion polymerase. All primers were synthesized from Eurofins or Integrated DNA Technologies. The heterologous genes listed in Supplementary Information were codon-optimized synthesized by GenScript. For assembling repair fragment, up-stream homologous arm, promoter, gene, terminator, and down-stream homologous arm were first amplified with primers carrying 5′-terminal 30-40 bp overhangs, respectively. The entire gene repair fragment was assembled by overlap PCR method^57^. High-efficiency yeast transformation was performed by using the LiAc/SS carrier DNA/PEG method^58^.

### Culture of engineered yeast and its metabolite detection

Three or more biologically independent colonies were inoculated into 1 mL Delft medium supplemented with 60 mg/L uracil within 14 mL culture tubes for 18 h at 30 °C, 220 rpm. The preculture was then transferred into 20 mL fresh Delft medium supplemented with 60 mg/L in shake flasks at an initial OD_600_ of 0.05 for 72 h. After fermentation, metabolites were extracted from culture. Briefly, 500 μL of culture was mixed with 500 μL of 30% acetonitrile (ACN), vortexed thoroughly, and then centrifuged for 5 min at 13,000 × *g*. If necessary, the supernatant was further diluted with 15% ACN to the suitable concentration for LC-MS/MS analysis. Samples were then stored at −20 °C until analysis.

One microliter of each sample from yeast fermentation was injected for analysis by LC-MS/MS system, consisting of a Shimadzu Nexera UHPLC system and a high-end hybrid triple quadrupole ion trap instrument (Sciex QTRAP 6500+) with Luna Omega 1.6 μm Polar C18 100 Å column (Phenomenex). Analytes were eluted at a constant flow rate of 400 μL min^-1^ with 0.1% formic acid as solvent A and ACN with 0.1% formic acid as solvent B. Analyte bisBIAs were separated with the following gradient method: 10% B from 0-0.1 min, 10% B to 45% B from 0.1-5 min, 45% B to 90% B from 5-5.5 min, held at 90% B from 5.5-7 min, 90% B to 10% B from 7-7.01 min, and held at 10% B from 7.01-9.5 min. For quantification of isotetrandrine in engineered strains, MS2 mode was used with detailed parameters (Product of 623.0 Da; from *m*/*z* 100 to 650; CE from 10 V to 47 V; DP 130 V; EP 10 V; CXP 17 V).

(*S*,*S*)-**17** and (*R*,*S*)-**17** analysis was performed by using a Waters Xevo G2-X QTOF system with an electrospray ionization source operating in positive mode. Full scanning was conducted over an *m*/*z* range of 100-800. The scanning time was 0.5 s, the detection time was 30 min, the low energy impact voltage was 6 V, and the high energy impact voltage was 15-40 V. Nitrogen gas was used as the solvent gas, and leucine enkephalin was used for real-time correction. All the target compounds were eluted at a constant flow rate of 300 µL/min by using column HSS T3 Column, 100 Å, 1.8 µm, 2.1 mm × 100 mm, with 0.04% formic acid as solvent A and methanol with 0.04% formic acid as solvent B. The gradient method was used as follows: 5% B to 10% B from 0-5.0 min, 10% B to 15% B from 5.0-20.0 min, 15% B to 25% B from 20.0- 23.0 min, 25% B to 40% B from 23.0-26.0 min, 40% B to 90% B from 26.0-27.0 min, held at 90% B from 27.0-28.5 min, 90% B to 5% B from 28.5-28.6 min, and held at 5% B from 28.6-30 min. MassLynx software was used for data acquisition and processing.

