## Supplementary Information for "Biosynthesis of bisbenzylisoquinoline alkaloids"

**The PDF file includes:**

Supplementary Figs. 1 to 29
Supplementary Tables 1 to 12
Synthesis of compounds
Compound structure analysis and assignment
The sequences of the genes identified in this work
Supplementary Figs. 30 to 67: NMR Spectrum
References

### Supplementary Figures

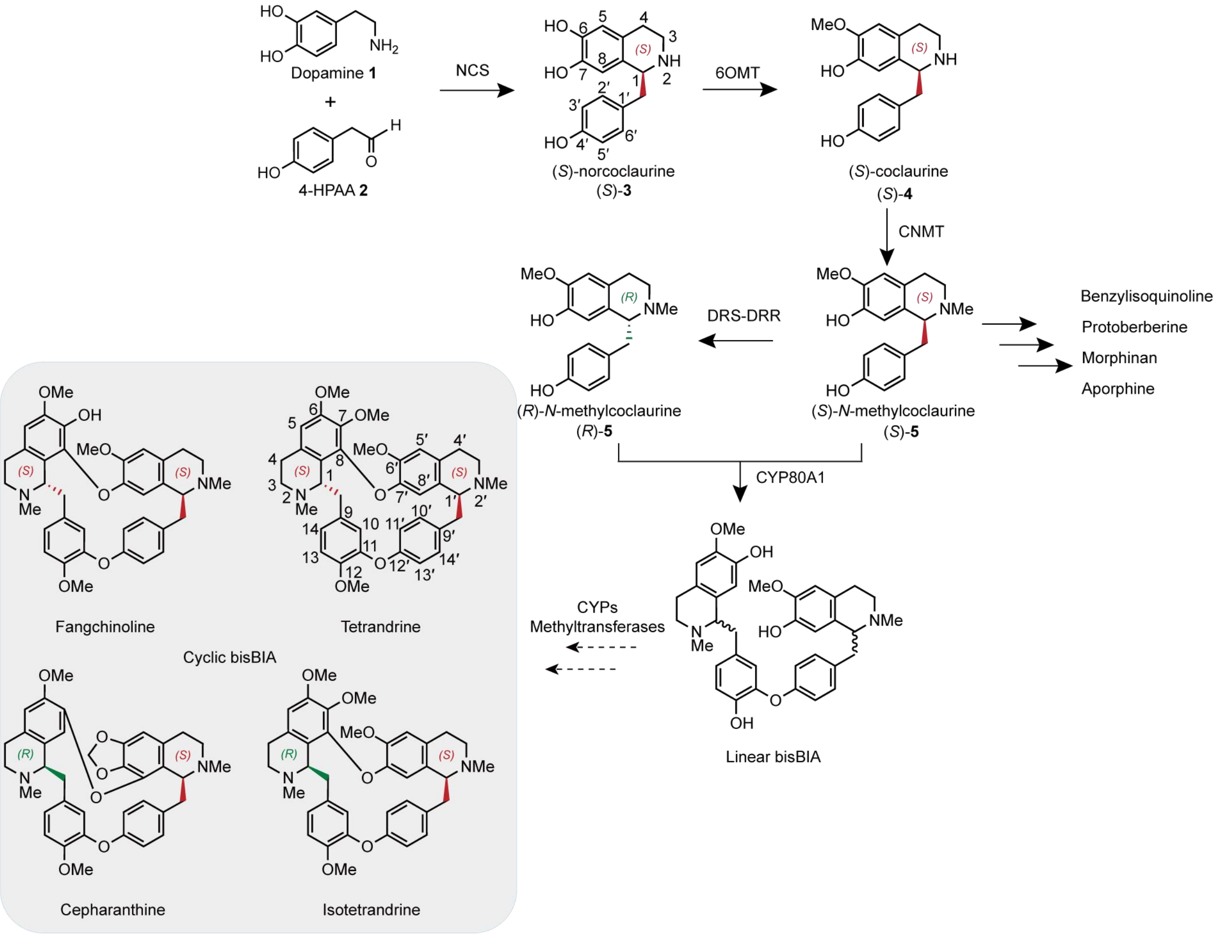


**Supplementary Fig. 1. Previously proposed biosynthetic pathway for bisBIAs [1].**
Dashed lines represent unknown steps in the bisBIA biosynthetic pathway. Common bioactive cyclic bisBIAs found in *S. tetrandra* are shown within the gray background [2]. 4-HPAA, 4-hydroxyphenylacetaldehyde; NCS, norcoclaurine synthase; 6OMT, norcoclaurine 6-*O*-methyltransferase; CNMT, coclaurine *N*-methyltransferase; DRS-DRR, dehydrococlaurine synthase-dehydrococlaurine reductase; CYP, cytochrome P450.

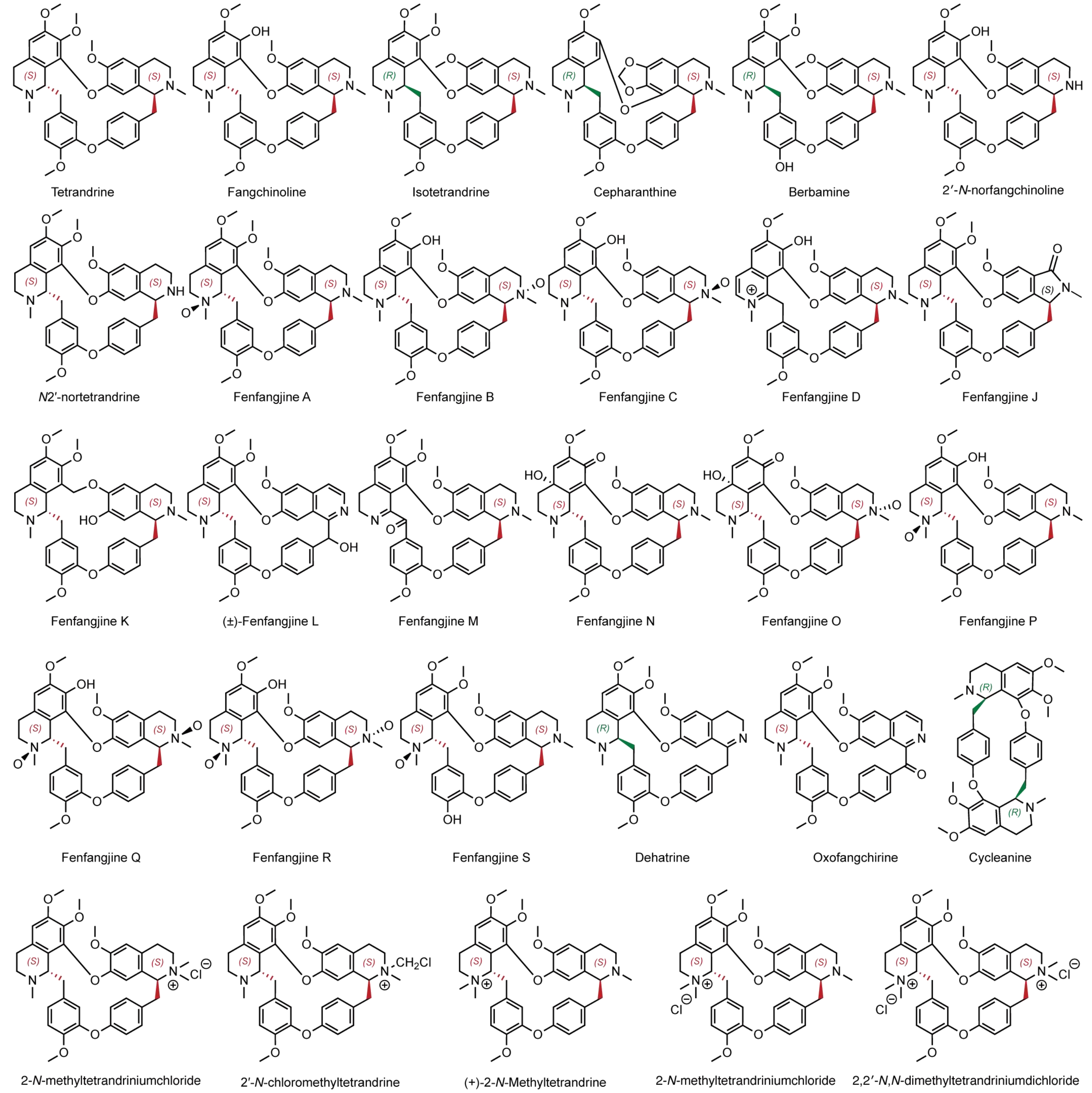


**Supplementary Fig. 2. Structures of known bisBIAs isolated and identified from *S. tetrandra*[2].** The structural diversity of bisBIA arises from a combination of factors, including different coupling modes, variations in coupling sites, linkages between 1-BIA stereoisomers of different configurations, and diverse methylation patterns.

a

b

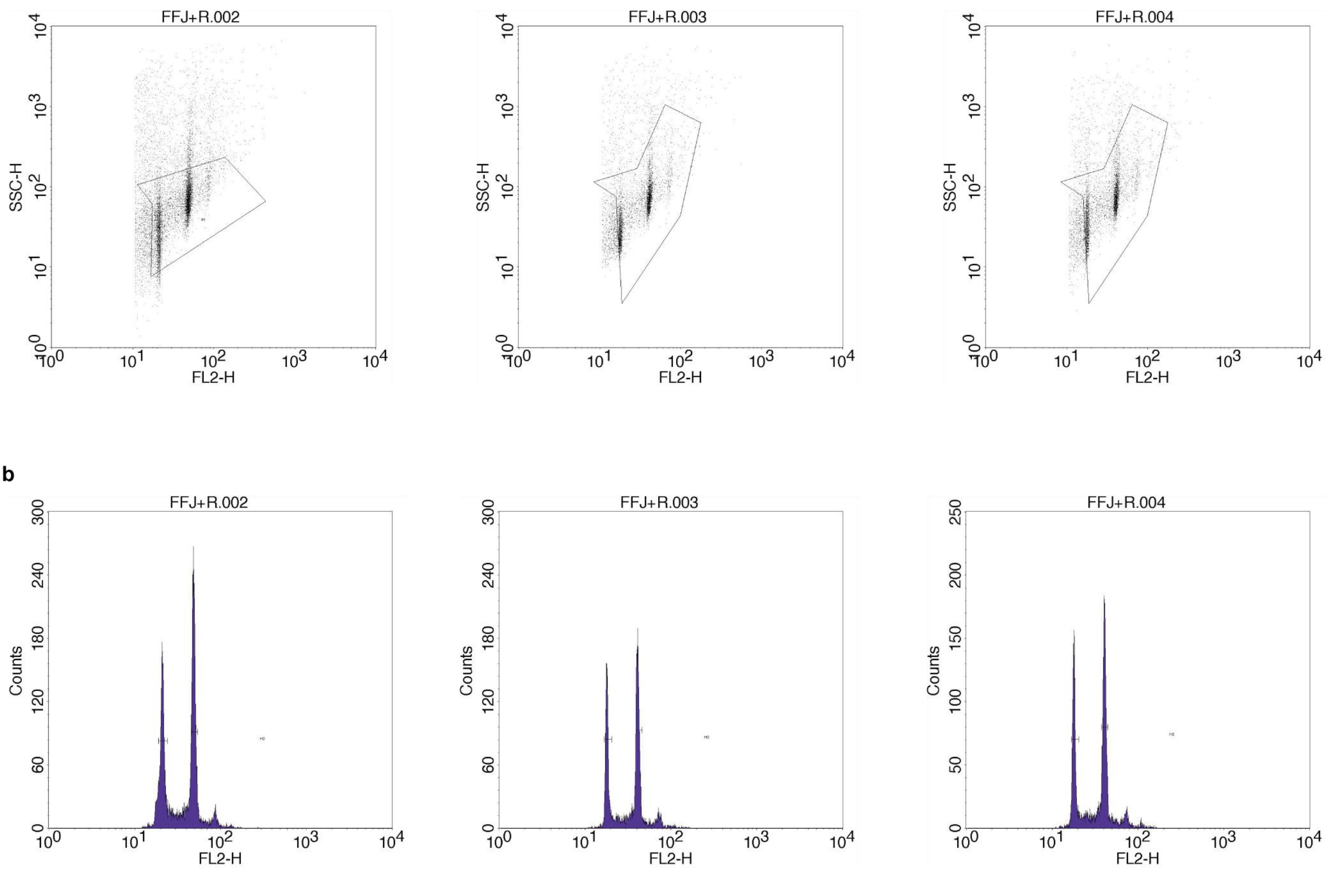


c

| Sample | Reference | Ratio (Sample/Ref) | Estimated Genome Size (Mb) |
|---|---|---|---|
| *Stephania tetrandra* | *Oryza sativa* (420 Mb) | 2.28 ± 0.02 | 959.07 ± 8.85 |

**Supplementary Fig. 3. Estimation of the genome size of *Stephania tetrandra* by flow cytometry. a**, Scatter plots of side scatter (SSC-H) versus fluorescence intensity (FL2-H) for nuclei isolated from *S. tetrandra* and the internal reference. **b**, Flow cytometric histograms showing the fluorescence peaks of *S. tetrandra* and *Oryza sativa* cv. Nipponbare. The x-axis represents the relative DNA content, and the y-axis represents the number of nuclei. **c**, Estimation of *S. tetrandra* genome size estimated based on the ratio of the mean fluorescence intensity between the sample and the internal standard in flow cytometry.

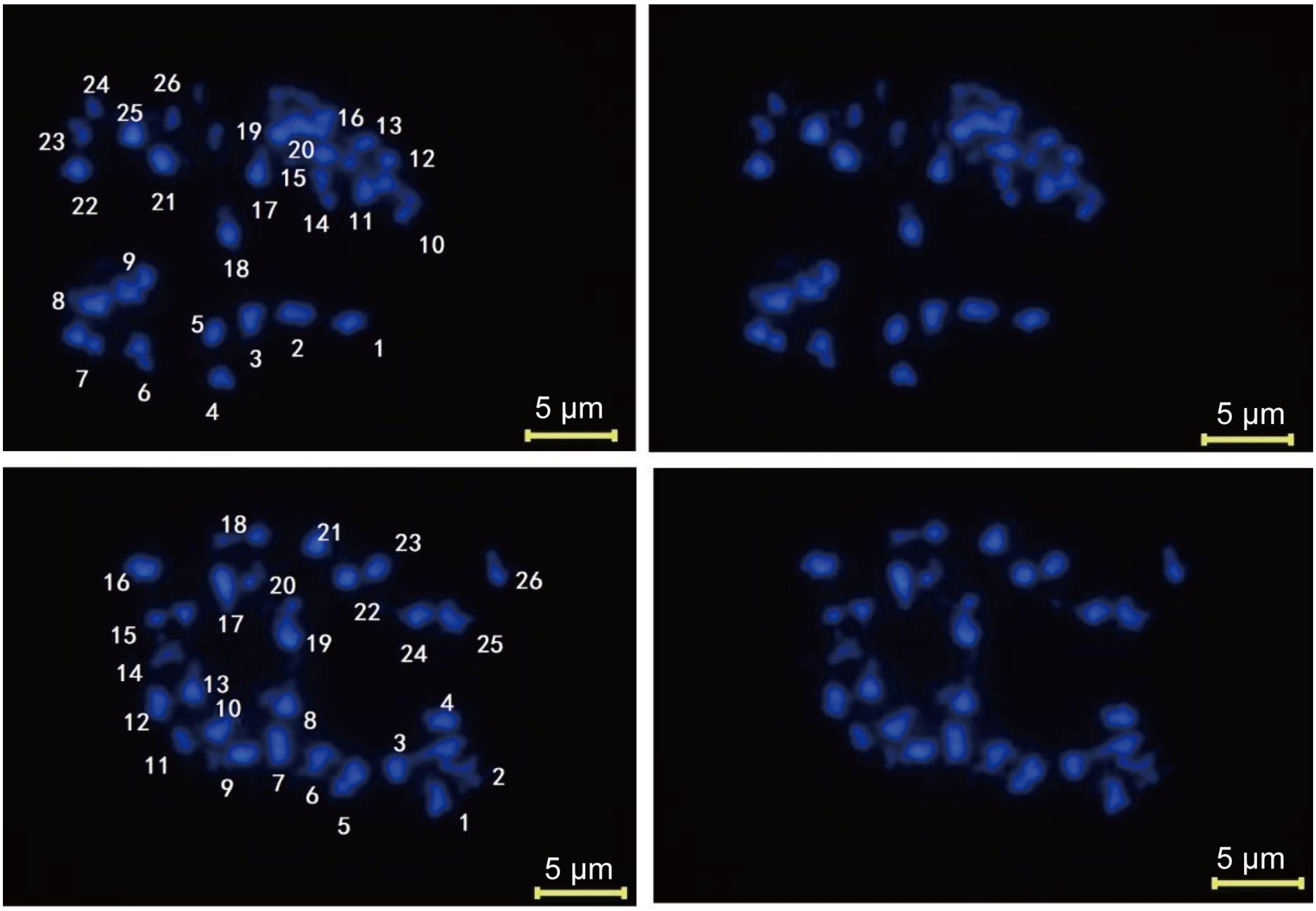


**Supplementary Fig. 4. Karyotype analysis of *S. tetrandra*.** Chromosomes were fluorescently stained using DAPI. Root-tip tissue samples of *S. tetrandra* were sampled and used to infer that *S. tetrandra* has 2n = 2x = 26 chromosomes.

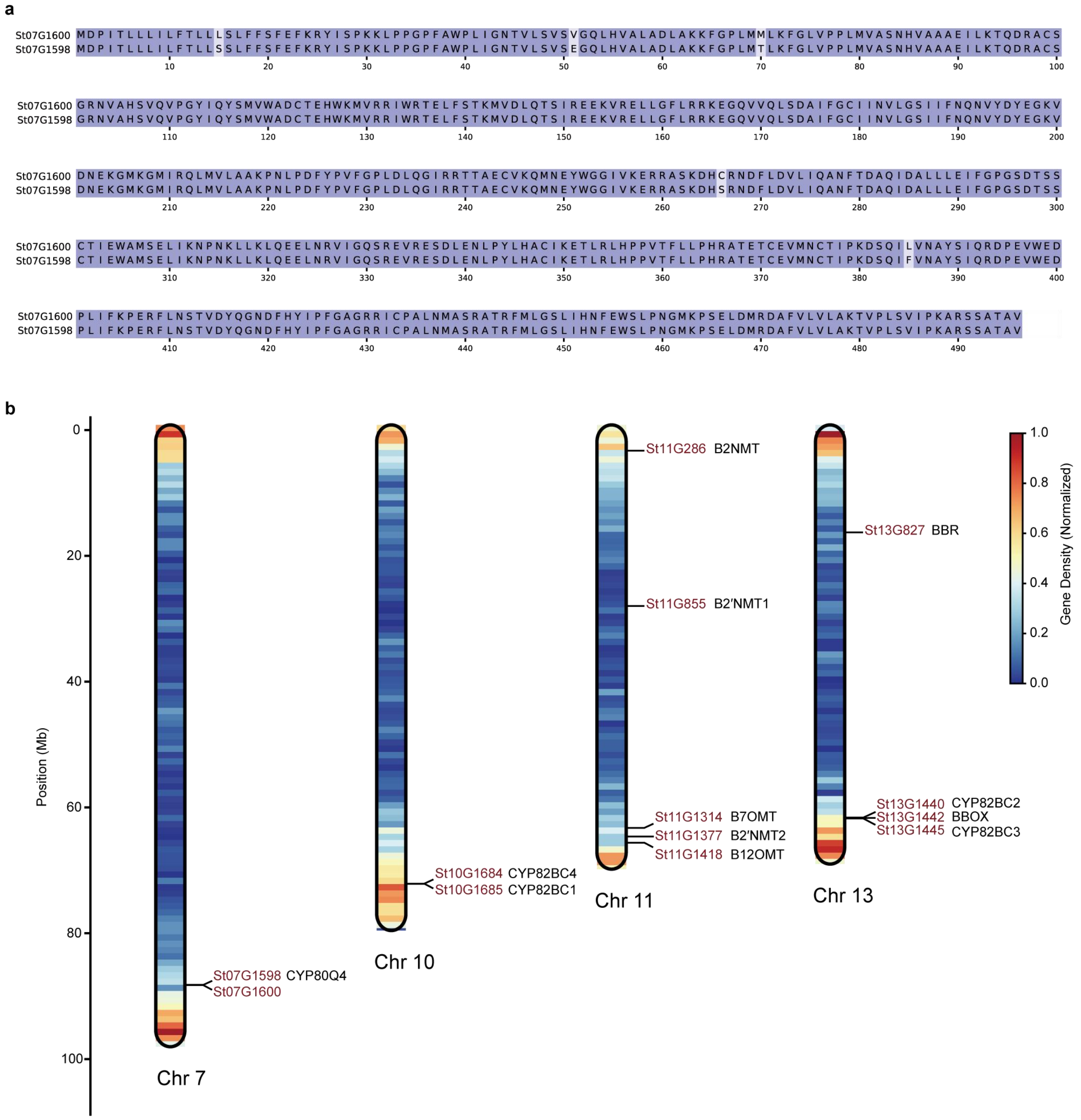


**Supplementary Fig. 5. Genomic organization and clustering of bisBIA biosynthetic genes in the *S. tetrandra* genome.** **a**, Pairwise protein sequence alignment of St07G1598 and St07G1600. These two CYP80 paralogs share 99% amino acid identity, suggesting a recent tandem duplication event within the *S. tetrandra*. **b**, Physical map illustrating the chromosomal distribution of identified bisBIA biosynthetic genes. The heat map along the chromosomes represents normalized gene density across the genome (scale bar, right). Chromosome 7: *St07G1598* and *St07G1600* are organized in a tandem arrangement, consistent with their high sequence homology shown in **a**. Chromosome 10: *CYP82BC1* and *CYP82BC4* reside within a tandem duplication array, reflecting the expansion of the CYP82 family in this species. Chromosome 13: A specialized metabolic gene cluster is identified on chromosome 13, harboring *CYP82BC2*, *CYP82BC3*, and the *BBOX* in close physical proximity. *BBR* is also localized on chromosome 13. Chromosome 11: Three methyltransferase genes — *B7OMT*, *B2'NMT2*, and *B12OMT* — are co-localized in a genomic cluster, suggesting coordinated transcriptional regulation of the early stages of alkaloid modification. *B2NMT* and *B2'NMT1* are also situated on chromosome 11, although they are located at a significant distance from the primary cluster.

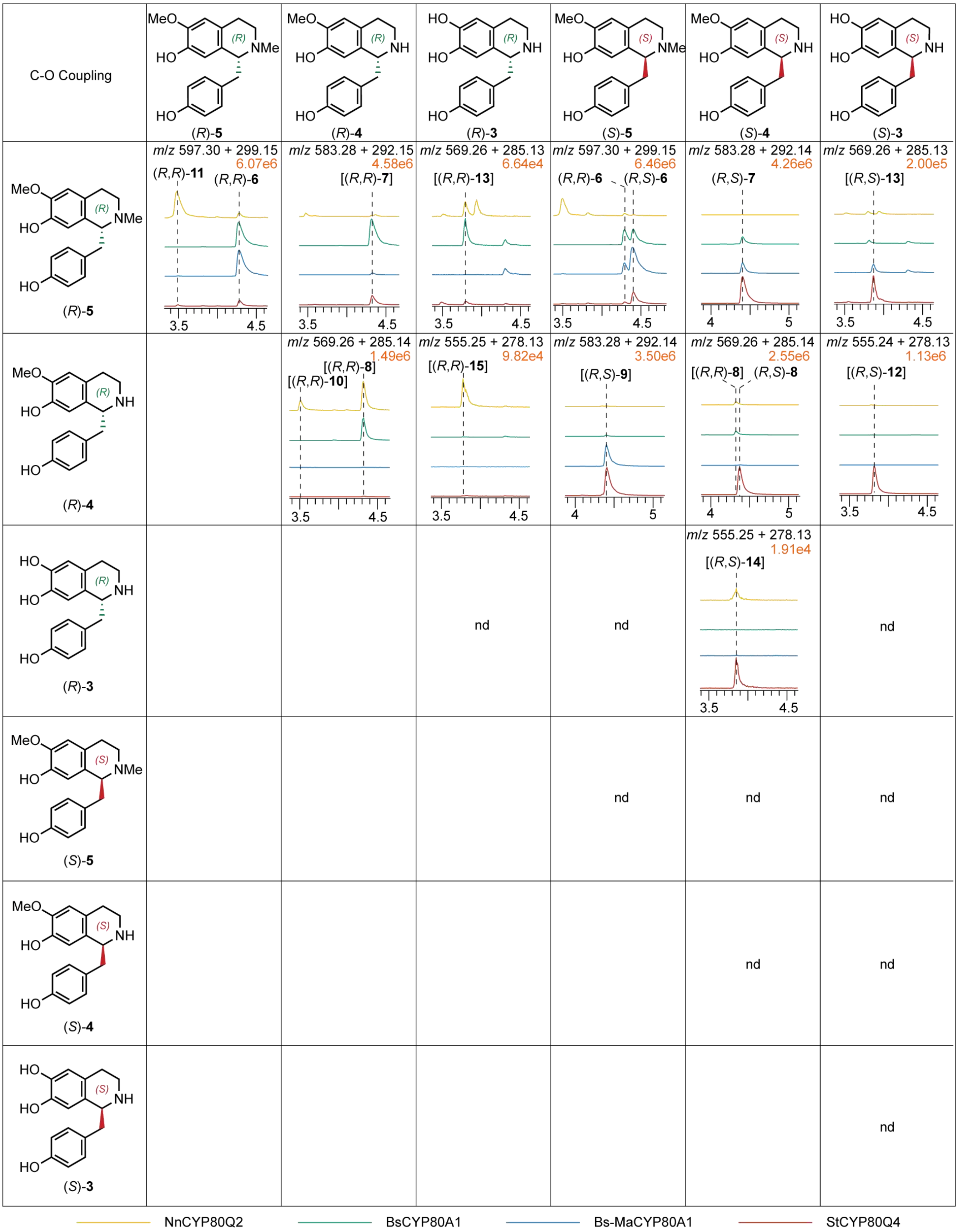


**Supplementary Fig. 6. Catalytic promiscuity of CYP80 bisBIA synthases.** Six 1-BIA precursors—the (*R*)- and (*S*)-enantiomers of norcoclaurine (**3**), coclaurine (**4**), and *N*-methylcoclaurine (**5**)—were evaluated in 21 pairwise combinations to assess the catalytic promiscuity of four bisBIA synthases (StCYP80Q4, BsCYP80A1, Bs-MaCYP80A1, and NnCYP80Q2)[3–5]. LC–MS analysis revealed that active coupling required at least one (*R*)-configured precursor (i.e., (*R*)-**3**, (*R*)-**4**, or (*R*)-**5**). In assays using heteroenantiomeric pairs, the reactions frequently yielded both the expected (*R*,*S*)-heterodimers and the (*R*,*R*)-homodimers (arising from (*R*)-monomer self-coupling), manifesting as dual product peaks in the corresponding chromatograms. As previously reported, utilizing (*R*)-5 as the sole substrate, NnCYP80Q2 exclusively catalyzes the formation of the "head-to-tail" coupled nelumboferine ((*R*,*R*)-11)[4], whereas BsCYP80A1 solely yields the "tail-to-tail" coupled guattegaumerine ((*R*,*R*)-6)[3]. Our profiling revealed that NnCYP80Q2 also produces minor amounts of (*R*,*R*)-6. Furthermore, StCYP80Q4 concurrently generates both (*R*,*R*)-6 and (*R*,*R*)-11, as confirmed by direct comparison with the products of the aforementioned enzymes. Details for the reactions yielding (*R*,*S*)-6, (*R*,*S*)-7, (*R*,*S*)-8, and (*R*,*S*)-9, as discussed in the main text, are provided in Supplementary Figs. 7 and 8. MS/MS spectra for the remaining products are presented in Supplementary Fig. 9. Product names enclosed in square brackets denote tentative assignment as bisBIAs based solely on MS data, pending rigorous structural elucidation. Each cell displays the characteristic dual-ionization $m/z$ values ($[M + H]^+$ and $[M + 2H]^{2+}$) of the detected bisBIAs, and the intensity of the dominant product peak (orange, scientific notation). All extracted ion chromatograms (EICs) are plotted on identical intensity scales; the $x$-axis represents retention time (min).

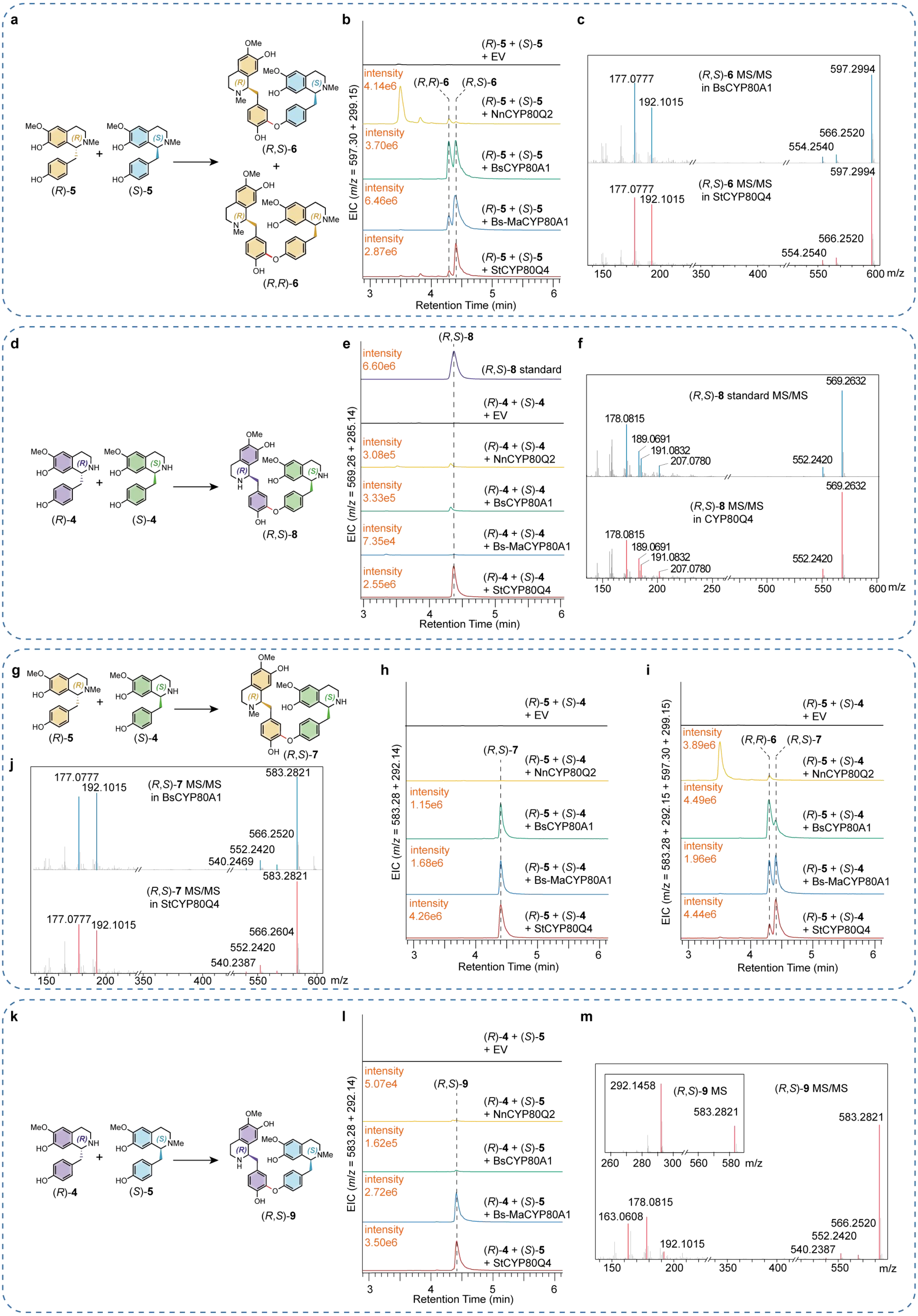
a
(R)-5
(S)-5
(R,S)-6
(R,R)-6
b
(R)-5 + (S)-5
+ EV
intensity
4.14e6
(R,R)-6
(R,S)-6
(R)-5 + (S)-5
+ NnCYP80Q2
intensity
3.70e6
(R)-5 + (S)-5
+ BsCYP80A1
intensity
6.46e6
(R)-5 + (S)-5
+ Bs-MaCYP80A1
intensity
2.87e6
(R)-5 + (S)-5
+ StCYP80Q4
EIC (m/z = 597.30 ± 299.15)
Retention Time (min)
c
177.0777
192.1015
(R,S)-6 MS/MS
in BsCYP80A1
597.2994
566.2520
554.2540
177.0777
192.1015
(R,S)-6 MS/MS
in StCYP80Q4
597.2994
566.2520
554.2540
m/z
d
(R)-4
(S)-4
(R,S)-8
e
(R,S)-8
intensity
6.60e6
(R,S)-8 standard
(R)-4 + (S)-4
+ EV
intensity
3.08e5
(R)-4 + (S)-4
+ NnCYP80Q2
intensity
3.33e5
(R)-4 + (S)-4
+ BsCYP80A1
intensity
7.35e4
(R)-4 + (S)-4
+ Bs-MaCYP80A1
intensity
2.55e6
(R)-4 + (S)-4
+ StCYP80Q4
EIC (m/z = 569.26 ± 285.14)
Retention Time (min)
f
569.2632
(R,S)-8 standard MS/MS
178.0815
189.0691
191.0832
207.0780
552.2420
569.2632
(R,S)-8 MS/MS
in CYP80Q4
178.0815
189.0691
191.0832
207.0780
552.2420
m/z
g
(R)-5
(S)-4
(R,S)-7
h
(R)-5 + (S)-4
+ EV
(R,S)-7
(R)-5 + (S)-4
+ NnCYP80Q2
intensity
1.15e6
(R)-5 + (S)-4
+ BsCYP80A1
intensity
1.68e6
(R)-5 + (S)-4
+ Bs-MaCYP80A1
intensity
4.26e6
(R)-5 + (S)-4
+ StCYP80Q4
EIC (m/z = 583.28 ± 292.14)
Retention Time (min)
i
(R)-5 + (S)-4
+ EV
intensity
3.89e6
(R,R)-6
(R,S)-7
(R)-5 + (S)-4
+ NnCYP80Q2
intensity
4.49e6
(R)-5 + (S)-4
+ BsCYP80A1
intensity
1.96e6
(R)-5 + (S)-4
+ Bs-MaCYP80A1
intensity
4.44e6
(R)-5 + (S)-4
+ StCYP80Q4
EIC (m/z = 583.28 ± 292.15 + 597.30 ± 299.15)
Retention Time (min)
j
177.0777
192.1015
(R,S)-7 MS/MS
in BsCYP80A1
583.2821
566.2520
552.2420
540.2469
583.2821
(R,S)-7 MS/MS
in StCYP80Q4
177.0777
192.1015
566.2604
552.2420
540.2387
m/z
k
(R)-4
(S)-5
(R,S)-9
l
(R)-4 + (S)-5
+ EV
intensity
5.07e4
(R,S)-9
(R)-4 + (S)-5
+ NnCYP80Q2
intensity
1.62e5
(R)-4 + (S)-5
+ BsCYP80A1
intensity
2.72e6
(R)-4 + (S)-5
+ Bs-MaCYP80A1
intensity
3.50e6
(R)-4 + (S)-5
+ StCYP80Q4
EIC (m/z = 583.28 ± 292.14)
Retention Time (min)
m
292.1458
(R,S)-9 MS
583.2821
(R,S)-9 MS/MS
583.2821
178.0815
163.0608
192.1015
566.2520
552.2420
540.2387
m/z

**Supplementary Fig. 7. Functional characterization of CYP80Q4 and its comparison with BsCYP80A1, Bs-MaCYP80A1, and NnCYP80Q2 using (*R*)- and (*S*)-stereoisomers of 4 and 5 as substrates.** *In vitro* enzyme assays using (*R*)- and (*S*)-configured of **4** and **5** as substrates demonstrate the high catalytic activity and distinct substrate specificity of CYP80Q4 in dimerization reactions compared to related CYPs. For each reaction set (enclosed in a blue dashed box), the figure displays the EICs, the MS/MS spectrum of the product(s), and a reaction schematic. **a-c**, With (*R*)-**5** and (*S*)-**5** as substrates, CYP80Q4, BsCYP80A1, and Bs-MaCYP80A1 all catalyzed the formation of both (*R,R*)-**6** and (*R,S*)-**6**. CYP80Q4 and Bs-MaCYP80A1 showed a clear preference for producing (*R,S*)-**6**. The overall product yield for CYP80Q4 was relatively low in this specific reaction. **d-f**, With (*R*)-**4** and (*S*)-**4** as substrates, only CYP80Q4 was capable of catalyzing the formation of the coupling product (*R,S*)-**8**. **g-j**, Catalytic activity assay using (*R*)-**5** and (*S*)-**4** as substrates. EICs show the formation of the coupling product (*R,S*)-**7** and the homo-dimerization product (*R,R*)-**6** from (*R*)-**5**. CYP80Q4 displayed the highest activity for the coupling of (*R*)-**5** and (*S*)-**4**, a reaction not catalyzed by NnCYP80Q2. **k-m**, With (*R*)-**4** and (*S*)-**5** as substrates, CYP80Q4 and Bs-MaCYP80A1 catalyzed the formation of a coupling product, which was tentatively identified as (*R,S*)-**9** based on MS/MS analysis.

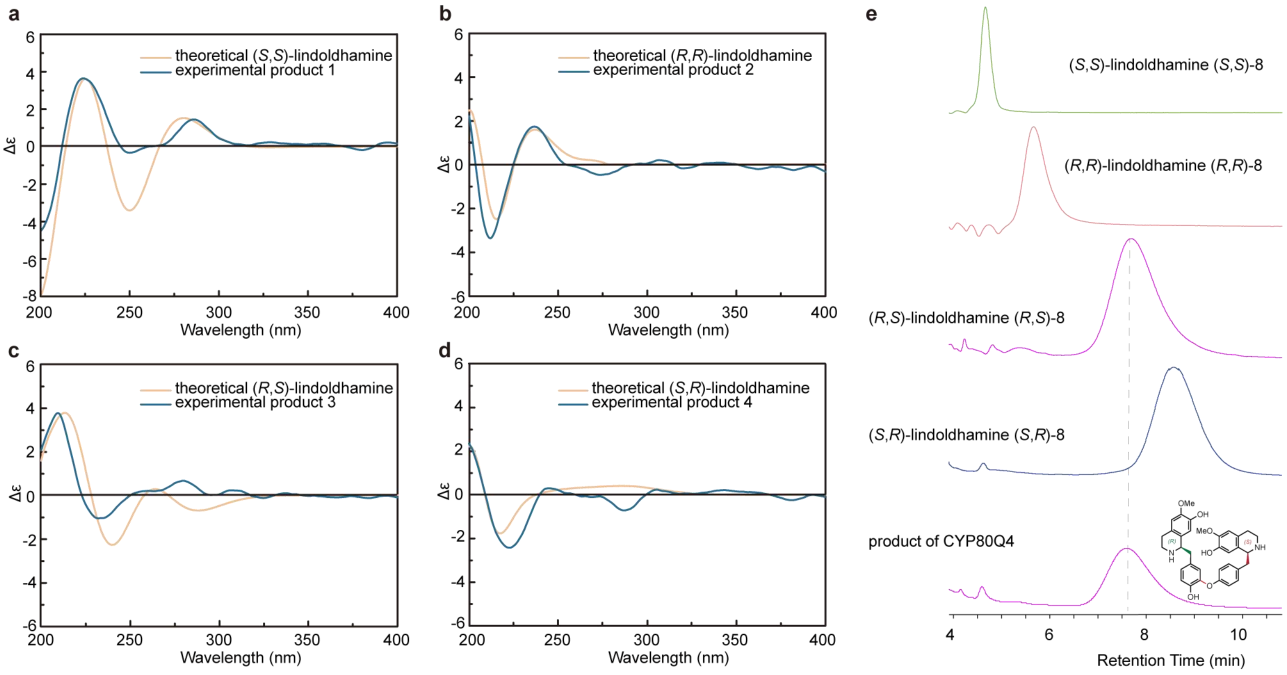


**Supplementary Fig. 8. Stereochemical assignment of (*R*,*S*)-8 via electronic circular dichroism (ECD) calculations and chiral analysis. a-d**, Determination of the absolute configurations of the four **8** stereoisomers. The synthetic isomers were separated by chiral chromatography. The experimental circular dichroism (CD) spectrum of each isomer was compared with its corresponding theoretically calculated ECD spectrum, enabling the assignment of their structures as (*R*,*R*)-, (*S*,*S*)-, (*R*,*S*)-, and (*S*,*R*)-**8**. **e**, Confirmation of the enzymatic product's identity. The product from the CYP80Q4-catalyzed reaction of (*R*)-**4** and (*S*)-**4** was co-injected with the four authentic lindoldhamine standards for chiral HPLC analysis. The enzymatic product co-eluted with the (*R*,*S*)-**8** standard, confirming its absolute configuration.

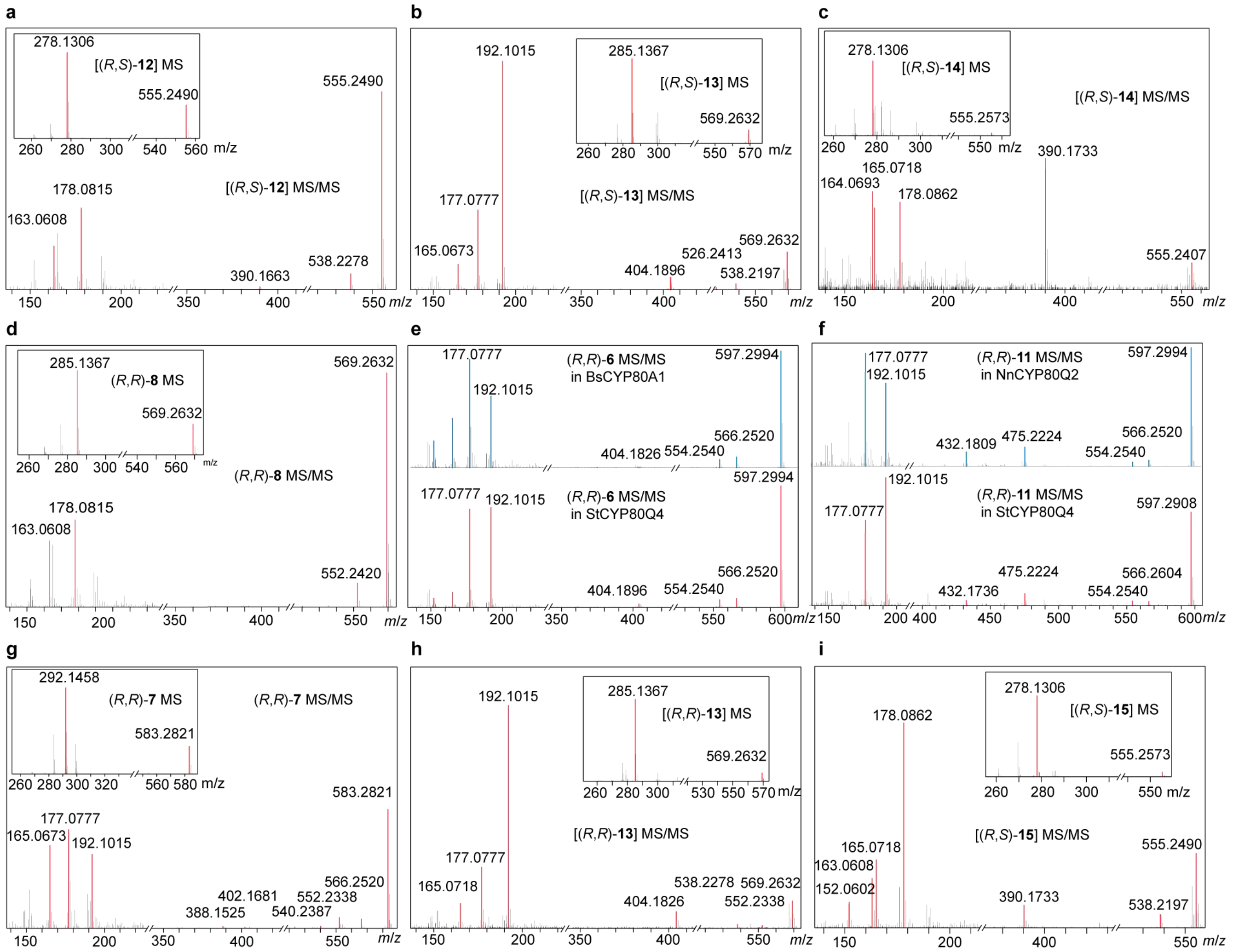


**Supplementary Fig. 9. MS/MS spectra of several bisBIA products generated via CYP80Q4-catalyzed coupling.** These spectra correspond to the enzymatic products derived from the specific substrate combinations evaluated in Supplementary Fig. 6: **a**, (*R*)-**4** and (*S*)-**3**; **b**, (*R*)-**5** and (*S*)-**3**; **c**, (*R*)-**3** and (*S*)-**4**; **d**, (*R*)-**4** and (*R*)-**4**; **e**, f, (*R*)-**5** and (*R*)-**5**; **g**, (*R*)-**5** and (*R*)-**4**; **h**, (*R*)-**5** and (*R*)-**3**; and **i**, (*R*)-**4** and (*R*)-**3**.

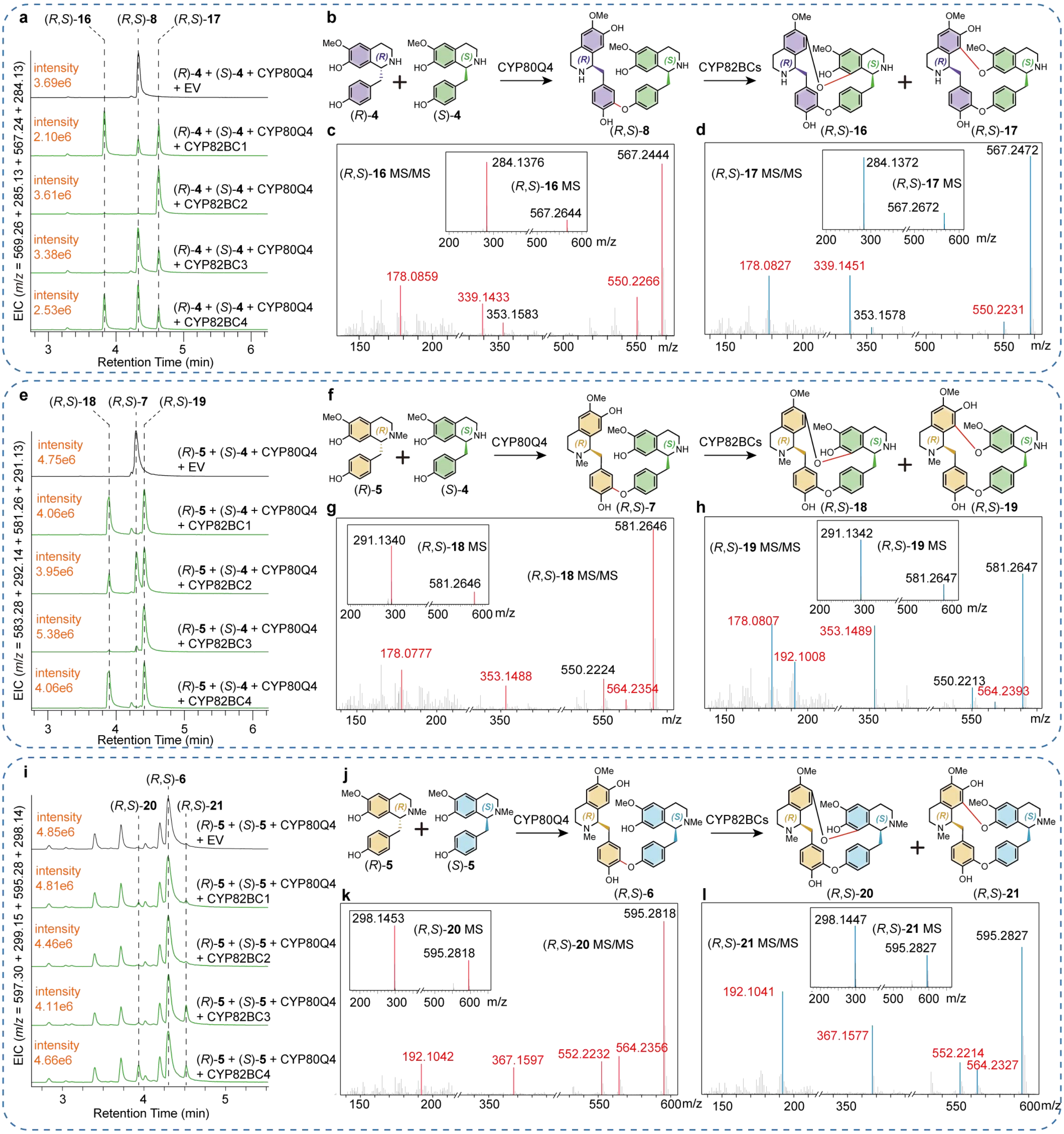


**Supplementary Fig. 10. Functional characterization of CYP82BC bisBIA-cyclizing enzymes.** The functions of four CYP82BCs were assayed in one-pot cascade reactions with CYP80Q4. Four different substrate pairs—(*R*)-**4**/(*S*)-**4**, (*R*)-**5**/(*S*)-**4**, (*R*)-**5**/(*S*)-**5**, and (*R*)-**4**/(*S*)-**5**—were evaluated; notably, the (*R*)-**4** and (*S*)-**5** pair failed to yield any cyclized products. In each cascade, CYP80Q4 initially catalyzes an intermolecular C–O coupling, and the resulting linear dimer is then further cyclized by a CYP82BC. For each reaction set (enclosed in a blue dashed box), the figure displays the EICs, MS/MS spectra of the final products, and a schematic of the reaction. Fragments highlighted in red font indicate the diagnostic ions used for structural assignment. **a-d**, With the (*R*)-**4**/(*S*)-**4** substrate pair, CYP82BCs acted on the CYP80Q4-produced linear intermediate (*R,S*)-**8** to form two isomeric cyclized products (*m/z* 567). The relative yields of these two products varied depending on the specific CYP82BC used. Based on NMR and MS/MS analyses, these final products were identified as (*R,S*)-**16** and (*R,S*)-**17**. **e-h**, With the (*R*)-**5**/(*S*)-**4** substrate pair, CYP82BCs converted the CYP80Q4 product (*R,S*)-**7** into two final products (*m/z* 581), later identified as (*R,S*)-**18** and (*R,S*)-**19** by NMR and MS/MS. The product ratio differed among the four CYP82BC enzymes. **i-l**, With the (*R*)-**5**/(*S*)-**5** substrate pair, CYP82BCs catalyzed the cyclization of the intermediate (*R,S*)-**6** to generate two products, (*R,S*)-**20** and (*R,S*)-**21** (*m*/*z* 595), as confirmed by MS/MS. Catalytic efficiencies for this specific substrate pair were markedly lower compared to the other two productive pairs.

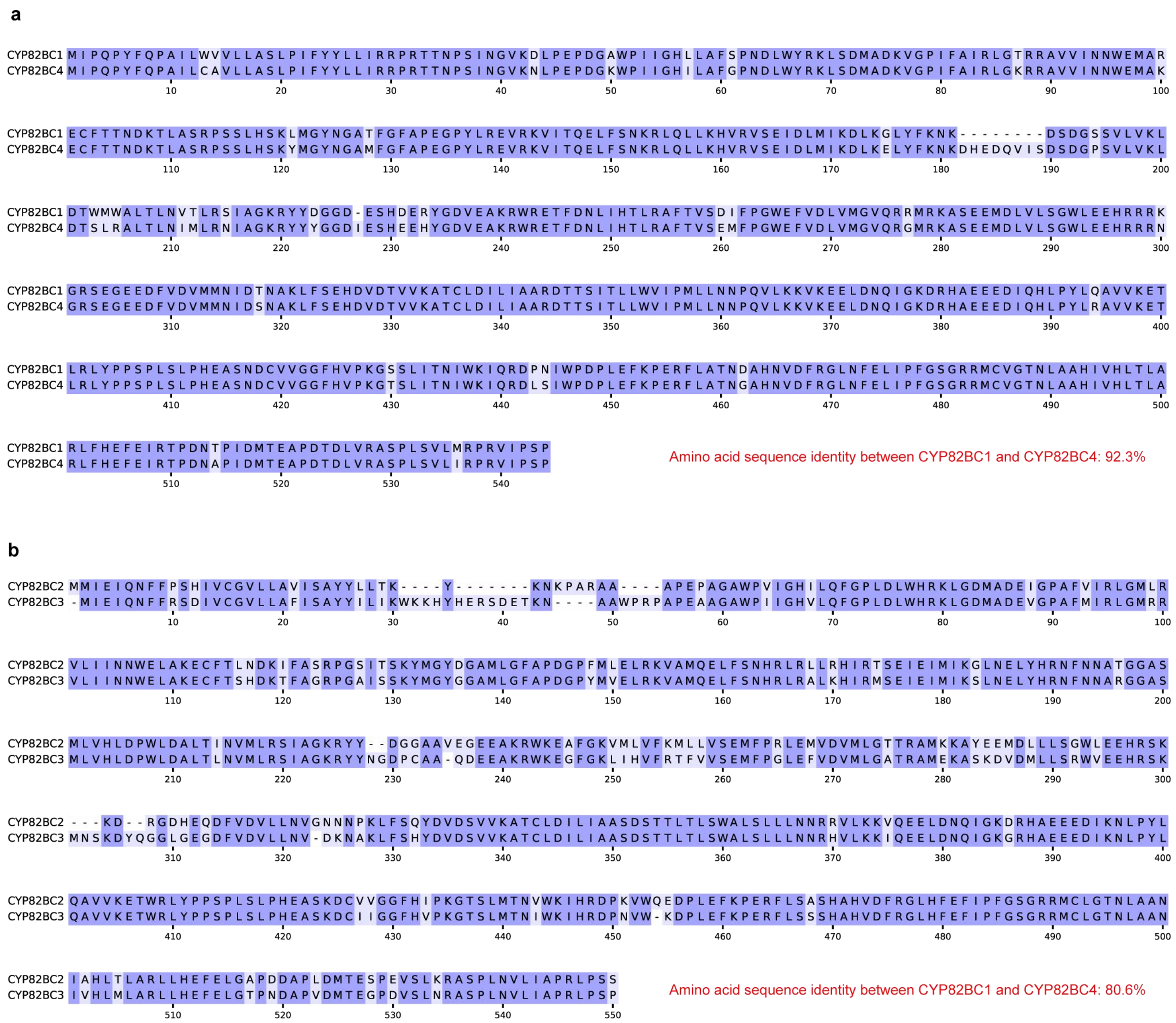


**Supplementary Fig. 11. Protein alignment of CYP82BCs. a**, Amino-acid identity between CYP82BC1 and CYP82BC4 is 92.3%. **b**, Amino-acid identity between CYP82BC2 and CYP82BC3 is 80.6%.

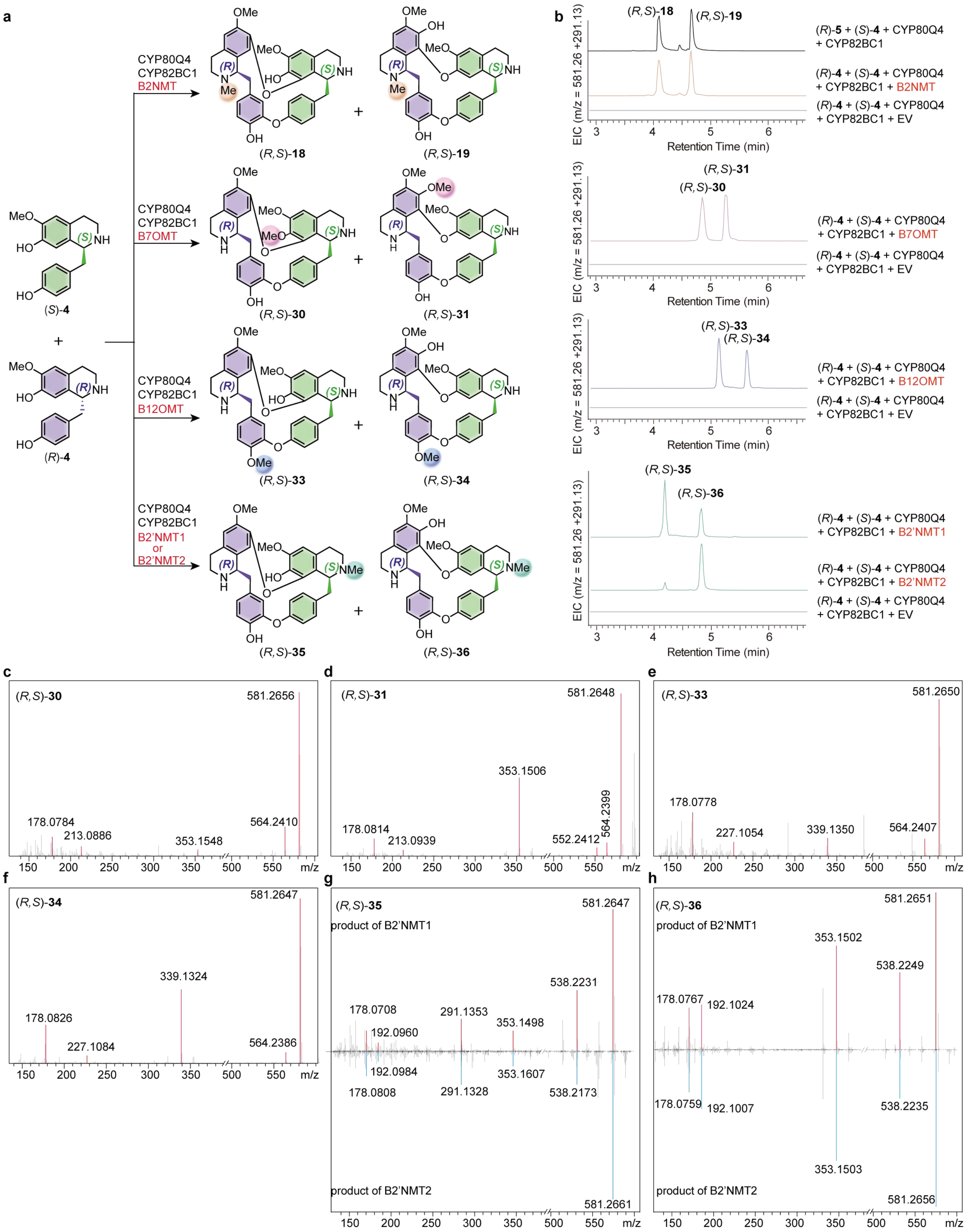


**Supplementary Fig. 12. Functional characterization of five regioselective MTs via *in vitro* cascade reactions. a**, Schematic representation of the enzymatic cascades. Reactions were initiated with the (*R*)-**4** and (*S*)-**4**, coupled with CYP80Q4, CYP82BC1, and individual candidate MTs (B2NMT, B7OMT, B12OMT, B2′NMT1, or B2′NMT2) to generate distinct monomethylated cyclic bisBIAs. Colored shading on the chemical structures highlights the specific methylation sites introduced by each corresponding MT. **b**, EICs showing the *in vitro* production of the newly methylated bisBIAs. The colors of the product traces are matched to the colored shading of the respective methylation sites in **a**. For B2NMT, the products (*R*,*S*)-**18** and (*R*,*S*)-**19** (orange trace) were confirmed by comparison with reference compounds generated via the CYP80Q4/CYP82BC1 cascade using (*R*)-**5** and (*S*)-**4** as substrates (black trace). Reactions with the empty vector (EV) served as negative controls. **c**–**f**, High-resolution MS/MS spectra of the *O*-methylated products. The B7OMT products (*R*,*S*)-**30** (**c**) and (*R*,*S*)-**31** (**d**) exhibit a diagnostic +14 Da shift in the dimeric isoquinoline fragment (*m*/*z* 353), localizing the methylation to O7 or O7′. In contrast, the B12OMT products (*R*,*S*)-**33** (**e**) and (*R*,*S*)-**34** (**f**) retain the unshifted isoquinoline fragment (*m*/*z* 339) but display a +14 Da shift in the diphenyl ether-associated fragment (*m*/*z* 227), confirming specific *O*12-methylation. **g**, **h**, Mirrored MS/MS spectra comparing the N2′-methylated products generated by the paralogous enzymes B2′NMT1 (top, red peaks) and B2′NMT2 (bottom, blue peaks). The identical fragmentation patterns confirm that both enzymes catalyze the formation of (*R*,*S*)-**35** (**g**) and (*R*,*S*)-**36** (**h**).

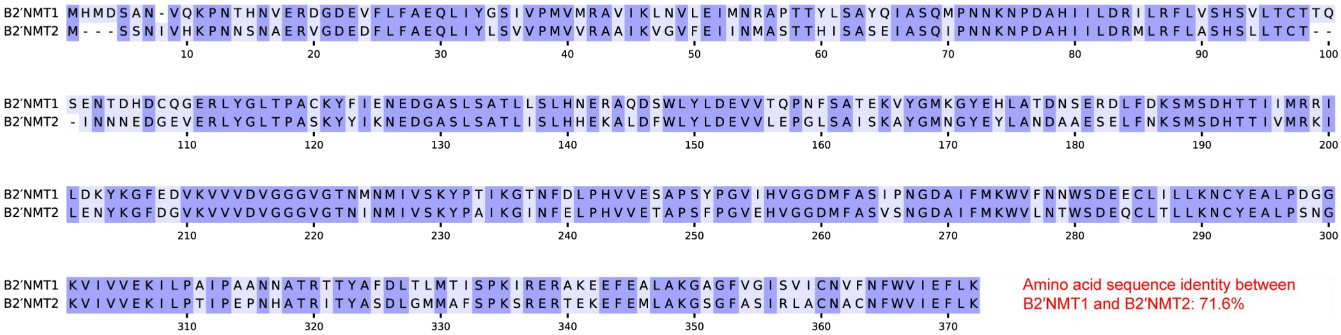


**Supplementary Fig. 13. Protein alignment of B2′NMT1 and B2′NMT2.** Amino-acid identity between B2′NMT1 and B2′NMT2 is 71.6%.

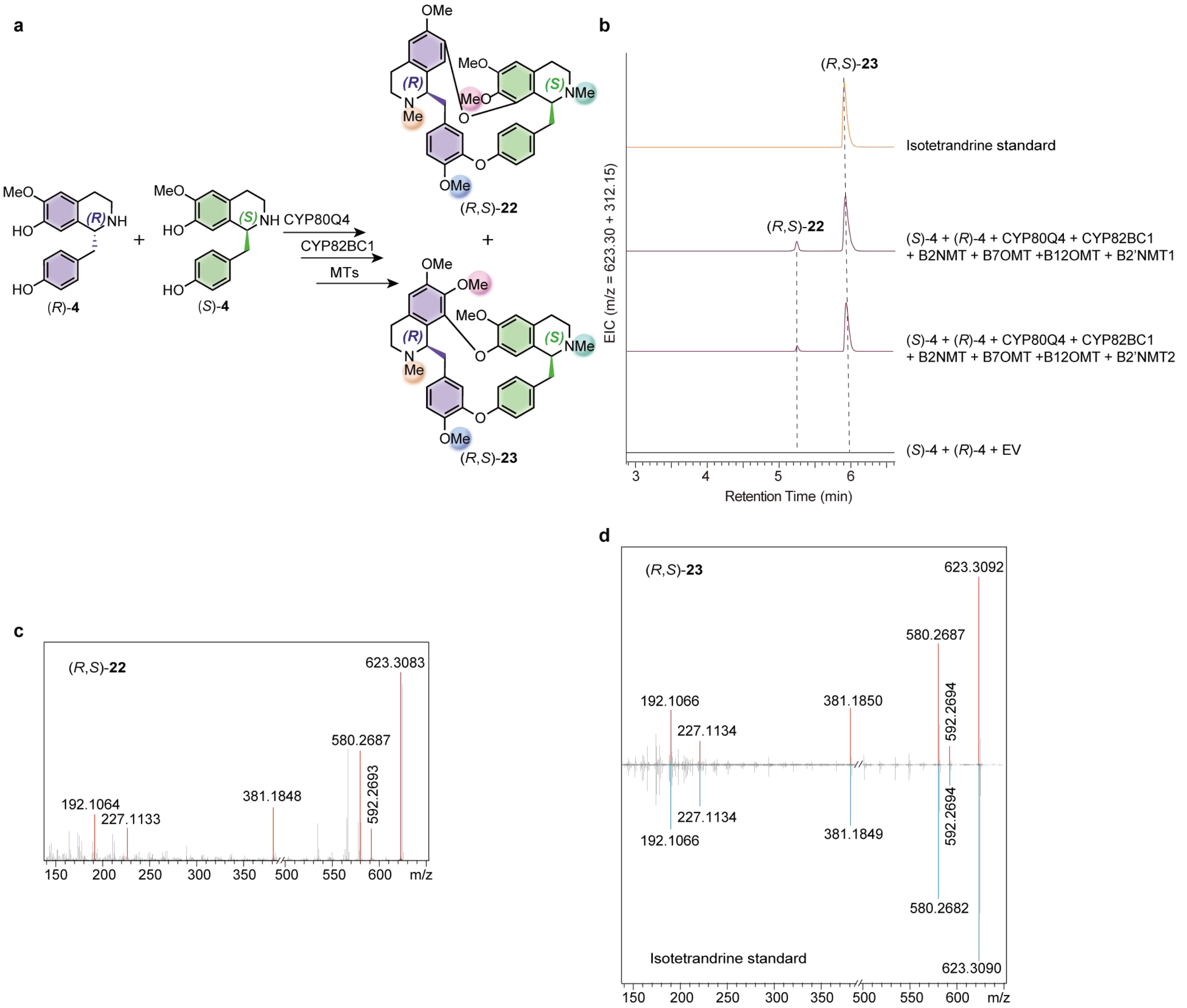


**Supplementary Fig. 14. *In vitro* reconstitution of the complete biosynthetic cascade yielding fully methylated (*R*,*S*)-bisBIAs. a**, Schematic representation of the multienzyme cascade. Co-incubation of the (*R*)-**4** and (*S*)-**4** with CYP80Q4, CYP82BC1, and a combination of four MTs (B2NMT, B7OMT, B12OMT, and either B2′NMT1 or B2′NMT2) yields the fully methylated products, obaberine ((*R*,*S*)-**22**) and isotetrandrine ((*R*,*S*)-**23**). **b**, EICs confirming the production of (*R*,*S*)-**22** and (*R*,*S*)-**23**. The enzymatic cascade reactions (purple traces) generate both products, with (*R*,*S*)-**23** perfectly co-eluting with the authentic isotetrandrine standard (orange trace). Reactions with the EV served as negative controls. **c**, MS/MS spectrum of the enzymatically produced (*R*,*S*)-**22**, assigned as obaberine based on MS/MS fragmentation and the established regioselectivities of the combined MTs (refer to the "Compound structure analysis and assignment" section for detailed structural elucidation). **d**, Mirrored MS/MS spectra confirming the structural identity of (*R*,*S*)-**23**. The fragmentation pattern of the enzymatically produced (*R*,*S*)-**23** (top, red peaks) identically matches that of the authentic isotetrandrine standard (bottom, blue peaks).

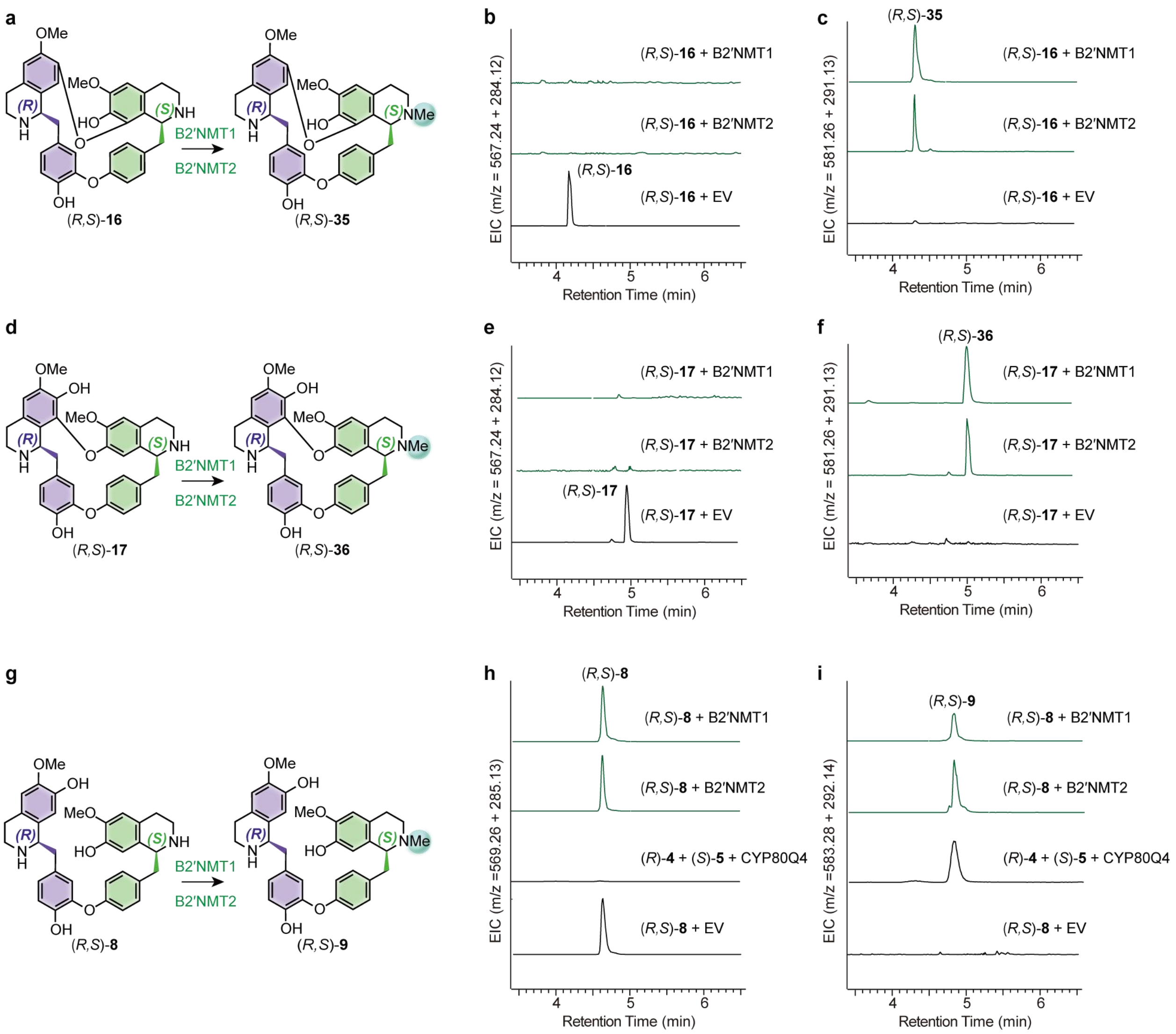


**Supplementary Fig. 15. Substrate promiscuity and biochemical characterization of the paralogous MTs B2′NMT1 and B2′NMT2.** Both enzymes catalyze regioselective methylation at the *N*2′ position and exhibit broad substrate tolerance, readily accepting both cyclic and linear bisBIA scaffolds. **a–c**, Assays using the cyclic scaffold (*R,S*)-**16**. Schematic representation of the reaction (**a**) and EICs demonstrating the depletion of the substrate (*R,S*)-**16** (**b**) and the corresponding formation of the product (*R,S*)-**35** (**c**). **d–f**, Assays using the cyclic scaffold (*R,S*)-**17**. Reaction schematic (**d**) alongside EICs confirming the consumption of (*R,S*)-**17** (**e**) and the production of (*R,S*)-**36** (**f**). **g–i**, Assays utilizing the linear (*R,S*)-**8**. Schematic representation (**g**) and EICs verifying the consumption of (*R,S*)-**8** (**h**) and the generation of (*R,S*)-**9** (**i**). Notably, in **i**, the enzymatically produced (R,S)-9 perfectly co-elutes with a reference standard generated *in vitro* via the CYP80Q4-catalyzed coupling of (*R*)-**4** and (*S*)-**5**. Throughout the figure, the green shading on the chemical structures highlights the newly introduced *N*2′-methyl groups, matching the green enzymatic product traces in the EICs. Reactions carrying the EV served as negative controls.

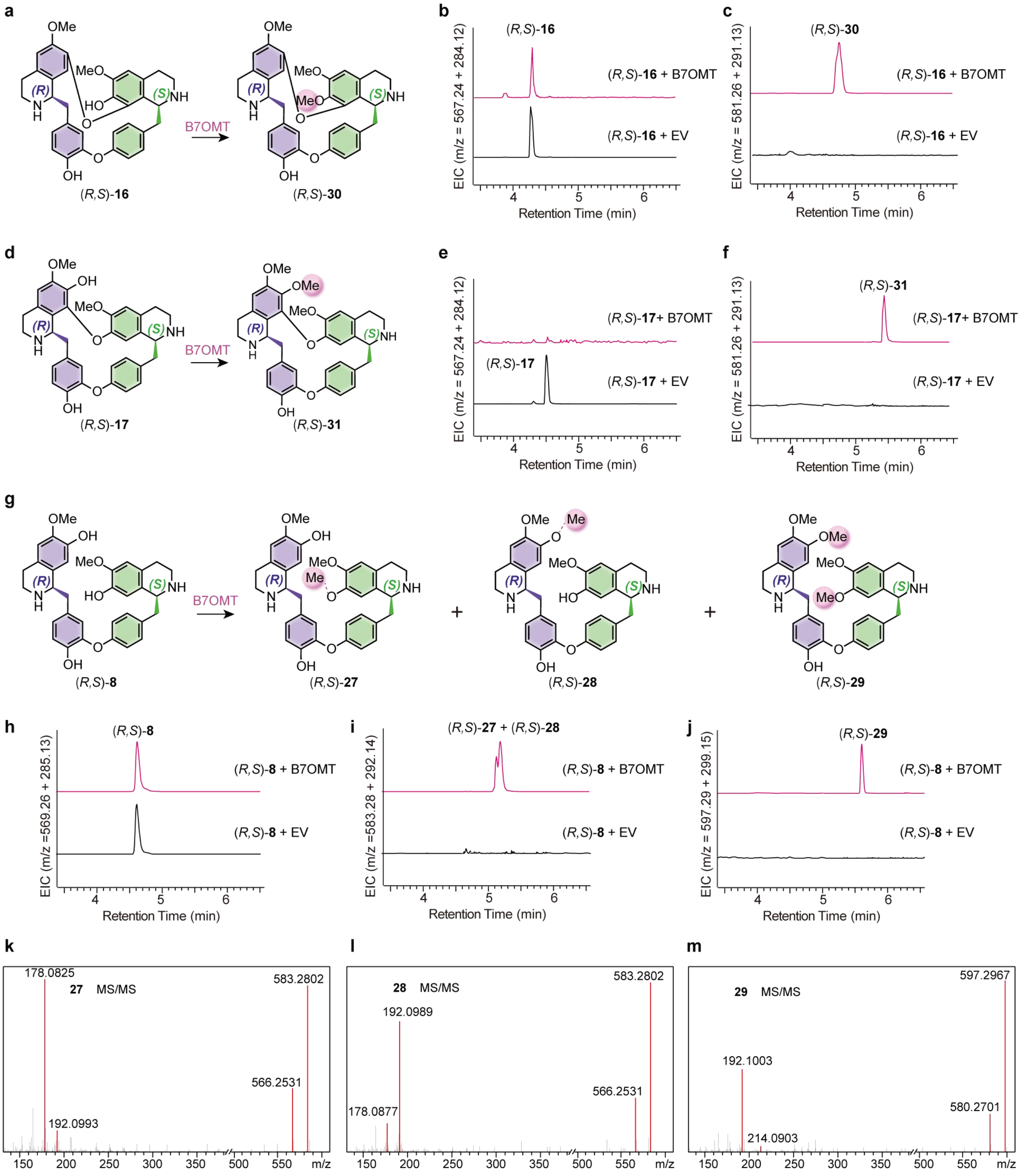


**Supplementary Fig. 16. Substrate promiscuity and biochemical characterization of the B7OMT.** The enzyme exhibits broad substrate tolerance, catalyzing *O*-methylation on both cyclic and linear bisBIA scaffolds. **a–c**, Assays using the cyclic scaffold (*R,S*)-**16**. Schematic representation of the reaction (**a**) and EICs demonstrating the consumption of (*R,S*)-**16** (**b**) and the formation of the monomethylated product (*R,S*)-**30** (**c**). **d–f**, Assays using the cyclic scaffold (*R,S*)-**17**. Reaction schematic (**d**) alongside EICs confirming the depletion of (*R,S*)-**17** (**e**) and the generation of (*R,S*)-**31** (**f**). **g–j**, Assays utilizing the linear (*R,S*)-**8**. Schematic representation of the reaction (**g**) and EICs verifying the consumption of (*R,S*)-**8** (**h**) to yield the monomethylated products (*R,S*)-**27** and (*R,S*)-**28** (**i**), as well as the dimethylated product (*R,S*)-**29** (**j**). In the chemical structures of (*R,S*)-**27** and (*R,S*)-**28**, the dashed bonds denote that while monomethylation at the dimeric isoquinoline moiety is confirmed by MS/MS, the precise regiochemical assignment to either the O7 or O7′ hydroxyl group cannot be unambiguously determined. **k–m**, High-resolution MS/MS spectra of the enzymatically produced (*R,S*)-**27** (**k**), (*R,S*)-**28** (**l**), and (*R,S*)-**29** (**m**) (refer to the “Compound structure analysis and assignment” section for detailed structural elucidation). Throughout the figure, the pink shading on the chemical structures highlights the newly introduced *O*-methyl groups, corresponding to the pink enzymatic product traces in the EICs. Reactions carrying the EV served as negative controls.

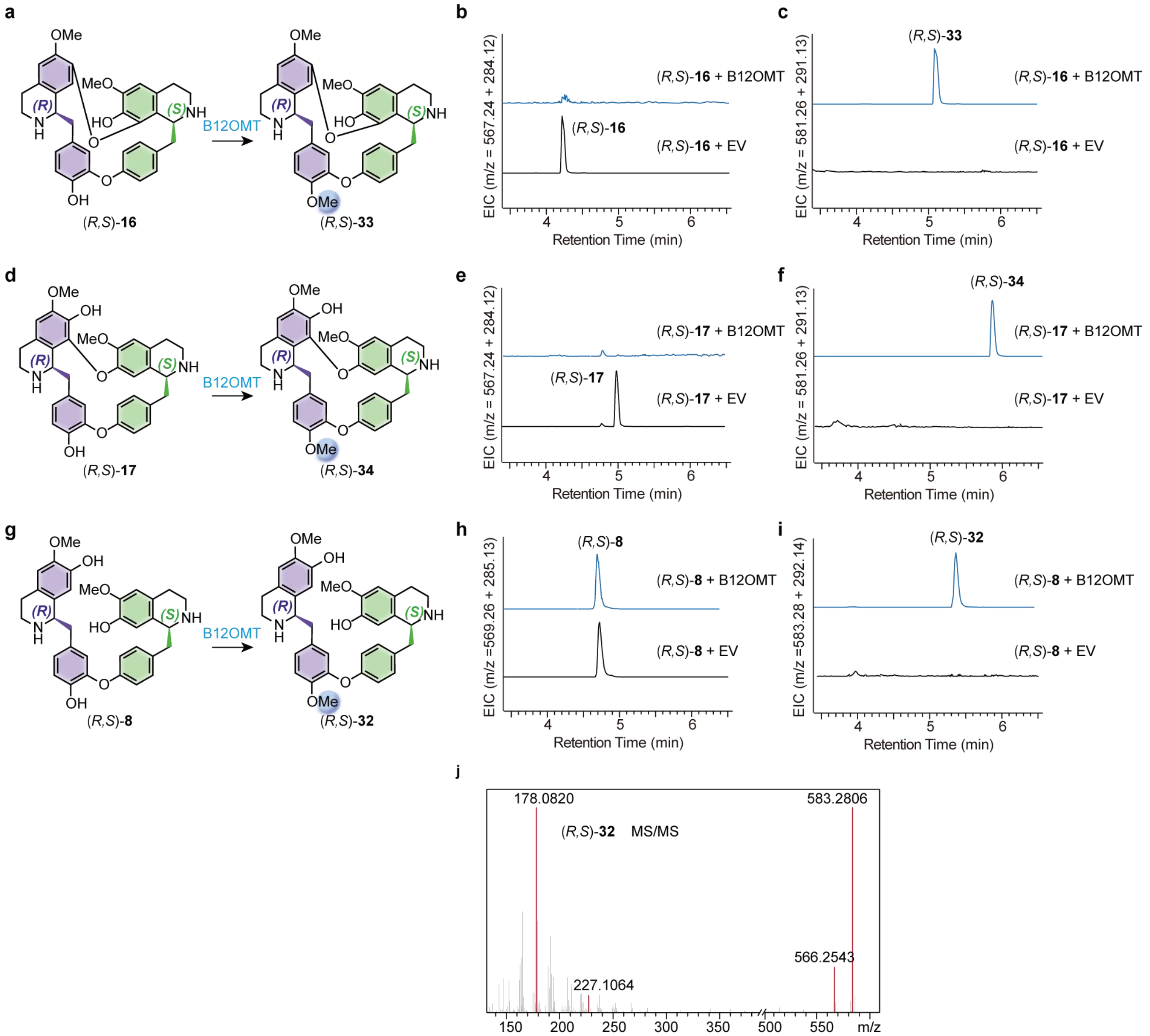


**Supplementary Fig. 17. Substrate promiscuity and biochemical characterization of the B12OMT.** The enzyme exhibits broad substrate tolerance, catalyzing strict regioselective methylation at the *O*12 position on both cyclic and linear bisBIA scaffolds. **a–c**, Assays using the cyclic scaffold (*R,S*)**-16**. Schematic representation of the reaction (**a**) and EICs demonstrating the consumption of (*R,S*)**-16** (**b**) and the formation of the monomethylated product (*R,S*)**-33** (**c**). **d–f**, Assays using the cyclic scaffold (*R,S*)**-17**. Reaction schematic (**d**) alongside EICs confirming the depletion of (*R,S*)**-17** (**e**) and the generation of (*R,S*)**-34** (**f**). **g–i**, Assays utilizing the linear (*R,S*)**-8**. Schematic representation of the reaction (**g**) and EICs verifying the consumption of (*R,S*)**-8** (**h**) to yield the monomethylated product (*R,S*)**-32** (**i**). **j**, MS/MS spectrum of the enzymatically produced (*R,S*)**-32**, whose structure and *O*12-methylation site were assigned based on diagnostic fragmentation patterns (refer to the "Compound structure analysis and assignment" section for detailed structural elucidation). Throughout the figure, the blue shading on the chemical structures highlights the newly introduced *O*12-methyl groups, corresponding to the blue enzymatic product traces in the EICs. Reactions carrying the EV served as negative controls.

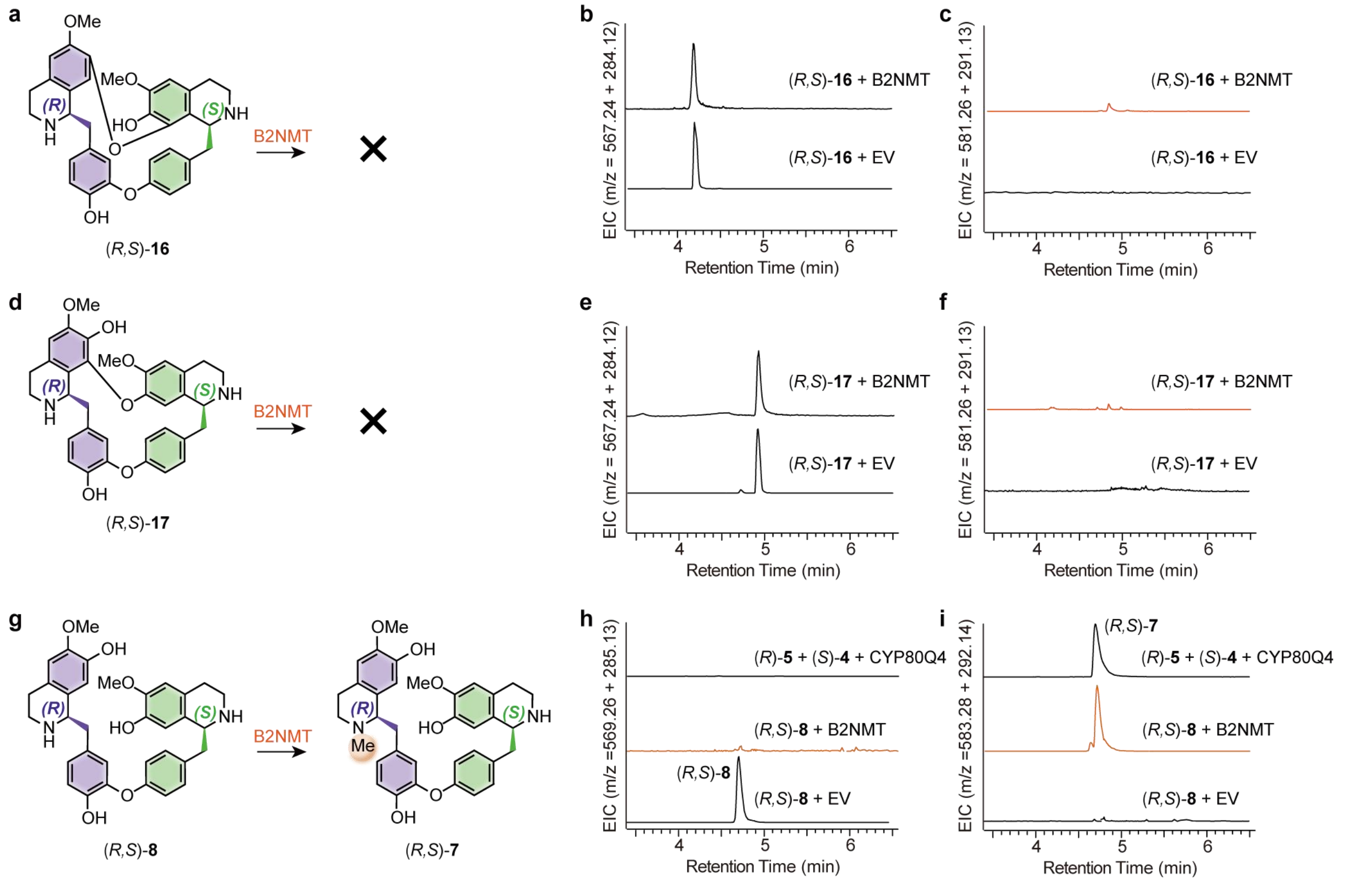


**Supplementary Fig. 18. Substrate specificity of the B2NMT.** In contrast to the promiscuous MTs, B2NMT exclusively methylates the linear bisBIA scaffold and exhibits no detectable activity toward cyclic substrates. **a–c**, Assays using the cyclic scaffold (*R*,*S*)-**16**. Schematic representation (**a**) and EICs confirming that the substrate (*R*,*S*)-**16** remains unconsumed (**b**) and no methylated products are formed (**c**). **d–f**, Assays using the cyclic scaffold (*R*,*S*)-**17**. Reaction schematic (**d**) alongside EICs showing a lack of substrate consumption (**e**) and an absence of product generation (**f**). **g–i**, Assays utilizing the linear (*R*,*S*)-**8**. Schematic representation of the reaction (g) and EICs demonstrating the efficient consumption of (*R*,*S*)-**8** (**h**) to yield the monomethylated product (*R*,*S*)-**7** (**i**). Notably, in **i**, the structural identity of the enzymatically produced (R,S)-7 (orange trace) is confirmed by its perfect co-elution with a reference product generated *in vitro* via the CYP80Q4-catalyzed coupling of (*R*)-**5** and (*S*)-**4** (top black trace). Throughout the figure, the orange shading on the chemical structure highlights the newly introduced *N*2-methyl group, corresponding to the orange enzymatic product traces in the EICs. Reactions carrying the EV served as negative controls.

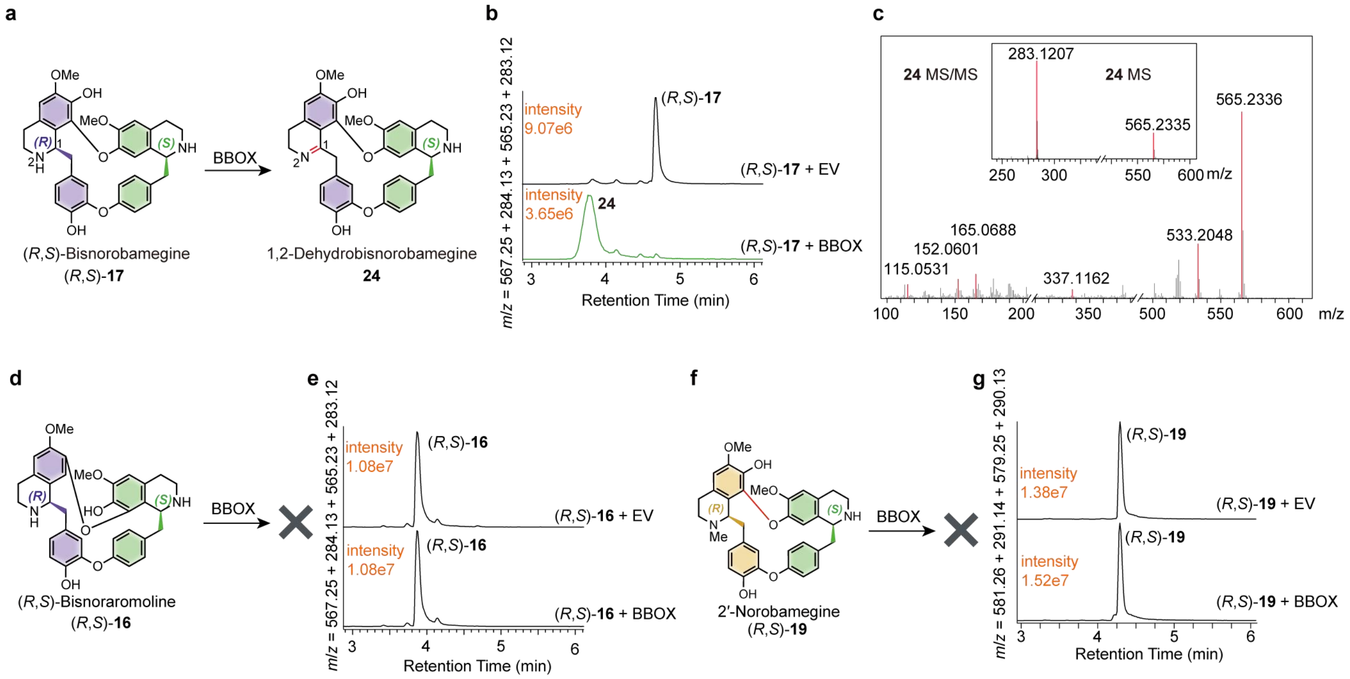


**Supplementary Fig. 19. Characterization and substrate specificity of the BBOX.** BBOX catalyzes the oxidative desaturation of specific cyclic bisBIAs to form planar imine intermediates but is inactive toward alternate regioisomers or *N*-methylated analogs. **a–c**, BBOX-catalyzed oxidation of (*R*,*S*)-**17** (*m*/*z* $[M + H]^+ = 567.25$; $[M + 2H]^{2+} = 284.13$). Schematic representation of the reaction (**a**) and EICs (**b**) demonstrating the conversion of (*R*,*S*)-**17** into the 1,2-dehydrobisnorobamegine intermediate (**24**, *m*/*z* $[M + H]^+ = 565.23$; $[M + 2H]^{2+} = 283.12$; green trace). **c**, MS (inset) and MS/MS spectra of **24**, displaying a diagnostic –2 Da mass shift relative to the substrate, consistent with dehydrogenation at the C1 position to form an imine. **d**, **e**, BBOX strictly discriminates macrocyclic regiochemistry. Reaction schematic (**d**) and EICs (**e**) confirming a complete lack of activity toward the C7–O–C8′ regioisomer (*R*,*S*)-**16** (*m*/*z* $[M + H]^+ = 567.25$; $[M + 2H]^{2+} = 284.13$), and no formation of the corresponding oxidized product (expected *m*/*z* $[M + H]^+ = 565.23$; $[M + 2H]^{2+} = 283.12$) **f**, **g**, BBOX requires a secondary amine. Reaction schematic (**f**) and EICs (**g**) revealing no detectable consumption of the *N*2-methylated analog (*R*,*S*)-**19** (*m*/*z* $[M + H]^+ = 581.26$; $[M + 2H]^{2+} = 291.14$), and no formation of the corresponding oxidized product (expected *m*/*z* $[M + H]^+ = 579.25$; $[M + 2H]^{2+} = 290.13$). In the EICs, the intensities of the highest peaks are annotated in orange. EV served as negative controls.

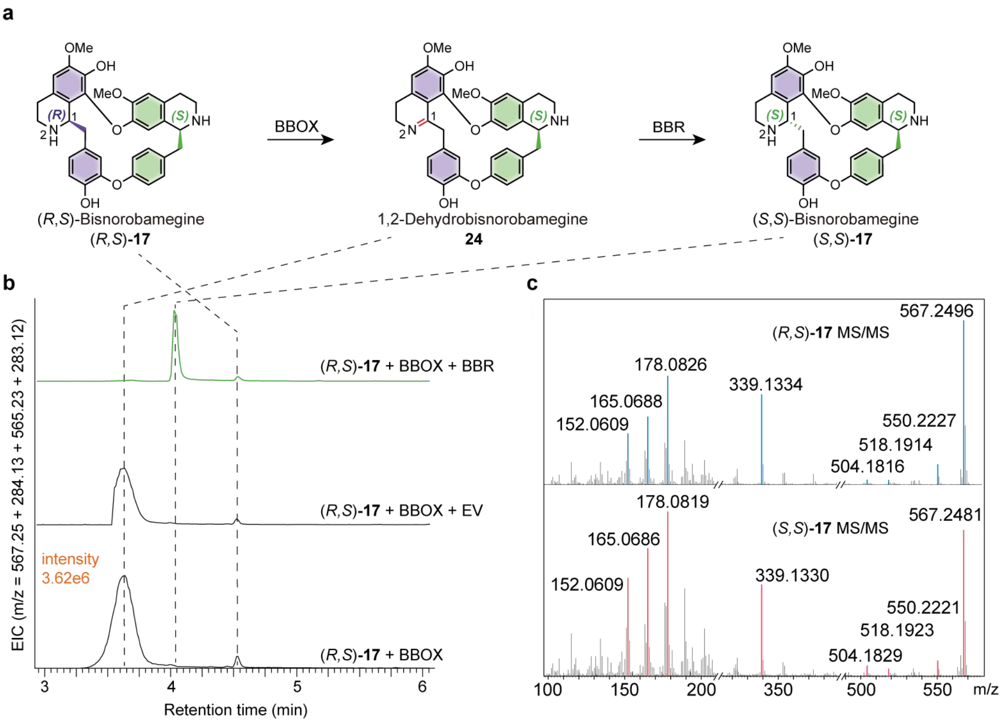


**Supplementary Fig. 20. Characterization of the BBR catalyzing stereochemical inversion. a**, Schematic representation of the two-step redox cascade. BBOX oxidizes (*R*,*S*)-**17** to the planar imine intermediate **24**, which is subsequently reduced by BBR to yield the stereoinverted diastereomer (*S*,*S*)-**17**. Dashed lines correlate the chemical structures to their respective chromatographic peaks in **b**. **b**, EICs of the enzyme assays. Incubation of (*R*,*S*)-**17** (*m*/*z* $[M + H]^+$ = 567.25; $[M + 2H]^{2+}$ = 284.13) with BBOX yields intermediate **24** (bottom trace). The subsequent addition of BBR results in the complete consumption of **24** (*m*/*z* $[M + H]^+$ = 565.23; $[M + 2H]^{2+}$ = 283.12) and the emergence of (*S*,*S*)-**17** (*m*/*z* $[M + H]^+$ = 567.25; $[M + 2H]^{2+}$ = 284.13) (top, green trace). Notably, the newly generated (*S*,*S*)-**17** elutes at a distinct retention time relative to the starting substrate (*R*,*S*)-**17**. The intensity of the highest peak is annotated in orange. Reactions with the EV in place of BBR served as negative controls (middle trace). **c**, Mirrored MS/MS spectra confirming the structural identity of the BBR product. The fragmentation pattern of the enzymatically produced (*S*,*S*)-**17** (bottom, red peaks) identically matches that of the (*R*,*S*)-**17** substrate (top, blue peaks), confirming that both compounds share an identical 2D macrocyclic bisBIA backbone despite their different stereoconfigurations.

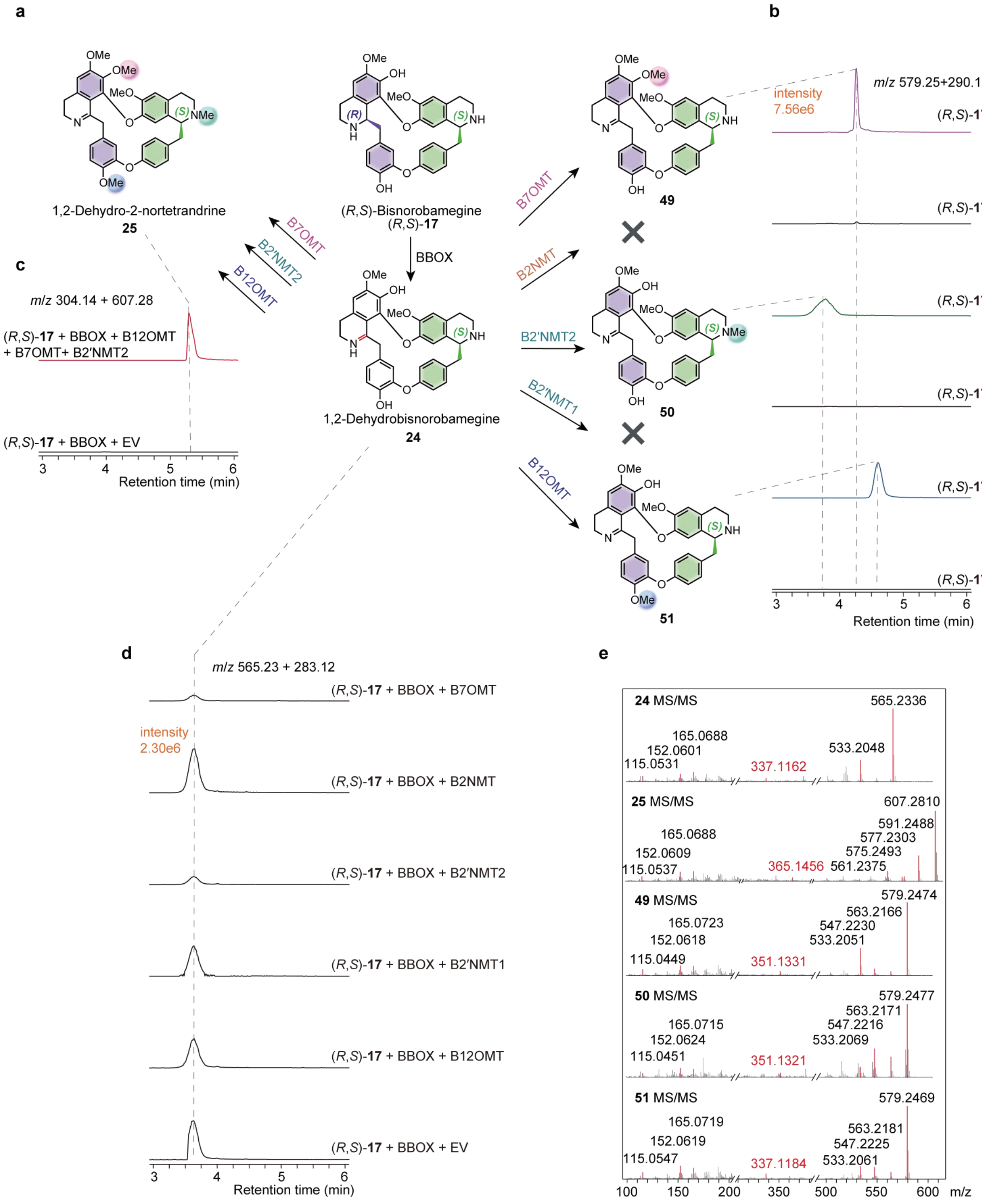


**Supplementary Fig. 21. Regioselective methylation of the BBOX-generated planar imine intermediate 24.** To demonstrate the flexibility of the stereochemical editing mechanism, the planar intermediate **24**—pre-generated in bulk via the BBOX-catalyzed oxidation of (*R*,*S*)-**17** and subsequently isolated—was utilized as a substrate for previously characterized MTs. **a**, Schematic representation of the consecutive oxidation–methylation cascade. The central iminium intermediate **24** is selectively methylated by individual MTs to yield monomethylated derivatives (**49**, **50**, **51**), or fully methylated to 1,2-dehydro-2-nortetrandrine (**25**) using an MT cocktail. Notably, B2NMT and B2′NMT1 exhibit no activity toward the oxidized scaffold. **b**, EICs monitoring the formation of monomethylated products. Individual incubations of **24** with B7OMT, B2′NMT2, and B12OMT successfully yield **49** (purple trace), **50** (green trace), and **51** (blue trace), respectively. In agreement with the schematic, B2NMT and B2′NMT1 fail to generate any detectable products. **c**, EIC demonstrating the efficient generation of the trimethylated product **25** (red trace) when **24** is co-incubated with a combined cocktail of B7OMT, B2′NMT2, and B12OMT. **d**, EICs tracking the remaining (unreacted) substrate **24** across the individual assays shown in **b**, with the substrate depletion profiles perfectly mirroring the product formation events. **e**, Comparative MS/MS spectra of **24**, the trimethylated **25**, and the monomethylated products **49**–**51**. Diagnostic fragments (annotated in red) confirm that the respective MTs maintain their strict regioselectivity on the planar oxidized scaffold (refer to the "Compound structure analysis and assignment" section for detailed structural elucidation). Throughout the figure, dashed lines correlate specific chromatographic peaks to their corresponding chemical structures. The intensities of the highest peaks are annotated in orange, and all traces within a given panel are plotted on identical intensity scales. Reactions carrying the EV served as negative controls.

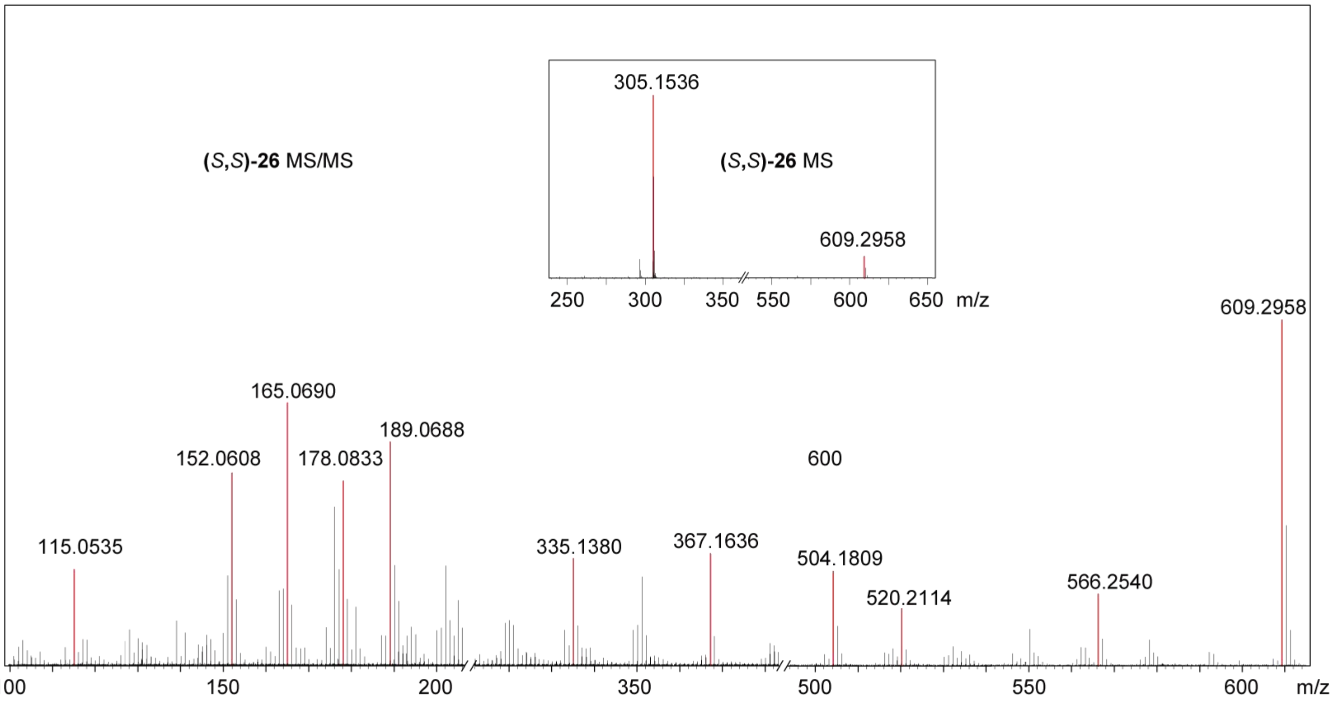


**Supplementary Fig. 22. MS/MS spectrum of the enzymatically generated (*S*,*S*)-26.**

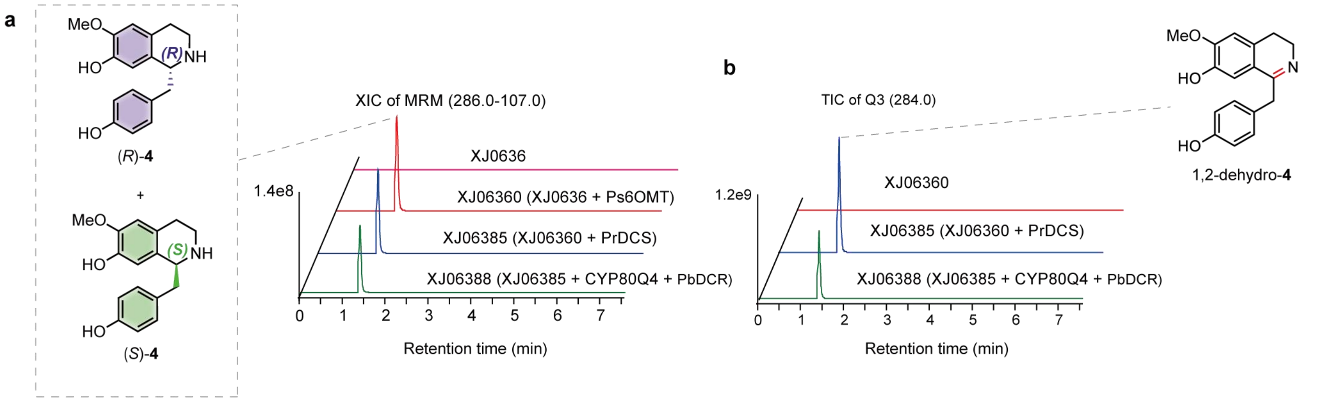


**Supplementary Fig. 23. Profiling of engineered yeast strains for the in vivo production of (*R*)-4 and (*S*)-4.** As demonstrated in Fig. 5b, the strain co-expressing CYP80Q4, PrDCS, and PbDCR successfully produces the linear intermediate (*R*,*S*)-**8**, confirming that the introduction of the PrDCS/PbDCR module efficiently converts a portion of (*S*)-**4** into (*R*)-**4**. **a**, XICs of MRM transitions (*m*/*z* 286.0–107.0) monitoring the accumulation of (*S*)-**4** and its epimer (*R*)-**4**. Starting from the chassis XJ06360, which exclusively produces (*S*)-**4** (red trace), the heterologous integration of *PrDCS* and *PbDCR* in strain XJ06388 (green trace) enables the detectable formation of both enantiomers, demonstrating the functional reconstitution of the stereochemical inversion branch. **b**, Identification of the transient dehydrogenated intermediate 1,2-dehydro-**4**. LC–MS analysis tracking the TIC of Q3 (m/z 284.0) reveals a massive accumulation of this intermediate in the XJ06360-derived strain XJ06385 (blue trace), which expresses *PrDCS* but lacks the reductase *PbDCR*. The subsequent integration of PbDCR in strain XJ06388 (green trace) resolves this metabolic bottleneck, driving the intermediate 1,2-dehydro-**4** toward the final product (*R*)-**4**. Dashed lines correlate the chemical structures to their peaks.

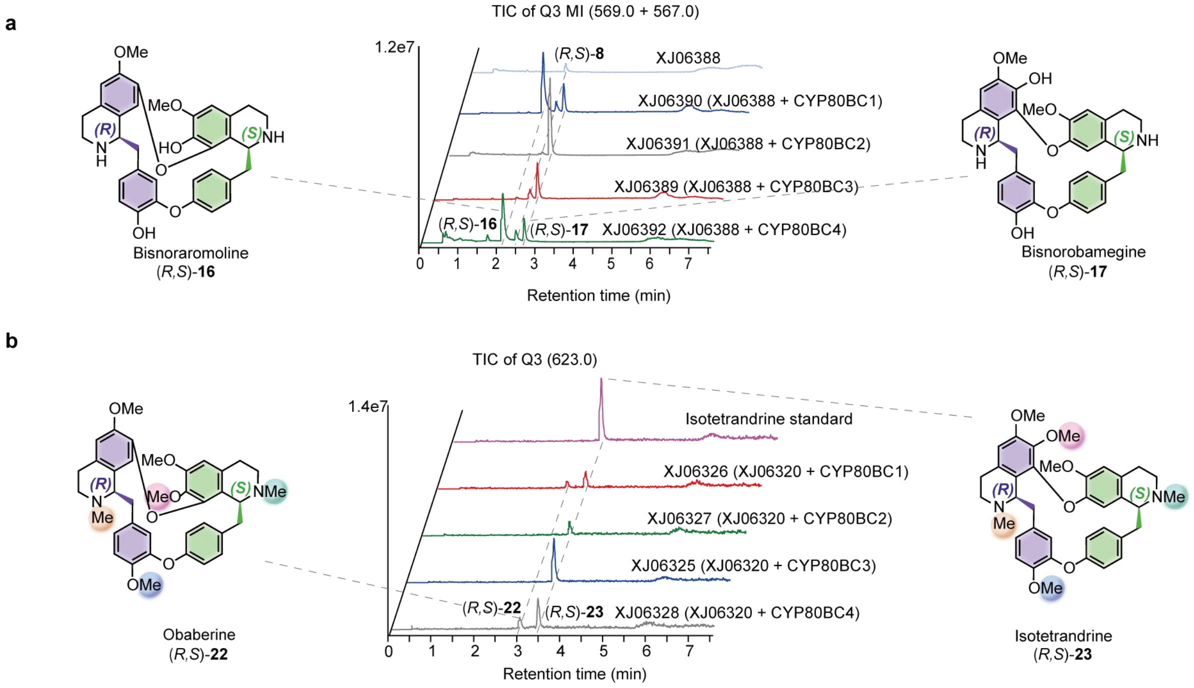


**Supplementary Fig. 24. Construction of yeast strains for cyclic (*R*,*S*)-bisBIA production. a**, Comparative profiling of CYP82BC paralogs for core macrocycle formation. The heterologous integration of individual CYP82BC enzymes (CYP82BC1–4) into the linear bisBIA-producing chassis XJ06388 successfully enabled the *in vivo* generation of the cyclic scaffolds (*R*,*S*)-**16** and (*R*,*S*)-**17**. The TIC of Q3 multiple ions (MI, *m*/*z* 569.0 + 567.0) highlights the consumption of the linear intermediate (*R*,*S*)-**8** (top blue trace) and the corresponding emergence of the cyclic products. **b**, Engineering and evaluation of strains for fully methylated (*R*,*S*)-bisBIA biosynthesis. To boost precursor supply, the copy numbers of *PrDCS*, *PbDCR*, and *CYP80Q4* were increased in the XJ06388 background, yielding the optimized chassis XJ06399. Subsequent integration of the tailoring MT module (*B2NMT*, *B12OMT*, *B7OMT*, and *B2'NMT2*) created strain XJ06320. The final integration of specific CYP82BC enzymes (CYP82BC1–4) into the XJ06320 background successfully drove the pathway to the fully methylated cyclic products (*R*,*S*)-22 and (*R*,*S*)-23. The TIC of Q3 (*m*/*z* 623.0) confirms their production, with (*R*,*S*)-23 matching the retention time of the authentic isotetrandrine standard (top purple trace). Throughout the figure, dashed lines correlate the chemical structures to their respective diagnostic chromatographic peaks.

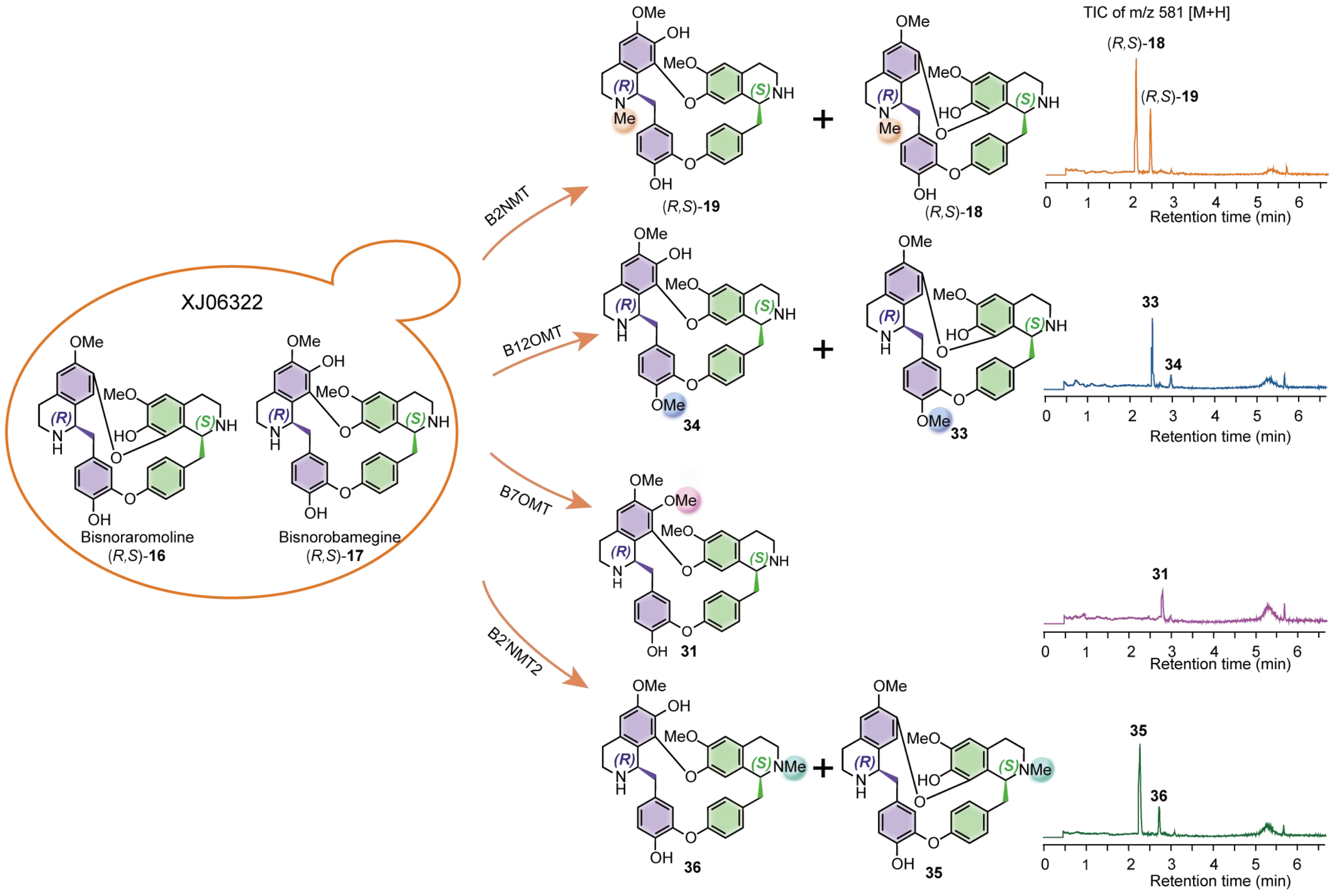


**Supplementary Fig. 25.** ***De novo*** **production of monomethylated bisBIA analogs via modular integration of single regioselective MTs** ***in vivo.*** The engineered yeast chassis XJ06322, which predominantly accumulates the core cyclic bisBIA scaffolds (*R*,*S*)-**16** and (*R*,*S*)-**17**, was utilized as a platform for targeted structural diversification. Heterologous integration and expression of individual MT genes (*B2NMT*, *B12OMT*, B*7OMT*, or *B2'NMT2*) in this background successfully yielded distinct profiles of monomethylated bisBIA derivatives. Specifically, the expression of B2NMT generated (*R*,*S*)-**18** and (*R*,*S*)-**19**; B12OMT produced (*R*,*S*)-**33** and (*R*,*S*)-**34**; B7OMT led to the formation of (*R*,*S*)-**31**; and B2′NMT2 yielded (*R*,*S*)-**35** and (*R*,*S*)-**36**. Total ion chromatograms (TICs) extracting the diagnostic mass for monomethylated products (*m*/*z* 581 $[M + H]^+$) confirm their efficient accumulation in the yeast cultures. Colored shading on the chemical structures highlights the distinct regiochemical methylation sites introduced by each enzyme, which perfectly match the colors of the corresponding chromatographic product traces (orange, blue, pink, and green).

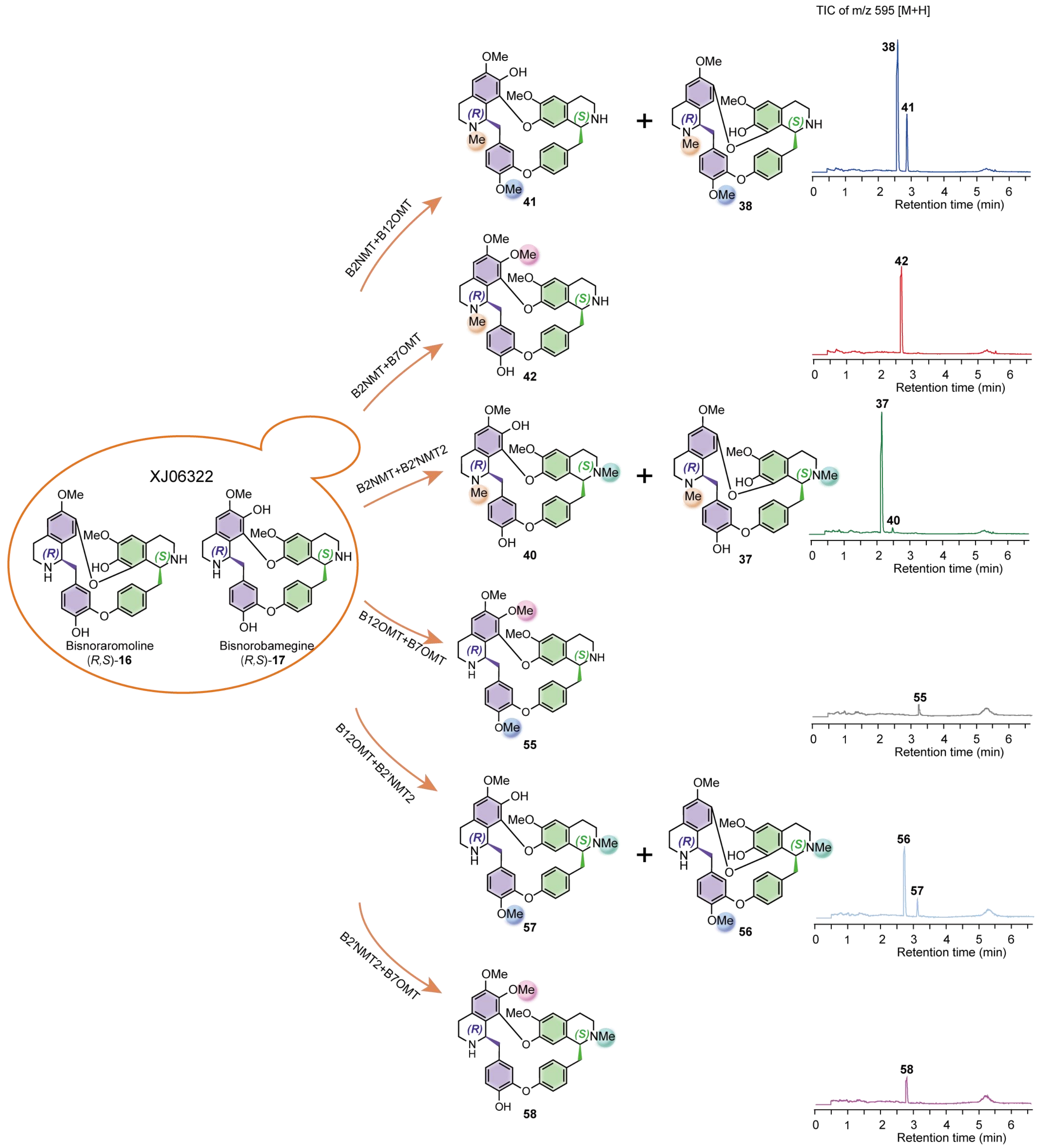


**Supplementary Fig. 26. *De novo* production of dimethylated bisBIA analogs via combinatorial pairwise integration of regioselective MTs *in vivo*.** Building upon the modular monomethylation platform, the engineered yeast chassis XJ06322—which accumulates the core cyclic scaffolds (*R*,*S*)-**16** and (*R*,*S*)-**17**—was further leveraged for higher-order structural diversification. Heterologous co-expression of the tailoring methyltransferase genes (*B2NMT*, *B12OMT*, *B7OMT*, and *B2'NMT2*) in six distinct pairwise combinations successfully directed the metabolic flux toward a library of dimethylated bisBIA derivatives. Specifically, the respective enzymatic combinations yielded the following products: *B2NMT* and *B12OMT* produced (*R*,*S*)-**38** and (*R*,*S*)-**41**; *B2NMT* and *B7OMT* generated (*R*,*S*)-**42**; *B2NMT* and *B2'NMT2* yielded (*R*,*S*)-**37** and (*R*,*S*)-**40**; *B12OMT* and *B7OMT* led to (*R*,*S*)-**55**; *B12OMT* and *B2'NMT2* produced (*R*,*S*)-**56** and (*R*,*S*)-**57**;

and *B2'NMT2* and *B7OMT* formed (*R*,*S*)-**58**. TICs extracting the diagnostic mass for dimethylated bisBIAs (*m*/*z* 595 $[M + H]^+$) confirm their efficient *in vivo* accumulation. Colored shading on the chemical structures explicitly delineates the distinct dual regiochemical methylation sites introduced by each specific MT pair.

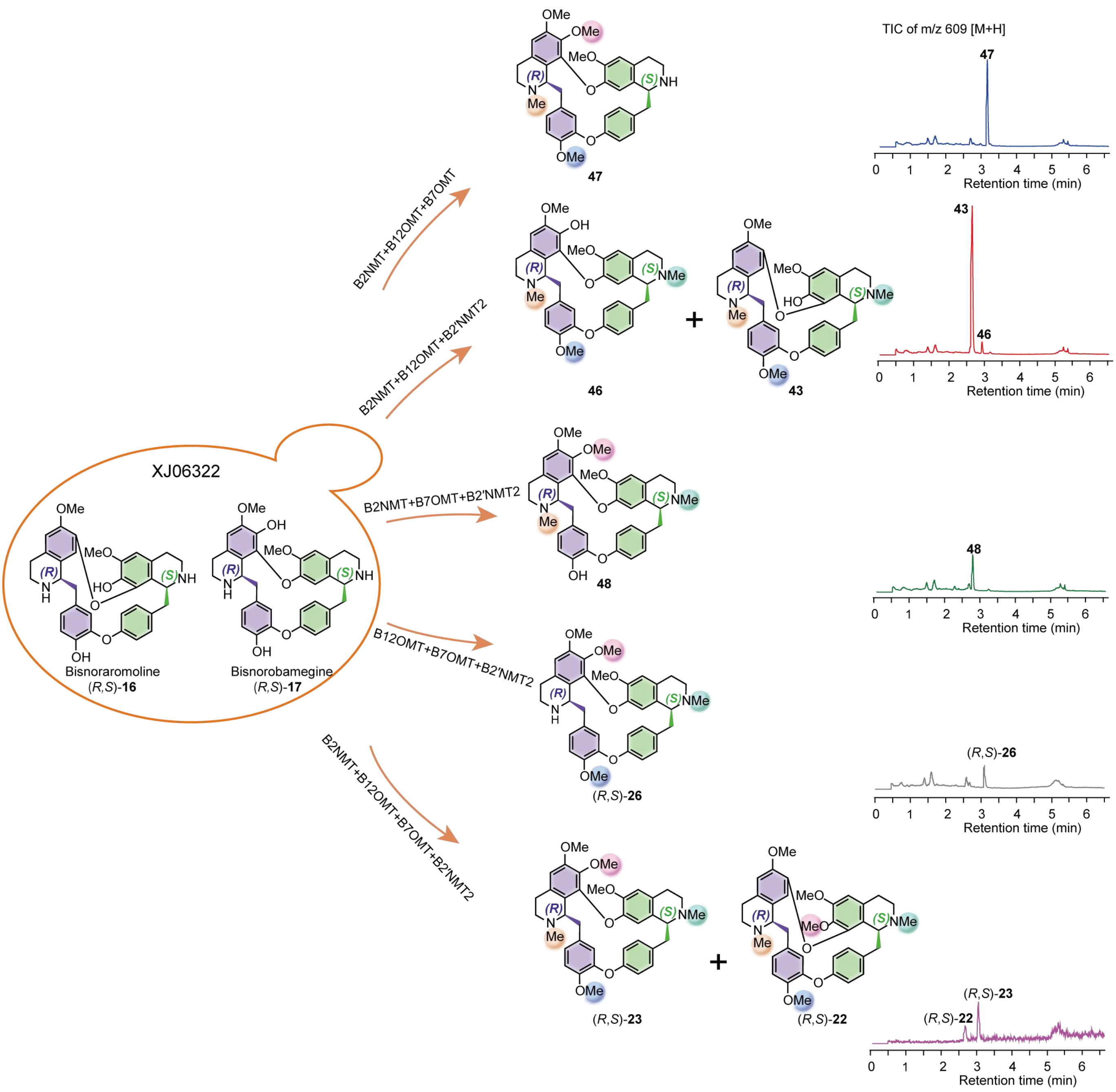


**Supplementary Fig. 27.** ***De novo*** **production of trimethylated and fully methylated bisBIA analogs via combinatorial integration of three or four MTs** ***in vivo*****.** Extending the modular biosynthesis platform to its maximum complexity, the engineered yeast chassis XJ06322—which predominantly produces the core cyclic scaffolds (*R*,*S*)-**16** and (*R*,*S*)-**17**—was leveraged for high-level structural diversification. Heterologous co-expression of the tailoring MT genes (*B2NMT*, *B12OMT*, *B7OMT*, and *B2'NMT2*) in higher-order combinations (groups of three or all four) successfully directed the metabolic flux toward a library of highly decorated bisBIA derivatives. Specifically, the co-expression of three MTs yielded distinct trimethylated products: the combination of *B2NMT*, *B12OMT*, and *B7OMT* produced **47**; *B2NMT*, *B12OMT*, and *B2'NMT2* generated **43** and **46**; *B2NMT*, *B7OMT*, and *B2'NMT2* yielded **48**; and *B12OMT*, *B7OMT*, and *B2'NMT2* formed (*R*,*S*)-**26**. Furthermore, the simultaneous integration of all four MTs successfully pushed the pathway to the fully methylated (tetramethylated) endpoints, yielding (*R*,*S*)-**22** and (*R*,*S*)-**23**. TICs extracting the diagnostic masses (e.g., m/z 609 $[M + H]^+$ for trimethylated analogs) confirm their efficient *in vivo* accumulation. Colored shading on the chemical structures explicitly delineates the complex, multi-site regiochemical methylation patterns achieved by these combinatorial MT ensembles, corresponding to the respective colored chromatographic product traces.

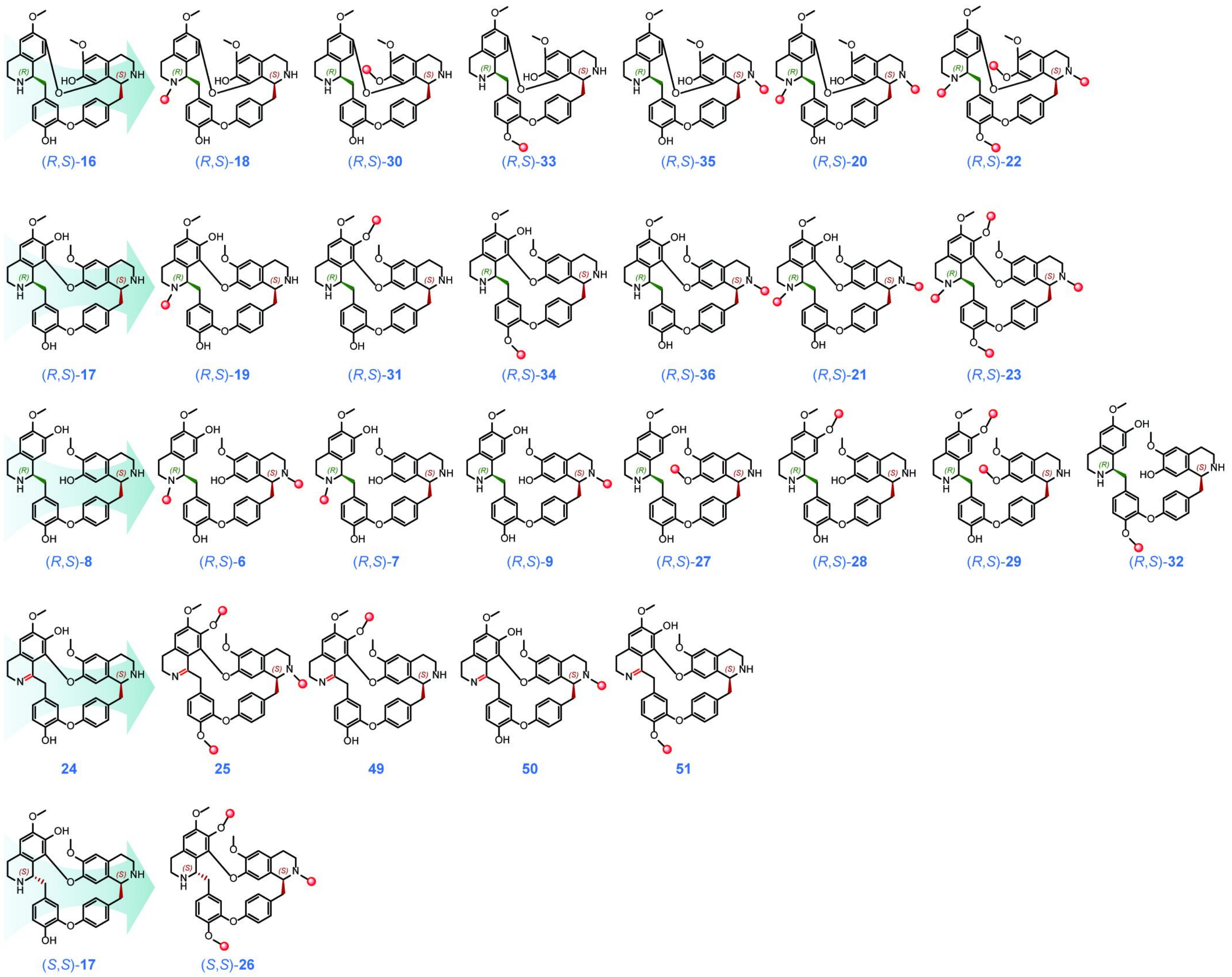


**Supplementary Fig. 28. Chemical space and structural diversity of bisBIAs accessed via the elucidated biosynthetic toolkit.** A comprehensive summary of the bisBIA analogs successfully generated *in vitro* and/or *in vivo* (in engineered yeast cell factories) in this study. The modular application of the characterized biosynthetic enzymes—including coupling CYPs, macrocyclizing CYPs, regioselective MTs, and the redox-mediated stereochemical inversion module (BBOX/BBR)—enables programmable access to a highly diversified structural library. The compounds are categorized based on their scaffolds (indicated by the large teal arrows): derivatives of the core cyclic scaffolds (*R*,*S*)-**16** (first row) and (*R*,*S*)-**17** (second row); analogs originating from the linear intermediate (*R*,*S*)-8 (third row); heavily decorated derivatives of the planar imine intermediate **24** (fourth row); and stereoinverted products derived from the (*S*,*S*)-configured scaffold (*S*,*S*)-17 (fifth row). In the chemical structures, the stereochemical bonds at the C1/C1′ positions are highlighted in green, while newly introduced modifications (such as regioselective *N*- and *O*-methylations or oxidative desaturation) are highlighted in red to trace the enzymatic decoration events.

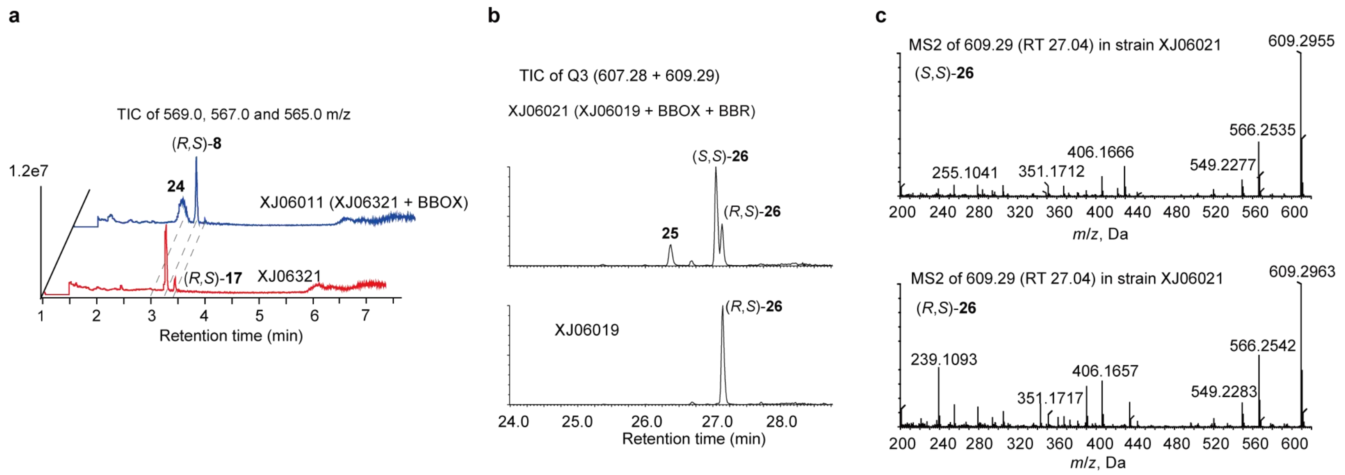


**Supplementary Fig. 29. *In vivo* biosynthesis of (*S*,*S*)-configured bisBIAs via stereochemical inversion in engineered yeast. a**, *In vivo* oxidation of the cyclic scaffold. TICs of MI, (*m*/*z* 569.0, 567.0, and 565.0 $[M + H]^+$) demonstrate that the chassis strain XJ06321 (bottom red trace) produces the cyclic scaffold (*R*,*S*)-**17** alongside the linear intermediate (*R*,*S*)-**8**. Heterologous integration of the oxidase gene BBOX (strain XJ06011, top blue trace) results in the targeted consumption of (*R*,*S*)-**17** and the corresponding accumulation of the dehydrogenated planar intermediate **24**. Dashed lines highlight the retention time correlation between the precursor and its oxidized product. **b**, LC–QTOF-MS analysis confirming functional stereochemical inversion. TICs of Q3 MI (*m*/*z* 607.28 and 609.29 $[M + H]^+$) reveal that while the precursor strain XJ06019 exclusively accumulates the (*R*,*S*)-26 diastereomer (bottom trace), co-expression of the complete redox module (BBOX and BBR) in strain XJ06021 effectively drives the *in vivo* generation of the stereoinverted product (*S*,*S*)-26, accompanied by the oxidized intermediate **25** (top trace). **c**, MS/MS spectra of (*S*,*S*)-26 (top) and (*R*,*S*)-26 (bottom) acquired from the engineered strain XJ06021. Both diastereomers exhibit virtually identical fragmentation patterns derived from the parent ion (*m*/*z* 609.29 $[M + H]^+$), confirming they share an identical 2D macrocyclic backbone despite their distinct stereoconfigurations.

### Supplementary Tables

**Supplementary Table 1.** ***S. tetrandra*** **genome assembly and annotation statistics**

| | |
|---|---|
| Pseudochromosomes | 13 |
| Total length (Mbp) | 924.44 |
| N50 (Mbp) | 2.07 |
| Predicted protein-coding genes | 22,226 |
| Complete and single-copy BUSCO | 91.8% |
| Illumina short-read mapping | 99.82% |

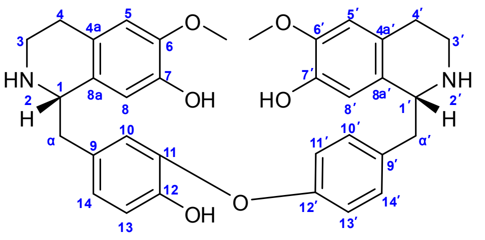


(*R*,*S*)-8

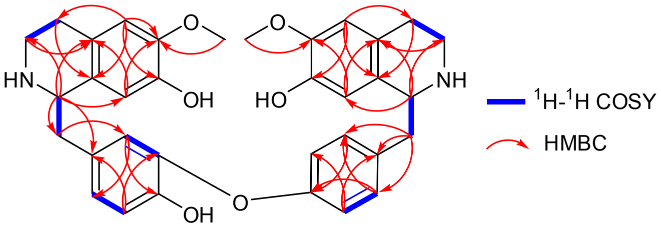


(*R*,*S*)-8

**Supplementary Table 2. $^{1}H$ and $^{13}C$ NMR data of (*R*,*S*)-8.**

| Pos. | $\delta_C$, type | $\delta_H$, mult. (*J* in Hz) |
|---|---|---|
| 1 | 55.5 | 4.54, m |
| 3 | 39.0 | 3.36, m |
| 4 | 24.5 | 2.92-2.81, m |
| 4a | 122.6 | |
| 5 | 112.3 | 6.73, s |
| 6 | 147.8 | |
| 7 | 145.3 | |
| 8 | 113.9 | 6.47, s |
| 8a | 124.5 | |
| α | 39.1 | 3.20, m<br>3.15, dd, (14.4, 6.6) |
| 9 | 127.4 | |
| 10 | 123.7 | 6.88, d, (1.8) |
| 11 | 142.7 | |
| 12 | 149.0 | |
| 13 | 117.9 | 6.97, d, (8.4) |
| 14 | 126.9 | 7.01, dd, (8.4, 1.8) |
| 1′ | 55.5 | 4.59, t, (6.0) |
| 3′ | 39.0 | 3.36, td, (12.0, 6.0) |
| 4′ | 24.5 | 2.92-2.81, m |
| 4a′ | 122.8 | |
| 5′ | 112.3 | 6.76, s |
| 6′ | 147.8 | |
| 7′ | 145.4 | |
| 8′ | 113.9 | 6.52, s |
| 8a′ | 124.7 | |
| α′ | 39.3 | 3.20, m<br>3.02, dd, (14.4, 7.8) |
| 9′ | 129.7 | |
| 10´ | 131.1 | 7.22, m |
| 11′ | 116.7 | 6.85, m |
| 12′ | 157.6 | |
| 13′ | 116.7 | 6.85, m |
| 14′ | 131.1 | 7.22, m |
| 6-$OCH_3$ | 56.1 | 3.75, s |
| 6′-$OCH_3$ | 55.3 | 3.76, s |

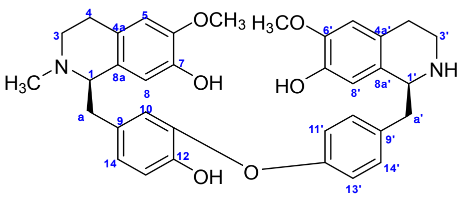


**(*R*,*S*)-7**

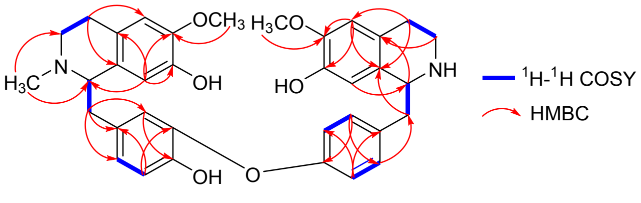


**(*R*,*S*)-7**

**Supplementary Table 3. $^{1}$H and $^{13}$C NMR data of (*R*,*S*)-7.**

| Pos. | $\delta_C$, type | $\delta_H$, mult. (*J* in Hz) |
|---|---|---|
| 1 | 65.9 | 4.36, m |
| 3 | 46.5 | 3.65, m<br>3.31, m |
| 4 | 23.3 | 3.05, m<br>2.91, m |
| 4a | 122.1 | |
| 5 | 112.8 | 6.71, s |
| 6 | 149.4 | |
| 7 | 146.1 | |
| 8 | 116.1 | 5.97, s |
| 8a | 123.7 | |
| α | 39.8 | 3.30, m<br>2.98, m |
| 9 | 128.6 | |
| 10 | 123.4 | 6.46, s |
| 11 | 145.3 | |
| 12 | 149.2 | |
| 13 | 118.4 | 6.95, m |
| 14 | 127.0 | 6.93, m |
| 1′ | 53.0 | 4.60, t, (8.0) |
| 3′ | 40.4 | 3.54, m<br>3.33, m |
| 4′ | 25.9 | 3.06, m |
| 4a′ | 123.7 | |
| 5′ | 112.6 | 6.78, s |
| 6′ | 149.2 | |
| 7′ | 146.5 | |
| 8′ | 114.5 | 6.33, s |
| 8a′ | 125.0 | |
| α′ | 42.5 | 3.22, m<br>3.11, m |
| 9′ | 113.0 | |
| 10´ | 131.9 | 7.21, d, (8.0) |
| 11′ | 119.4 | 6.89, d, (8.0) |
| 12′ | 158.6 | |
| 13′ | 119.4 | 6.89, d, (8.0) |
| 14′ | 131.9 | 7.21, d, (8.0) |
| 6-$OCH_3$ | 56.4 | 3.83, s |
| 6′-$OCH_3$ | 56.6 | 3.85, s |
| 2-$NCH_3$ | 40.6 | 2.88, s |

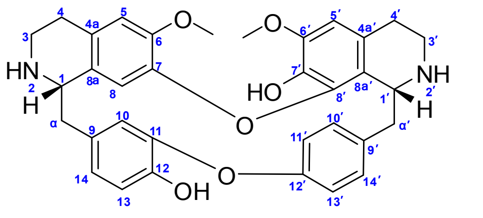


**(*R*,*S*)-16**

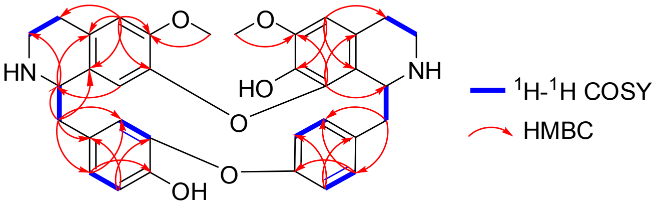


**(*R*,*S*)-16**

**Supplementary Table 4. $^1$H and $^{13}$C NMR data of (*R*,*S*)-16.**

| Pos. | $\delta_C$, type | $\delta_H$, mult. (*J* in Hz) |
|---|---|---|
| 1 | 54.4 | 4.17, m |
| 3 | 37.3 | 3.36-3.20, m |
| 4 | 28.3 | 2.48-2.31, m |
| 4a | 130.0 | |
| 5 | 112.5 | 6.53, s |
| 6 | 149.0 | |
| 7 | 144.8 | |
| 8 | 116.7 | 6.68, s |
| 8a | 127.2 | |
| α | 41.2 | 2.83-2.63, m |
| 9 | 127.7 | |
| 10 | 116.3 | 5.41, d, (1.8) |
| 11 | 141.9 | |
| 12 | 148.4 | |
| 13 | 115.9 | 6.70, d, (8.4) |
| 14 | 124.1 | 6.67, dd, (8.4, 1.8) |
| 1′ | 53.9 | 4.57, s |
| 3′ | 38.7 | 3.13, m<br>2.77, m |
| 4′ | 25.9 | 2.95-2.72, m |
| 4a′ | 121.8 | |
| 5′ | 106.0 | 6.53, s |
| 6′ | 147.7 | |
| 7′ | 135.0 | |
| 8′ | 141.5 | |
| 8a′ | 120.7 | |
| α′ | 42.2 | 3.31, m<br>2.94, m |
| 9′ | 138.1 | |
| 10´ | 131.3 | 6.79, m |
| 11′ | 121.0 | 6.19, m |
| 12′ | 152.3 | |
| 13′ | 122.4 | 6.96, m |
| 14′ | 128.8 | 7.58, m |
| 6-$OCH_3$ | 55.9 | 3.57, s |
| 6′-$OCH_3$ | 56.4 | 3.71, s |

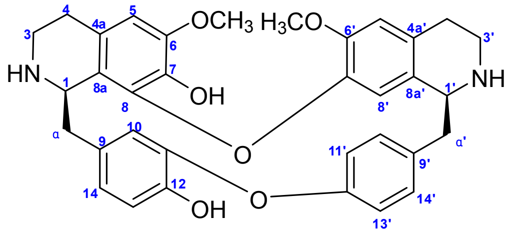


***(R,S)*-17**

**Supplementary Table 5. $^{1}H$ and $^{13}C$ NMR data of (*R*,*S*)-17.**

| Pos. | $\delta_C$, type | $\delta_H$, mult. ($J$ in Hz) |
|---|---|---|
| 1 | 56.6 | 4.70, s |
| 3 | 38.4 | 3.52, m<br>3.44, m |
| 4 | 26.2 | 3.24, m<br>3.15, m |
| 4a | 128.8 | |
| 5 | 112.9 | 6.30, s |
| 6 | 152.4 | |
| 7 | 145.3 | |
| 8 | 117.2 | |
| 8a | 117.4 | |
| α | 38.5 | 3.24, m<br>2.82, m |
| 9 | 133.5 | |
| 10 | 116.5 | 6.65, m |
| 11 | 150.2 | |
| 12 | 147.3 | |
| 13 | 124.5 | 6.65, m |
| 14 | 112.9 | 6.83, overlapped |
| 1′ | 60.8 | 4.51, s |
| 3′ | 49.5 | 3.80, overlapped<br>3.44, m |
| 4′ | 25.3 | 3.10, m<br>2.82, m |
| 4a′ | 133.5 | |
| 5′ | 106.5 | 6.51, s |
| 6′ | 150.3 | |
| 7′ | 136.7 | |
| 8′ | 120.6 | 6.14, s |
| 8a′ | 117.2 | |
| α′ | 37.9 | 3.24, m<br>2.97, m |
| 9′ | 131.8 | |
| 10′ | 117.2 | 6.84, overlapped |
| 11′ | 124.4 | 6.79, dd, (8.0, 2.0) |
| 12′ | 156.6 | |
| 13′ | 124.4 | 7.15, m |
| 14′ | 131.8 | 7.50, d, (8.0) |
| 6-$OCH_3$ | 56.3 | 3.63, s |
| 6′-$OCH_3$ | 56.6 | 3.80, overlapped |

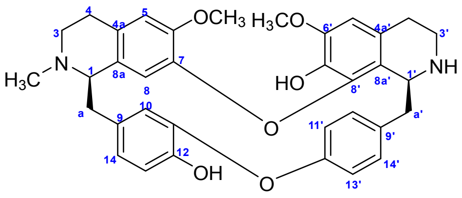


**(*R*,*S*)-18**

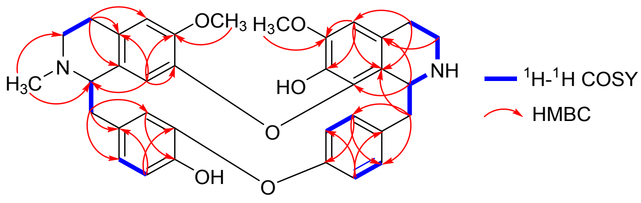


**(*R*,*S*)-18**

**Supplementary Table 6. [1]H and [13]C NMR data of (*R*,*S*)-18.**

| Pos. | $\delta_C$, type | $\delta_H$, mult. (*J* in Hz) |
|---|---|---|
| 1 | 63.7 | 3.45, m |
| 3 | 51.0 | 2.68, m<br>2.27, m |
| 4 | 28.4 | 3.32, m<br>2.27, m |
| 4a | 130.3 | |
| 5 | 111.2 | 6.46, s |
| 6 | 147.9 | |
| 7 | 143.9 | |
| 8 | 116.8 | 6.61, overlapped |
| 8a | 128.8 | |
| α | 37.0 | 2.97, m<br>2.72, m |
| 9 | 130.3 | |
| 10 | 116.5 | 5.28, s |
| 11 | 147.7 | |
| 12 | 143.8 | |
| 13 | 115.1 | 6.61, overlapped |
| 14 | 124.2 | 6.61, overlapped |
| 1′ | 53.0 | 4.69, br d |
| 3′ | 40.8 | 3.40, m<br>2.95, m |
| 4′ | 24.5 | 3.01, m<br>2.89, m |
| 4a′ | 127.8 | |
| 5′ | 105.6 | 6.50, s |
| 6′ | 147.3 | |
| 7′ | 134.8 | |
| 8′ | 141.5 | |
| 8a′ | 118.7 | |
| α′ | 36.4 | 3.42, m |
| 9′ | 136.3 | |
| 10′ | 131.1 | 6.90, m |
| 11′ | 120.8 | 6.25, br d |
| 12′ | 152.6 | |
| 13′ | 122.1 | 6.95, m |
| 14′ | 128.5 | 7.60, m |
| 6-$OCH_3$ | 55.3 | 3.57, s |
| 6′-$OCH_3$ | 56.1 | 3.71, s |
| 2-$NCH_3$ | 43.6 | 2.45, s |

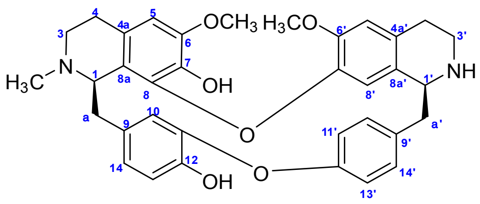


***(R,S)*-19**

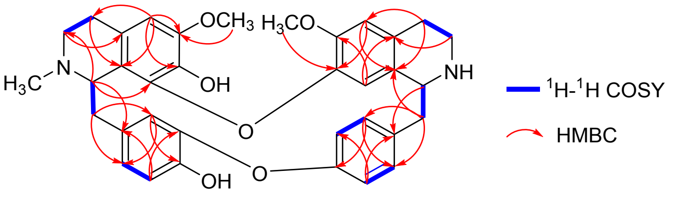


***(R,S)*-19**

**Supplementary Table 7. [1]H and [13]C NMR data of (*R*,*S*)-19.**

| Pos. | $\delta_C$, type | $\delta_H$, mult. ($J$ in Hz) |
|---|---|---|
| 1 | 56.6 | 4.70, m |
| 3 | 38.7 | 3.80, m<br>3.47, m |
| 4 | 25.4 | 3.23, m<br>3.11, m |
| 4a | 130.9 | |
| 5 | 112.9 | 6.82, s |
| 6 | 152.4 | |
| 7 | 145.4 | |
| 8 | 142.3 | |
| 8a | 118.1 | |
| α | 38.8 | 3.47, m<br>3.12, m |
| 9 | 136.5 | |
| 10 | 114.3 | 6.36, s |
| 11 | 150.0 | |
| 12 | 147.3 | |
| 13 | 125.0 | 6.83, m |
| 14 | 112.9 | 6.82, m |
| 1′ | 61.0 | 4.55, m |
| 3′ | 46.7 | 3.80, m<br>3.66, m |
| 4′ | 23.8 | 3.01, m<br>2.74, m |
| 4a′ | 136.5 | |
| 5′ | 106.2 | 6.52, s |
| 6′ | 150.3 | |
| 7′ | 149.9 | |
| 8′ | 120.3 | 6.06, s |
| 8a′ | 116.2 | |
| α′ | 37.7 | 3.24, m<br>2.91, m |
| 9′ | 133.7 | |
| 10′ | 117.2 | 6.83, m |
| 11′ | 123.1 | 6.57, dd, (8.0, 2.4) |
| 12′ | 157.0 | |
| 13′ | 123.1 | 7.14, d, (8.0) |
| 14′ | 131.6 | 7.49, d, (8.0) |
| 6-$OCH_3$ | 56.3 | 3.63, s |
| 6′-$OCH_3$ | 56.7 | 3.82, s |
| 2-$NCH_3$ | 42.4 | 2.66, s |

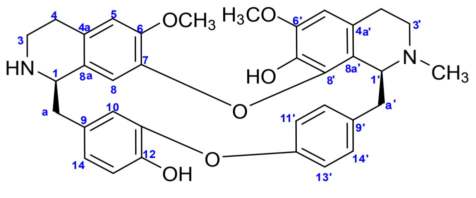


(*R*,*S*)-**35**

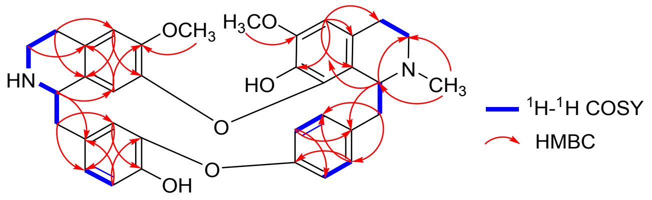


(*R*,*S*)-**35**

**Supplementary Table 8. $^{1}$H and $^{13}$C NMR data of (*R*,*S*)-35.**

| Pos. | $\delta_C$, type | $\delta_H$, mult. ($J$ in Hz) |
|---|---|---|
| 1 | 53.3 | 4.36, br s |
| 3 | 39.4 | 2.91, m<br>2.58, m |
| 4 | 26.4 | 2.55, m<br>2.43, m |
| 4a | 127.8 | |
| 5 | 112.0 | 6.56, s |
| 6 | 148.9 | |
| 7 | 144.9 | |
| 8 | 123.3 | 6.73, s |
| 8a | 124.7 | |
| α | 37.9 | 3.18, m<br>2.89, m |
| 9 | 126.6 | |
| 10 | 115.7 | 5.40, m |
| 11 | 148.2 | |
| 12 | 144.6 | |
| 13 | 116.3 | 6.73, m |
| 14 | 123.5 | 6.73, m |
| 1′ | 61.0 | 4.07, br s |
| 3′ | 44.6 | 3.08, m<br>2.83, m |
| 4′ | 23.9 | 2.89, m<br>2.58, m |
| 4a′ | 123.5 | |
| 5′ | 105.7 | 6.42, s |
| 6′ | 146.5 | |
| 7′ | 134.5 | |
| 8′ | 141.6 | |
| 8a′ | 122.5 | |
| α′ | 39.8 | 3.22, m<br>2.72, m |
| 9′ | 139.2 | |
| 10′ | 131.2 | 6.82, d, (8.0) |
| 11′ | 120.5 | 6.22, d, (8.0) |
| 12′ | 151.4 | |
| 13′ | 122.6 | 6.93, overlapped |
| 14′ | 128.1 | 7.40, overlapped |
| 6-$OCH_3$ | 55.9 | 3.70, s |
| 6′-$OCH_3$ | 48.9 | 3.58, s |
| 2′-*N*$CH_3$ | 41.6 | 2.58, s |

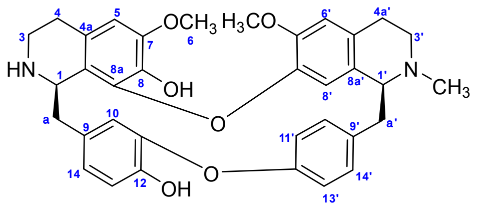


(*R*,*S*)-**36**

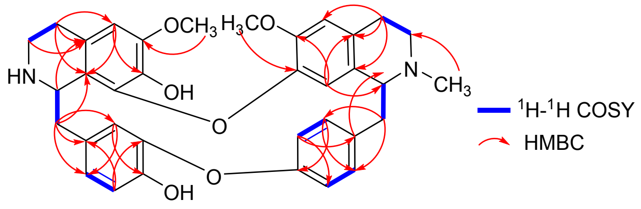


(*R*,*S*)-**36**

**Supplementary Table 9. $^{1}$H and $^{13}$C NMR data of (*R,S*)-36.**

| Pos. | $\delta_C$, type | $\delta_H$, mult. ($J$ in Hz) |
|---|---|---|
| 1 | 56.8 | 4.53, m |
| 3 | 38.4 | 3.60, m<br>3.33, m |
| 4 | 26.0 | 3.00, m<br>2.93, m |
| 4a | 136.7 | |
| 5 | 106.3 | 6.51, s |
| 6 | 150.3 | |
| 7 | 142.7 | |
| 8 | 146.3 | |
| 8a | 116.0 | |
| α | 37.8 | 3.28, m<br>2.83, m |
| 9 | 129.9 | |
| 10 | 121.3 | 6.12, m |
| 11 | 150.2 | |
| 12 | 147.4 | |
| 13 | 117.2 | 6.84, d (8.0) |
| 14 | 124.4 | 6.80, dd (8.0, 2.4) |
| 1′ | 64.6 | 4.51, m |
| 3′ | 46.5 | 3.84, m<br>3.36, m |
| 4′ | 24.8 | 3.22, m<br>3.15, m |
| 4a′ | 128.0 | |
| 5′ | 112.7 | 6.81, s |
| 6′ | 152.5 | |
| 7′ | 145.1 | |
| 8′ | 117.2 | 6.39, s |
| 8a′ | 124.4 | |
| α′ | 36.5 | 3.51, m<br>3.06, m |
| 9′ | 134.0 | |
| 10′ | 133.7 | 6.57, d, (8.0) |
| 11′ | 123.0 | 6.30, d, (8.0) |
| 12′ | 156.5 | |
| 13′ | 122.8 | 7.17, d, (8.0) |
| 14′ | 131.8 | 7.49, d, (8.0) |
| 6-$OCH_3$ | 56.2 | 3.60, s |
| 6′-$OCH_3$ | 56.6 | 3.79, s |
| 2′-$NCH_3$ | 41.6 | 2.92, s |

**Supplementary Table 10. Strains and plasmids used in this study**

| Strain ID | Genotype | Source |
| --- | --- | --- |
| IMX581 | MATa ura3-52 can1Δ::cas9-natNT2 TRP1 LEU2 HIS3 | [6] |
| XJ0636 | IMX581 (X-2:: TPI1p-EcaroL-pYX212t + ADH1t-ARO7*-PGK1p + TEF1p-ARO4*-CYC1t + FBA1t-MtPDH1-tHXT7p X-4:: CYC1t-ARO1-TPI1 + TDH3p-ARO2-ADH1t + TDH2t-ARO3-TEF1p XI-5:: IDP1t-CYP76AD5-TDH3p + CCW12p-DODC-TPS1t XI-3:: TEF1p-CjNCSΔ35-TDH2t ΔARI1:: LP3.T7 ΔADH6:: LP3.T7 ΔYPR1:: LP3.T7 ΔHFD1:: LP4.T9 ΔGRE2:: TDH3p-CjNCSΔ35-CYC1t XI-1:: TDH3p-CjNCSΔ35-CYC1t X-3:: FBA1p-PsAAAD*-TPS1t-ADH1t 106a:: FBA1p-AtPAT-TDH2t + pYX212t-MtncADH-TEF2p) | Our previous study |
| XJ06360 | XJ0636 (XII-2:: GPM1p-Ps6OMT-FBA1t) | This study |
| XJ06385 | XJ06360 (hfd1:: GPD-PrDCS-IDP1t + CCW12p-NCP1-CYC1t) | This study |
| XJ06388 | XJ06385 (416d:: GPD-CYP80Q4-IDP1t + CCW12p-PbDCR-CYC1t) | This study |
| XJ06389 | XJ06388 (XII-4:: CCW12p-CYP82BC1-PGI1t) | This study |
| XJ06390 | XJ06388 (XII-4:: CCW12p-CYP82BC2-PGI1t) | This study |
| XJ06391 | XJ06388 (XII-4:: CCW12p-CYP82BC3-PGI1t) | This study |
| XJ06392 | XJ06388 (XII-4:: CCW12p-CYP82BC4-PGI1t) | This study |
| XJ06398 | XJ06388 (ΔARI1:: GPD-PrDCS-IDP1t + CCW12p-PbDCR-CYC1t ΔADH6:: GPD-PrDCS-IDP1t + CCW12p-PbDCR-CYC1t ΔYPR1:: GPD-PrDCS-IDP1t + CCW12p-PbDCR-CYC1t) | This study |
| XJ06399 | XJ06398 (ΔYDR541c:: GPD-CYP80Q4-IDP1t + CCW12p-PbDCR-CYC1t) | This study |
| XJ06320 | XJ06399 (XI-2:: PDC1p-B2NMT-ENO2t + IDP1t-B12OMT-GPM1p + TPI1p-B7OMT-ADH1t + FBA1t-B2′NMT2-PGK1p) | This study |
| XJ06321 | XJ06399 (XII-4:: CCW12p-CYP82BC3-PGI1t) | This study |
| XJ06322 | XJ06399 (XII-4:: CCW12p-CYP82BC1-PGI1t) | This study |
| XJ06325 | XJ06320 (XII-4:: CCW12p-CYP82BC3-PGI1t) | This study |
| XJ06326 | XJ06320 (XII-4:: CCW12p-CYP82BC1-PGI1t) | This study |
| XJ06327 | XJ06320 (XII-4:: CCW12p-CYP82BC2-PGI1t) | This study |
| XJ06328 | XJ06320 (XII-4:: CCW12p-CYP82BC4-PGI1t) | This study |
| XJ06354 | XJ06322 (FgF20:: GPD-B2NMT-CYC1t) | This study |
| XJ06357 | XJ06322 (XII-1:: GPD-B12OMT-CYC1t) | This study |
| XJ06358 | XJ06322 (XII-1:: GPD-B7OMT-CYC1t) | This study |
| XJ06359 | XJ06322 (XII-1:: GPD-B2′NMT2-CYC1t) | This study |
| XJ06001 | XJ06322 (XII-1:: GPD-B12OMT-IDP1t + CCW12p-B7OMT-CYC1t) | This study |
| XJ06002 | XJ06322 (XII-1:: GPD-B12OMT-IDP1t + CCW12p-B2′NMT2-CYC1t) | This study |
| XJ06003 | XJ06322 (XII-1:: GPD-B2′NMT2-IDP1t + CCW12p-B7OMT-CYC1t) | This study |
| XJ06004 | XJ06322 (FgF20:: IDP1t-B12OMT-GPM1p + TPI1p-B7OMT-ADH1t + FBA1t-B2′NMT2-PGK1p) | This study |
| XJ06005 | XJ06354 (XII-1:: GPD-B12OMT-CYC1t) | This study |
| XJ06006 | XJ06354 (XII-1:: GPD-B7OMT-CYC1t) | This study |
| XJ06007 | XJ06354 (XII-1:: GPD-B2′NMT2-CYC1t) | This study |
| XJ06008 | XJ06354 (XII-1:: GPD-B12OMT-IDP1 + CCW12p-B7OMT-CYC1t) | This study |
| XJ06009 | XJ06354 (XII-1:: GPD-B12OMT-IDP1t + CCW12p-B2′NMT2-CYC1t) | This study |
| XJ06010 | XJ06354 (XII-1:: GPD-B2′NMT2-IDP1t + CCW12p-B7OMT-CYC1t) | This study |
| XJ06011 | XJ06321 (XII-1:: GPD-BBOX-CYC1t) | This study |
| XJ06016 | XJ06325 (XII-1:: GPD-BBOX-CYC1t) | This study |
| XJ06017 | XJ06325 (XII-1:: GPD-BBOX-IDP1t + CCW12p-BBR-CYC1t) | This study |
| XJ06019 | XJ06321 (FgF20:: IDP1t-B12OMT-GPM1p + TPI1p-B7OMT-ADH1t + | This study |

| | | |
|---|---|---|
| | FBA1t-B2′NMT2-PGK1p) | |
| XJ06020 | XJ06019 (XII-1:: GPD-BBOX-CYC1t) | This study |
| XJ06021 | XJ06019 (XII-1:: GPD-BBOX-IDP1t + CCW12p-BBR-CYC1t) | This study |
| Plasmid | Genotype | Source |
| pMEL10-XI-2 | 2μm, AmpR, KlURA3, SNR52p-XI-2-gRNA | Our lab |
| pMEL10-XII-1 | 2μm, AmpR, KlURA3, SNR52p-XII-1-gRNA | Our lab |
| pMEL10-XII-2 | 2μm, AmpR, KlURA3, SNR52p-XII-2-gRNA | Our lab |
| pMEL10-XII-4 | 2μm, AmpR, KlURA3, SNR52p-XII-4-gRNA | Our lab |
| pMEL10-LP3.T7 | 2μm, AmpR, KlURA3, SNR52p-LP3.T7-gRNA | This study |
| pMEL10-416d | 2μm, AmpR, KlURA3, SNR52p-416d-gRNA | This study |
| pMEL10-YDR541C | 2μm, AmpR, KlURA3, SNR52p-YDR541C-gRNA | This study |
| pMEL10-FgF20 | 2μm, AmpR, KlURA3, SNR52p-FgF20-gRNA | This study |

**Supplementary Table 11. List of primer pairs used to clone genes from *S. tetrandra***

| Target | Use | Primer sequence (5'–3') |
|---|---|---|
| *CYP80Q4* | Fwd | AGGAGAAAAAACCCCGGATCCATGGATCCAATCACACTGTTAC |
| | Rev | GAGTCGTATTACGGATCCTTAAACCGCAGTTGCTGAT |
| *CYP82BC1* | Fwd | AGGAGAAAAAACCCCGGATCCATGATCCCGCAACCTTACT |
| | Rev | GAGTCGTATTACGGATCCTCAAGGAGAAGGAATCACTC |
| *CYP82BC2* | Fwd | AGGAGAAAAAACCCCGGATCCATGATGATCGAGATCCAAAATTTCT |
| | Rev | GAGTCGTATTACGGATCCTCAGGAAGAAGGAAGCCG |
| *CYP82BC3* | Fwd | AGGAGAAAAAACCCCGGATCCATGATCGAGATCCAAAACTTCTT |
| | Rev | GAGTCGTATTACGGATCCTCAAGGAGAAGGAAGTCTCG |
| *CYP82BC4* | Fwd | AGGAGAAAAAACCCCGGATCCATGATCCCCCAACCTTACTT |
| | Rev | GAGTCGTATTACGGATCCTCAAGGAGAAGGAATCACTCG |
| *B12OMT* | Fwd | GCCATGGCTGATATCGGATCCATGGATTCCATTATTCGTGATCTA |
| | Rev | ACGGAGCTCGAATTCGGATCCTTATTTGAGAAATTCAATGACCCAC |
| *B7OMT* | Fwd | GCCATGGCTGATATCGGATCCATGGAGAAGTGTGAGAGGAA |
| | Rev | ACGGAGCTCGAATTCGGATCCTTATGGGTACACGACTAGAACA |
| *B2NMT* | Fwd | GCCATGGCTGATATCGGATCCATGACTAACAACGTTGGTTGT |
| | Rev | ACGGAGCTCGAATTCGGATCCTTATTTCTTTTTAAATAGAAAATGA |
| *B2'NMT1* | Fwd | GCCATGGCTGATATCGGATCCATGCATATGGATTCCGCTAATG |
| | Rev | ACGGAGCTCGAATTCGGATCCTCATTTGAGAAATTCTATGACCCA |
| *B2'NMT2* | Fwd | GCCATGGCTGATATCGGATCCATGAGTTCCAATATTGTGCACAAAC |
| | Rev | ACGGAGCTCGAATTCGGATCCTCATTTGAGAAATTCGATGACCCA |
| *BBOX* | Fwd | AGGAGAAAAAACCCCGGATCCATGGAGGGATGGGGC |
| | Rev | GAGTCGTATTACGGATCCTCAGTACGTGCTTTCCATTG |
| *BBR* | Fwd | GCCATGGCTGATATCGGATCCATGTGTTACCATCAACATTTCAACAT |
| | Rev | ACGGAGCTCGAATTCGGATCCTTATAGAAACCATTTCAAGTATTCCTC AACT |

**Supplementary Table 12. Primers used in the construction of engineered strains in this study**

| Primer name | Sequence (5′–3′) |
|---|---|
| XII-2 us-F | cggcatgcaaacatctacac |
| GPM1p-Ps6OMT-R | tgtttttattgtaatatgtgtg |
| GPM1p-Ps6OMT-F | aataatccaaacaaacacacatattacaataaaaacaatggaaactgtctcaaaaatcg |
| Ps6OMT-FBA1t-R | actcattaaaaaactatatcaattaatttgaattaacttagtaagggtaggcttctatg |
| Ps6OMT-FBA1t-F | gttaattcaaattaattgatat |
| FBA1t-XII-2 ds-R | cagtacaaggacgcgttaagaaaaatttcgagagagtcgccgatagtagagtaagctactatgaaagact |
| XII-2 ds-F | ctactatcggcgactctctcgaa |
| XII-2 ds-R | gagcgaacgtaagagaggttaatg |
| hfd1-us-F | caatgtcccagttatacgaatact |
| hfd1 us-LP4.T9-R | actctgtattgcgagctccattgtatggagtatccgtttatcccta |
| LP4.T9-GPD-F | atactccatacaatggagctcgcaatacagagtttaccgcatcttgcccttcattatcaatactcgccatt |
| GPD-p416-R | tcgaaactaagttctggtgttttaa |
| GPD-DRS_DRR-F | tctttttttagttttaaaacaccagaacttagtttcgaatggagtacagcgatttacaa |
| DRS-His-IDP1t-R | atgaaaaaaaaaagtggtagattgggctacgtaaattcgattaatggtgatggtgatgatgcaacattcttcttgttagc |
| IDP1t-r-v-R | tcgaatttacgtagcccaatcta |
| IDP1t-r-F | gatggtaatgatccgaacttgg |
| IDP1t-CCW12p-F | atcaaggttccccaagttcggatcattaccatcccacccatgaaccacacggttag |
| CCW12p-DODC-R | cattttgtttattgatatagtg |
| CCW12p-NCP1-F | tcttctgtcattcgcttaaacactatatcaataaacaaaatgccgtttggaatagacaac |
| NCP1-CYC1t-R | gagggcgtgaatgtaagcgtgacataactaattacatgattaccagacatcttcttggta |
| CYC1t-p416-F | tcatgtaattagttatgtcacgctta |
| CYC1t-LP4.T9-R | aacattacgtgtgcaacttcggctaaccatgttatctaaaaaagttgagggcaaattaaagccttcgagcg |
| LP4.T9-hfd1 ds-F | aacatggttagccgaagttgcacacgtaatgtttaaggttaattaa |
| hfd1-ds-R | gtttgaagcggcttacgattttc |
| 416d us-GAP-F | gttgataattagcgttgcctcatcaatgcgagatccgtttaaccggaccctcattatcaatactcgccatttc |
| GPD-CYP80Q4-F | tctttttttagttttaaaacaccagaacttagtttcgaatggaccctattacattattg |
| IDP1t-CYP80Q4-F | aaaaaaaagtggtagattgggctacgtaaattcgattaatggtgatggtgatgatgaacagcagtagcagaagatc |
| CCW12p-DRR-R | tcttctgtcattcgcttaaacactatatcaataaacaaaatggaatcttctggtgttcca |
| DRS_DRR-His-CYC1t-R | gcgtgaatgtaagcgtgacataactaattacatgattaatggtgatggtgatgatgagccttatcatcccataatt |
| CYC1t-416d ds-R | gaattccacatgttaaaatagtgaaggagcatgttcggcacacagtggacgcaaattaaagccttcgagcgtc |
| XII-4 us-F | gtatccggctgttccttcatagcc |
| XII-4 us-CCW12p-R | gccccttttggactaaccgtgtggttcatgggtggtgccatagtatgtgtgatggaa |
| CCW12p1-F | ccacccatgaaccacacggttag |
| CCW12p-CYP82BC3-F | tcttctgtcattcgcttaaacactatatcaataaacaaaatgatcgaaatccaaaatttc |
| CYP82BC3-PGI1t-R | gatttgtttacagatcctcttctgagatgagtttttgttctggagatggcaatcttggtg |
| Myc-PGI1t-F | gaacaaaaactcatctcagaagaggatctgtaaacaaatcgctcttaaatatatacc |
| XII-5-PGI1t-r-F | gtagtttagtgtttttcttcca |
| PGI1t-XII-4 ds-F | agatactcgcactggaagaaaaacactaaactacattccccattagagtcaaataa |
| XII-4 ds-R | tttctgccgtacctggatggtcat |
| CCW12p-CYP82BC1-F | atcttctgtcattcgcttaaacactatatcaataaacaaaatgattccacaaccatattt |
| CYP82BC1-PGI1t-R | gatttgtttacagatcctcttctgagatgagtttttgttctggtgatggaataactcttg |
| CCW12p-CYP82BC2-F | atcttctgtcattcgcttaaacactatatcaataaacaaaatgatgatcgaaatccaaaa |
| CYP82BC2-PGI1t-R | gatttgtttacagatcctcttctgagatgagtttttgttctgaagatggcaatcttggag |
| CCW12p- | atcttctgtcattcgcttaaacactatatcaataaacaaaatgattccacaaccatactt |

| | |
|---|---|
| CYP82BC4-F | |
| CYP82BC4-PGI1t-R | gatttgtttacagatcctcttctgagatgagtttttgttctggtgatggaataactcttg |
| LP3.T7-GPD-F | cgcatagacatacaagtggacagatgatgggtacgggcctctaatacatctcattatcaatactcgccatt |
| CYC1t-LP3.T7-R | gaatggtattctgaagctgcacatcgattcactaaagtgtggatgttcccgcaaattaaagccttcgagcg |
| ypr541-GPD-F | ctcttttatttgcatatactttttgtttatgttctatagcctaaattgtatcattatcaatactcgccatt |
| CYC1t-ypr541-R | gcttgattattttggcttcacataattgcgagattatgtgtggtatggcggcaaattaaagccttcgagcg |
| XI-2 us-F | taactcttcgtatgaggattttc |
| XI-2 us-PDC1p-R | acatcagcggaacatatgctcacccagtcgcatgttctatggcacatttttctgttg |
| XI-2 us-PDC1p-F | catgcgactgggtgagcatatgtt |
| PDC1p-B2NMT-F | tcataacctcacgcaaaataacacagtcaaatcaatcaaaatgactaacaatgtcggttg |
| B2NMT-ENO2t-R | aataagcagaaaagactaataattcttagttaaaagcacttcacttcttcttgaataaga |
| ENO2t-IDP1t-r-R | atcaaggttccccaagttcggatcattaccatcaggtatcatctccatctcccata |
| B12OMT-IDP1t-R | atgaaaaaaaaaagtggtagattgggctacgtaaattcgatcatttcaagaattcaataa |
| B12OMT-GPM1p-R | attcttcttaataatccaaacaaacacacatattacaataatggactccatcatcagaga |
| GPM1p-XI-2 ds-r-R | tagtcgtgcaatgtatgactttaa |
| GPM1p-TPI1p-F | tgctcacaaatcttaaagtcatacattgcacgactagtttaaagattacggatatttaa |
| ARO1-TPI1p-r-F | ttttagtttatgtatgtgttt |
| TPI1p-B7OMT-F | ttaaatctataactacaaaaaacacatacataaactaaaaatggaaaagtgtgaaagaaa |
| B7OMT-ADH1t-R | cttatttaataataaaaatcataaatcataagaaattcgctcatgggtagacaactagca |
| ADH1t-DIT1t-r-R | gcatatctacaattgggtgaaat |
| ADH1t-FBA1t-F | cgagcaaatgcctgcaaatcgctccccatttcacccaattgtagatatgccgtaagctactatgaaagactt |
| Ps6OMT-FBA1t-F | gttaattcaaattaattgatat |
| FBA1t-B2′NMT2-F | aatactcattaaaaaactatatcaattaatttgaattaactcacttcaagaattcgataa |
| B2′NMT2-PGK1p-R | aaggaagtaattatctactttttacaacaaatataacaaaatgtcttccaacattgtcca |
| PGK1p-CMT7p-R | tttgttatatttgttgtaaaaagt |
| PGK1p-r-R | acgcacagatattataacatctgc |
| PGK1p-XI-2-F | aaatgcctattgtgcagatgttataatatctgtgcgttacacacgactagcgctttcaga |
| XI-2 ds-R | atggttgaaaaggttacagagg |
| BBOX-IDP1t-R | TGAAAAAAAAAAGTGGTAGATTGGGCTACGTAAATTCGAtcagtaagtactttccataga |
| CCW12p-BBR-F | ATCTTCTGTCATTCGCTTAAACACTATATCAATAAACAAAatgtgttaccatcaacactt |
| AmpR-m-F1 | agccggtgagcgtggatcgcgcggtatcattgcagcact |
| AmpR-m-R1 | gcaatgataccgcgcgatccacgctcaccggctccagat |
| SNR52p-NCS-R | TGAACCTTCAACAGAGGTCTCTgatcatttatctttcactgcggagaag |
| SNR52p-NCS-F | gcagtgaaagataaatgatcAGAGACCTCTGTTGAAGGTTCACCAGAAT |
| NCS-gRNA-R | tattttaacttgctatttctagctctaaaacAGAGACCTGTTCAACCAATTTGTATCTG |
| vector-gRNA-F | accctcactaaagggaacaaaagctggagcttctgttttagagctagaaatagcaag |
| vector backbone-F | GATCATTTATCTTTCACTGCGGAGAAG |
| vector backbone-R | GTTTTAGAGCTAGAAATAGCAAGTTAAAATAAGGCTAGTC |
| BsaI-ari1-gRNA-F | AAAGGTCTCAGATCAATTAGCATAGGATTTTCCGgttttagagctagaaatagcaag |
| SNR52p-adh6-gRNA-R | AAAGGTCTCACGAGGTGAAGTGgatcatttatctttcactgcgga |
| SNR52p-adh6-gRNA-F | AAAGGTCTCACTCGAGAACTGTgttttagagctagaaatagcaag |
| SNR52p-ypr1-gRNA-R | AAAGGTCTCTaaacCCATTCCAGTGTTGGGTTTCgatcatttatctttcactgcgga |
| BsaI-ydc541c-gRNA-F | AAAGGTCTCAGATCGGCACCCAAAATAGATAGAGgttttagagctagaaatagcaag |

| | |
|---|---|
| SNR52p-aad3-gRNA-R | AAAGGTCTCTAAACCACCTATTACATTCCGCCTGgatcatttatctttcactgcgga |
| BsaI-gre2-gRNA-F | AAAGGTCTCAGATCTAGAATACGGAATTTTCTCGgttttagagctagaaatagcaag |
| SNR52p-hfd1-gRNA-R | AAAGGTCTCTaaacTTTAGATAATCAATATGCCTgatcatttatctttcactgcgga |
| LP3.T7-gRNA-F | tctccgcagtgaaagataaatgatcATTGTACCCCAGCGGCGGCGgttttagagctagaaatagcaagtt |
| LP3.T7-gRNA-R | aacttgctatttctagctctaaaacCGCCGCCGCTGGGGTACAATgatcatttatctttcactgcggaga |
| 416d-gRNA-F | gcagtgaaagataaatgatcTAGTGCACTTACCCCACGTTgttttagagctagaaatagc |
| 416d-gRNA-R | gctatttctagctctaaaacAACGTGGGGTAAGTGCACTAgatcatttatctttcactgc |
| FgF20-gRNA-F | gcagtgaaagataaatgatcGTTAGAGCTGTTACAAGTTAgttttagagctagaaatagc |
| FgF20-gRNA-R | gctatttctagctctaaaacTAACTTGTAACAGCTCTAACgatcatttatctttcactgc |

### Synthesis of compounds

synthesis of **lindoldhamine stereoisomers**

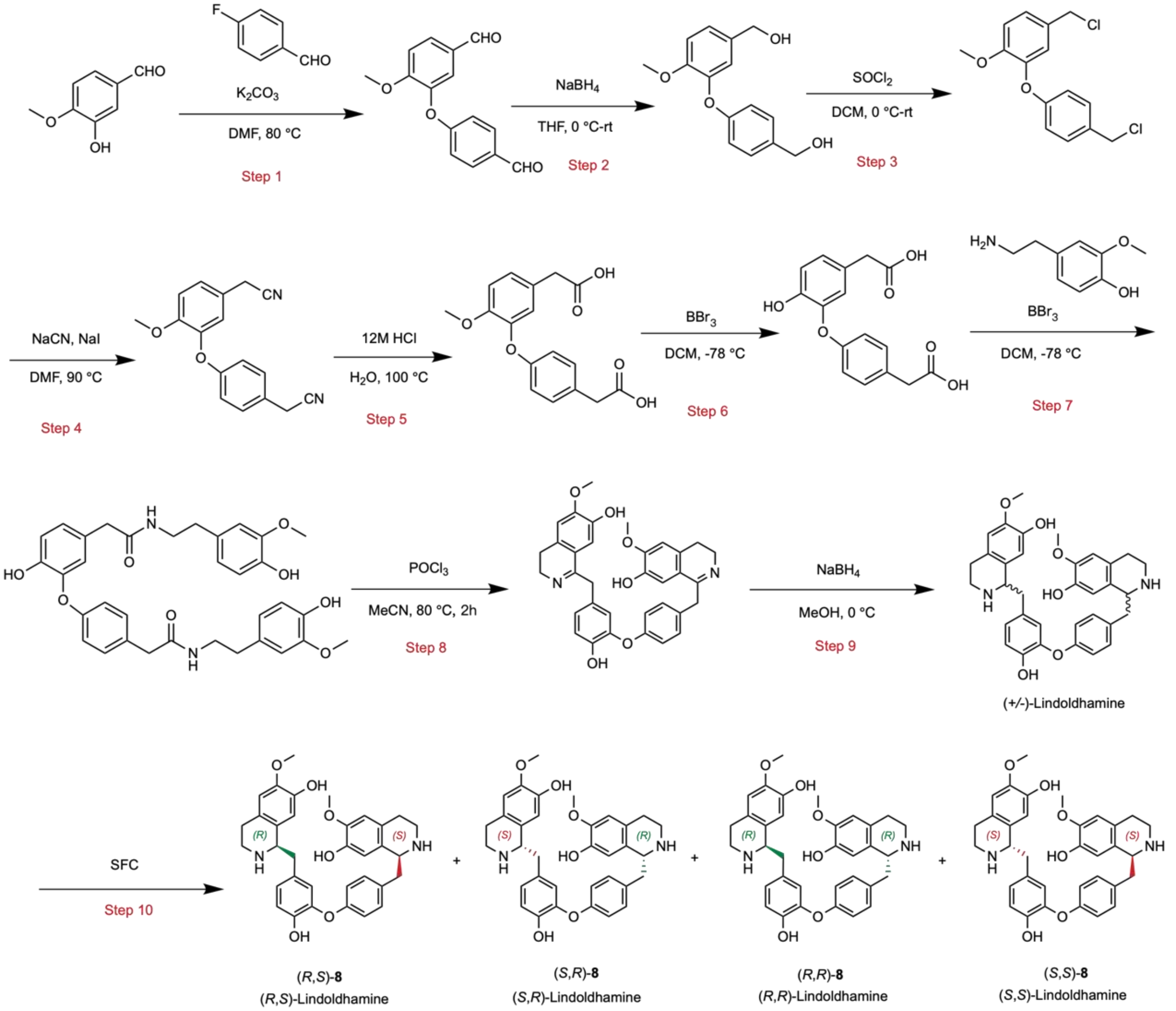


**Step 1. 3-(4-formylphenoxy)-4-methoxybenzaldehyde**

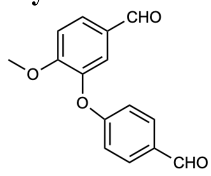

To a solution of 3-hydroxy-4-methoxybenzaldehyde (50 g, 0.33 mol, 1.0 equiv.) in DMF (200 mL) were added $K_2CO_3$ (113.54 g, 0.82 mol, 2.5 equiv.) and 4-fluorobenzaldehyde (40.78 g, 0.33 mol, 1.0 equiv.). The reaction mixture was stirred at 80 ℃ for 16 h. After completion, the mixture was diluted with water and extracted with EtOAc (800 mL × 3). The combined organic phases were washed with brine, dried over $Na_2SO_4$, filtered, and concentrated. The residue was purified by flash column chromatography (petroleum ether/ethyl acetate = 85:15) to afford 3-(4-formylphenoxy)-4-methoxybenzaldehyde (75 g, 80.2% yield) as an off-white solid.

**Step 2. (4-(5-(hydroxymethyl)-2-methoxyphenoxy)phenyl)methanol**

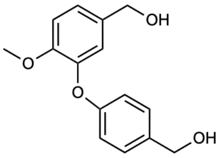

To a solution of 3-(4-formylphenoxy)-4-methoxybenzaldehyde (75 g, 0.29 mol, 1.0 equiv.) in THF (700 mL) was added $NaBH_4$ (16.6 g, 0.44 mol, 1.5 equiv.) portionwise at 0 ℃. The mixture was warmed to 25 ℃ and stirred for 3 h. After completion, the reaction was quenched with water and extracted with EtOAc (500 mL × 3). The combined organic phases were washed with brine, dried over $Na_2SO_4$, filtered, and concentrated under reduced pressure to afford (4-(5-(hydroxymethyl)-2-methoxyphenoxy)phenyl)methanol (70 g, 82.7% yield) as an off-white solid.

**Step 3. 4-(chloromethyl)-2-(4-(chloromethyl)phenoxy)-1-methoxybenzene**

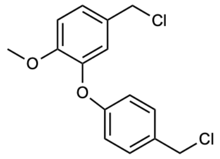


To a solution of (4-(5-(hydroxymethyl)-2-methoxyphenoxy)phenyl)methanol (70 g, 0.27 mol, 1.0 equiv.) in DCM (700 mL) at 0 ℃ was added $SOCl_2$ (63.98 g, 0.54 mol, 2.0 equiv.). The reaction mixture was stirred at 25 ℃ for 16 h. After completion, the solvent was removed under reduced pressure, and the residue was purified by flash column chromatography (petroleum ether/ethyl acetate = 95:5) to afford 4-(chloromethyl)-2-(4-(chloromethyl)phenoxy)-1-methoxybenzene (70 g, 78.8% yield) as an off-white solid.

**Step 4. 2-(4-(5-(cyanomethyl)-2-methoxyphenoxy)phenyl)acetonitrile**

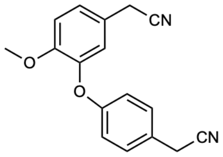


To a mixture of 4-(chloromethyl)-2-(4-(chloromethyl)phenoxy)-1-methoxybenzene (30 g, 0.10 mol) in DMF (250 mL) was added sodium cyanide (10.89 g, 0.22 mol) and sodium iodide (3.03 g, 0.02 mol), the mixture was stirred at 90 °C for 16 h. After completion, the mixture was diluted with water and extracted with EtOAc (500 mL × 3). The combined organic phases were washed with brine, dried over $Na_2SO_4$, filtered and concentrated. The residue was purified by flash column chromatography (PE/EA = 3:1) to afford 2-(4-(5-(cyanomethyl)-2-methoxyphenoxy)phenyl)acetonitrile (20 g, 64.1% yield) as an off-white solid.

**Step 5. 2-(4-(5-(carboxymethyl)-2-methoxyphenoxy)phenyl)acetic acid**

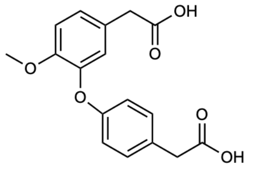


A solution of 2-(4-(5-(cyanomethyl)-2-methoxyphenoxy)phenyl)acetonitrile (20 g, 0.072 mol, 1.0 equiv.) in concentrated HCl (12 M, 400 mL) was stirred at 100 ℃ for 4 h. After completion, the mixture was diluted with water (1000 mL) and extracted with EtOAc (800 mL × 3). The combined organic phases were washed with brine (500 mL x 2), dried over $Na_2SO_4$, filtered, and concentrated. The residue was purified by flash column chromatography (DCM/MeOH = 10:1) to afford 2-(4-(5-(carboxymethyl)-2-methoxyphenoxy)phenyl)acetic acid (15 g, 59.4% yield) as a yellow solid.

**Step 6. 2-(4-(5-(carboxymethyl)-2-hydroxyphenoxy)phenyl)acetic acid**

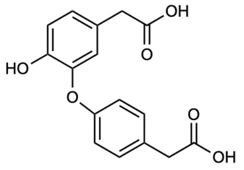


To a solution of 2-(4-(5-(carboxymethyl)-2-methoxyphenoxy)phenyl)acetic acid (15 g, 0.047 mol, 1.0 equiv.) in DCM (200 mL) was added $BBr_3$ (189.6 mL, 0.19 mol, 4.0 equiv., 1 M in DCM) at -78 ℃ under $N_2$ atmosphere, then it was allowed to warm to 25 ℃ and stirred for 2 h. After completion, the mixture was quenched with water (200 mL) and extracted with ethyl acetate (800 mL × 3), the organic layer was washed with brine (500 mL × 3), dried over $Na_2SO_4$, filtered and concentrated. The residue was purified by flash column chromatography (DCM/MeOH=10/1) to afford 2-(4-(5-(carboxymethyl)-2-hydroxyphenoxy)phenyl)acetic acid (10.2 g, 64.1% yield) as a yellow solid.

**Step 7. 2-(4-hydroxy-3-(4-(2-((4-hydroxy-3-methoxyphenethyl)amino)-2-oxoethyl)phenoxy)phenyl)-N-(4-hydroxy-3-methoxyphenethyl)acetamide**

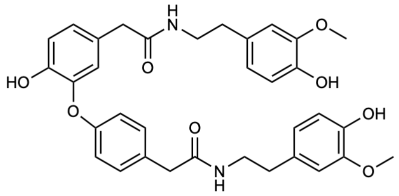

To a solution of 2-(4-(5-(carboxymethyl)-2-hydroxyphenoxy)phenyl)acetic acid (10.2 g, 0.034 mol, 1.0 equiv.) in DMF (100 mL) were added 4-(2-aminoethyl)-2-methoxyphenol (22.54 g, 0.13 mol, 3.8 equiv.), EDCI (16.8 g, 0.088 mol, 2.6 equiv.), HOBt (11.84 g, 0.088 mol, 2.6 equiv.), and DIEA (26.13 g, 0.20 mol, 5.9 equiv.). The reaction mixture was stirred at 25 ℃ for 16 h under $N_2$. After completion, the mixture was diluted with water (1000 mL) and extracted with EtOAc (1000 mL × 3). The combined organic phases were washed with brine (500 mL × 3), dried over $Na_2SO_4$, filtered, and concentrated. The residue was purified by flash column chromatography (DCM/MeOH = 10:1) to afford 2-(4-hydroxy-3-(4-(2-((4-hydroxy-3-methoxyphenethyl)amino)-2-oxoethyl)phenoxy)phenyl)-N-(4-hydroxy-3-methoxyphenethyl)acetamide (17.0 g, 75.7% yield) as a grey solid.

**Step 8. 1-(4-hydroxy-3-(4-((7-hydroxy-6-methoxy-3,4-dihydroisoquinolin-1-yl)methyl)phenoxy)benzyl)-6-methoxy-3,4-dihydroisoquinolin-7-ol**

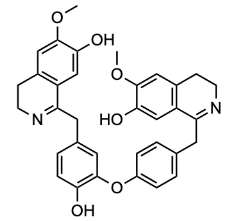

To a solution of 2-(4-hydroxy-3-(4-(2-((4-hydroxy-3-methoxyphenethyl)amino)-2-oxoethyl)phenoxy)phenyl)-N-(4-hydroxy-3-methoxyphenethyl)acetamide (12.1 g, 0.020 mol, 1.0 equiv.) in MeCN (200 mL) was added $POCl_3$ (30.82 g, 0.20 mol, 10 equiv.). The reaction mixture was stirred at 80 ℃ for 2 h. After completion, the solvent was removed under reduced pressure, and the residue was purified by flash column chromatography (DCM/MeOH = 10:1) to afford 1-(4-hydroxy-3-(4-((7-hydroxy-6-methoxy-3,4-dihydroisoquinolin-1-yl)methyl)phenoxy)benzyl)-6-methoxy-3,4-dihydroisoquinolin-7-ol (9.0 g, 71.1% yield) as a yellow solid.

**Step 9. Mixture of lindoldhamine stereoisomers**

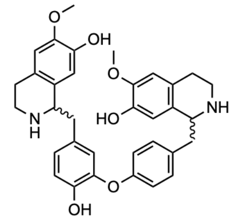

To a solution of 1-(4-hydroxy-3-(4-((7-hydroxy-6-methoxy-3,4-dihydroisoquinolin-1-yl)methyl)phenoxy)benzyl)-6-methoxy-3,4-dihydroisoquinolin-7-ol (9.0 g, 0.016 mol, 1.0 equiv.) in MeOH (150 mL) was added $NaBH_4$ (1.2 g, 0.032 mol, 2.0 equiv.) at 0 ℃. The reaction mixture was stirred at 0 ℃ for 2 h. After completion, the reaction was quenched with $H_2O$ (150 mL) and extracted with EtOAc (500 mL × 3). The combined organic phases were washed with brine (400 mL x 2), dried over $Na_2SO_4$, filtered, and concentrated. The residue was purified by preparative HPLC (Gemini column; ACN/H2O containing 0.05% $NH_3$) to afford a mixture of lindoldhamine stereoisomers (4.0 g, 42.1% yield) as an off-white solid.

**Step 10. Chiral separation of lindoldhamine stereoisomers**

The mixture of lindoldhamine stereoisomers (2.0 g) was separated by SFC using a CHIRALPAK IJ column (MeOH/ACN/DEA = 95:5:0.1, v/v/v), followed by a CHIRALPAK IC column (hexane/EtOH/DEA = 40:60:0.1, v/v/v), to afford (*R*,*S*)-lindoldhamine (203.09 mg, 10.2% yield) as an off-white solid. The other three stereoisomers, (*S*,*R*)-, (*R*,*R*)-, and (*S*,*S*)-lindoldhamine, were also obtained. Absolute configurations were assigned by combining chiral analysis with calculated and experimental ECD data (Supplementary Fig. 8), as described in the Methods section.

#### Compound structure analysis and assignment

**Cyclic bisBIAs:**

Because authentic standards for some methylated intermediates in the bisBIA pathway are commercially unavailable, we assigned the structures based on NMR, or MS/MS analysis and chromatographic retention time shifts on reversed-phase columns. The *N*2-methylation of (*R*,*S*)-**8** to (*R*,*S*)-**7** catalyzed by B2NMT was confirmed by comparison with the authentic (*R*,*S*)-**7** standard prepared from the aforementioned CYP80Q4 reaction, with (*R*,*S*)-**7** subsequently cyclized into two distinct cyclic bisBIA scaffolds. Similarly, NMR analysis confirmed that B2′NMT selectively methylates the N2′ position of both (*R*,*S*)-**16** and (*R*,*S*)-**17** to produce (*R*,*S*)-**35** and (*R*,*S*)-**36**. Conversely, the *O*-methylation sites targeted by B7OMT and B12OMT were deduced via diagnostic MS/MS fragmentation analysis.

A mass shift of +14 Da relative to the substrate indicates a single methylation event. Because *O*-methylation substantially increases hydrophobicity and prolongs chromatographic retention by ablating a strong hydrogen bond donor, whereas N-methylation exerts minimal impact on overall polarity due to the preservation of the amine's protonation state[7], we leveraged these retention time differences for initial categorization. Specifically, *O*-methylated products exhibited a distinct retention delay (~1.0 min in this work), whereas N-methylated derivatives showed minimal shifts (~0.1 min in this work) relative to their substrates. Specific methylation sites (*N*2, *N*2′, *O*7, *O*7′, and *O*12) were subsequently assigned by comparing their MS/MS spectra with those of authentic standards and NMR-validated intermediates. This approach is particularly effective for cyclic bisBIA scaffolds, which yield distinct diagnostic MS/MS fragments according to previous reports and our spectral analyses[8].

In positive-ion mode, bisBIAs typically produce both singly ($[M + H]^+$) and doubly ($[M + 2H]^{2+}$) protonated ions. The MS/MS fragmentation patterns are consistent across both C7-O-C8′ and C7′-O-C8 linked cyclic scaffolds, independent of the C1/C1′ absolute configurations[8]. Using isotetrandrine ((*R*,*S*)-**23**; $[M + H]^+ = m/z$ 623, $[M + 2H]^{2+} = m/z$ 312) as a representative example (Supplementary Fig. 14d), the MS/MS behavior of cyclic bisBIAs is characterized by the formation of abundant diagnostic A-type ions representing the dimeric isoquinoline moiety, typically corresponding to $[M + H - 242]^+$ or $[M + H - 228]^+$. These arise from preferential cleavage of the C1–Cα bond followed by *O*-demethylation of an aromatic methoxy group, yielding an A-type ion at *m*/*z* 381 for (*R*,*S*)-**23**. Comparing the A-type ion *m*/*z* of a product with that of isotetrandrine determines whether methylation has occurred within the dimeric isoquinoline core. The precise nominal mass loss (–242 vs. –228 Da) strictly depends on the methylation status of the *O*12 position. Further ether bond cleavage of the A-type ion generates the B-type mono-isoquinoline ion at m/z 192. Conversely, the remaining fragment from the initial C1–Cα cleavage yields the C-type diphenyl ether ion. Its mass (*m*/*z* 213 vs. 227) serves as a direct indicator of *O*12 methylation; for example, the *m*/*z* 227 peak in isotetrandrine confirms the presence of the *O*12 methyl group. Additionally, the neutral loss of methylamine from the *N*2′ position ($-NH_2CH_3$) yields D-type ions (*m*/*z* 592 for isotetrandrine), whereas the combined loss of both the *N*2′ methylamine and the *N*2 methyl group (–C2H5N) yields E-type ions (*m*/*z* 580). By comparing the D- and E-type ions of the products with those of isotetrandrine, we could assign *N*-methylation to either the *N*2 or *N*2′ position. The proposed MS/MS fragmentation pathway for the authentic standard isotetrandrine is illustrated below. Ultimately, the cascade reaction incorporating the complete suite of MTs produced isotetrandrine that perfectly matched the authentic standard, firmly validating the reliability of our enzymatic assignments and MS/MS-based deductions (Fig. 3, Supplementary Fig. 14a-d).

The following section details the structural assignment of the methylated products via comprehensive MS/MS analysis.

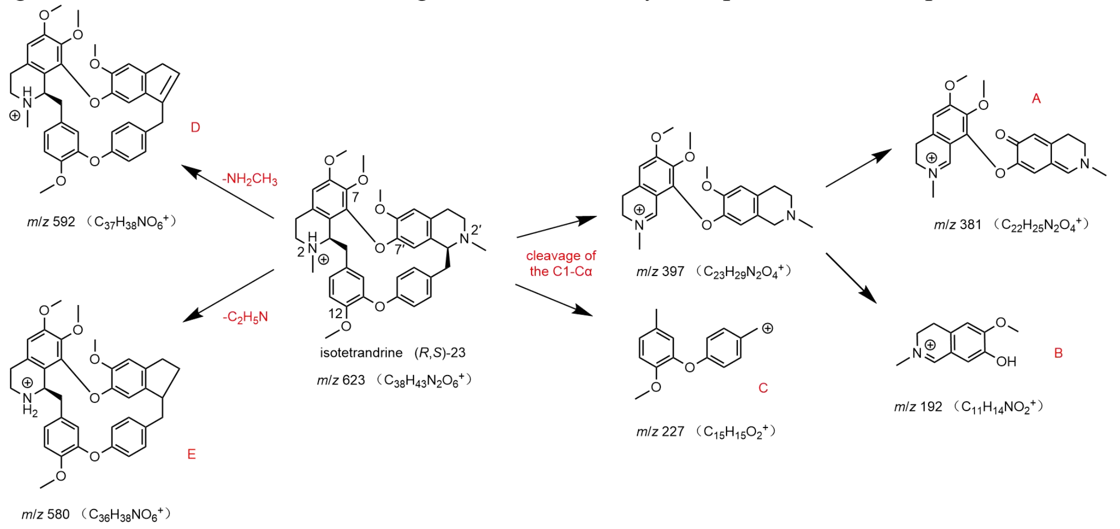

(*R*,*S*)-**30** and (*R*,*S*)-**31**:

In the cascade reaction, B7OMT-catalyzed methylation of the substrates (*R*,*S*)-**16** and (*R*,*S*)-**17** yielded the monomethylated products (*R*,*S*)-**30** and (*R*,*S*)-**31**, respectively, each exhibiting the expected +14 Da mass shift. In their MS/MS spectra (Supplementary Fig. 12c, d), the A-type ions (*m*/*z* 353) shifted by +14 Da relative to the substrates (Supplementary Fig. 10c, d), whereas the C-type ions remained unshifted at *m*/*z* 213. This localized the newly introduced methyl group to the dimeric isoquinoline core rather than the diphenyl ether moiety (*O*12). A pronounced chromatographic retention delay of ~1.0 min relative to the substrates strongly indicated *O*-methylation rather than *N*-methylation (Supplementary Fig. 16b, c, e, f). This assignment was further corroborated by the absence of characteristic *N*-methylation MS/MS fragments: the B-type (*m*/*z* 178) and E-type (*m*/*z* 564) ions remained consistent with the substrates, lacking the mass shifts associated with *N*2 (*m*/*z* 192) or *N*2′ methylation observed in isotetrandrine. Because the *O*7 and *O*7′ hydroxyls are the only available *O*-methylation sites on the dimeric isoquinoline core of these scaffolds, these data collectively indicate that B7OMT catalyzes methylation at either the *O*7 or *O*7′ position to generate (*R*,*S*)-**30** and (*R*,*S*)-**31**.

(*R*,*S*)-**33** and (*R*,*S*)-**34**:

In the cascade reaction, B12OMT-catalyzed methylation of the substrates (*R*,*S*)-**16** and (*R*,*S*)-**17** produced the monomethylated products (*R*,*S*)-**33** and (*R*,*S*)-**34**, respectively, each exhibiting the expected +14 Da mass shift relative to the substrates (Supplementary Fig. 10c, d). In their MS/MS spectra (Supplementary Fig. 12e, f), the A-type ions (*m*/*z* 339) remained consistent with the unmethylated dimeric isoquinoline core, shifting by –42 Da relative to isotetrandrine (corresponding to the absence of three methyl groups). Conversely, the C-type diphenyl ether fragments (*m*/*z* 227) exhibited a +14 Da shift, perfectly matching the *O*12-methylated pattern observed in isotetrandrine. This firmly localized the newly introduced methyl group to the *O*12 position on the diphenyl ether moiety. Coupled with a pronounced chromatographic retention delay of >1.0 min characteristic of *O*-methylation (Supplementary Fig. 15e, h), these data identify the products as the *O*12-methylated derivatives (*R*,*S*)-**33** and (*R*,*S*)-**34**.

(*R*,*S*)-**22**:

In the cascade reaction incorporating the complete MT suite, incubation with (*R*,*S*)-**16** yielded the product (*R*,*S*)-**22**, which exhibited a +56 Da mass shift relative to the substrate, indicative of four methylation events. Detailed MS/MS analysis revealed that the diagnostic A-, B-, C-, and E-type ions of (*R*,*S*)-22 perfectly matched the corresponding fragments of the isotetrandrine standard (Supplementary Fig. 14c, d). This comprehensive spectral alignment confirms that the methylation pattern is identical to that of isotetrandrine, identifying (*R*,*S*)-**22** as the fully methylated C7-O-C8′ terminal product, featuring methylation at the *N*2, *N*2′, *O*7′, and *O*12 positions.

Incubation of the BBOX-generated imine intermediate **24** with B2′NMT2, B7OMT, and B12OMT—either collectively or individually—yielded the corresponding trimethylated and distinct monomethylated products. Given the regioselectivity established for these MTs, we hypothesized that their targeted methylation sites on **24** would mirror those observed for the fully reduced scaffolds (*R*,*S*)-**16** and (*R*,*S*)-**17**. To definitively verify these assignments on the oxidized scaffold, we subjected the products to comprehensive MS/MS analysis.

**25**:

Following the BBOX-catalyzed oxidation of (*R*,*S*)-**17** to the intermediate **24** (m/z 565), the subsequent addition of B2′NMT2, B7OMT, and B12OMT yielded the trimethylated oxidized product **25** (*m*/*z* 607) (Supplementary Fig. 21a, b). In its MS/MS spectrum (Supplementary Fig. 21e), the A-type ion was observed at m/z 365 (corresponding to $[M + H − 242]^+$). This represents a –16 Da shift relative to the A-type ion of isotetrandrine, precisely matching a dimeric isoquinoline core lacking one methyl group (–14 Da) and two protons (–2 Da). Consequently, this fragmentation signature indicates the addition of exactly two methyl groups to the dimeric isoquinoline moiety, dictating that the third methylation occurs at the *O*12 position on the diphenyl ether ring. Coupled with the previously established regioselectivity of B2′NMT, catalytic methylation at the *N*2 position is highly improbable[9,10]. Collectively, these distinct fragmentation patterns unambiguously map the three methylations to the *N*2′, *O*7, and *O*12 positions, definitively validating the structure of **25**.

**49**:

Following the BBOX-catalyzed oxidation of (*R*,*S*)-**17** to intermediate **24** (*m*/*z* 565), subsequent incubation with B7OMT yielded product **49** (*m*/*z* 579). This +14 Da mass shift relative to **24** is indicative of a single methylation event (Supplementary Fig. 21a, b). A pronounced chromatographic retention delay of ~1.0 min suggested *O*-methylation rather than *N*-methylation (Supplementary Fig. 21b, d). In the MS/MS spectrum, the diagnostic A-type ion was observed at *m*/*z* 351, corresponding to the [M + H − 228]$^+$ fragment (Supplementary Fig. 21e). This specific neutral loss confirms that the *O*12 position remained unmethylated. Collectively, these data support the assignment of this compound as the *O*7-methylated oxidized derivative **49**.

**50:**
Following the BBOX-catalyzed oxidation of (*R*,*S*)-**17** to intermediate **24** (*m*/*z* 565), subsequent incubation with B2′NMT2 yielded product **50** (*m*/*z* 579). This +14 Da mass shift relative to **24** indicates a single methylation event (Supplementary Fig. 21a, b). A minimal chromatographic retention shift (<0.1 min) relative to the substrate strongly suggested *N*-methylation rather than *O*-methylation (Supplementary Fig. 21b, d). In the MS/MS spectrum, the diagnostic A-type ion was observed at *m*/*z* 351, corresponding to the [M + H − 228]$^+$ fragment. This specific loss confirms that the *O*12 position remained unmethylated, effectively narrowing the potential target sites to either the *N*2 or *N*2′ position (Supplementary Fig. 21e). Considering the intrinsic difficulty of imine *N*-methylation[9,10], alongside the established regioselectivity of B2′NMT, these data collectively identify the product as the *N*2′-methylated oxidized derivative **50**.

**51:**
Following the BBOX-catalyzed oxidation of (*R*,*S*)-**17** to intermediate **24** (m/z 565), subsequent incubation with B12OMT yielded product **51** (*m*/*z* 579). This +14 Da mass shift relative to **24** indicates a single methylation event (Supplementary Fig. 21a, b). A pronounced chromatographic retention delay of ~1.0 min suggested *O*-methylation rather than *N*-methylation (Supplementary Fig. 21b, d). In the MS/MS spectrum, the diagnostic A-type ion was observed at *m*/*z* 337, corresponding to the [M+H−242]$^+$ fragment (Supplementary Fig. 21e). This specific neutral loss localizes the methylation to the *O*12 position, confirming the identity of the product as the *O*12-methylated oxidized derivative **51**.

#### Linear bisBIAs
In addition to the cyclic bisBIAs, the structures of several linear bisBIA derivatives identified in this study required assignment. These include the CYP80Q4 and B2′NMT1/2 product (*R*,*S*)-**9**, as well as the linear MT products (*R*,*S*)-**27**, (*R*,*S*)-**28**, (*R*,*S*)-**29**, and (*R*,*S*)-**32**. The structural elucidation of these compounds was achieved through a combined analysis of chromatographic retention times, MS/MS fragmentation patterns, and the comparative substrate specificities of the characterized enzymes.

The MS/MS fragmentation behaviors of linear bisBIAs follow established pathways, as previously reported[11]. As illustrated below, we outline these characteristic cleavage patterns using daurisoline as a representative example. Linear bisBIAs typically undergo two primary modes of fragmentation: C–N bond cleavage within the isoquinoline rings and C1–Cα bond cleavage at the benzylic positions. In positive ion mode, protonated daurisoline ([M+H]$^+$, *m*/*z* 611) undergoes characteristic C–N bond scissions driven by the instability of the tertiary amines. Specifically, the loss of an N-linked methylamine moiety (–$NH_2CH_3$, 31 Da) yields the diagnostic fragment F (*m*/*z* 580), while the elimination of –$CH_3N{=}CH_2$ (43 Da) generates fragment J (*m*/*z* 568). Concurrently, characteristic C1–Cα benzylic cleavages produce diagnostic isoquinoline-derived fragments G and H (*m*/*z* 192 and 206, respectively) and the diphenyl ether-derived fragment I (*m*/*z* 213).

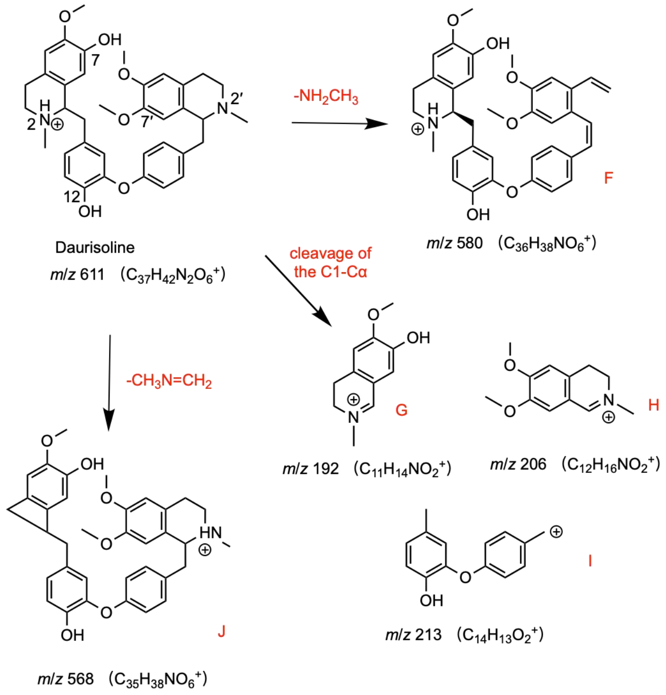


(*R*,*S*)-**9**:
The linear bisBIA (*R*,*S*)-**9** was initially generated via the CYP80Q4-catalyzed C–O phenol coupling of (*R*)-**4** and (*S*)-**5**. Subsequent incubation of (*R*,*S*)-**8** with either B2′NMT1 or B2′NMT2 yielded a product exhibiting a chromatographic retention time identical to that of (*R*,*S*)-9 (Supplementary Fig. 15h, i). A minimal retention time shift (~0.1 min) relative to the substrate (*R*,*S*)-**8** strongly supported its assignment as an *N*-methylated derivative. Taken together with the strict regioselectivity previously demonstrated by B2′NMT1 and B2′NMT2, we deduced that (*R*,*S*)-**9** represents the specific N2′-methylated product of the (*R*,*S*)-**8** scaffold.

(*R*,*S*)-**27**, (*R*,*S*)-**28**, and (*R*,*S*)-**29**:
Incubation of the linear substrate (*R*,*S*)-**8** with B7OMT yielded three distinct products: two exhibiting +14 Da mass shifts (monomethylated) and one displaying a +28 Da shift (dimethylated) (Supplementary Fig. 16g–j). Pronounced retention time delays of >0.5 min and >1.0 min relative to the substrate, coupled with the established preference of B7OMT for the *O*7 and *O*7′ positions in cyclic bisBIAs, strongly suggested *O*-methylation. In the MS/MS spectra of the monomethylated products (*R*,*S*)-**27** and (*R*,*S*)-**28** (Supplementary Fig. 16k, l), the simultaneous presence of both the H-type (*m*/*z* 192) and G-type (*m*/*z* 178) fragments indicated that a single methyl group was appended to either the H- or G-type moiety. Given that *O*7 or *O*7′ are the sole available sites on these fragments, (*R*,*S*)-**27** and (*R*,*S*)-**28** are identified as the distinct *O*7- and *O*7′-monomethylated isomers; however, their exact structural assignment to the two closely eluting chromatographic peaks remains unresolved. Conversely, the dimethylated product (*R*,*S*)-**29** lacked the *m*/*z* 178 fragment entirely, displaying only the *m*/*z* 192 ion (Supplementary Fig. 16m). This –14 Da shift for the H-type fragment, coupled with the unchanged G-type fragment relative to daurisoline, confirms the concurrent methylation of both the *O*7 and *O*7′ hydroxyl groups, definitively identifying (*R*,*S*)-**29**.

(*R*,*S*)-**32**:
Incubation of the linear substrate (*R*,*S*)-**8** with B12OMT yielded product (*R*,*S*)-**32**, which exhibited a +14 Da mass shift. A pronounced retention time delay of >0.5 min relative to the substrate indicated *O*-methylation (Supplementary Fig. 17g–i). Crucially, the MS/MS spectrum of (*R*,*S*)-**32** revealed an I-type fragment at *m*/*z* 227 (Supplementary Fig. 17j), representing a +14 Da shift relative to the corresponding I-type fragment of daurisoline. This diagnostic shift localizes the newly added methyl group to the *O*12 position. This assignment aligns perfectly with the strict regioselectivity exhibited by B12OMT toward cyclic bisBIAs, definitively confirming the structure of (*R*,*S*)-**32** as the *O*12-methylated derivative of (*R*,*S*)-**8**.

**FASTA nucleotide sequences for genes described in this manuscript. "y" stands for codon optimization according to preferences for *Saccharomyces cerevisiae*.**

>CYP80Q4
ATGGATCCAATCACACTGTTACTGATTCTCTTCACACTTCTCTCATCGCTCTTTTTTTCTTTCG
AGTTCAAACGTTACATATCCCCTAAAAAACTTCCTCCAGGCCCCTTCGCATGGCCTCTGATC
GGCAACACCGTTCTGAGCGTCAGCGAGGGCCAGCTCCACGTCGCGCTGGCCGACCTGGCCA
AGAAGTTCGGTCCCCTCATGATGCTCAAGTTCGGGCTCGTCCCGCCGCTGATGGTGGCCTCG
AACCACGTGGCGGCCGCGGAGATACTCAAGACGCAGGACCGCGCCTGTTCCGGCCGCAAC
GTGGCACACAGCGTGCAAGTGCCTGGTTACATTCAGTACTCCATGGTGTGGGCTGACTGCA
CCGAGCACTGGAAAATGGTGAGGAGGATATGGAGGACAGAGTTGTTCTCCACCAAGATGG
TGGACTTGCAGACCAGTATAAGGGAGGAGAAGGTAAGGGAATTGCTAGGGTTTTGAGGA
GGAAAGAAGGGCAAGTGGTGCAGCTCAGTGATGCCATATTTGGGTGCATTATCAATGTTTT
AGGGTCTATAATTTTTAACCAAAATGTGTATGATTATGAGGGGAAGGTTGATAATGAGAAG
GGCATGAAGGGTATGATTAGGCAGCTTATGGTGTTGGCTGCAAAACCTAATTTGCCTGATTT
TTATCCGGTTTTTGGTCCCCTGGACTTGCAAGGGATTAGAAGAACCACCGCTGAGTGTGTGA
AGCAGATGAATGAGTACTGGGGAGGGATAGTGAAGGAGAGGAGGGCAAGCAAAGATCAC
TCCAGGAATGACTTCTTGGATGTGTTGATCCAAGCCAACTTCACCGATGCGCAAATCGATG
CCTTGCTCTTGGAAATATTCGGACCTGGTTCGGACACTAGCTCATGTACCATAGAGTGGGCG
ATGTCCGAGCTTATTAAGAACCCTAACAAGCTATTGAAACTCCAAGAGGAGCTGAACAGAG
TGATTGGGCAGAGCAGGGAAGTGAGGGAATCAGATTTAGAAAATCTCCCATATCTCCATGC
GTGCATAAAGGAGACTTTGAGATTGCATCCACCAGTGACATTCCTTCTCCCTCACCGCGCAA
CTGAGACGTGTGAGGTAATGAACTGCACCATCCCTAAGGACTCACAAATTTTCGTGAACGC
TTATTCGATACAGAGGGATCCAGAGGTGTGGGAGGACCCATTGATCTTCAAGCCGGAGCGG
TTTTTGAACTCGACGGTGGACTATCAGGGCAACGACTTTCACTACATACCGTTCGGTGCTGG
TCGGAGAATATGCCCTGCACTCAACATGGCTTCTAGGGCCACCAGGTTTATGTTAGGCTCTT
TGATTCACAATTTCGAGTGGAGTTTGCCTAATGGAATGAAACCAAGCGAGTTGGACATGCG
AGACGCGTTCGTGTTGGTGCTTGCCAAGACAGTTCCTCTCAGTGTTATACCAAAGGCAAGA
TCATCAGCAACTGCGGTTTAA

>yCYP80Q4
ATGGACCCTATTACATTATTGTTGATCTTGTTTACCCTGTTATCTTCTTTATTCTTCTCTTTTG
AGTTCAAGAGATACATTTCTCCAAAAAAGTTGCCACCAGGTCCATTTGCTTGGCCATTGATT
GGTAATACAGTTTTGTCTGTTTCTGAAGGTCAATTACATGTTGCTTTAGCTGATTTGGCTAA
AAAATTTGGTCCATTGATGATGTTAAAGTTCGGTTTAGTTCCACCATTAATGGTTGCTTCTA
ATCATGTTGCTGCTGCTGAAATTTTAAAAACTCAAGATAGAGCTTGCTCTGGTAGAAATGTT
GCTCATTCTGTTCAAGTTCCAGGTTATATTCAATATTCTATGGTTTGGGCTGATTGTACAGA
ACATTGGAAAATGGTTAGAAGAATTTGGAGAACAGAATTATTCTCTACTAAGATGGTTGAT
TTGCAAACTTCTATTAGAGAAGAAAAGGTTAGAGAATTGTTGGGTTTTCTAAGAAGAAAAG
AAGGTCAAGTTGTTCAATTGTCTGATGCTATTTTTGGTTGTATTATCAACGTTTTGGGTTCTA
TTATCTTCAATCAAAACGTTTACGACTACGAAGGTAAAGTTGATAATGAAAAAGGTATGAA
GGGTATGATTAGACAATTGATGGTTTTAGCTGCTAAACCAAATTTGCCAGATTTTTATCCAG
TTTTTGGTCCATTAGATTTGCAAGGTATTAGAAGAACAACAGCTGAATGTGTTAAACAAAT
GAATGAATACTGGGGTGGTATTGTTAAAGAAAGAAGAGCTTCTAAAGACCATTCTAGAAAT
GATTTTCTGGATGTTTTGATCCAAGCTAATTTTACAGATGCTCAAATTGATGCTTTGTTGTTG
GAAATTTTCGGTCCAGGTTCTGATACATCTTCTTGTACAATTGAATGGGCTATGTCTGAATT
AATTAAGAATCCAAACAAGCTGTTGAAGTTGCAAGAAGAATTGAATAGAGTTATCGGTCAA
TCTAGAGAAGTTAGAGAATCTGATTTGGAAAATTTGCCATATTTGCATGCTTGTATTAAGGA
AACTTTGAGATTACATCCACCAGTTACTTTTCTATTGCCACATAGAGCTACTGAAACTTGTG
AAGTTATGAATTGTACTATCCCAAAAGATTCTCAAATCTTTGTTAACGCTTACTCTATTCAA
AGAGATCCAGAAGTTTGGGAAGATCCATTGATTTTTAAACCAGAAAGATTCTTGAACTCCA
CTGTTGATTATCAAGGTAATGATTTTCACTACATCCCATTTGGTGCTGGTAGAAGAATTTGT
CCAGCTTTAAATATGGCTTCTAGAGCTACAAGATTCATGTTAGGTTCTTTGATTCATAACTT

CGAATGGTCTTTGCCAAATGGTATGAAACCATCTGAATTGGATATGAGAGATGCTTTTGTTT
TGGTTTTGGCTAAAACAGTTCCATTGTCTGTTATTCCAAAAGCTAGATCTTCTGCTACTGCTG
TTTAA

>yBs-MaCYP80A1
ATGGACTATATTGTTGGTTTCGTAAGTATTTCTTTAGTTGCTCTGCTTTACTTCCTTTTGTTCAAA
CCTAAGCATACTAATCTACCTCCTTCTCCCCCAGCATGGCCGATAGTAGGTCATCTGCCAGACT
TGATCTCTAAAAACAGCCCACCATTCCTAGATTATATGTCTAACATCGCTCAGAAATATGGTCC
GTTGATTCATTTGAAGTTCGGTCTGCACTCTAGTATTTTTGCCTCTACTAAGGAAGCAGCTATGG
AGGTCTTACAGATTAACGACAAGGCTTTGAGCGGGAGACAGCCTCTTCCCTGCTTTAGGATAA
AGCCCCATATAGACTATTCTATCGTCTGGTCCGACTCTAATAGTTACTGGAAGAACGGACGTAA
GATACTTCACACCGAAATATTTTCCCAGAAAATGCTTCAAGCGCAGGAAAAAAATAGGGAGAG
GGTGGCAGAAAACCTGGTGAACTTTATCCGTACAAAAGTGGGTAACGTTGTCGAACTAAGGAG
CTGGCTATTTGGCTGTGCGCTGAATGTATTAGGCCATGTTGTATTTTCAAAGGATGTGTTCGGCT
ACTCAGATCAATCAGACGAAGTTGGTATGGATAAGCTGATTCATGGTATGATCATGACGGGCG
GAGACTTCGATGTAGCATCCTACTTCCCGTTCTTGGCAAGATTTGATATACACGGTCTTAAAAG
GAAAATGGATGAACAGTTTAAATTACTAATCAAGGTTTGGGAGGGAGAAGTATTAGCGAGGAA
AGCTAACCAGAACCCCGAGCCGAAGGATATGTTAGATGTGCTAATTGCGAATGACTTTAATGA
ACATCAGATAAATGCTATGTTCTTAGAAACGTTCGGTCCAGGGAGCGATACGAGCTCAGCAAC
AATAGAATGGGCGTTGGCACAATTGATAAAAAACCCAGACAAGTTGGCAAAACTAAGGGAAG
AGCTGGACAGGGTTGTAGGAAGATCATCCACGGTGAAGGAGAGCCACTTCTCACAGTTGCCAT
ATCTTCAGGCTTGCGTCAAAGAGACGATGAGACTGTATCCGTCCGTTACGATCATGATCCCACA
CCGTTGTATGGAAACGTGCCAAGTCATGGGCTATACTATTCCTAAAGGTATTGATGTTCATGTC
AATGCCCATGCGATTGGCAGGGACCCTAAGGATTGGAAAGACCCACTTAAGTTTCAGCCTGAG
AGGTTCTTAGACTCCGATATAGAGTACAATGGGAAGCAATTCCAGTTTATTCCGTTTGGTTCCG
GCAGAAGAATTTGCCCTGGATGGCCCCTGGCCGTTCGTACCATCCCGCTGGTCTTAGCGTCACT
GGTCCATGCATTCGATTGGGAATTACCTGATGGAGTTCCCAACGAAAAGCTTGACATGGAAGA
ATTGTTTACCCTAACTCTTTGCATGGCCAAGCCATTACGTGTAATCCCCAAGGTTCGTATATAA

>yBsCYP80A1
ATGGATTACATCGTTGGTTTTGTTTCTATTTCTTTGGTTGCTTTGTTGTATTTCTTGTTGTTTA
AGCCAAAGCATACTAATTTGCCACCATCTCCACCAGCTTGGCCAATTGTTGGTCATTTGCCA
GATTTGATTTCTAAAAATTCCCCACCATTTTTGGATTATATGTCTAATATCGCTCAGAAATA
CGGTCCATTAATTCATTTGAAATTCGGTTTGCATTCCTCTATTTTTGCTTCTACAAAAGAAGC
TGCTATGGAAGTTTTGCAAACTAATGATAAAGTCTTGTCTGGTAGACAACCATTACCATGTT
TTAGAATTAAGCCACATATCGATTACTCTATTTTGTGGTCTGATTCTAATTCTTACTGGAAA
AAAGGTAGAAAGATCTTGCATACAGAAATTTTTAGCCAAAAGATGTTGCAAGCTCAAGAAA
AGAATAGAGAAAGAGTTGCTGGTAATTTGGTTAATTTTATCATGACTAAGGTCGGTGACGT
TGTTGAATTAAGATCTTGGTTGTTTGGTTGTGCTTTAAATGTTTTGGGTCATGTTGTTTTTAG
CAAAGATGTTTTTGAGTACTCTGATCAATCTGATGAAGTTGGTATGGATAAATTAATCCATG
GTATGTTGATGACTGGTGGTGACTTTGATGTTGCTTCTTATTTTCCAGTTTTGGCTAGATTTG
ATTTGCATGGTTTAAAAAGAAAGATGGATGAACAATTCAAGTTGTTAATCAAGATCTGGGA
AGGTGAAGTTTTAGCTAGAAGAGCTAATAGAAATCCAGAACCAAAAGATATGTTGGATGTT
TTGATTGCTAACGATTTTAACGAACATCAAATCAATGCTATGTTCATGGAAACTTTTGGTCC
AGGTTCTGATACTAATTCTAATATTATCGAGTGGGCTTTAGCTCAATTAATTAAGAATCCAG
ATAAGTTGGCTAAGTTAAGAGAAGAATTAGATAGAGTTGTCGGTAGATCTTCTACAGTTAA
AGAATCTCATTTCTCTGAATTGCCATATTTGCAAGCTTGTGTTAAAGAAACTATGAGATTGT
ATCCACCAATTTCTATTATGATCCCACATAGATGTATGGAAACATGTCAAGTTATGGGTTAT
ACTATTCCAAAAGGTATGGATGTTCATGTTAATGCTCATGCTATTGGTAGAGATCCAAAAG
ATTGGAAAGATCCATTGAAATTTCAGCCAGAAAGATTTTTGGATTCTGATATTGAATACAAC
GGTAAACAATTCCAATTCATTCCATTTGGTTCTGGTAGAAGAATTTGTCCAGGTAGACCATT
GGCTGTTAGAATTATTCCATTAGTTTTGGCTTCTTTGGTTCATGCTTTTGGTTGGGAATTACC

AGATGGTGTTCCAAATGAAAAATTGGATATGGAAGAATTGTTCACATTGTCTTTGTGTATGG
CTAAACCATTGAGAGTTATTCCAAAAGTTAGAATTTAA

>yNnCYP80Q2
ATGGCTTTACTTGCTTTGTTCATCTTATTTTTGTTGTCAATCCTATCCTTGGTCCTATTTTTAA
AGCCATCTTCCAAGAAGTTACCACCTGGCCCATTCTCTTGGCCAATCATTGGTACCCAATTG
CCATCTCCAACCATGAAGCCAAACTTGGAATTATTCAAGTTGGCTCAAAGATACGGTCCAT
TGATGTTGTTCAGATTTGGTTTCGAAAACGTTGTGGTTGCTTCCAACCACGTCGCTGCTATG
GAAGTTTTGAAGAACCAAGACAGAGTCTTGTCCGGTAGATTCAAGGCTAACTCCGTCAGAG
TCAAGGGTTACATTGAATACTCCATGGTCTGGGCTGACTGTACCGATTACTGGAAGATGGTT
AGAAAAATCTTGAGAACTGAATTGTTCTCTACCAAGATGTTGGATGTTCATGCTCACGTTAG
AGAAGAAAAGGTCTCTGAATTGATGAAGTTCTTGAGAAGAAAAGAAGGTGAAGAAGTTAA
CTTCGTTGACGTTATCTTCGGTTGTATCCTAAACATGTTGGGTGCCTTGATCTACTCTAAGGA
TGTCTACGATTTCGAAGACAAGACTGACATTAACTTAGGTATGAAAGGTATGATCAGACAA
TTGATGATTTTGGCCGCTACTCCAAACATTGCTGACTTGTACCCAATTTTCTTCGATGGTTCC
GACTTCCAAGGTTTGAGAAAGGAATCTGCCGCCTGCGTTAAGAGAATGTCTGAATCCTGGG
CCGCCATTATCAACGAAAGAAGAAAGAACAACGATCATACTAAGAACGACTTGTTGCAAG
TTTTGTTGGATTCTGGTTTCTCTGATCCACAAATCGATGCTATCTTTTTGGAAACCTTCGGTC
CAGGTTCTGACACTTCTGCTTCTACTATCGAATGGGCTTTGGCTGAACTGTTGCGTAACCCA
GAAAAATTAGTCAAGTTGCACGAAGAGCTCGACAGAGTTATCGGTAGAAACAACACTGTTA
AGGACTCCGATTTGCCTAATTTGCCATACTTGCACGCTTGTGTCAAGGAAACATTGAGATTG
CACCCACCAGTTCCATTCTTGATTCCACACATTGCCTTGGAATCTTGTGAAGTTATGAACTA
CACCATTCCAAAGGGTAGCGAAGTTTTGGTCAATCTATATGCAATTGGTCGTGACCCAACC
ACCTGGGACAACCCAAATTCTTTCTTGCCAGAAAGATTCTTGAACAGTGAAGTCGACTACC
AAGGTAACCATTTCCAATACATCCCATTCGGAGCTGGCAGAAGAATGTGTCCAGGTATGAG
TTTAGGTACTCGTGTCGTCAGATTGATTCTAGCTGCTTTAGTTCACACTTTCGACTGGTCTTT
GCCAGGTGGTATGCACCAAGATGAATTGGACATGGCTGATAGATTTGGTGTTGGTTTCCAA
AAGGAAACTCCCTTGGTTGTCATCCCAACCCTCCGTAAATGA

>CYP82BC1
ATGATCCCGCAACCTTACTTTCAACCTGCAATCCTATGGGTCGTGCTCTTAGCTTCCCTCCCC
ATCTTCTACTATCTCCTGATCAGACGGCCCAGAACCACCAACCCCTCCATCAACGGCGTAA
AAGACCTCCCAGAACCAGATGGCGCGTGGCCGATCATCGGCCACCTCCTGGCCTTCAGCCC
CAATGACCTCTGGTACCGAAAGCTCAGCGACATGGCGGACAAGGTCGGCCCCATCTTCGCG
ATCCGCCTTGGCACGCGCCGAGCCGTGGTCATCAACAACTGGGAAATGGCAAGAGAGTGCT
TCACCACCAACGACAAAACGCTCGCGAGCCGCCCGTCTTCGTTGCACTCCAAGTTAATGGG
CTACAACGGCGCCACGTTCGGGTTCGCCCCAGAAGGGCCGTACCTGCGCGAGGTCCGGAAA
GTGATTACCCAAGAGCTCTTCTCCAACAAGCGTCTCCAGCTGTTGAAGCACGTGAGAGTGT
CGGAGATAGACCTGATGATTAAGGATCTCAAGGGGCTTTATTTCAAGAACAAAGACTCTGA
TGGCTCGTCCGTGTTGGTAAAGTTGGACACGTGGATGTGGGCTCTGACGCTGAACGTTACG
CTCAGAAGTATTGCTGGGAAGAGGTATTATGATGGTGGGGATGAAAGTCATGACGAACGCT
ATGGCGATGTTGAAGCTAAGAGGTGGAGGGAGACTTTTGATAATTTGATTCACACTCTGAG
GGCGTTTACGGTGTCGGACATATTTCCGGGGTGGGAGTTTGTGGATTTGGTGATGGGAGTG
CAGCGGCGGATGAGGAAGGCGAGTGAGGAGATGGACTTGGTGTTGAGTGGGTGGTTGGAG
GAGCATAGGAGGAGGAAGGGAAGGAGCGAGGGGGAGGAGGACTTCGTGGACGTGATGAT
GAACATTGACACCAACGCTAAGTTGTTCTCAGAGCATGACGTGGACACTGTTGTTAAAGCT
ACTTGTTTGGATATATTAATAGCAGCTAGAGACACAACATCCATCACCCTCCTATGGGTCAT
CCCCATGTTACTGAACAATCCTCAAGTTCTGAAGAAGGTGAAAGAAGAGTTGGACAATCAA
ATAGGCAAAGATAGGCACGCAGAAGAGGAAGACATCCAACACTTGCCTTACCTACAAGCA
GTTGTAAAGGAGACTTTGCGCTTATACCCTCCTTCCCCGCTTTCATTACCACATGAGGCATC
AAATGACTGCGTCGTTGGCGGGTTCCATGTCCCAAAAGGGTCTTCCTTAATCACAAATATTT
GGAAGATACAACGCGATCCCAATATTTGGCCAGACCCTTTGGAGTTCAAGCCGGAGAGATT

CCTTGCTACCAATGACGCCCATAATGTTGATTTTAGAGGTTTAAACTTTGAATTAATTCCTTT
CGGGTCGGGTAGAAGAATGTGTGTGGGAACAAACCTTGCCGCTCATATCGTGCACCTAACG
CTTGCTCGTTTGTTCCATGAGTTTGAGATCAGAACTCCTGACAATACACCAATTGATATGAC
CGAAGCTCCGGACACTGATCTCGTTAGAGCAAGCCCTCTCAGTGTACTTATGCGCCCACGA
GTGATTCCTTCTCCTTGA

>yCYP82BC1
ATGATTCCACAACCATATTTCAACCAGCTATTTTGTGGGTTGTTTGTTAGCATCTTTGCCA
ATTTTCTATTACTTGTTGATCAGAAGACCAAGAACTACAAACCCATCAATTAATGGTGTTAA
GGATTTGCCAGAACCAGATGGTGCATGGCCAATTATTGGTCATTTGTTGGCTTTTTCTCCAA
ACGATTTGTGGTACAGAAAGTTGTCAGATATGGCTGATAAGGTTGGTCCAATCTTCGCAATC
AGATTGGGTACTAGAAGAGCTGTTGTTATTAATAACTGGGAAATGGCAAGAGAATGTTTCA
CTACAAACGATAAGACATTAGCTTCTAGACCATCTTCATTGCATTCAAAGTTGATGGGTTAC
AACGGTGCTACTTTTGGTTTTGCACCAGAAGGTCCATACTTGAGAGAAGTTAGAAAAGTTA
TTACACAAGAATTGTTTTCTAATAAGAGATTGCAATTGTTGAAGCACGTTAGAGTTTCAGAA
ATTGATTTGATGATCAAGGATTTGAAGGGTTTGTACTTCAAGAATAAGGATTCTGATGGTTC
TTCAGTTTTGGTTAAGTTGGATACTTGGATGTGGGCATTGACTTTGAACGTTACATTGAGAT
CTATCGCTGGTAAAAGATACTACGATGGTGGTGACGAATCACATGATGAAAGATACGGTGA
CGTTGAAGCAAAGAGATGGAGAGAAACATTCGATAATTTGATCCATACTTTGAGAGCTTTT
ACAGTTTCAGATATCTTTCCAGGTTGGGAATTTGTTGATTTGGTTATGGGTGTTCAAAGAAG
AATGAGAAAAGCTTCTGAAGAAATGGATTTGGTTTTATCAGGTTGGTTAGAAGAACACAGA
AGAAGAAAAGGTAGATCTGAAGGTGAAGAAGATTTTGTTGATGTTATGATGAACATCGATA
CTAACGCAAAGTTATTTTCAGAACATGATGTTGATACTGTTGTTAAGGCTACATGTTTGGAT
ATCTTGATCGCTGCAAGAGATACTACATCTATCACATTGTTGTGGGTTATTCCAATGTTGTT
GAACAACCCACAAGTTTTGAAGAAAGTTAAGGAAGAATTGGATAACCAAATCGGTAAAGA
TAGACATGCAGAAGAAGAAGATATCCAACATTTGCCATATTTGCAAGCTGTTGTTAAGGAA
ACTTTGAGATTGTACCCACCATCTCCATTGTCATTACCACATGAAGCTTCTAATGATTGTGT
TGTTGGTGGTTTTCATGTTCCAAAAGGTTCTTCATTGATCACTAACATCTGGAAGATTCAAA
GAGATCCAAATATTTGGCCAGATCCATTGGAATTCAAACCAGAAAGATTTTTAGCTACAAA
CGATGCACATAACGTTGATTTCAGAGGTTTGAACTTCGAATTGATCCCATTCGGTTCTGGTA
GAAGAATGTGTGTTGGTACTAATTTGGCTGCACATATTGTTCATTTGACATTGGCTAGATTG
TTCCATGAATTCGAAATTAGAACTCCAGATAATACACCAATTGATATGACTGAAGCACCAG
ATACAGATTTGGTTAGAGCTTCTCCATTGTCAGTTTTAATGAGACCAAGAGTTATTCCATCA
CCA

>CYP82BC2
ATGATGATCGAGATCCAAAATTTCTTTCCATCTCATATTGTATGCGGCGTGCTCTTAGCCGT
GATCTCAGCTTACTATCTCCTCACCAAGTATAAGAACAAGCCGGCCAGGGCGGCCGCACCA
GAACCGGCCGGTGCCTGGCCGGTCATCGGCCATATCCTTCAGTTCGGACCCCTCGACCTCTG
GCACCGAAAGCTCGGCGACATGGCCGACGAGATCGGACCGGCCTTCGTGATCCGCCTCGGC
ATGCTTCGCGTCCTCATCATAAACAACTGGGAGCTAGCCAAGGAGTGCTTCACATTGAACG
ACAAAATATTTGCGAGCCGTCCAGGCTCAATAACTTCCAAGTACATGGGCTACGACGGGGC
CATGTTGGGGTTCGCCCCGGACGGGCCCTTCATGCTCGAATTGCGAAAGGTAGCAATGCAA
GAGCTTTTCTCTAACCATAGGCTTCGTTTGCTGAGGCACATAAGGACGTCGGAAATCGAAA
TCATGATCAAGGGCCTAAATGAACTTTATCATAGAAATTTTAATAATGCTACGGGTGGTGCT
TCAATGCTAGTGCACTTGGATCCGTGGTTGGATGCACTGACGATAAACGTGATGCTTCGGA
GCATCGCGGGCAAGCGATACTACGATGGAGGAGCCGCGGTGGAAGGCGAGGAGGCGAAG
AGGTGGAAGGAAGCGTTTGGGAAAGTGATGCTCGTCTTCAAGATGCTTTTGGTGTCGGAGA
TGTTTCCTCGGCTGGAGATGGTTGATGTGATGCTGGGCACAACCCGTGCCATGAAGAAGGC
CTACGAGGAGATGGATTTGTTGTTGAGTGGATGGTTGGAGGAGCATCGGAGTAAAAAGGAC
CGTGGGGATCACGAACAAGACTTTGTAGATGTGCTGCTCAACGTTGGCAACAACAATCCTA
AGTTATTCTCTCAATACGACGTAGACTCCGTTGTTAAAGCCACTTGTTTGGACATATTGATA

GCCGCGAGCGACTCGACGACGCTGACCCTATCGTGGGCTCTATCCTTGTTACTCAACAATCG
TCGCGTCCTAAAGAAGGTCCAAGAAGAGCTGGACAATCAGATCGGGAAAGACAGGCACGC
GGAGGAAGAGGACATCAAAAACCTGCCCTACCTACAAGCCGTGGTGAAGGAGACTTGGCG
CTTGTACCCTCCTTCGCCGCTTTCACTGCCACACGAGGCCTCAAAAGATTGCGTTGTTGGCG
GATTCCATATCCCAAAGGGCACGAGCTTAATGACAAATGTTTGGAAGATACATCGCGATCC
AAAGGTTTGGCAAGAAGACCCGTTAGAGTTCAAGCCCGAGAGGTTCCTTAGTGCTTCTCAT
GCCCATGTTGATTTTCGAGGTCTACACTTCGAATTCATCCCTTTTGGATCAGGTAGAAGAAT
GTGTCTTGGAACAAACCTCGCTGCAAATATAGCGCACCTGACTCTTGCTCGTTTGCTTCATG
AATTTGAGCTGGGAGCTCCTGATGATGCACCTTTAGACATGACCGAAAGTCCGGAAGTAAG
TCTCAAAAGAGCAAGCCCTCTCAATGTGCTTATCGCTCCGCGGCTTCCTTCTTCCTGA

>yCYP82BC2
ATGATGATCGAAATCCAAAATTTCTTTCCATCTCATATTGTTTGTGGTGTTTTGTTGGCAGTT
ATTTCAGCTTACTACTTGTTGACTAAGTACAAGAATAAGCCAGCTAGAGCTGCAGCTCCAG
AACCAGCAGGTGCTTGGCCAGTTATTGGTCATATCTTGCAATTCGGTCCATTGGATTTGTGG
CATAGAAAATTGGGTGACATGGCAGATGAAATTGGTCCAGCTTTCGTTATCAGATTGGGCA
TGTTGAGAGTTTTGATTATTAACAACTGGGAATTGGCAAAGGAATGTTTCACTTTGAACGAT
AAGATTTTTGCTTCTAGACCAGGTTCTATCACATCAAAGTACATGGGTTACGATGGTGCAAT
GTTAGGTTTTGCTCCAGATGGTCCTTTTATGTTGGAATTGAGAAAGGTTGCTATGCAAGAAT
TATTTTCTAACCATAGATTGAGATTGTTAAGACATATTAGAACATCAGAAATCGAAATCATG
ATCAAGGGTTTGAATGAATTGTACCATAGAAACTTCAACAACGCAACTGGTGGTGCTTCTA
TGTTAGTTCATTTGGATCCATGGTTGGATGCATTGACAATTAATGTTATGTTGAGATCAATC
GCTGGTAAAAGATATTACGATGGTGGTGCAGCTGTTGAAGGTGAAGAAGCAAAAAGATGG
AAAGAAGCTTTCGGTAAAGTTATGTTGGTTTTTAAAATGTTGTTAGTTTCTGAAATGTTCCC
AAGATTGGAAATGGTTGATGTTATGTTGGGTACTACAAGAGCAATGAAGAAAGCTTACGAA
GAAATGGATTTGTTGTTGTCTGGTTGGTTGGAAGAACATAGATCTAAGAAAGATAGAGGTG
ACCATGAACAAGATTTTGTTGATGTTTTATTGAATGTTGGTAACAACAACCCAAAGTTGTTT
TCTCAATACGATGTTGATTCAGTTGTTAAGGCAACTTGTTTGGATATCTTGATTGCAGCTTCT
GATTCAACTACATTGACATTGTCTTGGGCTTTGTCATTGTTGTTGAACAACAGAAGAGTTTT
GAAGAAAGTTCAAGAAGAATTGGATAACCAAATCGGTAAAGATAGACATGCAGAAGAAGA
AGATATTAAAAATTTGCCATATTTGCAAGCTGTTGTTAAAGAAACTTGGAGATTATACCCAC
CATCTCCATTGTCATTACCACATGAAGCTTCTAAAGATTGTGTTGTTGGTGGTTTCCATATCC
CAAAGGGTACTTCATTGATGACAAACGTTTGGAAGATCCATAGAGATCCAAAGGTTTGGCA
AGAAGATCCATTGGAATTCAAACCAGAAAGATTTTTGTCTGCATCACATGCTCATGTTGATT
TCAGAGGTTTGCATTTCGAATTCATTCCATTTGGTTCTGGTAGAAGAATGTGTTTAGGTACT
AATTTGGCAGCTAACATCGCACATTTGACATTGGCTAGATTGTTGCATGAATTCGAATTAGG
TGCACCAGATGATGCTCCATTGGATATGACAGAATCTCCAGAAGTTTCATTGAAGAGAGCA
TCACCATTAAACGTTTTGATTGCTCCAAGATTGCCATCTTCA

>CYP82BC3
ATGATCGAGATCCAAAACTTCTTTCGATCTGATATTGTATGTGGCGTGCTCTTAGCCTTCAT
CTCAGCCTACTATATTCTCATCAAGTGGAAGAAACACTACCACGAACGCTCCGATGAAACA
AAGAATGCCGCGTGGCCGCGGCCAGCGCCGGAAGCAGCCGGTGCCTGGCCGATCATCGGC
CACGTCCTTCAGTTCGGACCCCTCGACCTCTGGCACCGGAAACTCGGCGACATGGCTGACG
AGGTCGGACCGGCCTTCATGATCCGTCTCGGCATGCGCCGCGTCCTCATCATAAACAACTG
GGAACTAGCCAAGGAGTGCTTCACATCGCACGACAAAACATTTGCGGGCCGTCCAGGCGCA
ATAAGTTCCAAGTACATGGGCTACGGCGGGGCCATGTTGGGCTTCGCCCCTGATGGGCCCT
ACATGGTCGAGTTGCGAAAGGTGGCAATGCAAGAGCTCTTCTCTAACCATAGGCTTCGTGC
GTTGAAGCACATAAGGATGTCCGAAATCGAAATCATGATCAAAAGCCTAAATGAACTTTAT
CATAGAAATTTTAATAATGCTAGGGGTGGTGCTTCAATGCTAGTGCACTTGGATCCGTGGTT
GGATGCATTGACGTTAAACGTGATGCTTCGGAGCATCGCCGGCAAGCGATACTACAACGGA

GATCCGTGCGCGGCGCAAGACGAGGAGGCGAAGAGGTGGAAGGAAGGGTTTGGGAAGCTG
ATTCATGTTTTCAGGACGTTTGTGGTGTCGGAGATGTTTCCGGGGCTGGAATTTGTGGACGT
GATGCTGGGCGCAACGCGTGCCATGGAGAAGGCTAGTAAGGATGTGGACATGTTGTTGAGT
AGATGGGTGGAGGAGCATCGAAGTAAGATGAATAGTAAGGACTACCAAGGGGGTCTTGGT
GAAGGAGATTTTGTTGATGTGTTGCTCAACGTTGACAAGAATGCCAAGTTGTTCTCTCACTA
CGATGTTGATTCTGTTGTTAAAGCCACTTGCTTGGACATATTGATAGCAGCCAGCGATTCAA
CAACACTAACCCTATCATGGGCTCTCTCCTTGTTACTTAATAATCGACATGTCTTAAAGAAG
ATCCAAGAAGAGTTGGACAATCAAATTGGCAAAGGCAGGCACGCAGAAGAAGAAGACATC
AAGAACCTGCCCTACCTACAAGCAGTTGTGAAGGAGACTTGGCGCTTGTACCCTCCTTCAC
CACTTTCATTGCCACATGAGGCCTCGAAAGATTGCATTATTGGTGGATTCCATGTTCCAAAG
GGCACAAGCTTAATGACAAATATTTGGAAGATACATCGCGATCCCAATGTTTGGAAAGATC
CTTTAGAGTTCAAACCCGAAAGGTTTCTTAGTAGCTCTCATGCGCATGTTGATTTTAGAGGT
TTACATTTCGAATTCATCCCTTTTGGGTCAGGTAGAAGAATGTGCCTCGGAACAAACCTTGC
TGCGAATATCGTGCATCTGATGCTTGCTCGTTTGCTTCATGAATTTGAGCTTGGAACCCCTA
ATGATGCTCCCGTAGATATGACTGAAGGCCCGGATGTGAGTCTCAATAGAGCAAGCCCTCT
CAATGTACTTATCGCTCCGAGACTTCCTTCTCCTTGA

>yCYP82BC3
ATGATCGAAATCCAAAATTTCTTTAGATCAGATATCGTTTGTGGTGTTTTGTTGGCTTTTATTTC
TGCATACTACATCTTGATTAAATGGAAGAAACATTACCATGAAAGATCTGATGAAACTAAAAA
TGCTGCATGGCCAAGACCAGCTCCAGAAGCTGCAGGTGCATGGCCAATTATTGGTCATGTTTTG
CAATTCGGTCCATTGGATTTGTGGCATAGAAAATTGGGTGACATGGCTGATGAAGTTGGTCCAG
CTTTTATGATCAGATTGGGTATGAGAAGAGTTTTGATTATTAACAACTGGGAATTGGCTAAGGA
ATGTTTCACTTCACATGATAAGACATTTGCTGGTAGACCAGGTGCAATTTCTTCAAAATATATG
GGTTACGGTGGTGCTATGTTGGGTTTTGCACCAGATGGTCCATACATGGTTGAATTGAGAAAGG
TTGCTATGCAAGAATTGTTTTCTAACCATAGATTAAGAGCATTGAAACATATTAGAATGTCAGA
AATCGAAATCATGATCAAGTCTTTGAATGAATTGTACCATAGAAACTTCAACAACGCTAGAGG
TGGTGCATCAATGTTAGTTCATTTGGATCCATGGTTGGATGCTTTGACATTGAACGTTATGTTGA
GATCTATCGCAGGTAAAAGATACTACAACGGTGACCCATGTGCTGCACAAGATGAAGAAGCTA
AGAGATGGAAGGAAGGTTTCGGTAAATTGATCCATGTTTTTAGAACTTTCGTTGTTTCTGAAAT
GTTTCCAGGTTTAGAATTTGTTGATGTTATGTTGGGTGCTACAAGAGCAATGGAAAAGGCATCA
AAGGATGTTGATATGTTGTTGTCTAGATGGGTTGAAGAACATAGATCAAAGATGAACTCTAAG
GATTATCAAGGTGGTTTAGGTGAGGGTGACTTTGTTGATGTTTTGTTGAACGTTGATAAGAACG
CTAAGTTGTTTTCACATTACGATGTTGATTCTGTTGTTAAGGCAACTTGTTTGGATATCTTGATT
GCTGCATCAGATTCTACTACATTAACATTGTCATGGGCTTTGTCTTTGTTGTTGAACAACAGAC
ATGTTTTGAAGAAAATTCAAGAAGAATTGGATAACCAAATCGGTAAAGGTAGACATGCTGAAG
AAGAAGATATTAAAAATTTGCCATATTTGCAAGCAGTTGTTAAAGAAACTTGGAGATTATACC
CACCATCACCATTATCTTTGCCACATGAAGCTTCAAAGGATTGTATCATCGGTGGTTTCCATGT
TCCAAAGGGTACTTCTTTGATGACAAACATCTGGAAGATCCATAGAGATCCAAACGTTTGGAA
GGATCCATTGGAATTCAAACCAGAAAGATTTTTGTCTTCATCTCATGCTCATGTTGATTTCAGA
GGTTTGCATTTCGAATTCATTCCATTCGGTTCAGGTAGAAGAATGTGTTTAGGTACAAATTTGG
CTGCAAACATCGTTCATTTGATGTTGGCTAGATTATTGCATGAATTTGAATTGGGTACTCCAAA
TGATGCACCAGTTGATATGACAGAAGGTCCAGATGTTTCATTGAACAGAGCTTCTCCATTAAAC
GTTTTGATTGCACCAAGATTGCCATCTCCA

>CYP82BC4
ATGATCCCCCAACCTTACTTTCAACCTGCAATCCTATGCGCCGTGCTCTTAGCTTCGCTCCCC
ATCTTCTACTATCTCCTGATCAGACGGCCCAGAACCACCAACCCCTCCATCAACGGCGTAA
AAAACCTCCCAGAACCAGATGGCAAGTGGCCGATCATCGGCCACATCCTGGCCTTCGGCCC
CAATGACCTCTGGTACCGAAAGCTCAGCGACATGGCGGACAAGGTTGGCCCCATCTTCGCC
ATCCGCCTCGGCAAGCGCCGCGCCGTGGTCATCAACAACTGGGAAATGGCAAAAGAGTGCT
TCACCACCAACGACAAAACGCTCGCGAGCCGCCCATCTTCGTTGCACTCCAAGTACATGGG

CTACAACGGCGCCATGTTCGGGTTCGCCCCCGAAGGGCCGTACCTGCGCGAGGTCCGGAAA
GTGATTACCCAAGAGCTCTTCTCCAACAAGCGTCTCCAGCTGTTGAAGCATGTGAGAGTGT
CGGAGATAGACCTGATGATTAAGGATCTCAAAGAGCTTTATTTCAAGAACAAAGATCATGA
GGATCAGGTAATCAGTGACTCTGATGGGCCGTCAGTGTTGGTAAAGTTAGACACGTCGCTG
CGTGCTCTCACGCTGAACATTATGCTCAGGAATATTGCTGGGAAGAGGTATTACTATGGTG
GGGATATTGAAAGTCACGAGGAACATTATGGCGATGTTGAAGCGAAGAGGTGGAGGGAGA
CTTTTGATAATTTGATTCACACTCTGAGGGCGTTTACGGTGTCGGAGATGTTTCCGGGGTGG
GAGTTTGTGGATTTGGTGATGGGAGTCCAGCGGGGATGAGGAAGGCGAGTGAGGAGATG
GATTTGGTGTTGAGTGGGTGGTTGGAGGAGCATAGGAGGAGGAATGGGAGGAGCGAGGGG
GAGGAGGACTTCGTGGACGTGATGATGAACATTGACTCGAACGCTAAGTTGTTCTCAGAGC
ATGACGTAGACACTGTCGTTAAAGCTACTTGTTTGGATATATTAATAGCGGCTAGAGACAC
AACATCCATCACCCTCCTATGGGTCATCCCCATGTTACTGAACAATCCTCAAGTTCTGAAGA
AGGTGAAAGAAGAGTTGGACAATCAAATAGGCAAAGATAGGCACGCAGAAGAGGAAGAC
ATCCAACACTTGCCTTACCTACGAGCAGTTGTAAAGGAGACTTTGCGCTTATACCCACCTTC
CCCGCTTTCATTACCACATGAGGCATCAAATGACTGTGTCGTTGGAGGGTTCCATGTCCCAA
AAGGGACTTCCTTAATCACAAATATTTGGAAGATACAACGCGATCTCAGTATTTGGCCAGA
CCCTTTGGAGTTCAAGCCAGAGAGATTCCTTGCTACCAATGGCGCCCATAATGTCGATTTTA
GAGGTTTAAACTTTGAATTAATTCCTTTCGGGTCGGGTAGAAGAATGTGTGTGGGAACAAA
CCTTGCCGCTCATATCGTGCACCTAACGCTTGCTCGTTTGTTCCATGAGTTTGAGATCAGAA
CTCCTGACAACGCACCAATTGATATGACCGAAGCTCCAGACACTGATCTCGTCAGAGCAAG
CCCTCTCAGTGTGCTTATACGCCCACGAGTGATTCCTTCTCCTTGA

>yCYP82BC4
ATGATTCCACAACCATACTTTCAACCAGCTATTTTGTGTGCAGTTTTGTTAGCTTCTTTGCCAAT
TTTCTATTACTTGTTGATCAGAAGACCAAGAACTACAAACCCATCAATTAATGGTGTTAAAAAT
TTGCCAGAACCAGATGGTAAATGGCCAATTATTGGTCATATCTTGGCTTTCGGTCCAAACGATT
TGTGGTACAGAAAGTTGTCTGATATGGCAGATAAGGTTGGTCCAATCTTCGCTATCAGATTGGG
TAAAAGAAGAGCAGTTGTTATTAATAACTGGGAAATGGCTAAGGAATGTTTCACTACAAACGA
TAAGACTTTGGCATCTAGACCATCTTCATTGCATTCAAAGTACATGGGTTACAATGGTGCAATG
TTTGGTTTTGCTCCAGAAGGTCCATACTTGAGAGAAGTTAGAAAAGTTATTACACAAGAATTGT
TTTCTAATAAGAGATTGCAATTGTTGAAGCATGTTAGAGTTTCAGAAATTGATTTGATGATCAA
GGATTTGAAGGAATTGTACTTCAAGAATAAGGATCATGAAGATCAAGTTATTTCTGATTCAGAT
GGTCCATCTGTTTTGGTTAAGTTGGATACTTCATTGAGAGCTTTGACATTGAACATCATGTTGA
GAAACATTGCAGGTAAAAGATATTACTATGGTGGTGACATTGAATCTCATGAAGAACATTACG
GTGACGTTGAAGCTAAGAGATGGAGAGAAACTTTCGATAATTTGATCCATACTTTGAGAGCTTT
TACTGTTTCTGAAATGTTTCCAGGTTGGGAATTTGTTGATTTGGTTATGGGTGTTCAACGTGGTA
TGAGAAAAGCTTCTGAAGAAATGGATTTGGTTTTATCAGGTTGGTTAGAAGAACATAGAAGAA
GAAATGGTAGATCTGAAGGTGAAGAAGATTTCGTTGATGTTATGATGAACATCGATTCTAACG
CTAAGTTGTTTTCAGAACATGATGTTGATACTGTTGTTAAGGCAACATGTTTGGATATCTTGATC
GCTGCAAGAGATACTACATCAATCACTTTGTTGTGGGTTATTCCAATGTTGTTGAACAACCCAC
AAGTTTTGAAGAAAGTTAAGGAAGAATTGGATAACCAAATCGGTAAAGATAGACATGCTGAA
GAAGAAGACATCCAACATTTGCCATACTTAAGAGCAGTTGTTAAGGAAACATTGAGATTGTAT
CCACCATCTCCATTGTCATTACCACATGAAGCTTCTAATGATTGTGTTGTTGGTGGTTTTCATGT
TCCAAAAGGTACTTCTTTGATCACAAACATCTGGAAGATTCAAAGAGATTTGTCAATTTGGCCA
GATCCATTGGAATTCAAACCAGAAAGATTTTTAGCTACTAACGGTGTTCATAACGTTGATTTCA
GAGGTTTGAACTTCGAATTGATCCCATTCGGTTCTGGTAGAAGAATGTGTGTTGGTACTAATTT
GGCTGCACATATTGTTCATTTGACATTGGCTAGATTGTTCCATGAATTCGAAATTAGAACACCA
GATAATGCACCAATTGATATGACTGAAGCTCCAGATACAGATTTGGTTAGAGCATCTCCATTGT
CAGTTTTGATCAGACCAAGAGTTATTCCATCACCA

>B2NMT

ATGACTAACAACGTTGGTTGTGGCGAAGTTTTGAAGCCTGGAAAGGCCGAGCTGATTGAGA
AGTTGGTGCTGGGGCTGATACCAGATGAAGAGGTCAAACGCCTCATAAGGGATCAACTCGA
AAGGCGTATTCAGTGGATCTACAACCACAGCTGTGAACAACGCTTCTCTGGGCTCCACAAC
TTTGTTCAGTCTTTGCGACAGACGAGTCACGTCACGGAGACAAATGATTTCAATCCTGATAT
ATATGACATCCCCATCGCTTTTGCGAAGCTTATTAACGGAAGAGCATTGAAATTGAGTTGGT
GTTACTTTGAAGATAAATCAGTCCCATTGGATGATGCTGAGGAAGCAATGTTGGGTTTATAT
TGCGAAAGAGCACAAATAAGAGACGGTGATCAAATTCTAGACCTTGGTTGCGGATATGGAT
CTCTCGCCATTTATATTGCTCGCAAGTATCGTCACTGCCATGTTACTGGAATTACAGATACA
AAGTCCCAGAAAAAGTTCATGGAAGAGCAATGCAAGAGCCAAAACTTGAACAATGTGGAG
GTCATACTGGGGGACATTACCAAAGTTGAGTTGAACAAAGAATTTGACCGTGTAATGGTTA
TAGAAGTCTTTGAGCATATCAAGAATTATGAAGTACTTCTAAAGAAGATATCGAAATGGAT
GAAGGAAGATGGACTTCTTTTCGTGGAAAACATATGCCACAAGAACTTTTCGTACCAAATG
AAGCCTCTCAATGAAGAGGATTGGATTGAAGAAATGATCTTTCCTGATGAGATTGTAACCG
TGGCATCTGCTGACTTGCTGCTGTATTTCCAGAAGGATGTTTCAATTGTGAACCATTGGGTC
CTTAATGGCAAGCACATATCCCGCTCAAGTGAAGAATGGCTGAAGAGACTAGATGATAATG
CTGATGCCGCAAAAGCAATTATCAAGGACTTCTTAGGCAGTGAAGATGAAGCAGTGAAGTG
GATTAATCAGTGGAGATTAAATTTTTACATGGAATAGAGCAAGGCGGATTCCATAATGGT
GAAGAATGGATGGTAGCTCATTTTCTATTTAAAAAGAAATAA

>yB2NMT
ATGACTAACAATGTCGGTTGTGGTGAAGTCTTGAAGCCAGGTAAGGCTGAATTAATTGAAA
AGTTGGTTTTGGGTTTGATTCCAGATGAAGAAGTCAAGCGTTTGATCAGAGATCAATTGGA
AAGAAGAATCCAATGGATTTACAACCATAGTTGTGAACAAAGATTCTCAGGTCTACATAAT
TTTGTTCAATCCTTGAGACAAACCTCTCATGTCACTGAAACCAACGACTTCAACCCAGACAT
CTACGACATTCCTATTGCTTTCGCTAAGTTAATCAACGGTCGTGCCCTTAAGTTGTCTTGGT
GTTACTTTGAAGACAAGTCTGTTCCATTAGATGATGCTGAAGAAGCCATGTTGGGCTTATAC
TGTGAAAGAGCTCAAATTCGTGACGGTGATCAAATCTTGGACTTGGGTTGCGGTTACGGTTC
TTTGGCCATTTACATCGCTAGAAAGTACAGACACTGTCACGTCACTGGTATCACTGACACCA
AGTCCCAAAAGAAATTCATGGAAGAACAATGTAAGTCTCAAAACTTGAACAACGTTGAAGT
TATCCTGGGTGACATTACTAAGGTCGAATTGAACAAGGAATTCGACAGAGTTATGGTCATT
GAAGTTTTCGAACACATCAAGAACTACGAAGTGTTGTTGAAAAAGATCTCCAAGTGGATGA
AGGAAGATGGTTTGTTGTTCGTTGAAAACATCTGTCACAAGAACTTCTCTTACCAAATGAAA
CCATTGAACGAAGAAGACTGGATCGAAGAAATGATTTTCCCAGACGAAATCGTTACCGTCG
CTTCTGCTGATTTGCTATTGTATTTCCAAAAGGATGTTTCCATCGTTAACCACTGGGTTTTGA
ATGGTAAGCACATTTCCAGATCTTCTGAAGAATGGTTGAAGAGATTAGATGACAACGCTGA
CGCTGCTAAAGCTATCATCAAGGACTTCTTGGGTTCCGAAGATGAAGCCGTCAAATGGATT
AACCAATGGAGATTGAACTTTTTGCACGGTATTGAACAAGGTGGTTTCCACAACGGTGAAG
AATGGATGGTTGCTCACTTCTTATTCAAGAAGAAGTGA

>B2′NMT1
ATGCATATGGATTCCGCTAATGTGCAAAAGCCCAACACCCACAATGTTGAGAGAGACGGAG
ACGAAGTCTTCTTGTTTGCGGAACAACTGATCTACGGGTCTATTGTACCGATGGTGATGCGA
GCTGTCATCAAGCTCAACGTTTTGGAGATCATGAATAGAGCACCAACGACTTATCTATCAG
CATATCAGATTGCGTCTCAGATGCCTAACAACAAAAACCCTGATGCGCACATCATCCTTGA
TCGGATACTTCGTTTTCTCGTTAGTCATTCAGTGCTCACTTGCACCACCCAATCAGAGAACA
CTGATCATGATTGTCAAGGCGAAAGGTTATATGGCCTCACACCCGCTTGCAAGTACTTCATC
GAGAATGAGGACGGGGCCTCACTTTCTGCGACCTTGCTTTCCTTGCACAACGAGAGAGCGC
AAGATTCCTGGCTTTACCTAGATGAAGTAGTTACTCAACCCAATTTTTCAGCAACTGAAAAG
GTCTATGGAATGAAAGGCTACGAGCATTTAGCCACAGACAATTCTGAGAGAGACTTATTTG
ACAAATCAATGTCTGATCACACAACAATTATTATGAGAAGGATTCTTGATAAGTACAAAGG
TTTTGAAGACGTAAAAGTTGTTGTTGATGTGGGTGGTGGAGTCGGAACCAACATGAACATG
ATTGTTTCCAAGTACCCTACAATTAAGGGCACTAATTTTGATCTGCCGCATGTCGTTGAATC

TGCACCGTCATATCCAGGTGTCATACATGTTGGAGGAGATATGTTTGCAAGTATTCCAAATG
GAGATGCCATTTTCATGAAGTGGGTTTTCAACAATTGGAGCGATGAAGAATGCTTGATATT
GTTAAAAAATTGTTACGAAGCCCTGCCAGATGGAGGAAAGGTGATTGTTGTAGAGAAGATA
CTTCCAGCAATTCCTGCAGCTAATAATGCAACAAGAACAACATATGCCTTTGATTTGACCTT
GATGACAATCTCACCAAAAATAAGGGAGAGAGCTAAAGAAGAATTTGAGGCCTTGGCTAA
AGGTGCTGGGTTTGTTGGAATTAGCGTGATTTGTAATGTTTTTAACTTTTGGGTCATAGAAT
TTCTCAAATGA

>yB2′NMT1
ATGCACATGGACTCTGCTAACGTTCAAAAGCCAAATACTCACAATGTGGAAAGAGACGGTG
ACGAAGTTTTCTTGTTTGCTGAACAATTGATCTACGGTTCTATCGTTCCAATGGTTATGCGTG
CTGTCATCAAGTTGAACGTTTTGGAAATTATGAACAGAGCTCCAACTACTTACTTGTCTGCT
TACCAAATCGCCTCTCAAATGCCAAACAACAAGAACCCAGACGCTCACATTATTTTAGACA
GAATCTTGAGATTCTTGGTTTCTCACTCTGTCCTAACCTGTACCACTCAATCTGAAAACACT
GACCACGATTGCCAAGGTGAAAGATTGTACGGTTTGACTCCAGCTTGTAAGTACTTCATCG
AAAACGAAGATGGTGCCAGTCTTTCCGCAACCTTATTGTCCTTGCACAACGAGAGAGCTCA
AGATTCTTGGTTATATTTGGATGAAGTCGTTACCCAACCAAACTTTAGCGCCACTGAAAAG
GTCTACGGTATGAAAGGTTACGAACATTTGGCCACCGACAACTCCGAAAGAGATTTGTTCG
ACAAGTCCATGTCTGATCACACAACTATCATCATGAGAAGAATTTTAGATAAATACAAGGG
TTTCGAAGACGTCAAGGTTGTTGTCGATGTTGGTGGTGGTGTTGGTACTAACATGAACATGA
TTGTTTCCAAGTATCCTACCATCAAGGGTACCAACTTCGACTTGCCACACGTCGTTGAATCC
GCTCCATCTTACCCAGGTGTCATTCATGTTGGAGGTGACATGTTCGCTTCCATCCCAAACGG
TGATGCTATCTTCATGAAGTGGGTTTTCAACAACTGGTCTGACGAAGAATGTTTGATTTTGT
TGAAGAACTGTTACGAAGCCTTACCTGACGGTGGTAAGGTTATTGTCGTAGAAAAGATTTT
GCCAGCCATTCCAGCTGCTAATAACGCTACCAGAACCACCTACGCTTTCGATCTAACTTTGA
TGACTATCTCTCCAAAGATCAGAGAACGTGCCAAGGAAGAATTTGAAGCTTTGGCTAAGGG
TGCTGGTTTCGTCGGTATTTCAGTTATCTGTAACGTCTTCAACTTCTGGGTCATTGAATTCTT
GAAATGA

>B2′NMT2
ATGAGTTCCAATATTGTGCACAAACCCAACAATAGCAACGCCGAGAGAGTCGGAGACGAA
GACTTTTTGTTTGCAGAGCAACTCATCTACCTATCTGTTGTACCAATGGTGGTGAGAGCTGC
TATCAAGGTCGGTGTGTTCGAGATCATCAACATGGCCTCTACAACACATATCTCAGCATCA
GAGATCGCGTCGCAGATCCCTAACAACAAAAATCCAGACGCCCATATCATCCTTGATCGAA
TGCTTCGGTTCCTCGCTAGTCATTCACTACTCACTTGCACCATCAACAACAATGAAGACGGT
GAAGTGGAGAGGCTGTATGGCTTGACACCGGCTTCCAAGTACTACATCAAAAATGAAGATG
GTGCCTCGCTCTCTGCTACCTTGATTTCCTTGCATCACGAGAAGGCATTGGATTTCTGGCTTT
ACTTGGATGAGGTGGTTCTTGAACCGGGGCTTTCAGCGATTAGTAAGGCCTATGGAATGAA
TGGGTACGAGTACTTAGCCAACGATGCGGCGGAGAGCGAGTTGTTCAACAAGTCAATGTCA
GATCATACGACGATTGTTATGAGGAAGATTCTTGAGAACTATAAAGGGTTTGATGGCGTGA
AAGTTGTGGTTGATGTAGGTGGTGGGGTTGGAACCAACATTAATATGATTGTTTCCAAGTAT
CCGGCAATTAAGGGCATCAATTTCGAGTTGCCTCACGTGGTTGAAACCGCGCCTTCCTTCCC
TGGTGTTGAACACGTTGGAGGAGATATGTTTGCTAGTGTTTCAAATGGAGATGCCATTTTCA
TGAAATGGGTTCTCAACACATGGAGTGATGAACAATGCTTGACATTGTTGAAGAACTGCTA
CGAGGCCCTTCCGAGCAACGGAAAAGTGATTGTCGTGGAGAAGATACTTCCAACAATTCCT
GAACCGAATCATGCTACAAGAATTACTTATGCCTCAGATTTGGGAATGATGGCATTCTCGC
CGAAATCAAGGGAACGAACTGAGAAAGAGTTCGAGATGTTAGCGAAAGGGTCTGGATTCG
CTAGCATTAGATTGGCGTGTAATGCTTGTAACTTCTGGGTCATCGAATTTCTCAAATGA

>yB2′NMT2
ATGTCTTCCAACATTGTCCACAAGCCAAATAACTCTAACGCCGAACGTGTTGGTGACGAAGAC
TTCTTGTTTGCTGAACAATTGATCTACTTGTCCGTTGTTCCAATGGTTGTTAGAGCTGCTATCAA

GGTCGGTGTCTTCGAAATCATCAACATGGCCTCCACCACTCACATTTCAGCTTCTGAAATTGCT
TCTCAAATCCCAAACAACAAAAACCCAGACGCTCACATTATCTTGGACAGAATGCTAAGATTC
TTGGCCTCTCACTCTTTATTGACCTGTACTATCAACAACAATGAAGACGGTGAAGTTGAAAGAT
TATACGGTTTGACACCAGCTTCTAAATATTACATTAAGAACGAAGATGGTGCTTCTTTGTCTGC
TACCTTGATTTCTTTGCATCACGAAAAGGCTTTGGATTTCTGGTTATACTTGGACGAAGTCGTCT
TGGAACCTGGACTGTCCGCTATCTCCAAGGCCTACGGTATGAACGGTTACGAATACTTGGCTAA
CGATGCCGCCGAATCTGAATTATTCAACAAGTCCATGAGCGACCACACTACCATCGTTATGAG
AAAGATTTTGGAAAACTATAAGGGTTTCGATGGCGTTAAGGTTGTCGTCGATGTTGGTGGTGGT
GTTGGTACCAACATTAATATGATCGTTTCTAAGTACCCAGCCATCAAGGGTATTAACTTTGAGT
TGCCACACGTTGTCGAAACTGCTCCATCTTTCCCAGGTGTCGAACATGTTGGTGGTGACATGTT
CGCAAGTGTCTCCAACGGTGACGCTATTTTCATGAAATGGGTGTTGAACACTTGGTCTGATGAA
CAATGTTTGACTTTGTTGAAGAACTGTTACGAAGCTTTGCCATCCAATGGTAAGGTCATCGTCG
TTGAAAAAATCTTGCCAACCATTCCAGAACCAAACCACGCTACCAGAATCACTTACGCTTCCG
ATCTAGGTATGATGGCTTTCTCCCCTAAGTCTCGTGAAAGAACTGAAAAGGAATTTGAAATGCT
TGCTAAGGGTTCTGGTTTCGCCTCCATAAGATTAGCTTGCAACGCTTGTAACTTCTGGGTTATCG
AATTCTTGAAGTGA

>B12OMT
ATGGATTCCATTATTCGTGATCTAAGTAGCAATGGCAACGGAGACGTCCGCGACGCAGGTT
ACTTGTTCGCGAAGGGACTGGTGAATGCATGCCTTCTACCAATGGTGATGCGAGCTGCTAT
CAAGCTCAATGTGTTTGAGATCATGAACAACGCATCGAAAACTCATCTCTCACCCTCTCACA
TTGCCTCTCAACTCCCTAATAACAAAAACCCAAATGCGCAGTTCGTCCTCGATCGAATGCTT
CGCTTTCTCGCTAGCCATTCAGTTCTCACTTGCACCACAAAAGAGAGTGTTAACAATAGTAA
CAATGGTGAAGTCGAAAGGTTGTACGGCTTGACTCCAGCTTCCAAGTACTTCATCAAAAAT
GAAGATGGAGCCTCACTTGCTGCTTCCCTCTTGGCTTCTACAGACAAGTTGATGTTGGAAAC
CTGTTATTACTTGGATGGTGTTGTTCTCGAACCAGATTTTTCAGTCGATGAGAAGGTTTTTG
GAATGAGTGCCTACCAATATTTTGCCCAAGACCCAGAATTGAACGAGTTGTGCAACAAAAC
CATGTCCGACGAAACTGCAATCACCATGAAGAGGATTCTTGACAAGTACAAAGGGTTTGAT
GGTCTCAAAGTTGTGGTTGATGTGGGTGGTGGGATTGGAACTAACATTAACTTAATTGTTTC
CAAGTACCCTACTATTAAAGGCATCAATTTCGATTCTCCTCATGTGGTTGAAACCGCACCGT
CCTACCCAGGTGTTGAACATGTCGGAGGGGACATGTTTGTTAGTGTTCCAAAAGGAGATGC
CATTTTCATGAAGTGGGTACTTCACAATTGGAAAGATGAGCAGTGCTTGACATTGTTGAAG
AAGTGTCATGAAGCTCTACCGAAGGGAGGAAAGGTGATTGTCGTAGAGGGGTTACTTCCAG
AAGTTCCTACGCCTGACAACGCTACGAAAGATATGTGCGCGTTAGATATAATTATGACAAT
GTCCTTCGGTGCAATGGAGAGAACTGAAAAAGAGTTTGAGACCTTGGCGAAAGTGTCTGGA
TTTGCTGACATTAGGTTGGTATGCAATGCTTGTAATCTGTGGGTCATTGAATTTCTCAAATA
A

>yB12OMT
ATGGACTCCATCATCAGAGACTTGTCTTCTAACGGTAACGGTGATGTCAGAGATGCTGGTTACT
TGTTTGCCAAGGGCCTCGTTAACGCTTGTCTACTTCCAATGGTTATGAGAGCTGCCATTAAATT
AAACGTTTTTGAAATCATGAACAATGCTTCTAAGACTCATTTGTCCCCTTCCCACATTGCTTCTC
AATTGCCAAACAACAAAAACCCAAACGCCCAATTCGTCTTAGACAGAATGTTGAGATTCTTGG
CCTCTCACTCTGTTTTGACTTGTACCACCAAGGAATCTGTCAACAACTCCAACAACGGTGAAGT
TGAACGTTTGTACGGTTTAACTCCAGCTTCCAAGTACTTCATCAAGAACGAAGACGGTGCCTCC
TTGGCTGCTTCTTTGTTGGCTTCCACTGACAAGTTGATGTTGGAAACCTGTTACTACCTAGATGG
TGTTGTCTTGGAACCAGATTTCTCCGTTGACGAAAAGGTTTTCGGTATGTCTGCTTACCAATACT
TTGCTCAAGACCCAGAGTTGAACGAATTGTGTAACAAGACCATGTCCGATGAAACTGCTATCA
CTATGAAGAGAATCTTAGATAAGTACAAGGGTTTCGACGGTTTGAAGGTGGTCGTCGATGTTG
GTGGTGGTATTGGTACCAACATTAATTTAATTGTCAGCAAATATCCAACTATCAAGGGTATCAA
CTTCGATTCTCCTCATGTCGTTGAAACCGCTCCAAGTTACCCAGGTGTCGAACACGTCGGTGGT
GACATGTTCGTTTCTGTTCCAAAGGGTGACGCTATTTTCATGAAATGGGTTTGCACAACTGGA

AGGATGAACAATGTTTGACCCTGTTGAAGAAGTGTCACGAAGCTTTACCAAAGGGTGGTAAGG
TTATCGTTGTTGAAGGTTTGTTGCCAGAAGTCCCAACCCCAGACAACGCCACTAAGGACATGTG
TGCTTTGGACATTATTATGACCATGTCTTTCGGTGCTATGGAACGTACAGAAAAGGAATTCGAA
ACTTTGGCTAAGGTCTCTGGTTTCGCTGATATCAGATTGGTCTGTAATGCATGCAACTTGTGGG
TTATTGAATTCTTGAAATGA

>B7OMT
ATGGAGAAGTGTGAGAGGAACGCAAAAGATCATTCCTTAGCAAGGAAAGAAGATCAAGAA
GAGTTGATCAGAGGTCAAGTGCAGGTATGGAATCACATGTTCAGCTACTTTGAGACAATCA
TGCTCAGGTTGGCAATTCAGCTAGGCATACCTGACCTAATCCACGACCACGGCAGTCCCAT
AACGCTGACCGAACTGGCTAGCAAATTGCCAATCGAAACCCTAAACCTAGACAGGTTCAAG
CAAACATTGAAGTTCATGGTGCATGTGAATCTGTTCACGGAAACAACAGATGAAGTGAGCG
GAGAGACCAAGTACGGTCTCAGTCCAGCAACAAAGCTCCTCCTCAAAACTAATGCTGGTAA
GAATAACAAGAGCTTGGCCAAGTTGGTGTTGCTGAATACAGACCCAACAGAGCTTTCTGTG
GCTGGGCGTCTACTTGAGAGCTTAGGAGGGACGAAGTCGTGTATCGAGTTGTTTTTGGGAT
GGACAGAACGAAAGCCATCGACGAGATGCAGACGGATGCGGAGTGGAATGCTATGGCGAT
CGACGGAATGGACAGTGGCACTGGGATCATGGTGGACGCTCTAGTTGAGGGGCTGAGAAG
AGAGAAGGTCATTGATGAGGGTGCTGCGTCGCTTGTGGATGTCGGCGGAAACTCCGGTGCT
GTAGCAAAGGCGATTGCCAATGCATTCCCACATCTCAAGTGCTCTGTGCTGGATACGGTTC
AGATTGTTAAAAGCGTACCCAAGGACCCTCGAGTGGAGTTCGTCGCTGGTGACATGTTTAT
TGCTATCCCTAATGCGGACATTGTGCTGTTGAAGAGTGTTTTGCATGACTATGAAGATGATG
TGTGCATAAAGATTCTGAAGAAGTGCAAAGAAGCAATAAATCCAACAAAGGGGAAGCTGC
TACTGGTTGACATTGTGGTGGACAGTGAGAACACGCCAGAGTTCTCCCGAGCAAGAATGGG
TATGGCCGTGAGCATGATGATTGCCGGTGGAAAAGAAAGGAGTACAAAAGAGTGGGAAAA
TCTCATCTACGGAGCTGGTTTTAGTCGGTATAAGATAATACCAATTGTAGCCGTTGAATCTG
TTCTAGTCGTGTACCCATAA

>yB7OMT
ATGGAAAAGTGTGAAAGAAACGCCAAGGACCATAGTTTGGCCCGTAAGGAAGACCAAGAAGA
ATTAATTAGAGGTCAAGTTCAAGTCTGGAACCACATGTTTTCTTACTTCGAAACCATTATGTTG
AGATTAGCCATCCAATTGGGTATTCCAGACTTGATTCACGATCACGGTTCCCCTATCACCTTAA
CTGAATTGGCTTCCAAGTTGCCAATCGAAACTTTGAATTTGGACAGATTCAAGCAAACTTTGAA
GTTCATGGTCCACGTTAACTTATTCACTGAAACTACTGATGAAGTTTCTGGTGAAACCAAGTAC
GGGTTGTCTCCAGCTACTAAATTGCTGTTGAAGACCAACGCCGGTAAGAACAACAAGTCTTTG
GCTAAATTGGTTTTGTTGAACACTGACCCAACCGAACTTTCCGTTGCTGGTAGATTGTTGGAAT
CTTTGGGTGGTACTAAGTCTTGTATTGAATTGTTCTTCGGTATGGACAGAACCAAAGCTATCGA
TGAAATGCAAACGGATGCTGAGTGGAACGCAATGGCCATTGATGGTATGGATTCCGGTACCGG
TATCATGGTTGACGCTTTGGTTGAAGGTTTGAGAAGAGAAAAGGTTATCGACGAAGGTGCTGC
TTCTTTGGTCGATGTTGGTGGTAACTCTGGCGCTGTCGCTAAGGCTATTGCTAACGCTTTCCCAC
ATTTGAAATGCTCTGTCTTAGACACCGTCCAAATCGTCAAGTCCGTTCCAAAGGACCCAAGAGT
TGAATTCGTTGCTGGTGACATGTTCATCGCCATTCCAAATGCTGACATTGTCTTATTGAAATCCG
TTTTGCACGACTACGAAGATGACGTCTGTATCAAGATTCTAAAGAAGTGTAAGGAAGCTATCA
ACCCAACAAAGGGTAAGTTGTTATTGGTTGACATCGTTGTTGATTCTGAAAACACCCCAGAATT
TTCTCGTGCTAGAATGGGTATGGCTGTTTCCATGATGATTGCTGGTGGTAAGGAAAGATCAACT
AAAGAATGGGAAAACTTGATCTATGGTGCCGGTTTCTCTCGTTACAAGATCATCCCAATTGTCG
CTGTCGAATCCGTGCTAGTTGTCTACCCATGA

>BBOX
ATGGAGGGATGGGGCCTGGTCTCAACGTTTTCATCTGTTTTTCTTCTGGTCTGCTCGGTGCTATC
ATGTTGTTCAGCCTCTTCAATCTCAACAGCTCTAGAGAATAATTTCCTTGAATGTCTGTCCTACT
CCAATGTACAAGCGTACACCCCTGACAGTGCTCCCTACACATGGATTCTACAATCTTCCGTTCA

AAACCTCAGATTCGCATCACCCACGACGCCAAAACCTCAATTCATAGTCACTCCTTCCCATGAG
TCCCAGGTTCCAGCAATTGTTGTTTGTTCTAGGAAACATGGGTTGCAAATAAGAGTTCGGAGTG
GAGGCCACGACTTCGAAGGCCTCTCCTACACATCACACGTTCCATTCGTGATGATCGACCTCGT
CAATCTTCGAACCATCAATGTTGACATAGAAAACAATAGCGCGTGGGTTCAGGCTGGGGCGAC
CCTTGGCGAAGTTTACTATCGAATTGCAGAGAAAAGCCCGATCCATGGTTTCCCAGCTGGTTTC
TCCTCCACTGTTGGTGTTGGTGGGCATATTAGCGGAGGCGGATGTGGTGCCATGGCGAGAATG
CACGGCATGGCGGCAGATCATGTTATTGACGCTCGACTAGTCAATGCGGAGGGAGAAATTCTT
GATAGGGAAAGCATGGGAGAAGGTTTGTTCTGGGCCATTCGAGGAGGAGGCGCCGCAAGTTTC
GGGGTCGTCCTTGCATTCAAGATCAAACTAGTCCAAGTTCCTCCTGTTGTGACAGCTTCAAAGG
TGAATATGACCTTAGAACAAGGTGCACTTGAGATAATTCACAAATGGCAATCTACTGCAAGTT
ATGAGGTTCCCAAAGAGCTTTTCATTCAAGTTTTTATTTATCCTATGACGACTAACAATAATAG
TACTGCCACTATAGAAGGATTTGGAAGAATTTTGCAGGTTGAATTTCAAATGATGTTTCTAGGA
CCAAAGGAGGAGCTTGTTCATTTGATGCAAGACAAGTTTCCCGAATTGGGAATAAAGCAGAAG
CACTGTAAGGAGATGAGTTGGATAGAATCTGCAATTTTTTTCGGCGCATTTCCTAATGGCGCAC
CTCTAGAAGTGTTACTAGATCGAACTCTGCCAAACAAGGATGCCTTCAAAACAAAATCCGACT
TCGTGACAAAGCCGATGTCGAAAGTTGAATTGAAACAAATCTTCAACAAGTCCTTAGAAGAAG
AAAAGCTTAAGGTGGTGTTGAGTCCATGGGGAGGAAGGATGAGTGAGATTTTAGAAGATGAG
ATTGCTTTTCCACACAGGGGTCACAACTTATACATGATTGAGTATTCAGTTGTGTGGGATGAAG
ATGAAGAAAGGACTAAAGCTTCTCAAAAGTACATAAGTTGGATTAGAGAATTGTATCTTTACA
TGAGTCCTTATGTTTCCAAATCTCCAAGAGCCGCATATTTCAATACTAGAGATCTTGATTTGGG
ACAAAACAAGATTAATTCGACTCCAACATACTGTGAAGCAAGGATTTGGGGAGAAAAGTACTT
CAAGGGTAACTTTGAGAAATTAGTGAAGGTGAAGAGTGAGGTTGATCCTGATAATTTTTTCAGT
AATGAGCAGAGTATCCCTCCGATGACACCCCAATCAATGGAAAGCACGTACTGA

>yBBOX
ATGGAAGGTTGGGGTTTGGTTTCTACTTTCTCCTCTGTTTTCTTGCTAGTCTGCTCTGTCTTGTCT
TGTTGTTCTGCTTCTTCTATTAGCACTGCTTTGGAAAATAACTTTTTGGAATGTTTGTCCTACTCT
AACGTCCAAGCTTACACCCCAGATTCCGCCCCATACACCTGGATTTTGCAATCTTCGGTTCAAA
ACTTGAGATTCGCCTCTCCAACTACCCCAAAGCCACAATTCATTGTCACTCCATCTCACGAATC
ACAAGTCCCAGCCATAGTTGTCTGTTCCAGAAAGCACGGTCTGCAAATCAGAGTTAGATCTGG
TGGTCATGACTTTGAAGGTTTGTCCTACACTTCCCATGTTCCATTCGTCATGATTGACTTGGTCA
ACTTGCGTACTATCAACGTTGATATTGAAAATAACTCTGCTTGGGTTCAAGCTGGCGCTACTTT
GGGTGAAGTCTACTACAGAATTGCTGAAAAATCTCCAATCCACGGTTTCCCTGCTGGTTTCTCC
TCTACTGTCGGAGTCGGTGGTCACATTAGTGGTGGTGGTTGTGGTGCTATGGCCCGTATGCACG
GTATGGCTGCTGACCACGTCATTGATGCTAGATTAGTTAACGCTGAAGGTGAAATTCTTGATAG
GGAATCCATGGGTGAAGGCTTATTCTGGGCGATCCGTGGTGGTGGTGCAGCCTCTTTCGGTGTT
GTTTTGGCTTTCAAGATCAAGTTAGTCCAGGTTCCACCAGTTGTTACTGCTTCCAAGGTTAACAT
GACTTTGGAACAAGGTGCTTTGGAAATCATTCACAAGTGGCAATCTACCGCCTCATACGAAGT
CCCAAAGGAATTGTTTATCCAAGTCTTCATCTACCCAATGACCACTAACAACAACTCTACAGCT
ACCATAGAAGGTTTTGGTAGAATCTTGCAAGTTGAATTCCAAATGATGTTCTTGGGTCCAAAAG
AAGAATTGGTTCATTTGATGCAAGACAAGTTCCCAGAATTAGGTATCAAGCAAAAGCACTGTA
AGGAAATGTCCTGGATTGAATCTGCTATCTTCTTTGGTGCTTTTCCAAACGGTGCTCCATTGGA
AGTCTTATTGGACAGAACCTTGCCAAACAAGGACGCTTTCAAGACCAAGTCAGATTTCGTCAC
CAAGCCAATGTCTAAGGTTGAATTGAAGCAAATCTTCAACAAGTCTTTAGAAGAAGAAAAGTT
GAAAGTTGTCCTATCTCCATGGGGTGGTAGAATGTCCGAAATTTTGGAAGATGAAATTGCCTTC
CCTCACAGAGGTCACAACTTATACATGATCGAATACTCTGTTGTCTGGGACGAAGACGAGGAA
AGAACTAAGGCTTCCCAAAAGTACATCTCTTGGATCAGAGAATTGTACTTGTATATGTCTCCAT
ACGTTTCCAAATCCCCAAGAGCTGCTTATTTCAACACCAGAGACTTGGATTTGGGTCAAAACAA
AATTAACTCCACCCCAACCTACTGTGAAGCCAGAATCTGGGGCGAAAAGTACTTCAAGGGTAA
CTTCGAAAAGTTGGTGAAGGTTAAGTCTGAAGTTGACCCAGACAACTTCTTCTCCAATGAACA
ATCCATCCCACCTATGACTCCTCAATCTATGGAAAGTACTTACTGA

>BBR
ATGTGTTACCATCAACATTTCAACATAAGTAGTCACCATTCTTCCATCTCTCTTTCTCATCAATT
TAGTCGATCAAGATTAATGGCGAGTGAGAAGAGCAGAGTGCTTGTGTTGGGTGGTACAGGGTA
CCTAGGCAAGAGACTGGTGAAGGCAAGCCTACGCGAAGGCCATCCCACTTATGTGCTTCAGAG
GCATGAGAATTGCATGGACGTCAACAAGATTCAGCTTCTCCTCAAATTCAAGAAGCGAGGGGC
AAAGCTTATTGAAGGGTCGCTCTCGGATCATGAGAGTCTTGTGCATGCTGTGAAGCAAGTTGAT
GTCGTTATTTCCGCCATATCTCAGTATGACGCGAGTACCATCGCCGCGAAGTTGGAGCACAAG
ATCATCCATGCCATCGAAGAAGCTGGAAATGTCAAGAGATTCGTGCCGGCTGAGTACTGCATG
GATCCGGCAAGGATGTTCCATACACTTGAGCCTGCAAAAGCCTTTGATGTAAGAATGATGGCC
GTGAGAGAAGCTGTTCAACGTGCCAAGATACCATACACTTCGGTCGCTGCTAACCTGTTTGCTT
CAGTCTTCGTACCTAATCTCTGCCAACTAGATACGTGCATGCCTCCAAAGGATAAAGTGTGCAT
ATTTGGAGATGGCAATGCAAAAGGAATTTTTTTGGATGAAGATGATATAGCAACATATACAAT
CAAAACTATAGATGATCCACGCACAGTGAACAAAACTCTCCACATCATGCCGTCAGAAAACAT
CGTCTCCCAAAATGAGTTGATTGATATTTGGGAGAAGCTAAGTGGAAAAAAGCTTAACAAGTC
AATTACCTCTCTCGAAGACTTCTTGGCAATGATGAAAGGAGAGGATTACAATACTCAATTTCTG
ATTGCCCATCTCCACCCCATCTACTACAAGAATTGCCTAACAAATTTTGAAATAAGCGATGGAG
AGGAGGAAGCTTCAAAGCTCTATCCAAATGTCAAGTATACCCGAGTTGAGGAATACTTGAAAT
GGTTTCTATAA

> yBBR
ATGTGTTACCATCAACACTTCAACATTTCCTCCCACCACTCCTCTATTTCGTTGTCCCATCAATT
TAGTCGTTCCAGATTGATGGCGTCTGAAAAGTCTAGAGTTTTGGTCTTGGGTGGTACCGGTTAC
TTGGGTAAGAGATTGGTTAAAGCTTCATTGAGAGAAGGTCACCCAACTTACGTCTTGCAAAGA
CATGAAAACTGCATGGATGTTAACAAGATCCAATTATTGTTGAAGTTCAAGAAAAGAGGTGCT
AAATTGATCGAAGGTTCTTTGTCTGACCACGAATCTTTGGTCCACGCTGTCAAGCAAGTTGACG
TGGTTATCTCTGCTATCTCTCAATACGATGCCTCCACTATCGCCGCCAAGCTAGAACACAAGAT
TATTCACGCCATTGAAGAAGCTGGTAACGTTAAGAGATTCGTTCCAGCTGAATACTGTATGGAC
CCTGCTAGAATGTTCCACACTTTGGAACCAGCTAAGGCTTTCGATGTTAGAATGATGGCTGTCA
GAGAAGCTGTTCAAAGAGCTAAAATCCCATATACTTCTGTTGCTGCTAATTTATTTGCTTCCGTT
TTCGTCCCAAACTTGTGTCAATTGGACACTTGTATGCCACCAAAGGACAAGGTCTGTATTTTCG
GTGACGGTAACGCTAAGGGTATCTTCTTAGATGAAGATGACATTGCTACCTACACCATCAAGA
CCATTGACGACCCAAGAACCGTTAACAAGACTTTGCACATCATGCCATCTGAAAACATTGTTTC
CCAAAACGAATTGATTGATATCTGGGAAAAGTTGTCTGGTAAGAAGTTGAACAAATCTATCAC
CTCTTTAGAGGATTTCTTGGCCATGATGAAGGGCGAAGACTACAACACTCAATTCTTGATTGCT
CACTTGCATCCAATCTACTACAAGAACTGTCTTACCAACTTCGAAATCTCTGATGGTGAAGAAG
AAGCCTCCAAGTTGTACCCAAATGTCAAGTACACACGTGTCGAAGAATATTTAAAGTGGTTCCT
ATG

### NMR spectrum

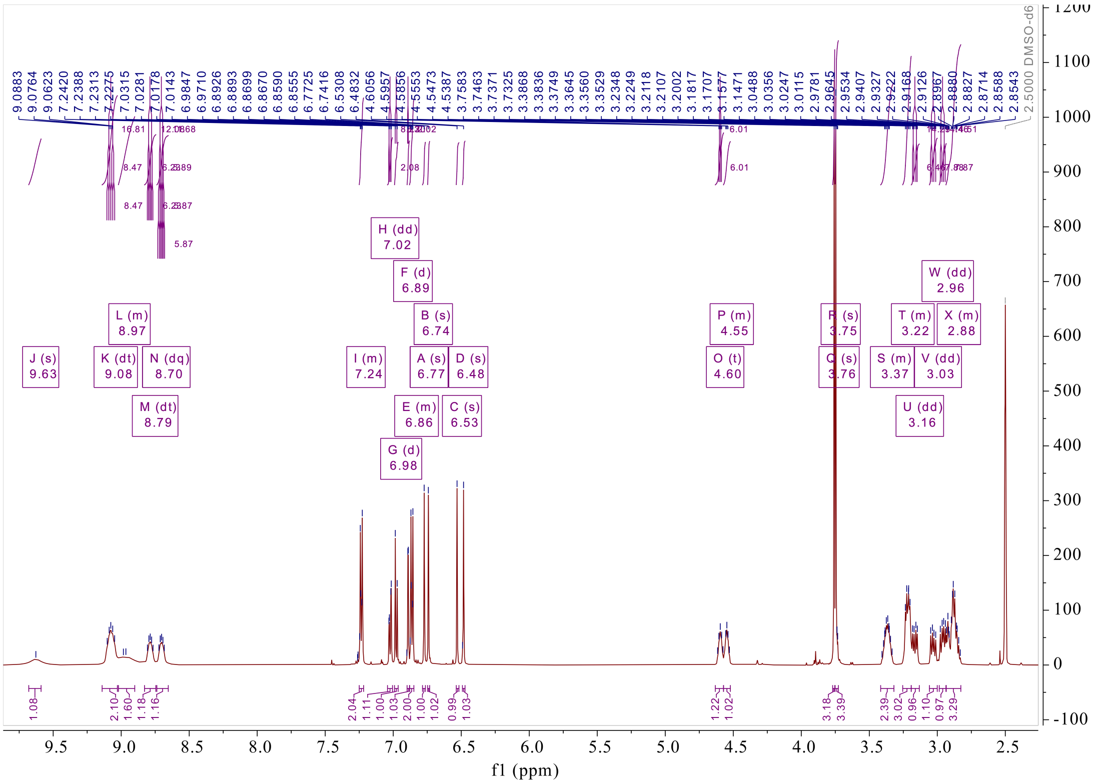


**Supplementary Fig. 30. $^{1}$H NMR spectrum (600 MHz) of (*R*,*S*)-8 in DMSO-$d_6$**

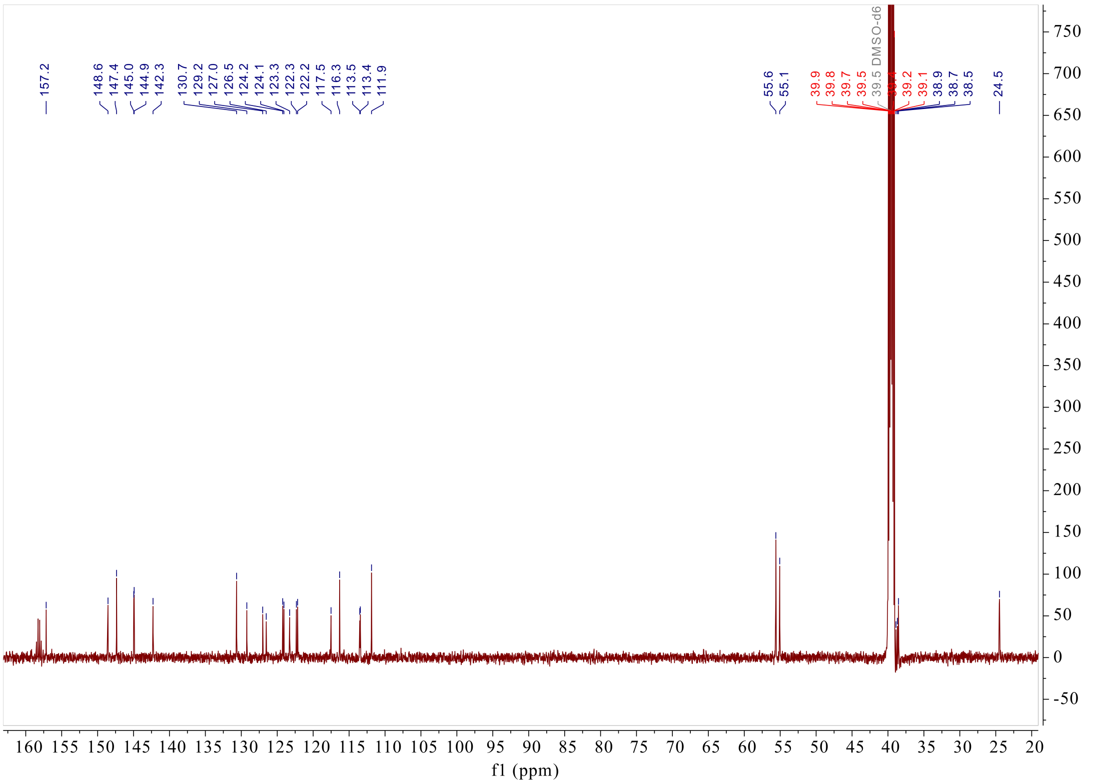


**Supplementary Fig. 31. $^{13}$C NMR spectrum (150 MHz) of (*R*,*S*)-8 in DMSO-$d_6$**

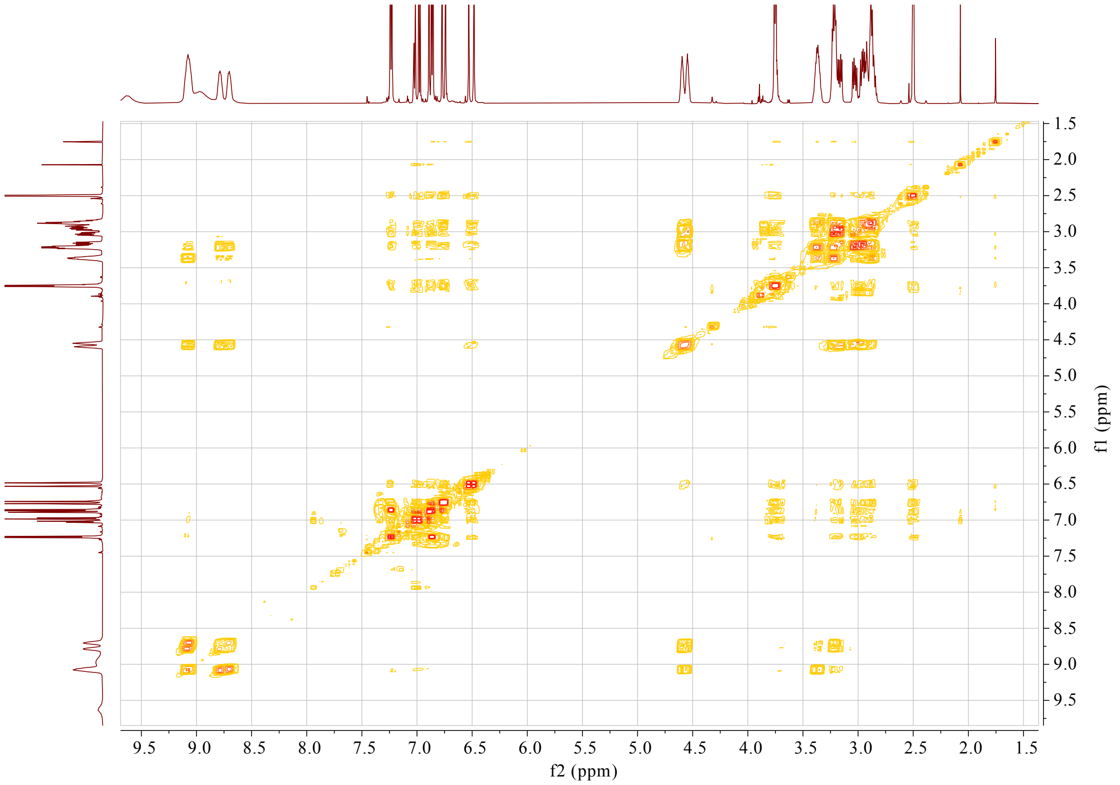


**Supplementary Fig. 32. $^{1}H$-$^{1}H$ COSY spectrum (600 MHz) of (*R*,*S*)-8 in DMSO-$d_6$**

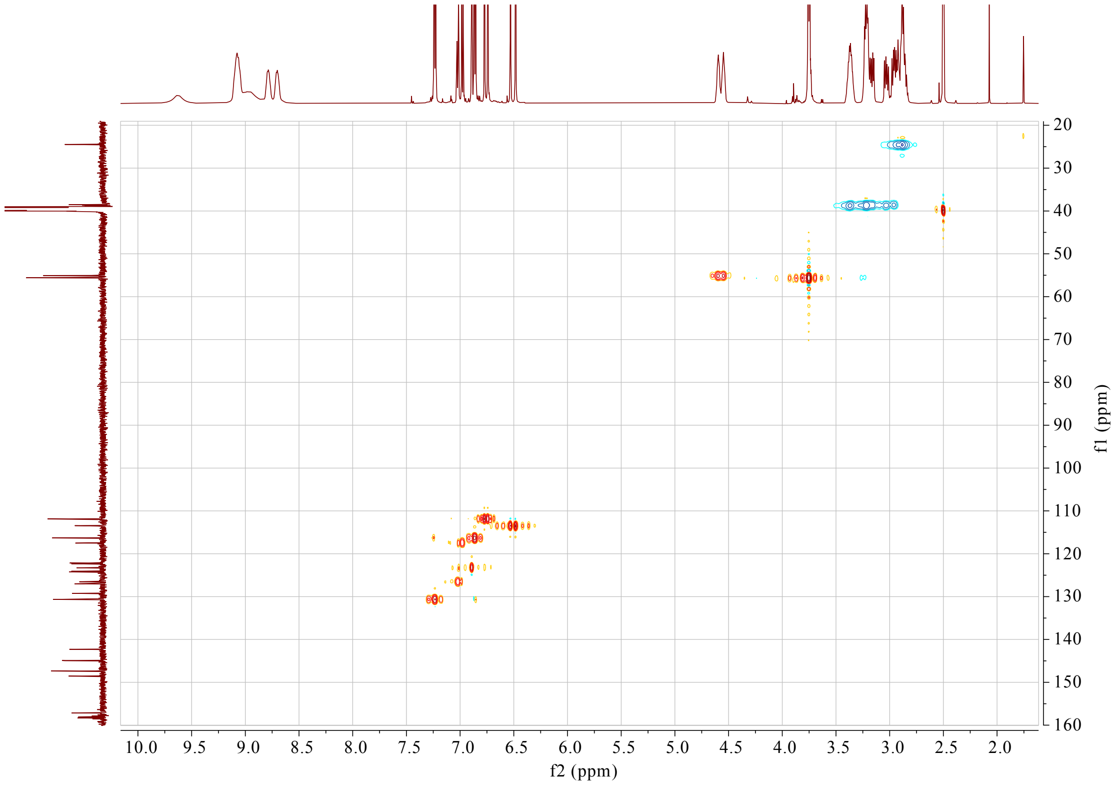


**Supplementary Fig. 33. HSQC spectrum (600 MHz) of (*R*,*S*)-8 in DMSO-$d_6$**

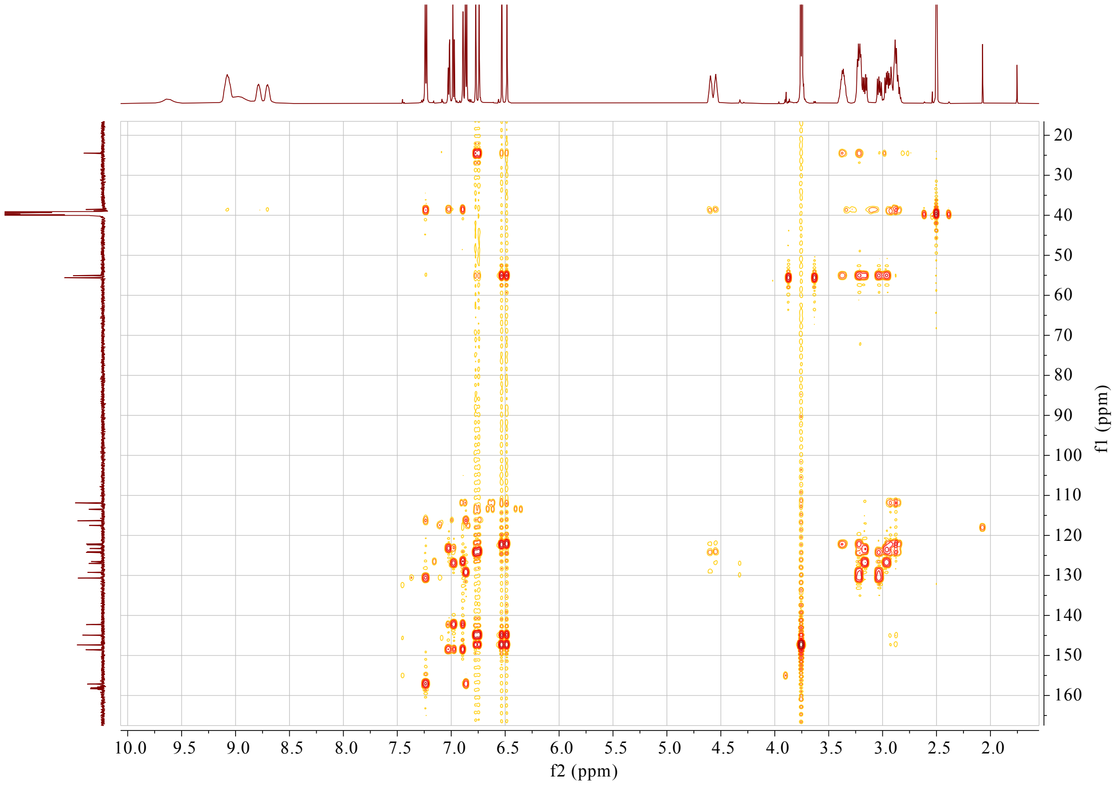


**Supplementary Fig. 34. HMBC spectrum (600 MHz) of (*R*,*S*)-8 in DMSO-$d_6$**

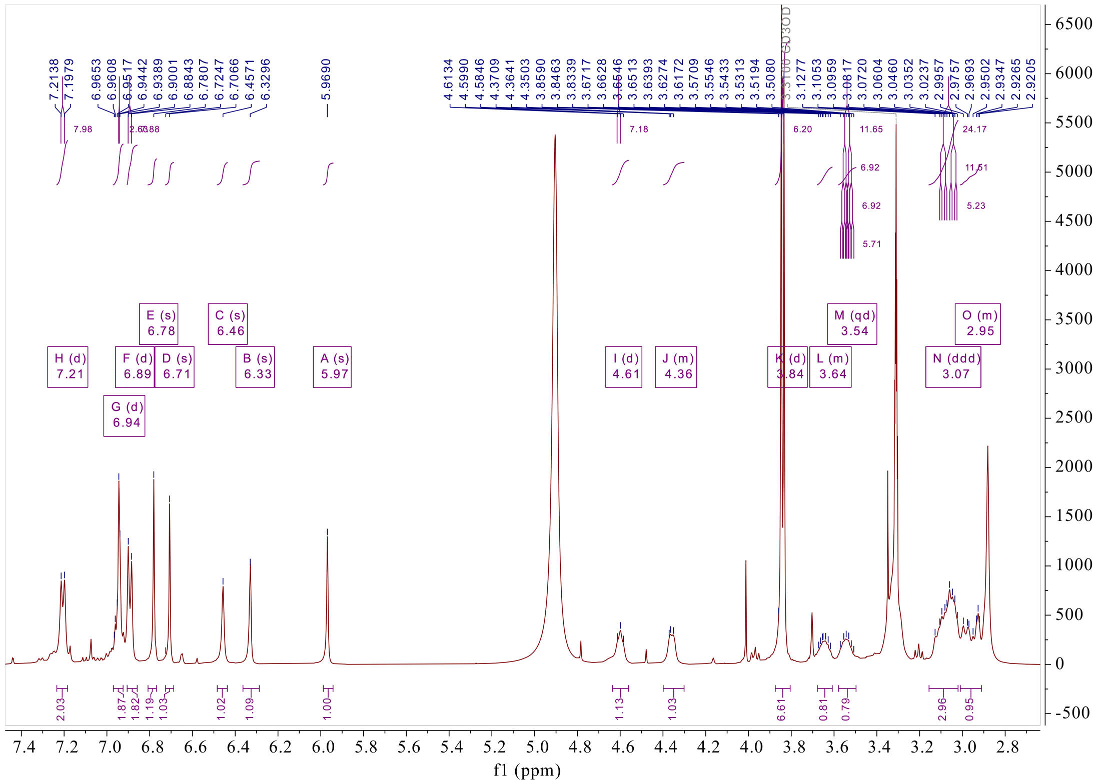


**Supplementary Fig. 35. $^{1}$H NMR spectrum (500 MHz) of (*R*,*S*)-7 in DMSO-$d_6$**

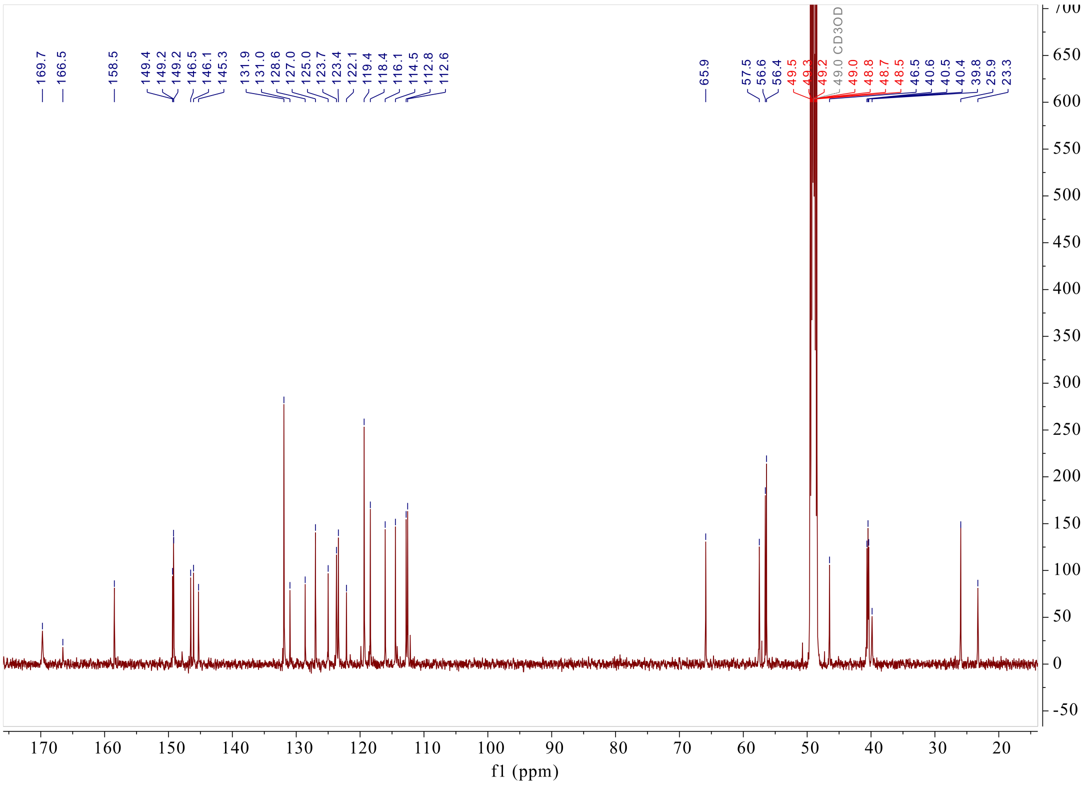


**Supplementary Fig. 36. $^{13}$C NMR spectrum (125 MHz) of (*R*,*S*)-7 in DMSO-$d_6$**

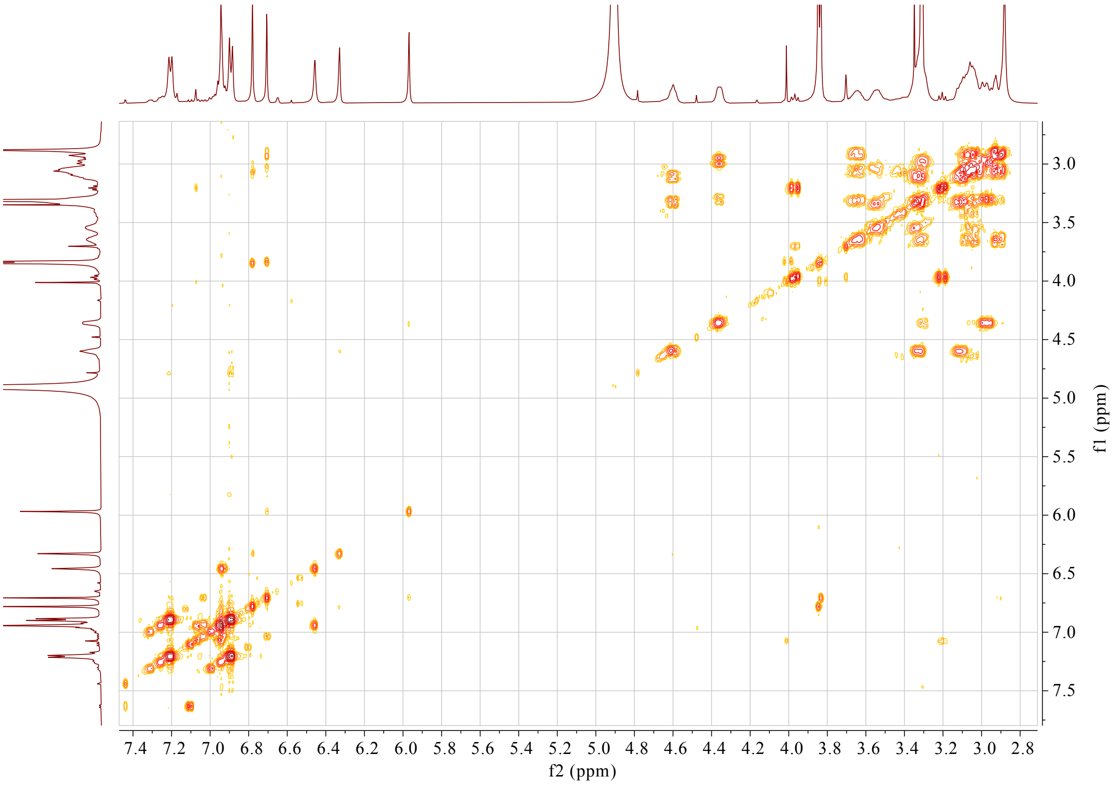


**Supplementary Fig. 37. $^{1}$H-$^{1}$H COSY spectrum (500 MHz) of (*R*,*S*)-7 in DMSO-$d_6$**

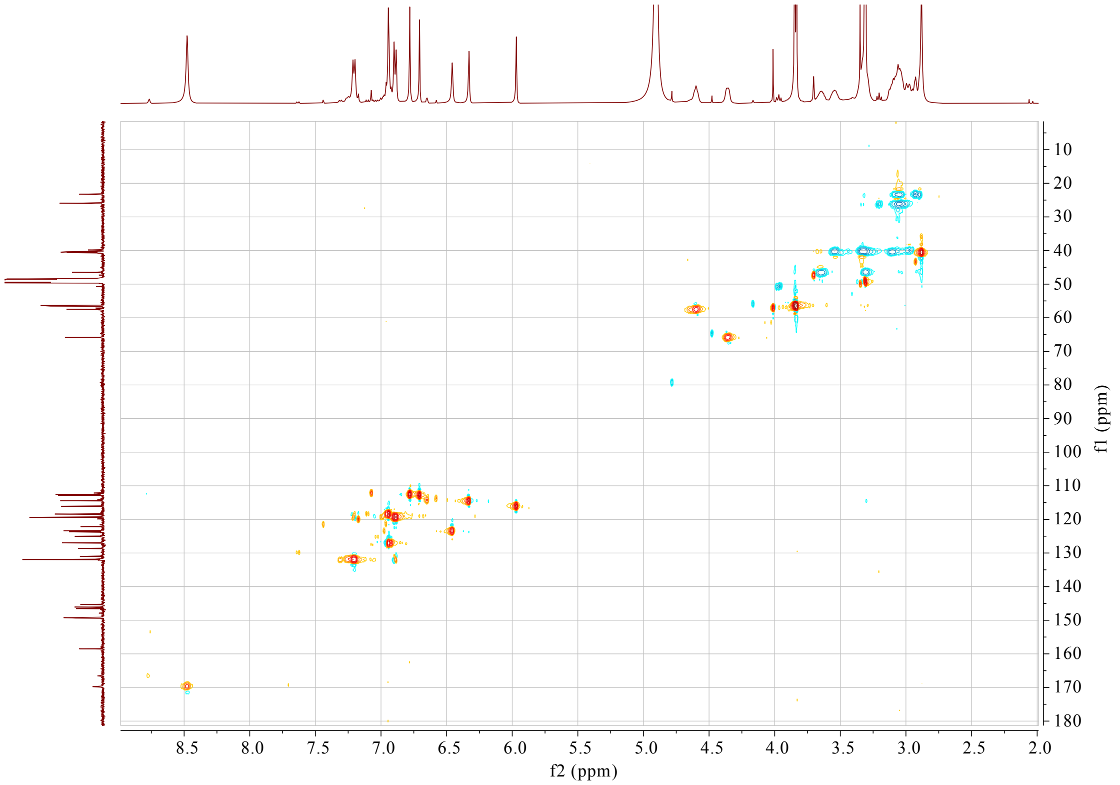


**Supplementary Fig. 38. HSQC spectrum (500 MHz) of (*R*,*S*)-7 in DMSO-$d_6$**

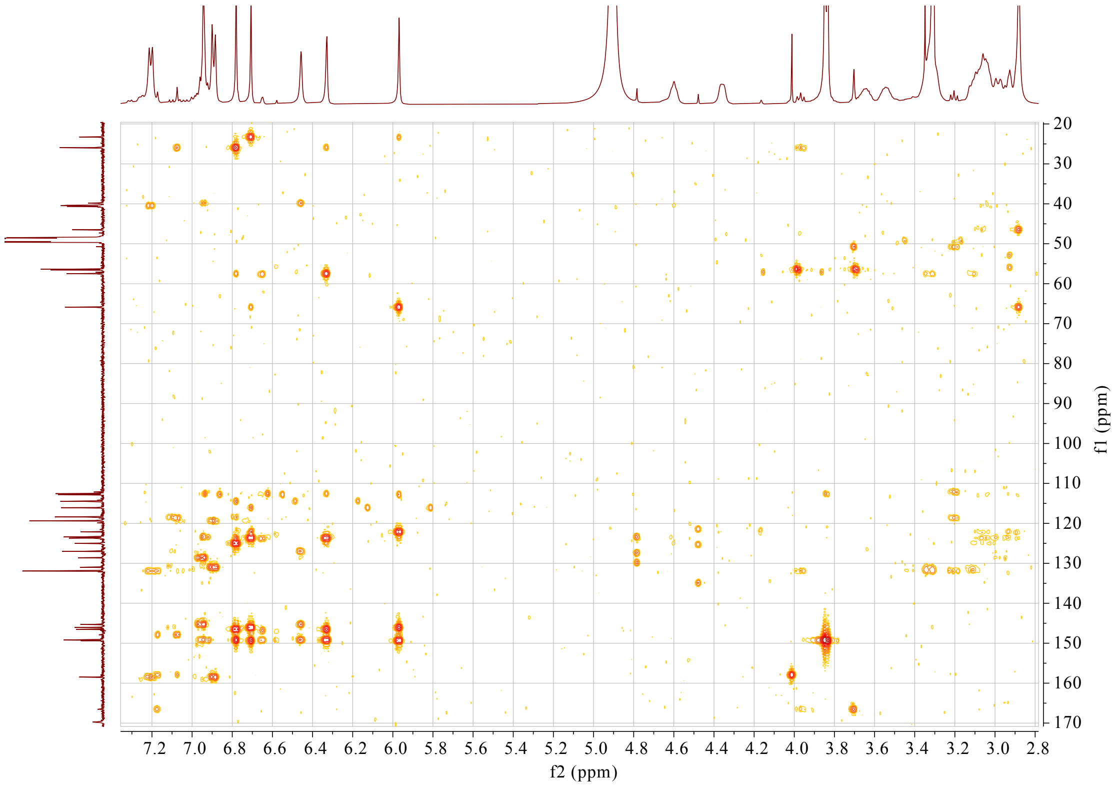


**Supplementary Fig. 39. HMBC spectrum (500 MHz) of (*R*,*S*)-7 in DMSO-$d_6$**

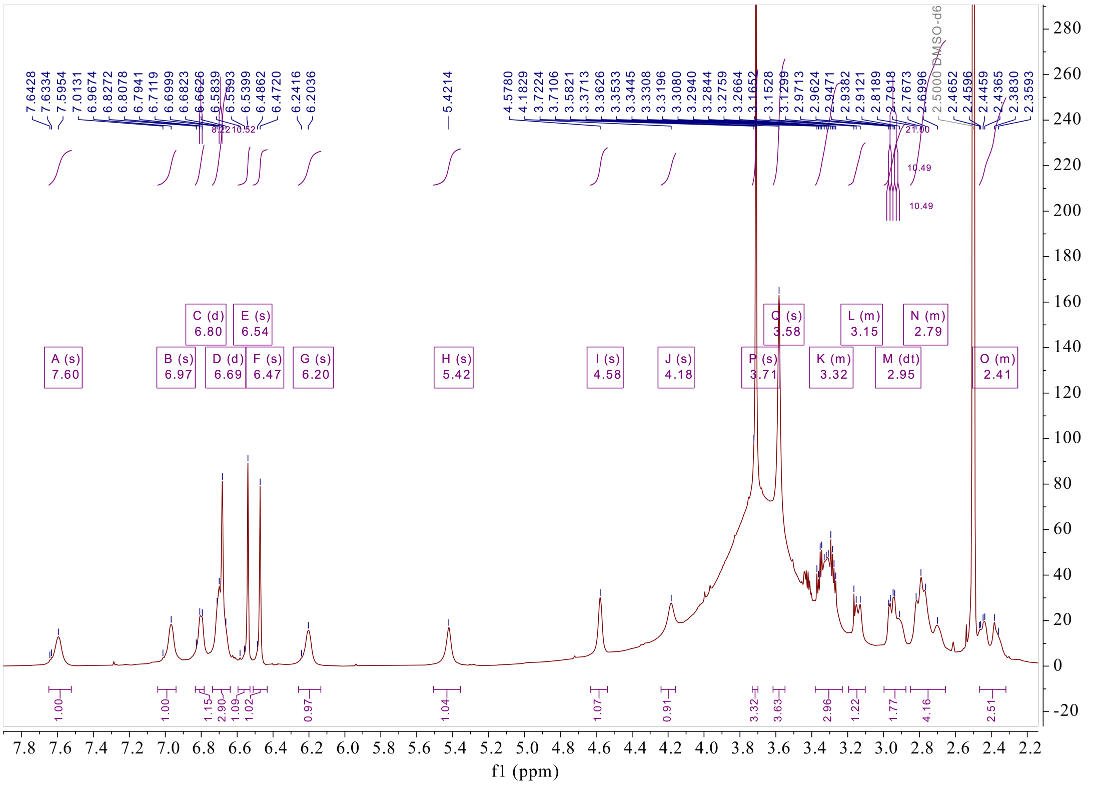


Supplementary Fig. 40. [1]H NMR spectrum (600 MHz) of (*R*,*S*)-16 in DMSO-$d_6$

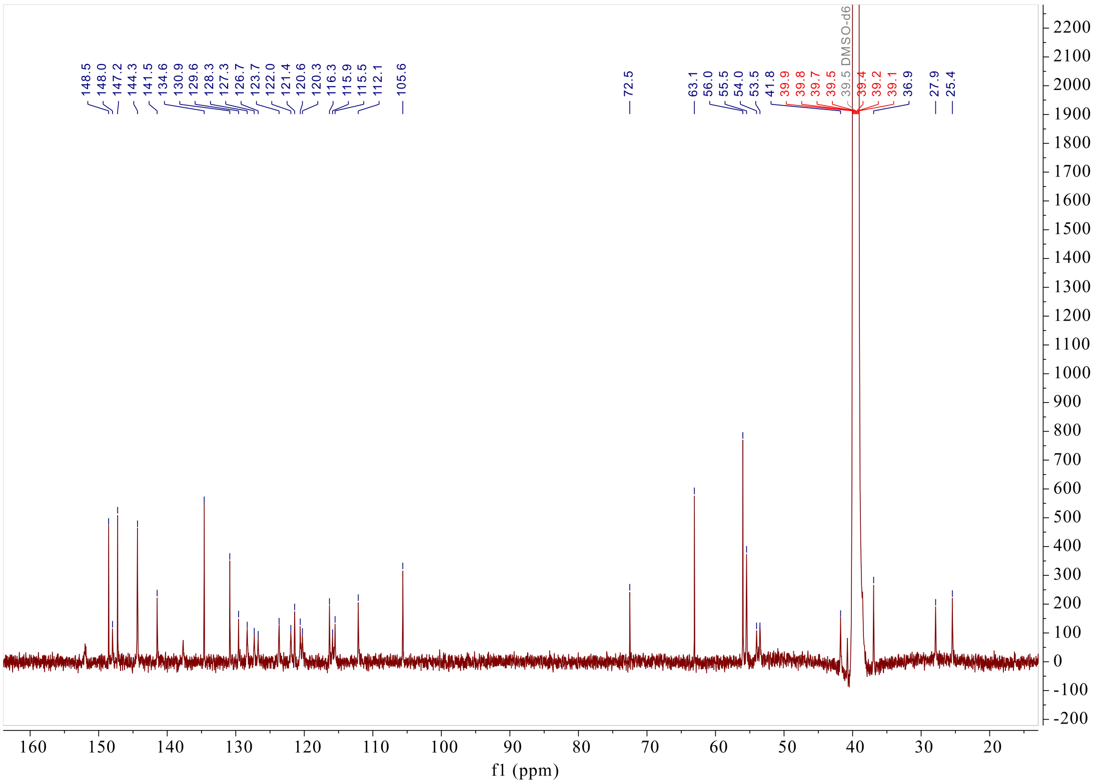


Supplementary Fig. 41. [13]C NMR spectrum (150 MHz) of (*R*,*S*)-16 in DMSO-$d_6$

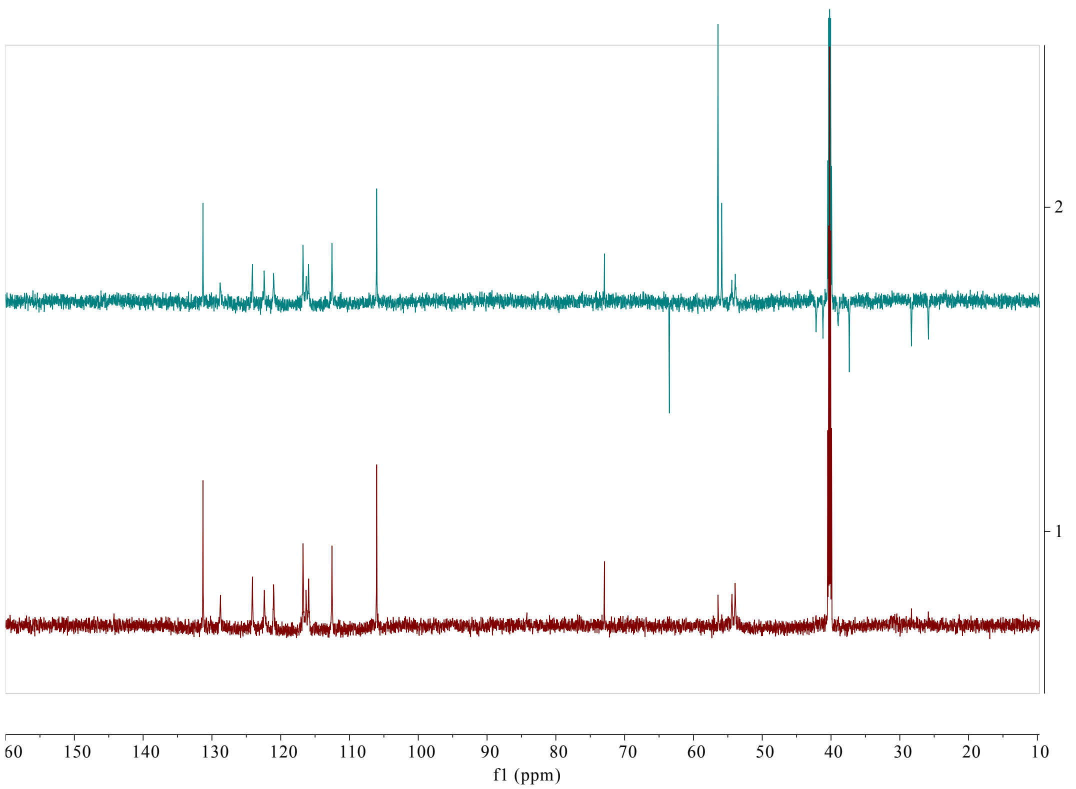


**Supplementary Fig. 42. Dept spectrum (150 MHz) of (*R*,*S*)-16 in DMSO-$d_6$**

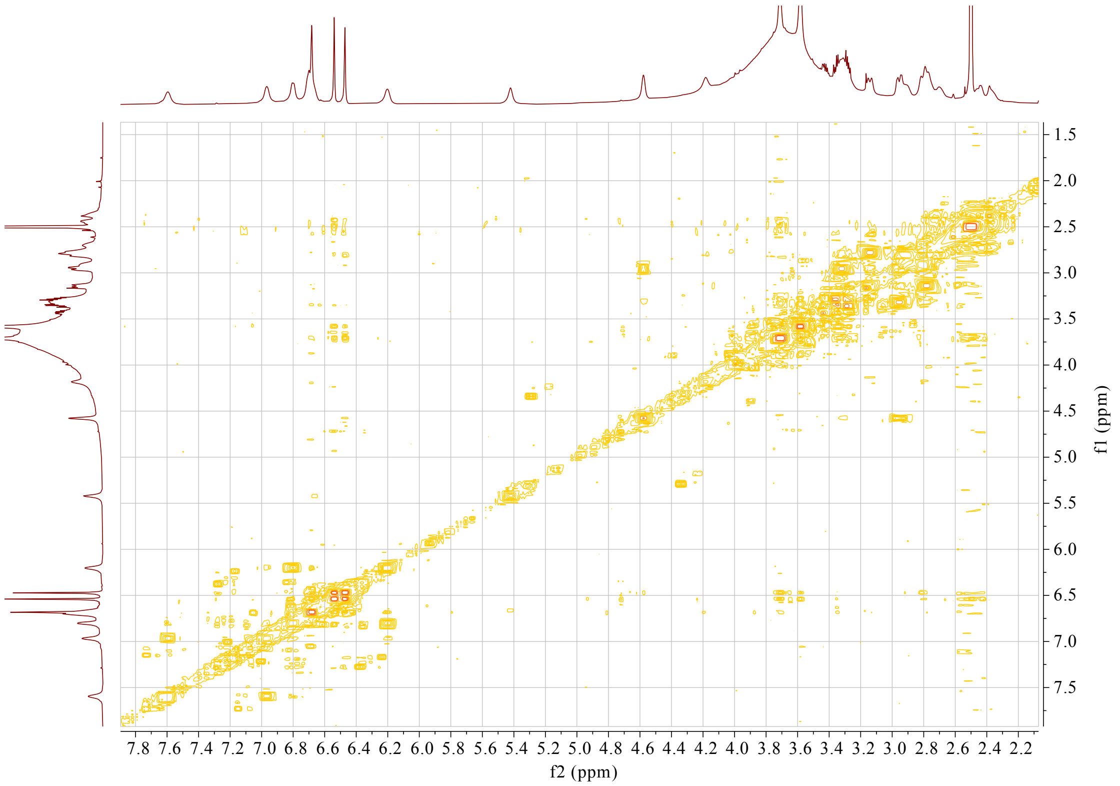


**Supplementary Fig. 43. $^1$H-$^1$H COSY spectrum (600 MHz) of (*R*,*S*)-16 in DMSO-$d_6$**

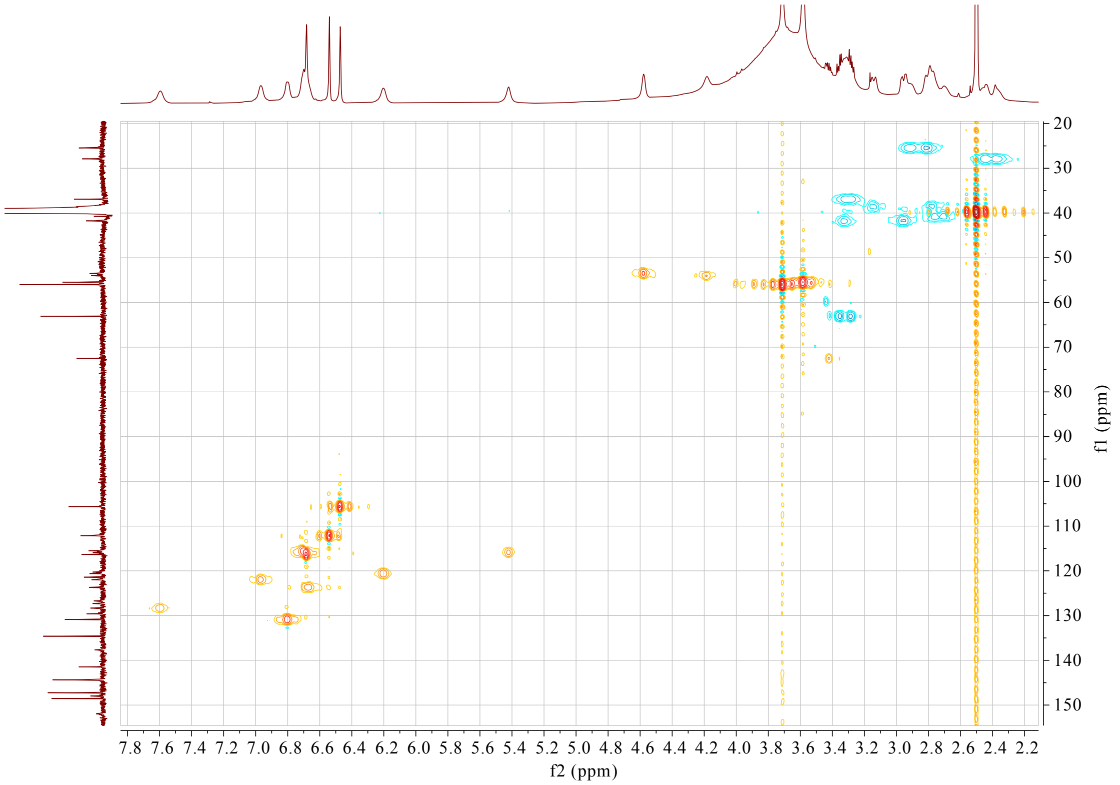


**Supplementary Fig. 44. HSQC spectrum (600 MHz) of (*R*,*S*)-16 in DMSO-$d_6$**

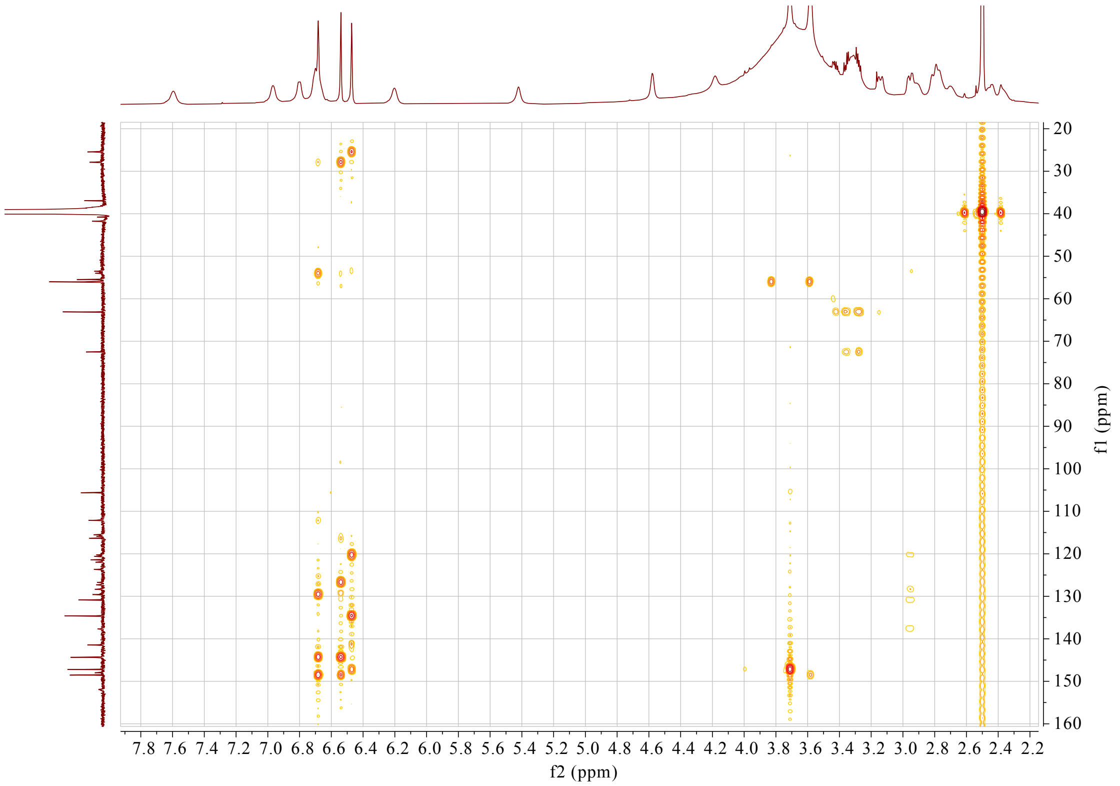


**Supplementary Fig. 45. HMBC spectrum (600 MHz) of (*R*,*S*)-16 in DMSO-$d_6$**

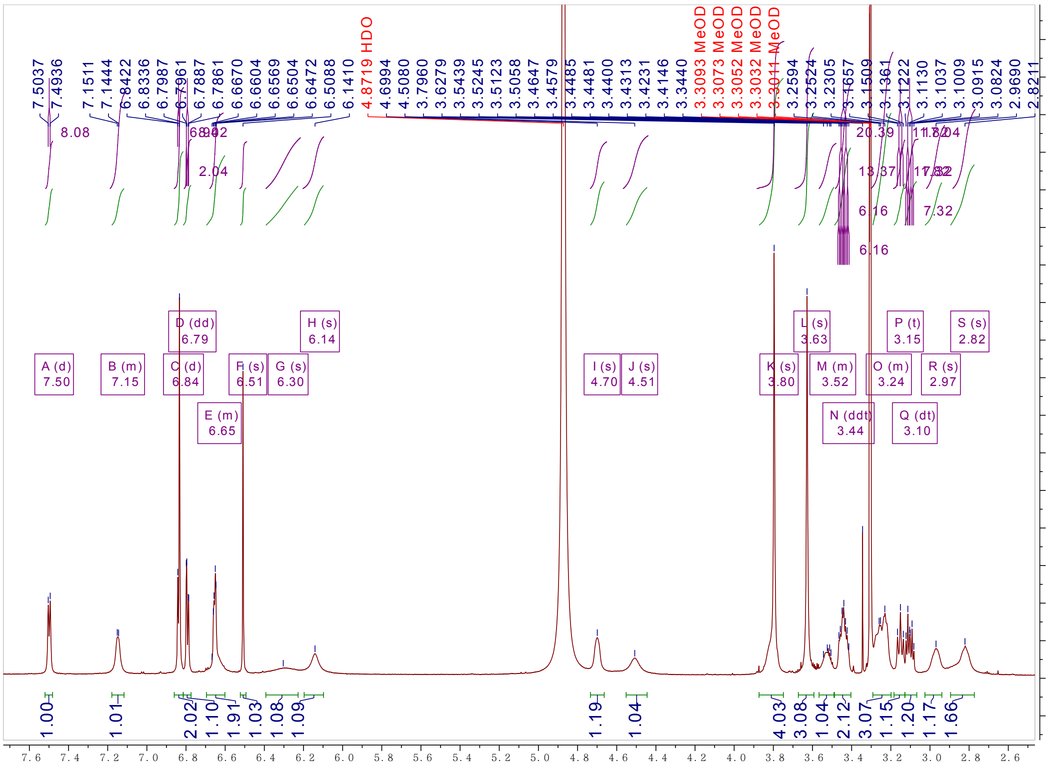


1459 **Supplementary Fig. 46. $^{1}$H NMR spectrum (800 MHz) of compound (*R*,*S*)-17 in DMSO-$d_6$**

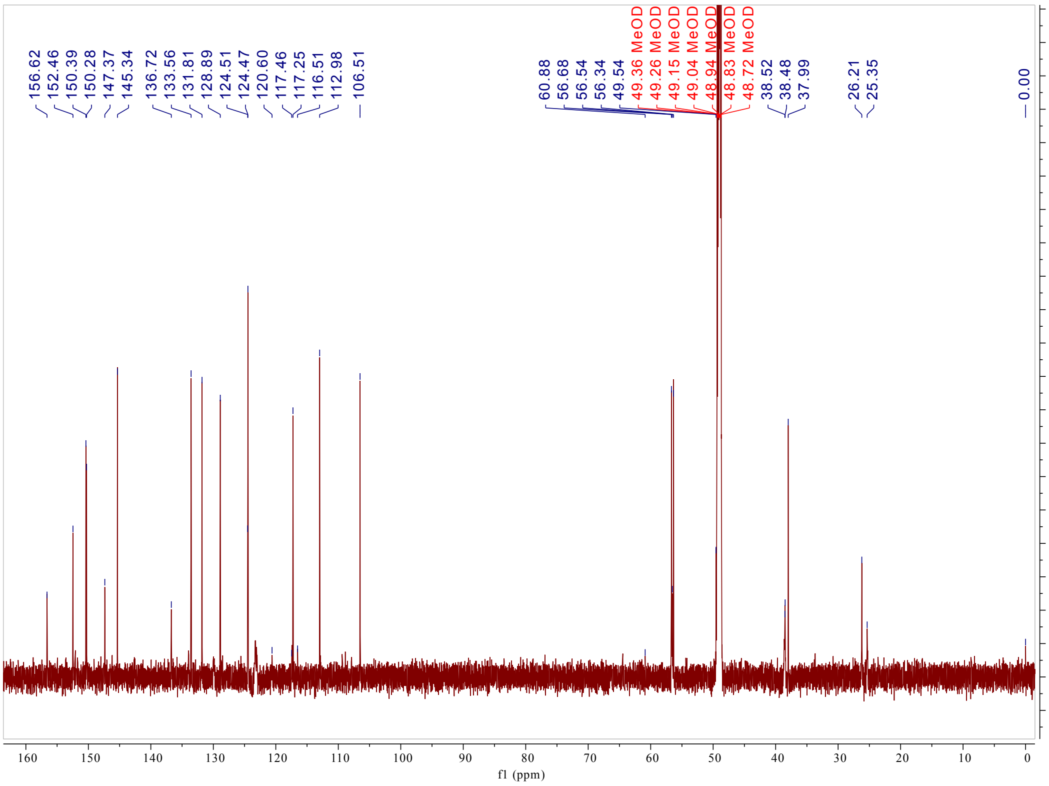


**Supplementary Fig. 47. $^{13}$C NMR spectrum (200 MHz) of compound (*R*,*S*)-17 in DMSO-$d_6$**

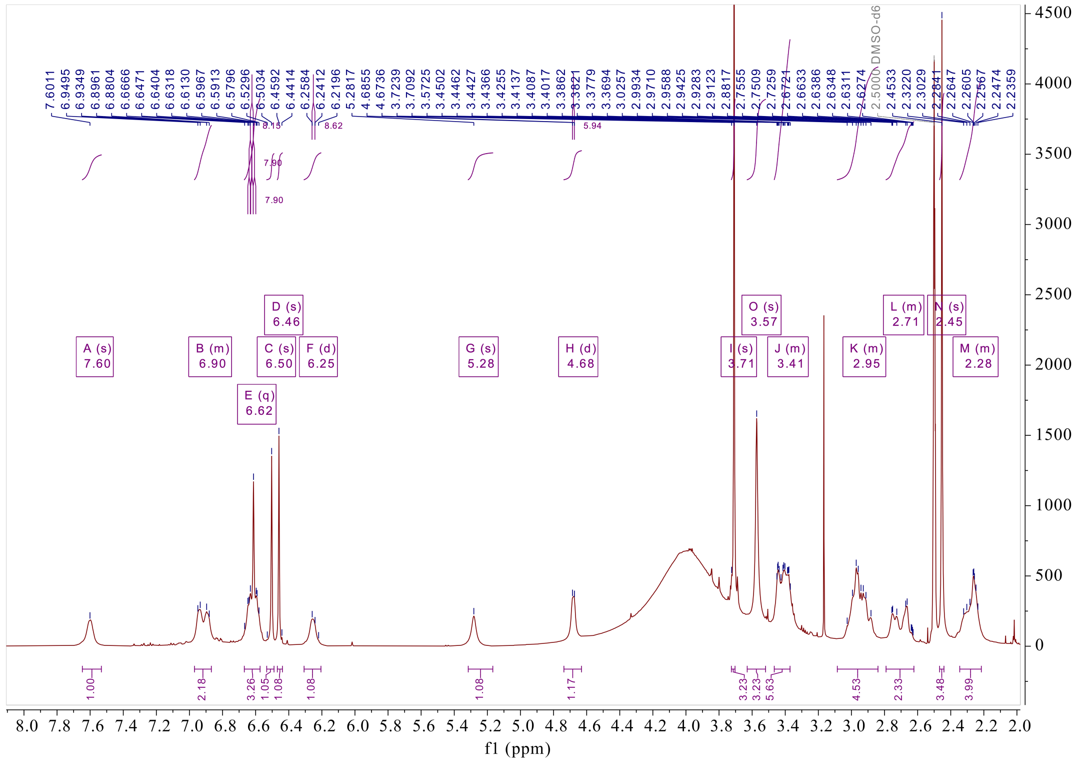


**Supplementary Fig. 48.** $^{1}$**H NMR spectrum (500 MHz) of compound (*R*,*S*)-18 in DMSO-*d*$_6$**

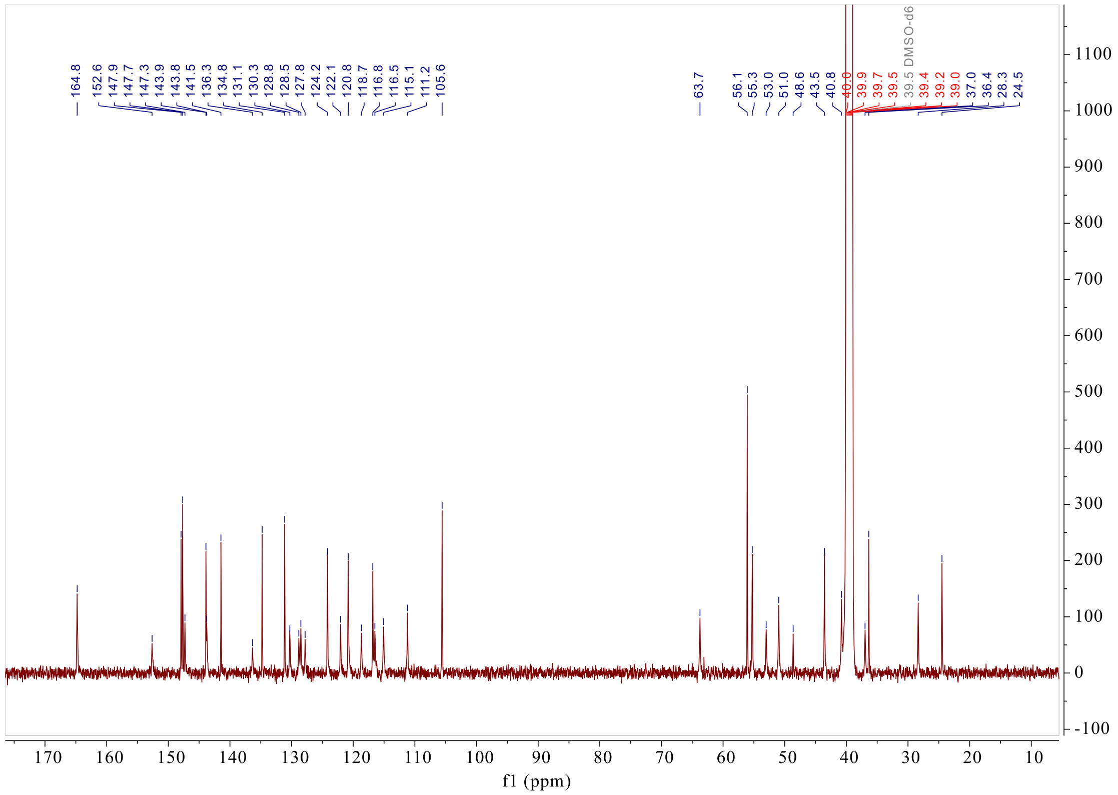


**Supplementary Fig. 49.** $^{13}$**C NMR spectrum (125 MHz) of compound (*R*,*S*)-18 in DMSO-*d*$_6$**

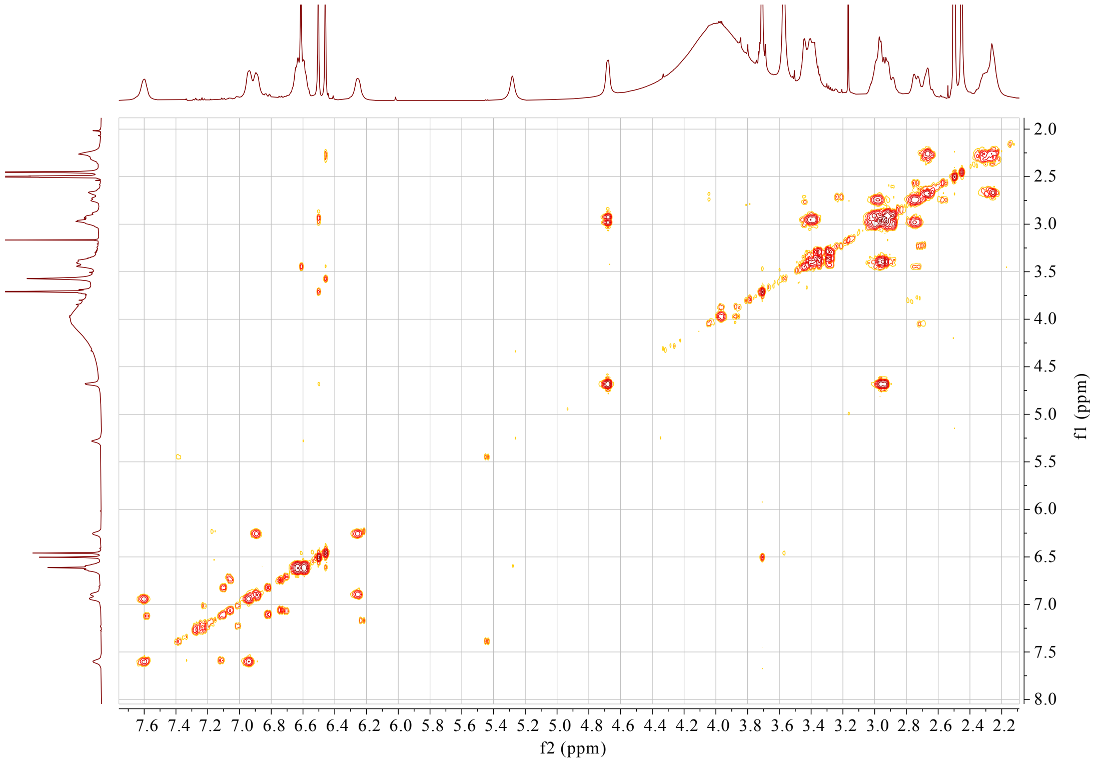


**Supplementary Fig. 50. $^{1}H$-$^{1}H$ COSY spectrum (500 MHz) of compound (*R*,*S*)-18 in DMSO-$d_6$**

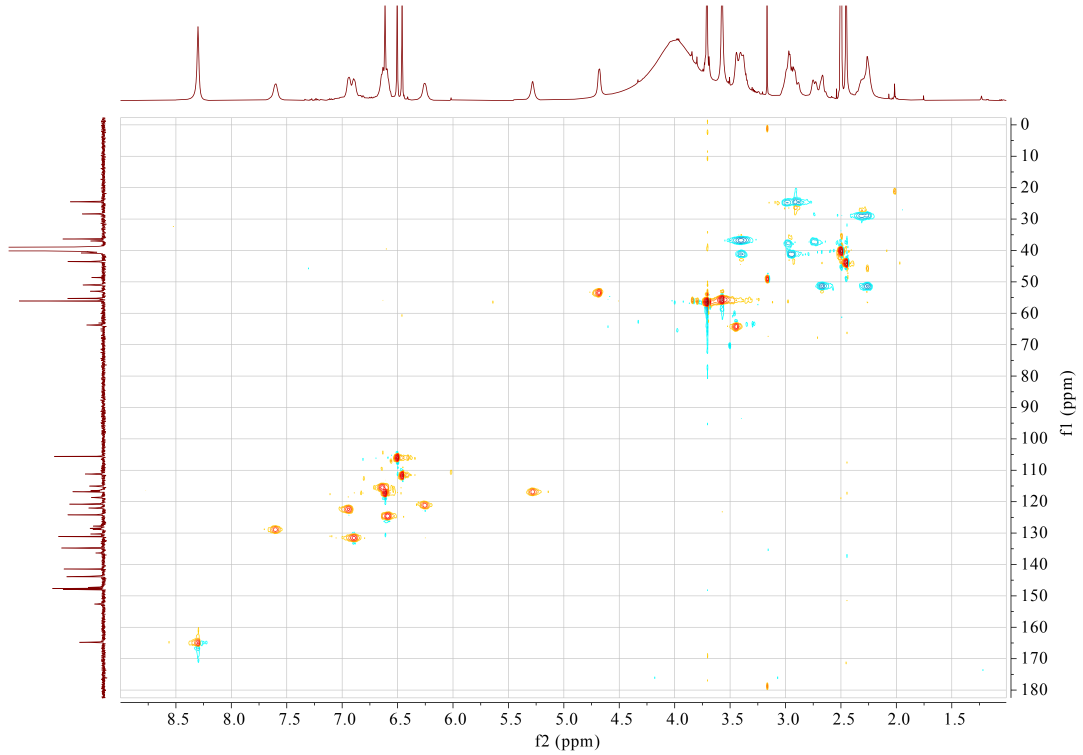


**Supplementary Fig. 51. HSQC spectrum (500 MHz) of compound (*R*,*S*)-18 in DMSO-$d_6$**

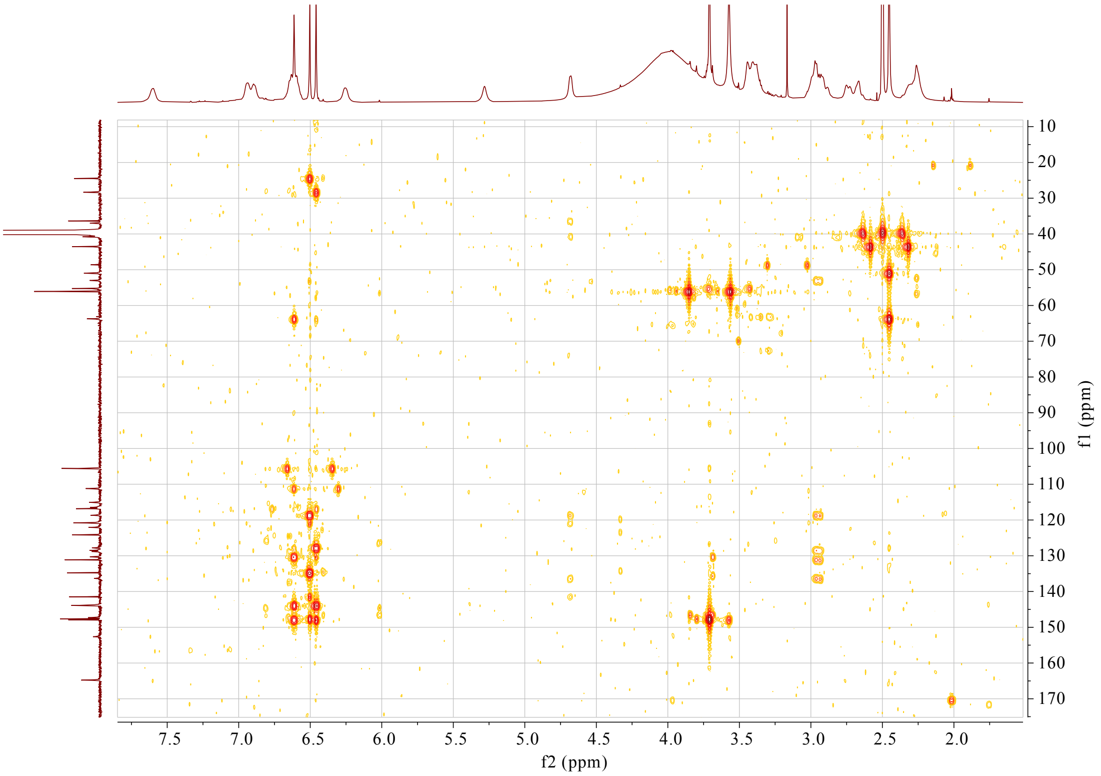


**Supplementary Fig. 52. HMBC spectrum (500 MHz) of compound (*R*,*S*)-18 in DMSO-$d_6$**

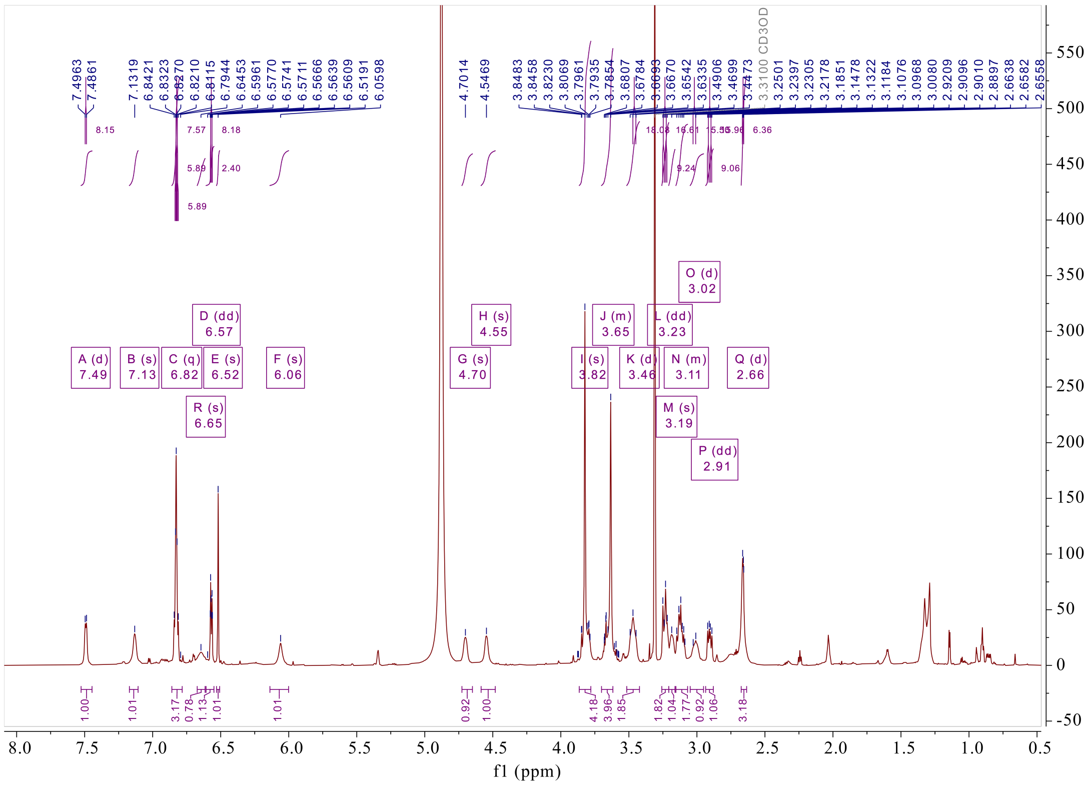


**Supplementary Fig. 53. $^{1}$H NMR spectrum (500 MHz) of compound (*R*,*S*)-19 in $CD_3OD$**

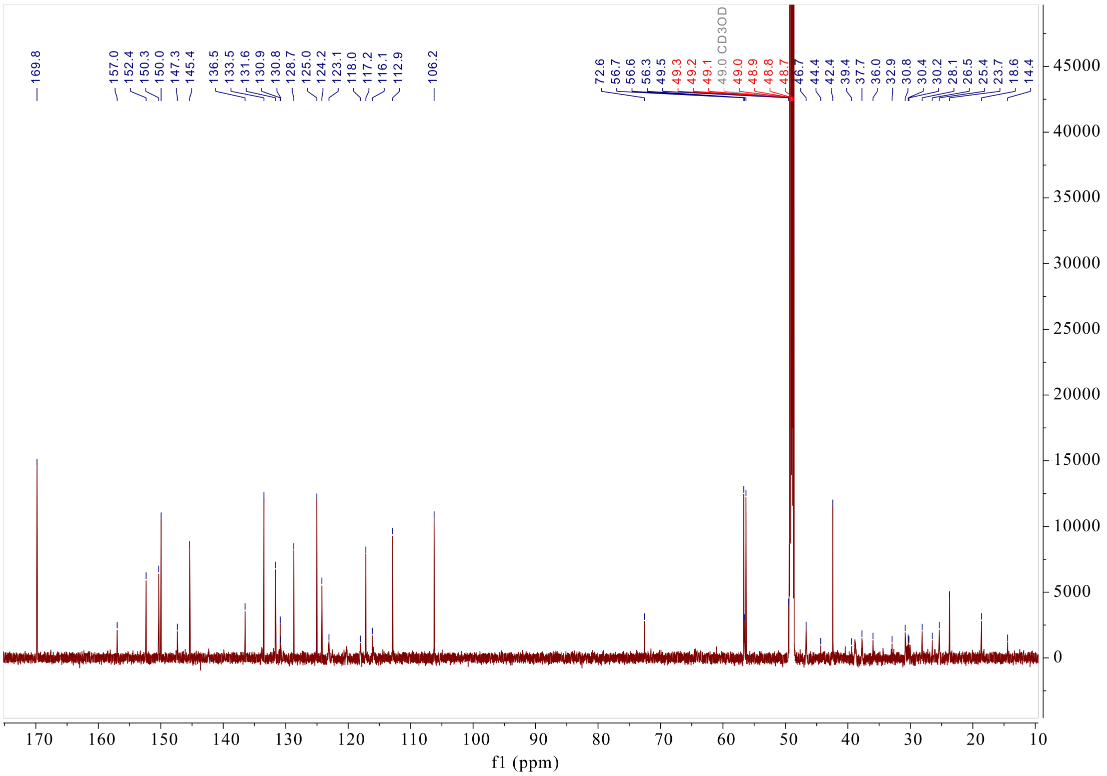


**Supplementary Fig. 54.** $^{13}$**C NMR spectrum (125 MHz) of compound (*R*,*S*)-19 in** $CD_3OD$

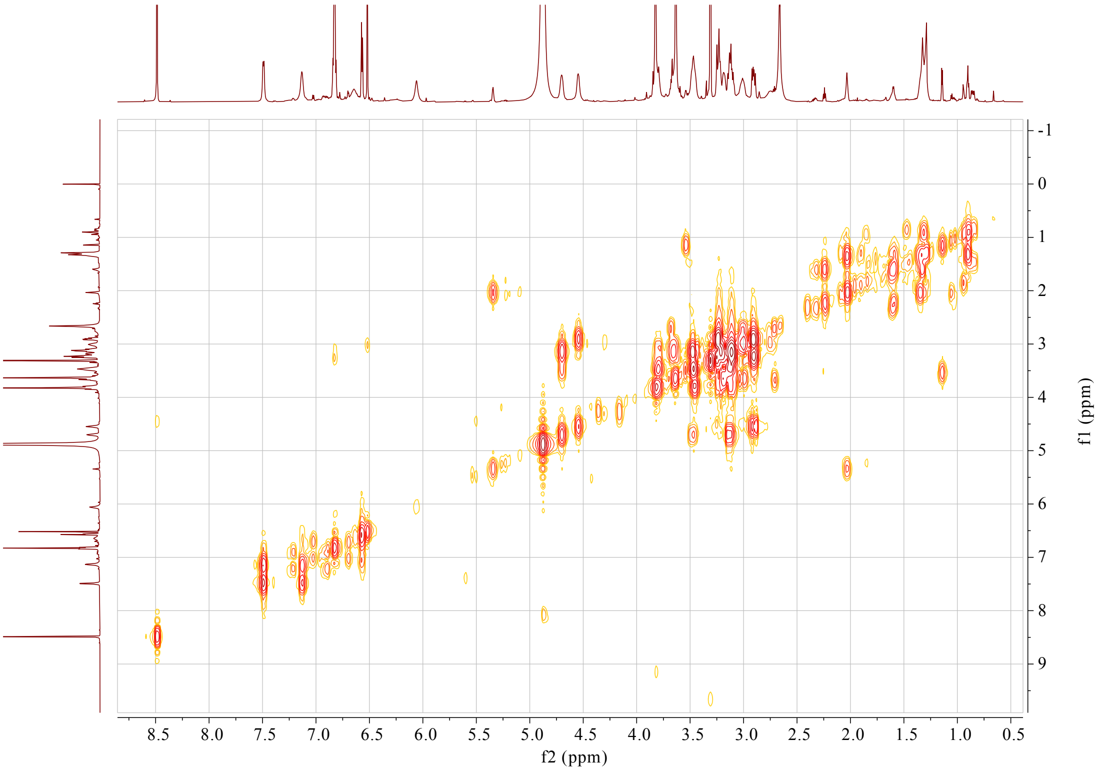


**Supplementary Fig. 55.** $^{1}$**H-**$^{1}$**H COSY spectrum (500 MHz) of compound (*R*,*S*)-19 in** $CD_3OD$

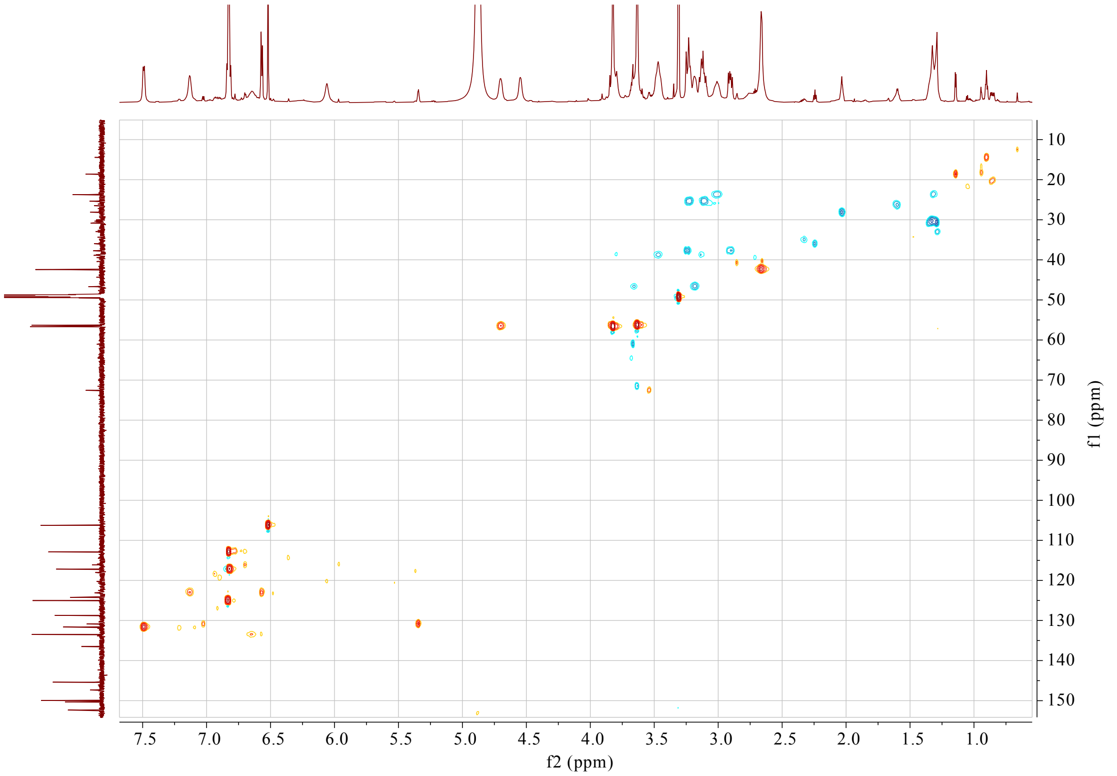


**Supplementary Fig. 56. HSQC spectrum (500 MHz) of compound (*R*,*S*)-19 in $CD_3OD$**

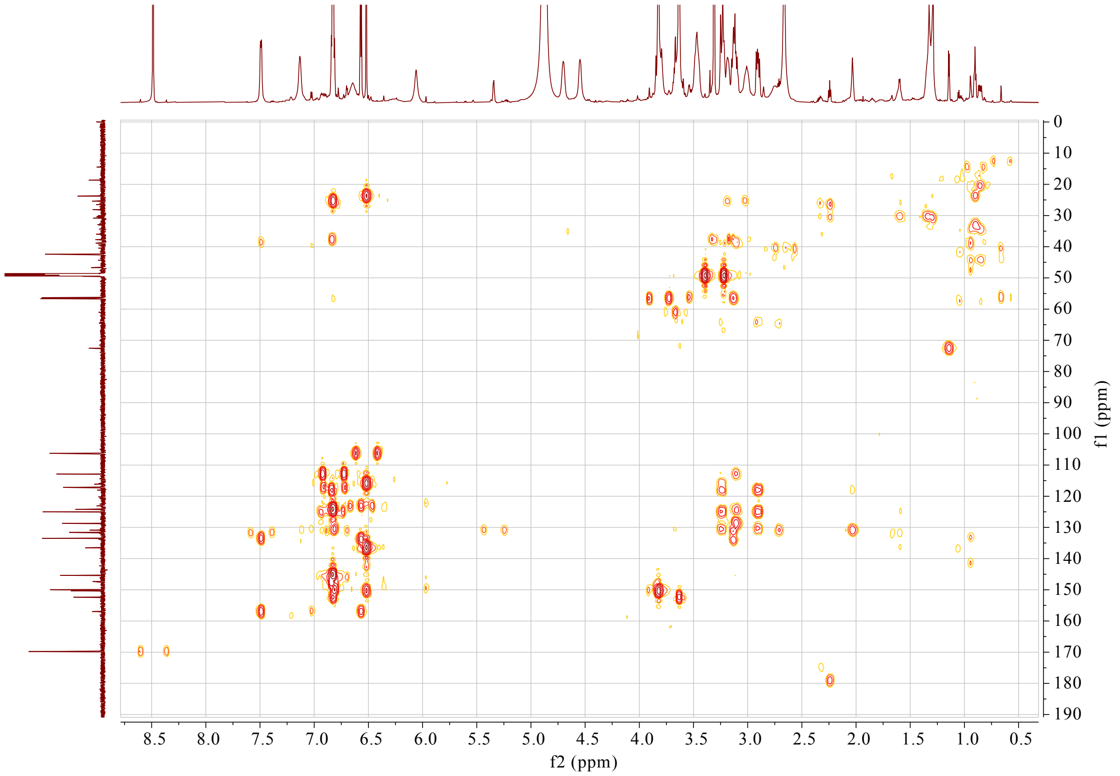


**Supplementary Fig. 57. HMBC spectrum (500 MHz) of compound (*R*,*S*)-19 in $CD_3OD$**

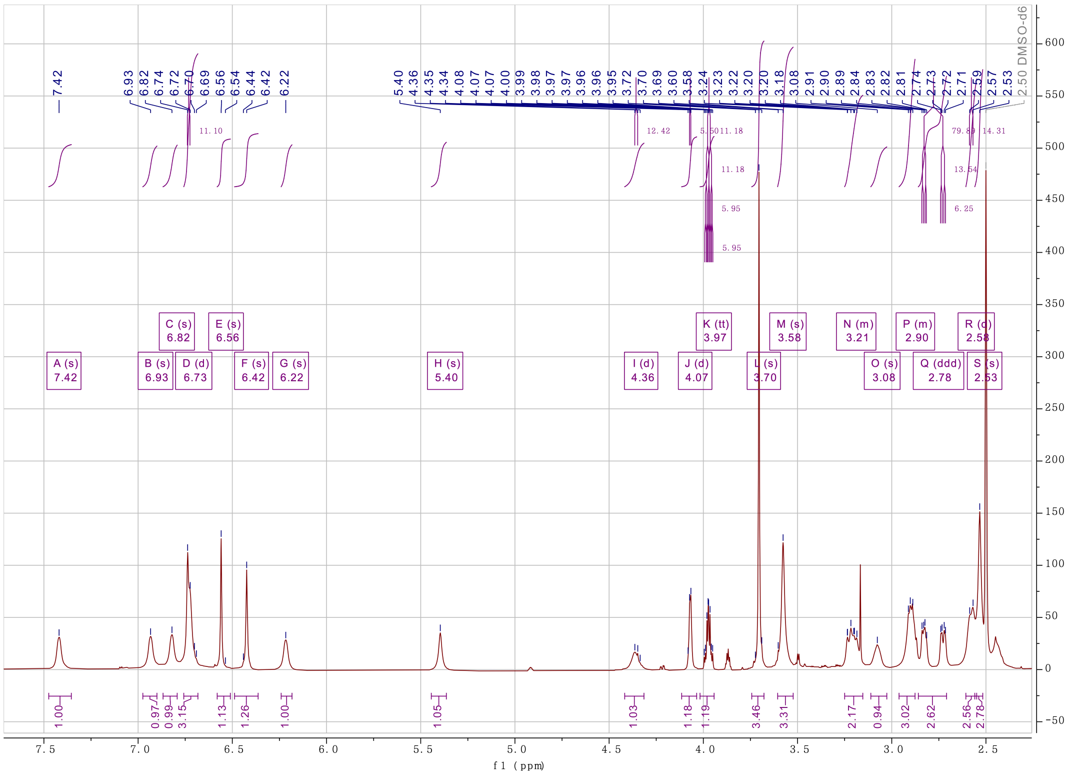


Supplementary Fig. 58. $^{1}$H NMR spectrum (800 MHz) of compound (*R*,*S*)-35 in DMSO-$d_6$

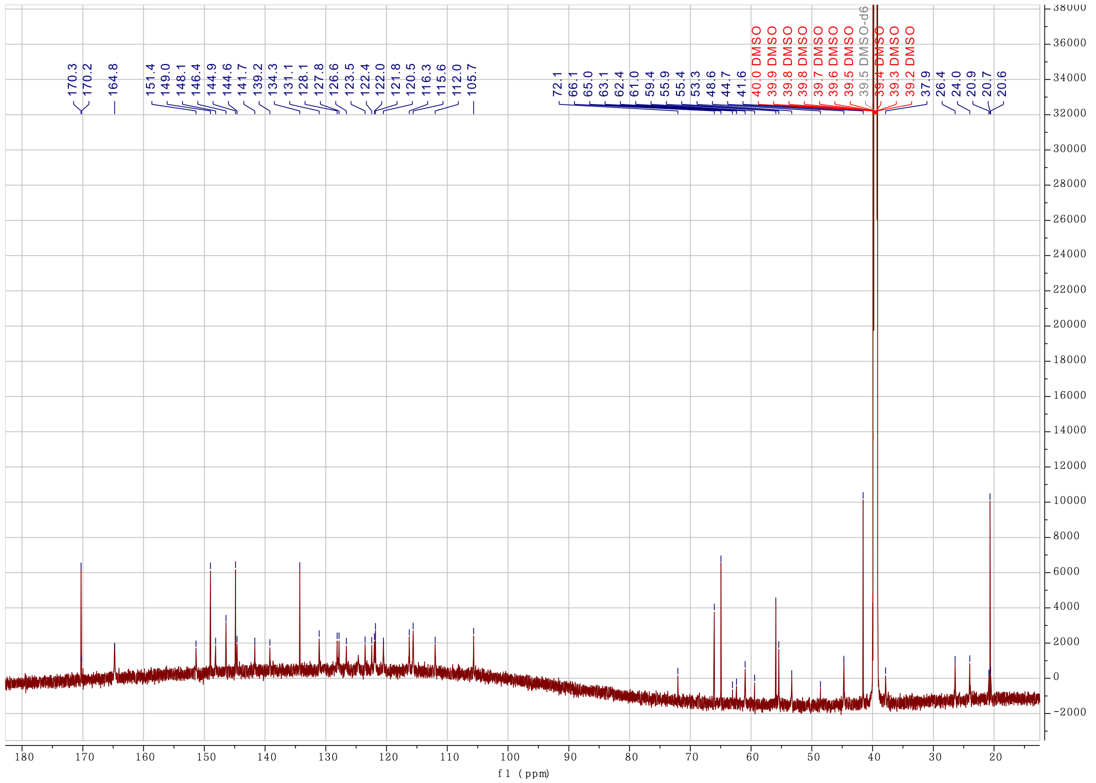


Supplementary Fig. 59. $^{13}$C NMR spectrum (200 MHz) of compound (*R*,*S*)-35 in DMSO-$d_6$

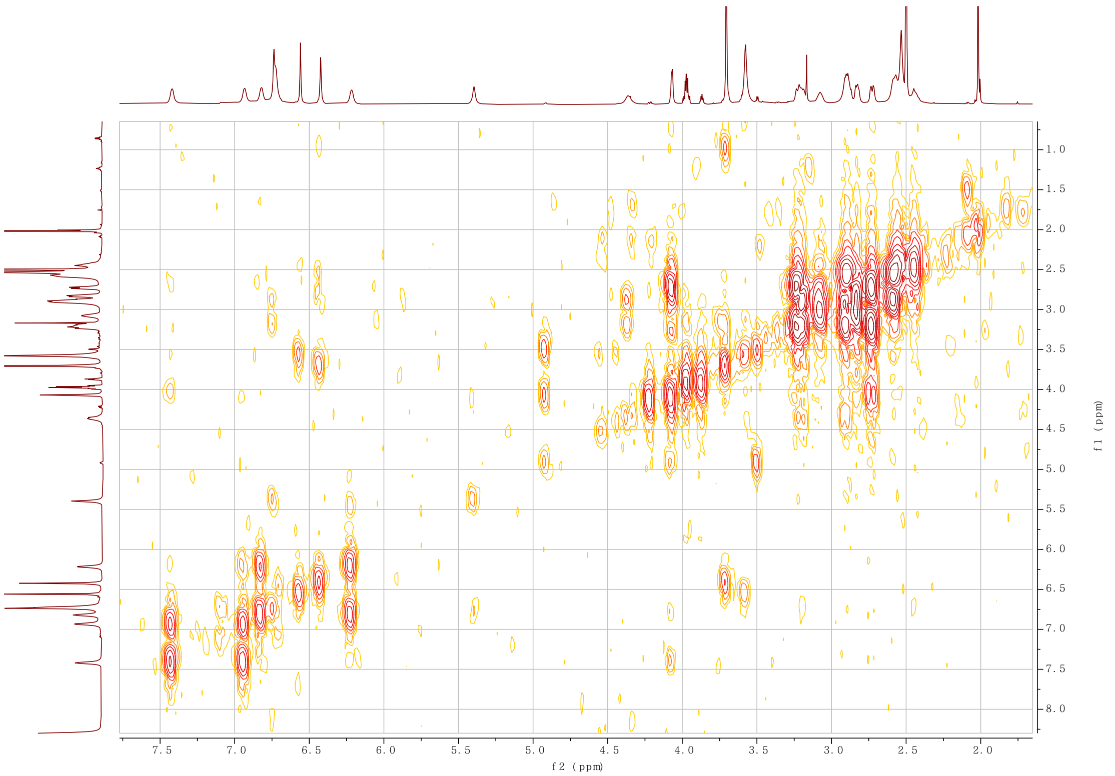


**Supplementary Fig. 60. $^{1}H$-$^{1}H$ COSY spectrum (800 MHz) of compound (*R*,*S*)-35 in DMSO-$d_6$**

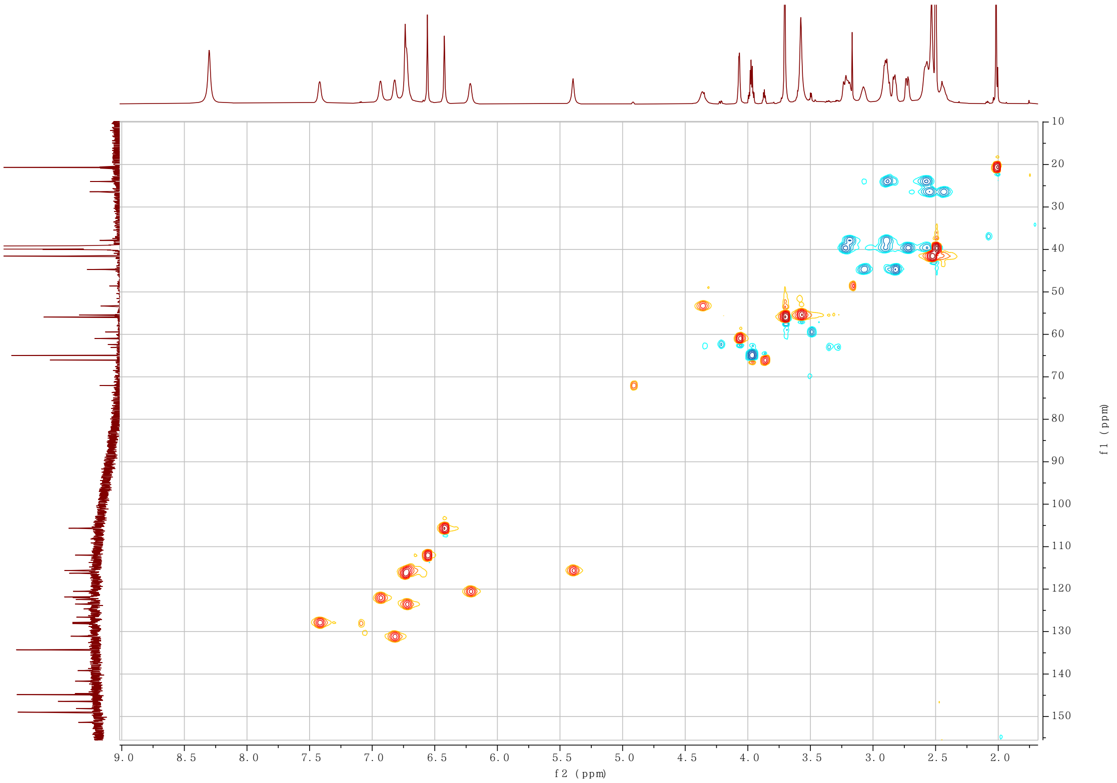


**Supplementary Fig. 61. HSQC spectrum (800 MHz) of compound (*R*,*S*)-35 in DMSO-$d_6$**

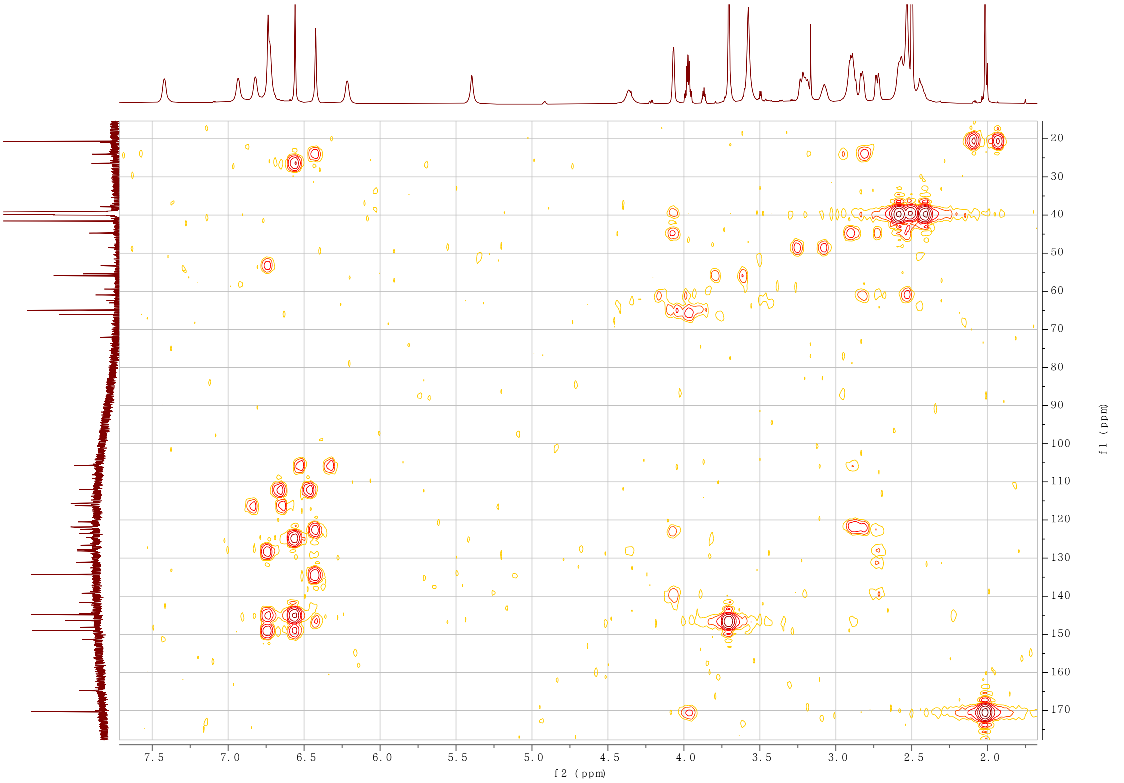


**Supplementary Fig. 62. HMBC spectrum (800 MHz) of compound (*R*,*S*)-35 in DMSO-$d_6$**

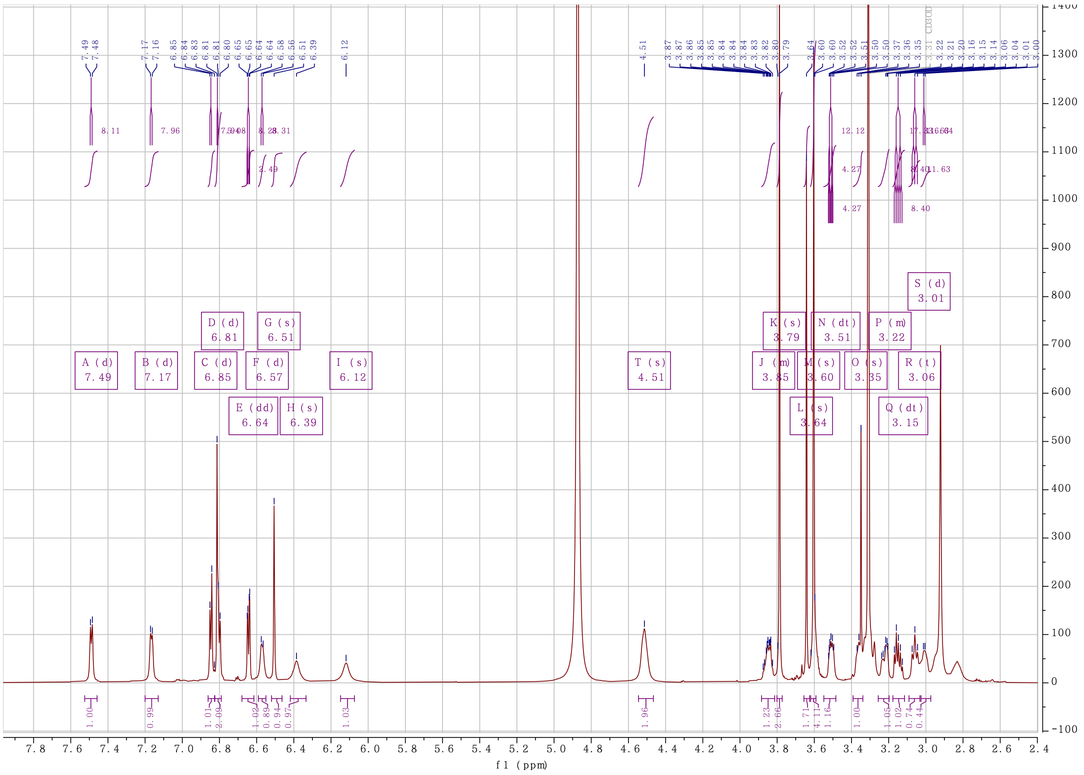


**Supplementary Fig. 63. $^{1}$H NMR spectrum (800 MHz) of compound (*R*,*S*)-36 in $CD_3OD$**

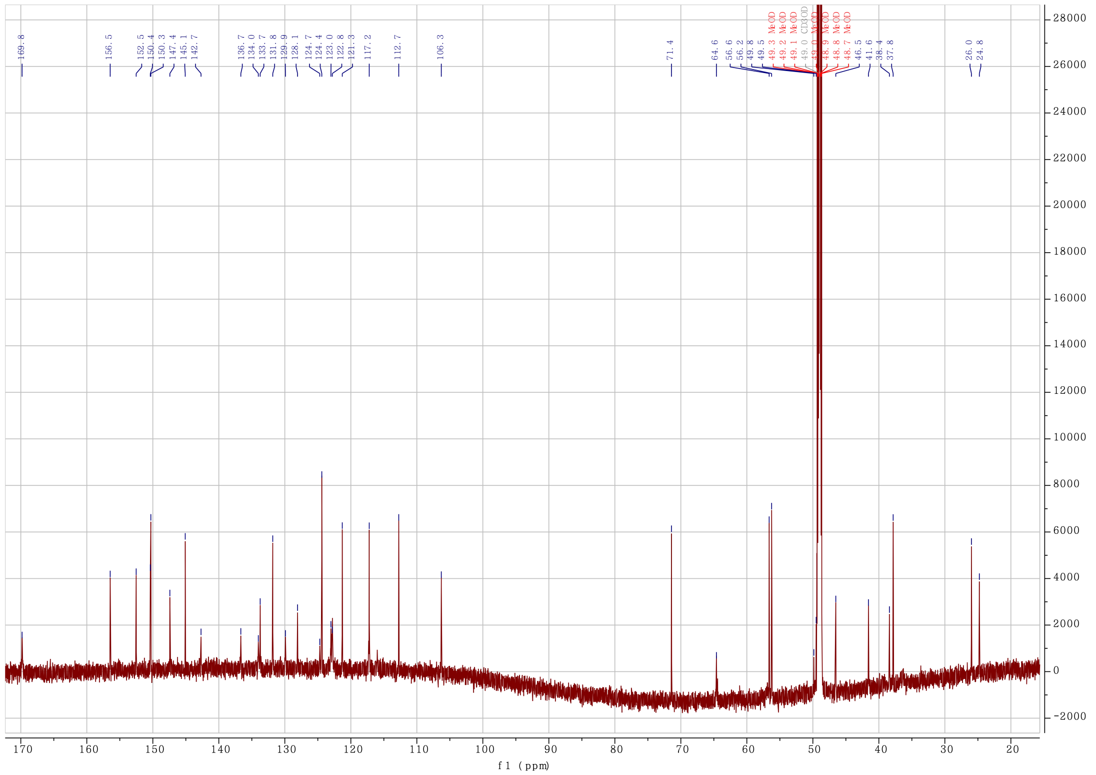


**Supplementary Fig. 64. $^{13}$C NMR spectrum (200 MHz) of compound (*R*,*S*)-36 in $CD_3OD$**

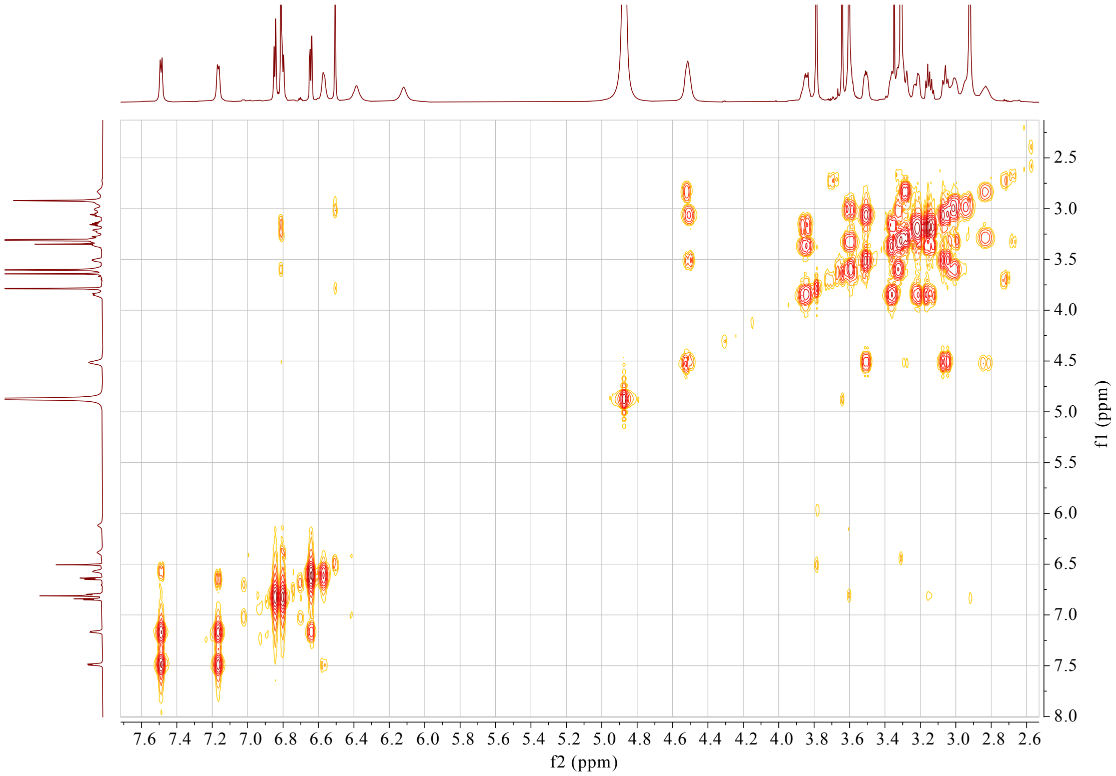


**Supplementary Fig. 65. $^{1}$H-$^{1}$H COSY spectrum (800 MHz) of compound (*R*,*S*)-36 in $CD_3OD$**

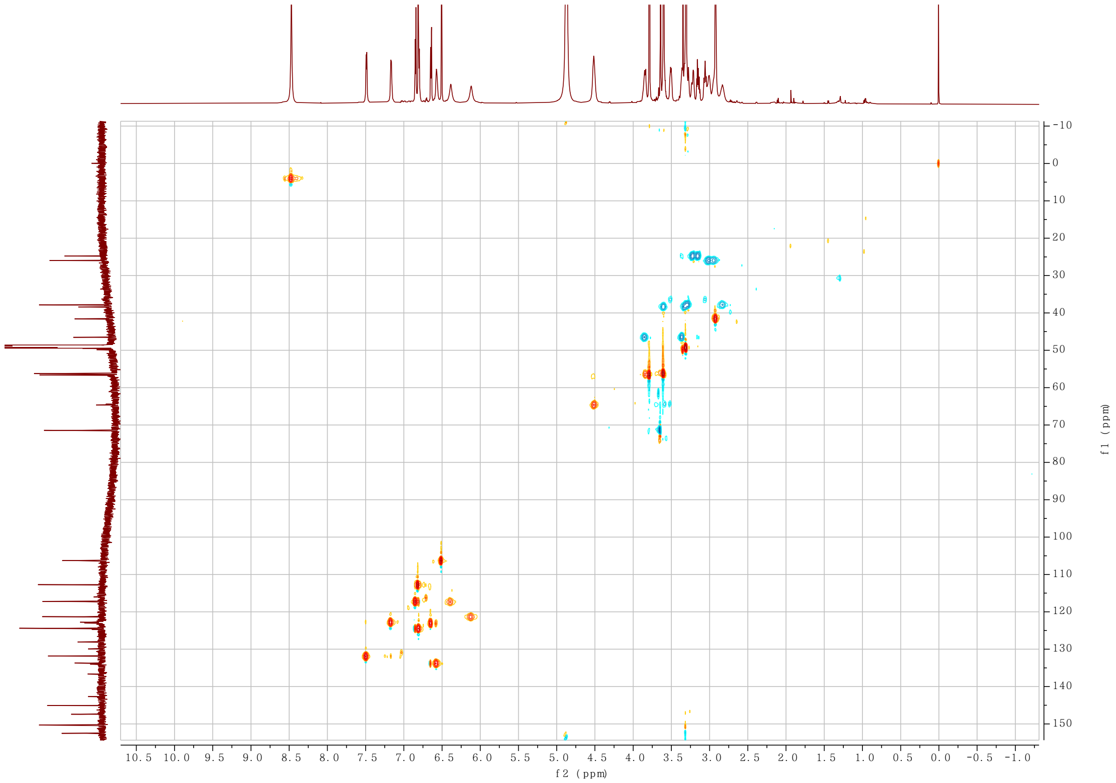


**Supplementary Fig. 66. HSQC spectrum (800 MHz) of compound (*R*,*S*)-36 in $CD_3OD$**

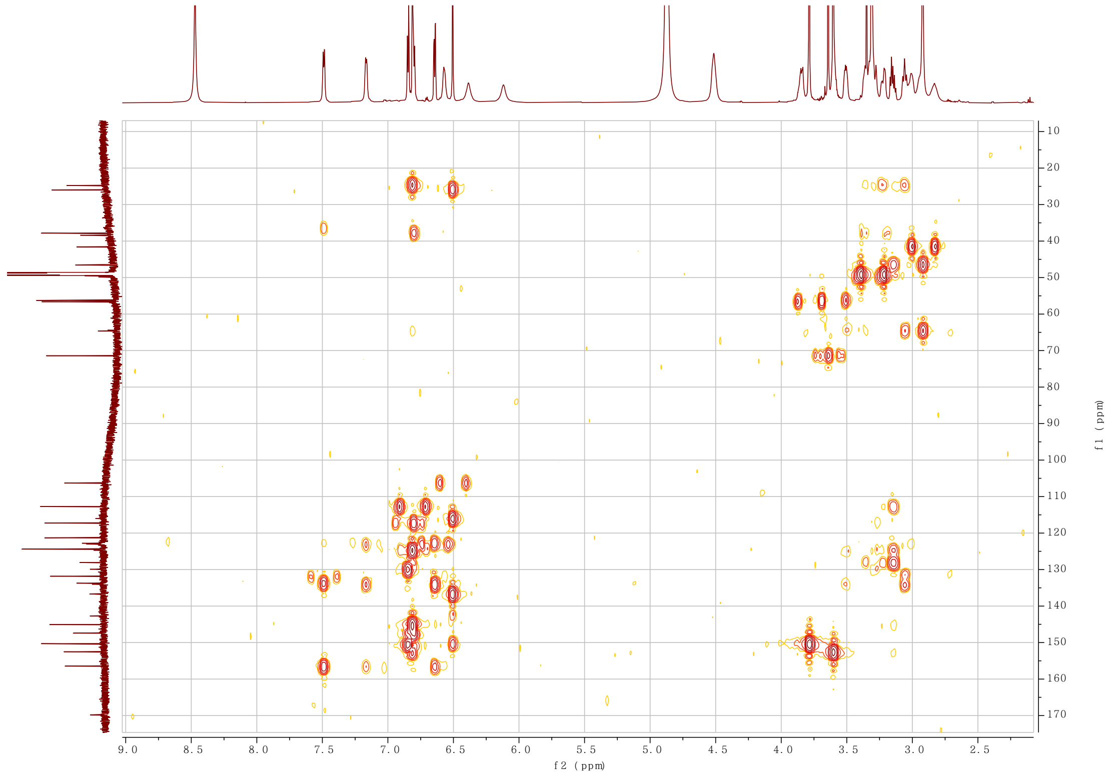


**Supplementary Fig. 67. HMBC spectrum (800 MHz) of compound (*R*,*S*)-36 in $CD_3OD$**
